# Covalent Protein Painting Reveals Structural Changes in the Proteome in Alzheimer Disease

**DOI:** 10.1101/2020.01.31.929117

**Authors:** Casimir Bamberger, Sandra Pankow, Salvador Martínez-Bartolomé, Michelle Ma, Jolene Diedrich, Robert A. Rissman, John R. Yates

## Abstract

The 3D structures of aberrant protein folds have been visualized in exquisite detail, yet no method has been able to quantitatively measure protein misfolding across a proteome. Here, we present Covalent Protein Painting (CPP), a mass spectrometry-based structural proteomics approach to quantify the accessibility of lysine ε-amines for chemical modification at the surface of natively folded proteins. We used CPP to survey 2,645 lysine residues in the proteome of HEK293T cells *in vivo* and found that mild heat shock increased rather than decreased lysine accessibility for chemical modification. CPP was able to differentiate patients with Alzheimer disease (AD) or Lewy body disease (LBD) or both from controls based on relative accessibility of lysine residues K147, K137, and K28 in Tubulin-β, Succinate dehydrogenase, and amyloid-β peptide, respectively. The alterations of Tubulin-β and Succinate dehydrogenase hint to broader perturbations of the proteome in AD beyond amyloid-β and hyper-phosphorylated tau.

## Introduction

AD is a neurodegenerative disorder marked by progressive loss of cognition and other important mental functions. While the cause for AD remains unclear, age is the strongest risk factor for its onset (https://www.alz.org/alzheimers-dementia/what-is-alzheimers/causes-and-risk-factors). A breakdown of the blood-brain barrier and continuous neuronal cell death contributes to cognitive and behavioral decline in AD patients ^1^. The deposition of neurofibrillary tangles and senile plaques precede neuronal cell death and are disease defining hallmarks of AD ^2^. Tangles consist of macromolecular aggregates of tau protein whereas plaques mainly contain aggregated amyloid-β peptide. However, neuronal cell death is not linked to amyloid-β fibril formation in tauopathies or in dementias with Lewy bodies, which yield large aggregates of tau protein and α-synuclein, respectively. While the onset and progression of these neurodegenerative diseases differ, they are all characterized by an accumulation of misfolded proteins. Cells normally recognize proteins with incorrect folds and attempt to re-fold or to pass them on to proteasomal degradation. Chaperone proteins like heat shock proteins play a key role in recognizing misfolded proteins and in refolding client proteins. Chaperones are part of the proteostasis network which encompasses a highly diverse group of proteins that keep the proteome in homeostasis ^3^. If the proteostasis network fails to recognize and remove misfolded proteins, conformationally altered proteins accumulate and can cause cell death. In age correlated neurodegenerative diseases additional changes might impinge on protein conformation homeostasis and it is tempting to propose an insufficiency in removing misfolded proteins with increasing organismal age as a molecular explanation for the onset of neurodegeneration. However, the large number of protein conformations and interactions present in cells makes it experimentally challenging to trace and pinpoint where, when, and why specific proteins misfold and persist in a misfolded state.

In an initial step to monitor protein conformer homeostasis we attempted to measure the degree of protein misfolding *in vivo*. We therefore developed Covalent Protein Painting (CPP), a structural proteomics approach to quantify changes in protein fold or altered protein-protein interaction for any protein in a proteome. CPP directly determines the relative surface accessibility of amino acid side chains by measuring the molar fraction of a chemical functionality that is accessible for chemical modification on the surface of proteins. Here, we targeted the ε-amine of lysine which is a primary amine. Instead of assessing changes in 3D-structure with *in vitro* labeling techniques like “rates of oxidation” (SPROX) ^4,5^, we adopted a standard formaldehyde-based tissue fixation protocol ^6^ to dimethylate all lysine ε-amines that are sufficiently solvent exposed to be chemically modified. Dimethylation leaves a smaller chemical footprint than other chemically more complex reagents used for *in vitro* labeling of lysine residues in highly purified protein complexes such as succimidylanhydride ^7^, diethylpyrocarbonate ^8^, or “Tandem Mass Tags” (TMT) ^9^.

In CPP, solvent exposed primary amines are chemically dimethylated with very high yields within seconds because the addition of each methyl moiety is a two-step reaction that is only rate limited by the initial formation of the hydroxymethylamine ^10^. Subsequently, the labeling reaction is quenched, labeling reagents are removed, and proteins are denatured and proteolytically digested with an endoprotease that is insensitive to lysine. The remaining, previously non-accessible lysine residues which become accessible as a result of proteolysis are labeled in a second labeling step with a set of isotopically different dimethyl moieties. When measured with mass spectrometry, this label allows CPP to directly determine the fraction of protein molecules that was accessible for chemical modification at a specific lysine site based on the relative intensities of the isotope labeled peptides that include that site. The covalent attachment of the label and its *in vivo* applicability sets CPP apart from other approaches for determining protein structure, such as Protein Painting which non-covalently “coats” the protein’s surface with small molecules in order to limit tryptic cleavage ^11^ or limited proteolysis which takes advantage of a differential availability of amino acid sequences for non-specific proteolytic digestion ^12^. Like other methods, CPP enables an unbiased discovery of structural changes caused by misfolding and altered protein-protein interactions in cells and tissues.

Using CPP we show that heat shock of HEK293T cells preferentially increased surface accessibility of lysine sites for chemical modification and that it significantly altered surface accessibility at 14 of 2,645 different lysine sites. Finally, we used CPP to differentiate between patients with neurodegenerative diseases and controls based on an altered lysine accessibility in Tubulin-β, Succinate dehydrogenase, and amyloid-β peptide in postmortem collected brain tissue samples.

## Results

### 3D proteome with CPP

We used ^13^CH_3_ isotope-defined formaldehyde and sodium cyanoborohydride to dimethylate solvent exposed lysine ε-amines in the proteome of HEK293T cells *in vivo* (Figure 1A and Extended data figure 1, Materials and Methods). Proteins were denatured and digested with the lysine-insensitive endoprotease Chymotrypsin. After digestion newly exposed primary amines in peptides were dimethylated with CDH_2_ formaldehyde and sodium cyanoborodeuteride. Following reversed-phase chromatography of peptides, mass spectra of peptide fragment ions were acquired in highest resolution (*R* 120.000) on an Orbitrap Fusion mass spectrometer in order to differentiate and quantify ^13^CH_3_ from CDH_2_-labeled peptides ^13^. Despite the reduced scan speed of the mass spectrometer at its highest resolution settings, CPP surveyed 385 lysine residues in 246 different proteins with 2,297 individual measurements from a total of six replicate experiments which included exchange of isotope and alternative combinations between the first versus second labeling step (first: second label, CDH_2_: ^13^CH_3_, ^13^CH_3_: CDH_2_, ^13^CHD_2_: CD_3_, CD_3_: ^13^CHD_2_). Each measure of relative abundance of the first to the second isobaric label at a lysine site yielded a ratio *R* per lysine site ^13^. *R* values were converted into percentiles of relative accessibility (*% accessibility* = 100 * *R* / (1 + *R*)) which reflects the proportion of protein or proteoform molecules in which a specific lysine site was accessible for chemical modification.

**Figure 1.**
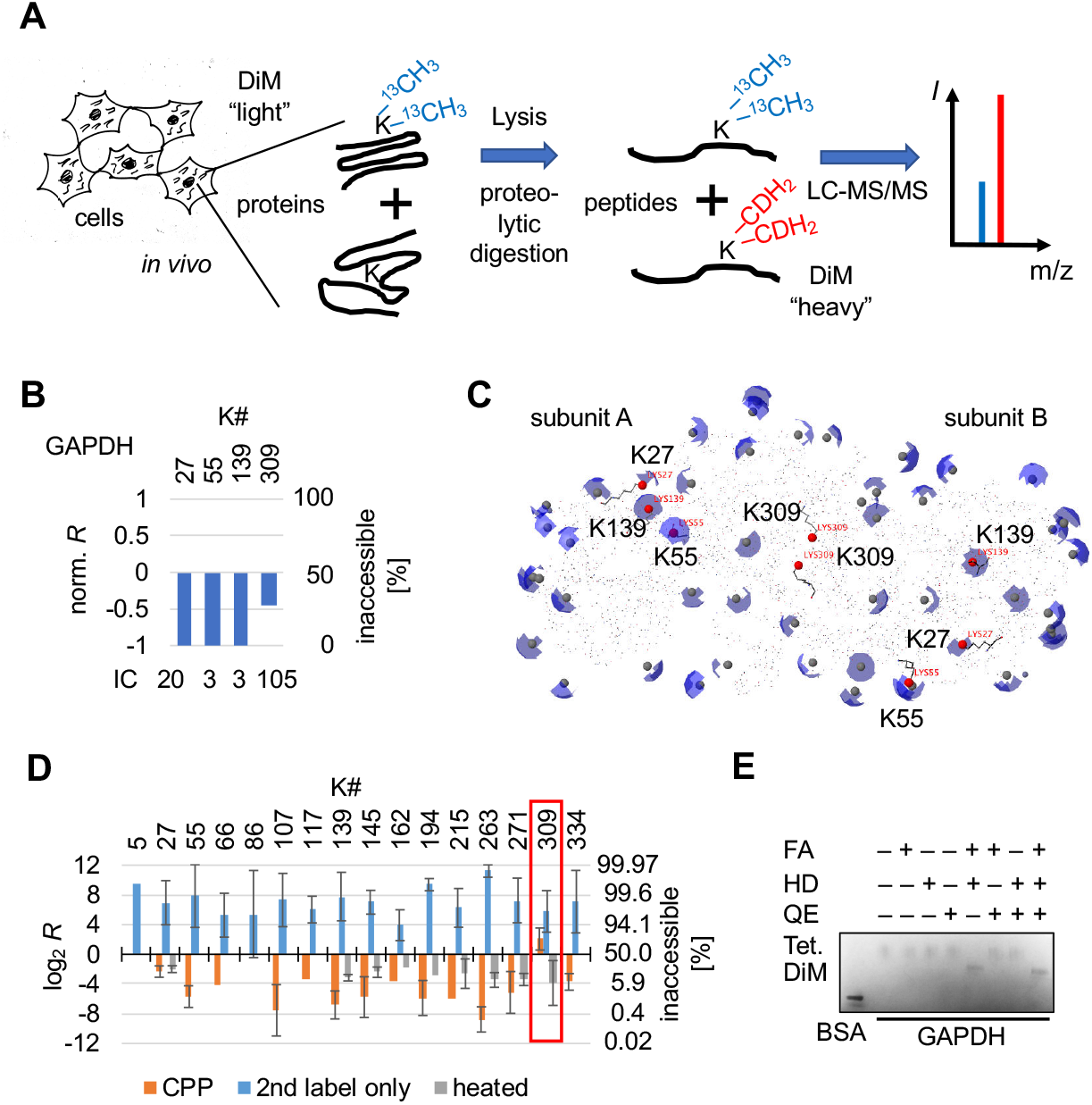
Covalent Protein Painting (CPP) determines whether the ε-amino group of lysine is accessible for chemical modification. **(A)** The schematic displays the workflow of CPP. Reductive alkylation labels lysine residues in proteins with isotope-defined “light” dimethyl moieties in cells *in vivo*. Following digestion into peptides with a lysine-insensitive protease (Chymotrypsin), newly solvent exposed lysine residues are modified with isotope-defined “heavy” dimethyl moieties. Bottom up mass spectrometry is used to analyze the ratio of light to heavy isotope labeled peptide molecules per lysine site. **(B)** Lysine residue K309 of human GAPDH is only partially accessible for chemical dimethylation in HEK293T cells. Proteins in HEK293T cells were covalently modified with CPP using isobaric isotopologue methyl moieties with ^13^CH_3_ for light and CDH_2_ for heavy, and the relative surface accessibility determined as described in (A). Numbers above the bars indicate the position of the lysine residue in GAPDH. The y-axis is the ratio of light to heavy fragment ion counts normalized to the total number of ion counts shown below each bar. A ratio of *R* = 1:1 (log_2_(1) = 0) indicates that the lysine site was accessible for chemical modification in 50 %, *R* > 0 in > 50%, and *R* < 0 in < 50% of protein molecules. Ion counts (IC) denotes the sum of fragment ion peaks. **(C)** One GAPDH dimer of the homo-tetramer (PDB: 4wnc) is displayed. Partial spheres (blue) highlight solvent accessible surface area (SASA) of each individual lysine ε-amine (grey sphere). Lysine residues that were assayed with CPP are highlighted in red in (B). GAPDH#K309 resides within the contact surface of two GAPDH monomers in the GPADH dimer. **(D)** The bar graph shows accessibility of different GAPDH lysine sites for chemical dimethylation in highly purified, native GAPDH tetramers (orange), heat denatured GAPDH (grey), and when the initial labeling step was omitted. A red box highlights CPP results obtained for GAPDH#K309. **(E)** Blue-native gel® electrophoresis of GAPDH indicates stability of the homo-tetramer following chemical dimethylation. GAPDH was pre-incubated with labeling reagents formaldehyde (FA), sodium cyanoborohydride (HD), and the quencher ammonium bicarbonate (QE). Bovine serum albumin (BSA, 66 kD) was included as molecular size indicator. Tetrameric GAPDH protein complexes migrated distinctively faster following CPP. Error bars are standard deviation (σ). Abbreviations: DiM, dimethyl moieties; Tet., homo-tetramers.

Consistent with lysine being the most solvent accessible amino acid in proteins, initial dimethylation labeled 337 out of 385 lysine sites (87.5 %) in > 95 % of protein or proteoform molecules. The remaining 47 lysine sites were either completely inaccessible (13 sites) or accessible in ≤ 95 % of protein molecules (34 sites, Extended data table 1). Several different lysine sites of the metabolic enzyme Glyceraldehyde 3-phosphate dehydrogenase GAPDH were quantified in the dataset and lysine sites GAPDH#K27, GAPDH#K55, and GAPDH#K139 were measured as completely accessible in all GAPDH molecules. In contrast, lysine residue GAPDH#K309 was accessible for chemical modification in < 75 % of GAPDH molecules (Figure 1B). Crystal structures of GAPDH show that GAPDH#K309 is participates in the protein-protein interface of two homo-dimers within the GAPDH homo-tetramer ^14^ (Figure 1C, Extended data figure 2).

Additional *in vitro* experiments showed that GAPDH#K309 was accessible in < 20 % (and conversely, inaccessible in > 80 %) of recombinantly expressed, highly purified human GAPDH tetramers whereas an additional 13 lysine sites were solvent exposed in, on average, 97 % of GAPDH molecules (σ = ±1.8 of log_2_R, Figure 1D and Extended data Figure 3). The remaining 12 of 26 total lysine sites in GAPDH were either not detected or peptides harbored more than one lysine residue upon endo-proteolytic digestion with Chymotrypsin which precluded a site-specific quantitation based on chromatographic elution profiles ^15^.

Next, we tested whether CPP detected protein unfolding and misfolding. Purified human GAPDH was heat denatured (95 °C, 5 min) and subjected to CPP. All lysine residues, including GAPDH#K309, were now accessible in at least > 80 %, and on average in 87 % (σ = ±0.7, log_2_R) of GAPDH molecules (Figure 1D, grey bars). Heat denaturation overall lowered lysine accessibility from 97 % of molecules in native GAPDH to 87 % suggesting that random protein aggregation following heat denaturation rendered lysine residues inaccessible in 10% of protein molecules. As an additional control, we omitted the first labeling step, endoproteolytically digested non-modified GAPDH, and dimethylated all lysine residues (Figure 1D, blue bars). As expected, in this control lysine sites were accessible for labeling on average in 99.3 % (σ = ±1.9, log_2_*R*) of GAPDH molecules as expected. The residual 0.7% reflected most likely random chemical noise picked up during mass spectrometric data acquisition and quantification of elution profiles.

Native gel electrophoresis showed that chemical dimethylation did not affect the tertiary structure of GAPDH (Figure 1E). Dimethylated GAPDH homo-tetramers (147 kD) migrated as a sharp signal slightly below non-modified GAPDH homo-tetramers but well above bovine serum albumin (BSA, 66.5 kD). The molecular weight of BSA is close to the calculated molecular weight of GAPDH dimers that were not observed. The signal intensity did not diminish, suggesting that GAPDH tetramers did not disassemble upon dimethylation. A comparison of the results to the solvent accessible surface area (SASA) of lysine ε-amines in crystal structures of GAPDH indicated that ε-amines required a SASA of > 1 Å^2^ in order to be chemically modified with CPP. Because the CPP results were congruent with the actual fold and tertiary structure of GAPDH and based on the results of additional experiments (Supplementary Information), we conclude that CPP measures the proportion of protein molecules in which a specific lysine residue was accessible for chemical modification.

A rise in temperature from 37 °C to 42 °C for 15 min is a physical stress that leads to protein unfolding or misfolding in eukaryotic cells and elicits a coordinated cellular response of the proteostasis network to limit proteome-wide damage ^16^. We applied CPP to HEK293T cells *in vivo* to find out whether heat preferentially misfolds a specific subset of proteins or whether it leads to wide-spread random unfolding of proteins in the proteome. We used multidimensional protein identification technology (MudPIT) ^17^ on an Orbitrap Velos mass spectrometer to survey the proteome in three biological replicates of HEK293T control cells following heat shock (42 °C, 15 min). The experiment yielded 16,081 different peptide measurements covering a total of > 7,000 different lysine sites of which 2,645 were quantified at least twice per condition in 979 different protein groups and proteoforms (Extended data table 2).

2,484 of 2,645 lysine residues were accessible for chemical modification in > 33 % of protein molecules in control HEK293T cells (Extended data figure 4 and Extended data network 1). Individual lysine sites were accessible for labeling in 94.4 % of protein molecules on average and accessibility normally distributed from 99.7 % to 50 % (2σ-interval of Gaussian fit, R^2^ = 0.9958, Figure 2A). The remaining 161 lysine residues that were accessible for chemical modification in ≤ 33 % of protein molecules clustered in a distinct second peak in the frequency distribution plot (Figure 2A Inset). A positive control in CPP represents the exogenously added endoprotease Chymotrypsin that was not present during initial labeling. Following proteolytic digestion, endoproteolytic peptides of Chymotrypsin are labeled in the second labeling step only, and thus none of its peptides can be quantified as accessible for chemical modification. Lysine residue K54 of the endoprotease Chymotrypsin (CRBT1#K54) was measured as “accessible” in 0.16 % of molecules most likely due to random chemical noise in mass spectra.

**Figure 2.**
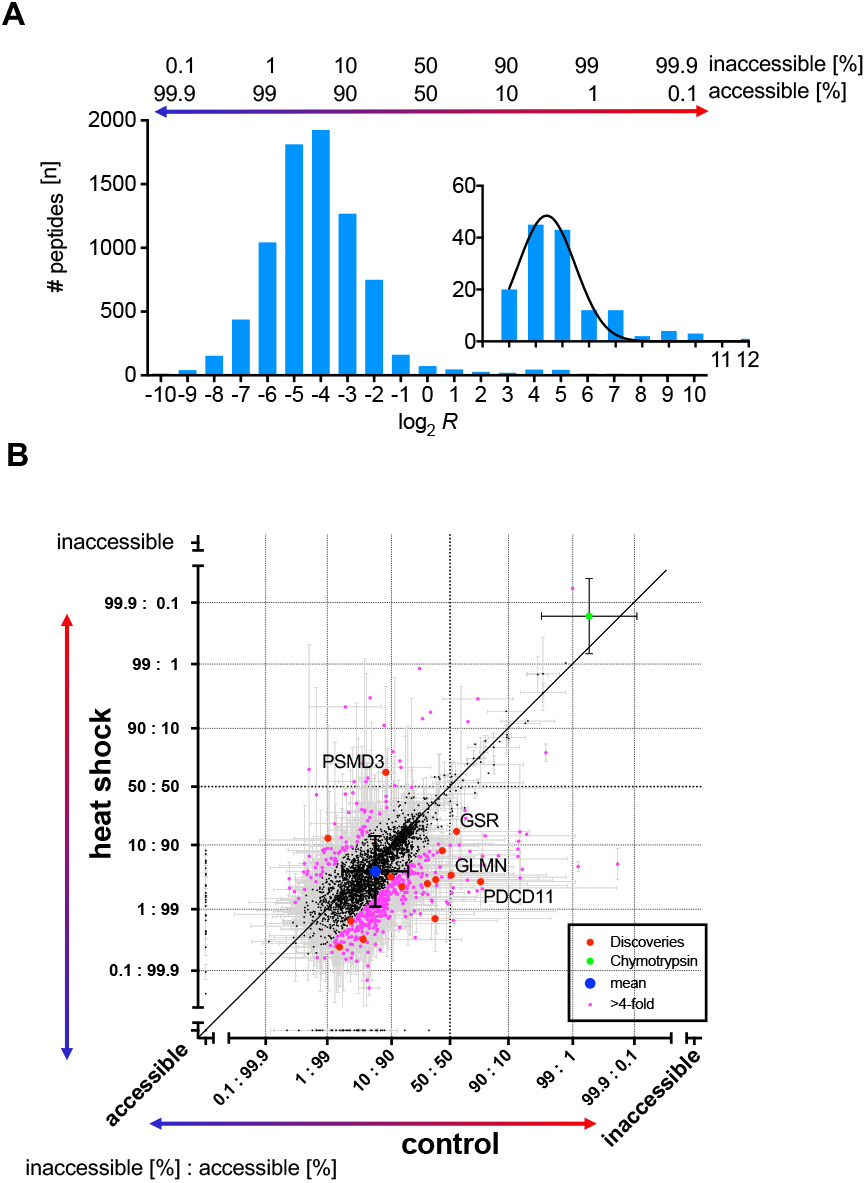
CPP quantified protein unfolding in HEK293T cells upon mild heat shock *in vivo*. (**A**) The frequency plot shows the distribution of the relative proportion of protein molecules in which a lysine site was accessible for chemical modification with CPP in the proteome of HEK293T cells. log_2_*R* values were binned by integer and the frequency distribution of lysine sites inaccessible for chemical modification in a majority of protein molecules are highlighted in the inset. The black line is a Gaussian fit. The relative number of protein molecules [%] in which lysine sites were accessible to chemical modification is indicated on top of the bar graph. (**B**) The scatterplot compares the relative number of protein molecules in which a lysine residue was accessible for chemical modification in control to heat shock-treated HEK293T cells. The units “accessible and “inaccessible” on the scale bar indicate lysine sites that were measured as either completely accessible or inaccessible in control or heat shock. Pink dots highlight individual lysine residues that differed between heat shock and control by Δ > 2. Dots in red indicate lysine sites that passed the discovery threshold of *q* < 0.01. CRBT1#K54 in Chymotrypsin is highlighted in green and the overall mean shown in blue. The 45° angled line (black) denotes no change between control and heat shock. Error bars are standard deviation (σ).

Heat shock altered relative surface accessibility in 461 of 2,645 lysine sites by > σ (≥ 4-fold, pink dots in the scatter plot in Figure 2B) and these lysine sites were more likely to become more accessible (369 sites) than inaccessible (92 sites) for chemical modification, indicating that heat shock preferentially unfolded proteins or weakened protein-protein interactions. Notably, the fractional change in the number of protein molecules with increased accessibility was < 20 % for the majority of the 369 lysine sites (Extended data figure 5). Thus, CPP revealed that most proteins are likely reversibly unfolded upon heat shock, reflecting increased entropy in the HEK293T proteome at elevated temperatures.

Heat shock not only preferentially increased the relative proportion of protein molecules in which a lysine site was accessible, it also significantly altered accessibility for chemical modification in 14 lysine sites (*q-*value < 0.01, red dots in the scatter plot in Figure 2B). For 4 of the 14 proteins, heat shock flipped the proportion of protein molecules from predominantly accessible (> 50 %) to inaccessible (< 50 %) for chemical modification or vice versa (Extended data table 3). K273 in the 26S proteasome non-ATPase regulatory subunit 3, PSMD3, (PSMD3#K273) was the only lysine site in the dataset which was accessible in the majority of PSMD3 molecules in control (91.7 %) and ended up accessible in only 36.8 % of PSMD3 molecules upon heat shock. All three additional lysine sites turned from predominantly inaccessible to accessible in > 50 % of protein molecules upon heat stress. PDCD11#K1402 in programmed cell death protein 11 (RRP5 homolog NFkB binding protein, NFBP) shifted from accessible in 24.0 % to accessible in 97.3 % of PDCD11 molecules upon heat shock. In mitochondrial Glutathione reductase, GSR, lysine site GSR#K501 changed from accessible in 43.9 % to accessible in 84.3 % of GSR molecules, and GLMN#K507 in Glomulin turned from accessible in 49.9 % to accessible in 96.5 % of GLMN molecules upon heat exposure. In summary, CPP revealed that mild heat shock increases entropy in the proteome based on a rise in the number protein molecules in which lysine was accessible for chemical modification. In a few proteins, heat shock led to alterations in protein conformation or protein-protein interaction that were potentially irreversible.

Prolonged heat shock causes extensive post translational protein modifications (PTM) on lysine, including ubiquitinylation and sumoylation ^18^. Thus, we determined if ubiquitinylation and sumoylation in response to heat shock occurred at lysine sites that were also quantified by CPP. 68 proteins were either ubiquitinylated or sumoylated in control and heat shock-exposed cells, with more PTM occurrences following heat shock. Ubiquitinylated or sumoylated proteins were overall enriched for the Gene Ontology term “protein folding” (*p*-value = 12.9), and several ubiquitinylation sites were identified with > 5 spectral counts (SpC) across all biological replicates, including tubulin-α(TBA1B, 13 SpC), splicing factor (U2AF2, 13 SpC), and heterogeneous nuclear ribonucleoprotein R (HNRNPR, 9 SpC), and apoptosis inhibitor 5 (API5, 5 SpC). Proteins ubiquitinylated only in heat shock-exposed cells included the ubiquitin-like modifier-activating enzyme 1 (UBA1, 13 SpC), transcription intermediary factor 1β (TRIM28, 8 SpC), and heat shock 70 kD protein (HSP70, 5 SpC). Overall, CPP covered 20 proteins and lysine sites out of 68 proteins that were ubiquitinylated or sumoylated. Differences in accessibility for chemical dimethylation were below 2-fold for most of these lysine sites. An exception was lysine site TBA1B#K430 in Tubulin-α1B, which was sumoylated upon heat shock and showed 3.4-fold more TBA1B molecules that were inaccessible for chemical dimethylation. Likewise, TPIS1#K256 in triosephosphate isomerase 1 was sumoylated and displayed 3.8-fold more TPIS1 molecules in which TPIS1#K256 was inaccessible after heat shock. PTM modification does not necessarily decrease the proportion of molecules in which PTM modified lysine site was accessible for chemical modification; HSP74#K84 of heat shock protein 70 kD protein 4 was inaccessible in 3.6-fold less HSP74 molecules despite being ubiquitinylated upon heat shock. As expected, a subset of lysine sites quantified with CPP were also PTM modified. CPP does not quantify changes in PTM because CPP tests only non PTM-modified lysine residues. One exception is naturally occurring lysine dimethylation, which can influence CPP results depending on the choice of isotope-defined reagents used in the design of the experiment (Supplementary Information).

### Differentiating neurodegenerative diseases based on 3D alterations

Next, we tested CPP as a potential conformational diagnostic tool to measure protein misfolding in neurodegenerative diseases. We analyzed prefrontal cortex samples of controls and 10 patients that were diagnosed with AD, LBD or with severe diffuse LBD in addition to AD (AD-dLBD, Extended data table 4). CPP was applied to whole tissue lysate and to the pellet formed following ultracentrifugation (UC) to enrich for protein aggregates ^19^. Overall, the experiment quantified 559, 342, and 303 lysine sites in lysate, pellet, and supernatant, respectively. Only lysine sites in amyloid precursor protein APP, APP#K699, mitochondrial succinate dehydrogenase SDHB, SDHB#K137, and tubulin-β TUBB, TUBB#K174 were significantly altered between controls and AD, AD-dLBD or LBD in the lysate (*q*-value < 0.05, Table 1). TUBB#K174 was overall accessible in 99.2 % to 97.3 % tubulin-βmolecules. In AD, AD-dLBD and LBD samples TUBB#K174 was on average accessible for chemical modification in 0.6 % fewer molecules than in controls. SDHB#K137 was on average accessible for chemical modification in 21.4 % fewer protein molecules in AD, AD-dLBD, and LBD. Inaccessibility for chemical modification increased > 2-fold in patient-derived samples over controls, in which SDHB#K137 was measured in half of all samples.

**Table 1.**
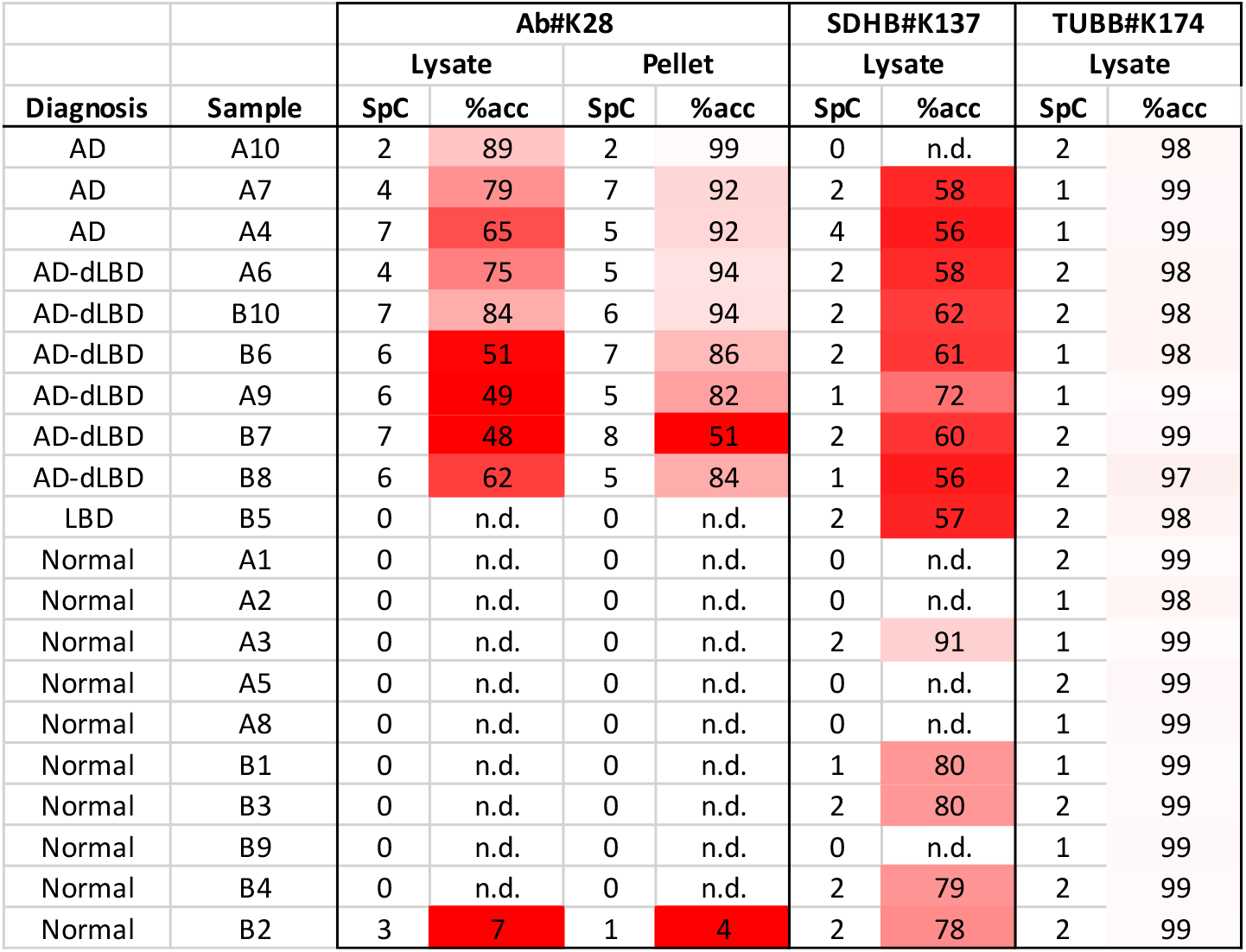
Amyloid-β misfold in AD and LBD patient prefrontal cortex. The table shows the accessibility in percent for Ab#K28 and SDHB#K137, and TUBB#K174 lysine sites in 10 patient and 10 control samples. Note that Ab#K28 was not detected in the supernatant following UC in any of the samples whereas it was present in UC pellet as well as initial lysate of patients diagnosed with AD. Abbreviations: SpC, spectrum count; %acc, percentile of peptide or protein molecules in which the lysine site was accessible for chemical dimethylation.

Lysine site K699 in a chymotryptic peptide of APP was almost exclusively present in AD and AD-dLBD samples. APP is the precursor protein of amyloid-β peptides Aβ_1-40_ and Aβ_1-42_ which are the naturally occurring endoproteolytic cleavage products of APP that form large peptide aggregates or plaques in AD patients ^20^. APP#K699 matches Aβ#K28 in the amyloid-β peptides Aβ_1-40_ and Aβ_1-42_. We infer chymotryptic APP#K699 peptides were derived from naturally accumulated amyloid-β peptides rather than from APP for a number of reasons. First, additional peptides that matched APP only and not amyloid-β peptides Aβ_1-40_ and Aβ_1-42_ were not detected in any of the samples. Second, peptides were present only in lysate and UC pellet but not UC supernatant, suggesting that only aggregated proteins or peptides yielded sufficient amounts of chymotryptic cleaved Aβ peptide for mass spectrometric detection. Third, Aβ peptides were not detected in an LBD only patient sample which is consistent with the observation that Aβ plaques are absent in LBD only diseased patients. Fourth, peptides originating from Aβ were not detected in 9 out of 10 control samples from cognitively unimpaired patients in any of the three different sample preparations (lysate, UC pellet and UC supernatant). The only control sample in which peptides originating from Aβ were detected was B2 where Aβ#K28 was detected and inaccessible for chemical modification in almost all chymotryptic peptide molecules. We further consider sample B2 as outlier because it was derived from an individual who did not show symptoms of neurodegeneration. Aβ#K28 accessibility varied between 48 % and 89 % in the brain tissue lysate AD and AD-dLBD patient samples. Aβ#K28 was accessible between 65 % to 89 % of peptide molecules in patients diagnosed with AD only whereas it was accessible between 48 % to 75 % of peptide molecules in patients diagnosed with AD-dLBD. Thus, in AD-dLBD patients Aβ#K28 was on average accessible in fewer (but not significantly fewer) molecules than in AD only patient samples.

Biochemical purification of protein aggregates impacted CPP results. Aβ#K28 was consistently accessible for chemical modification in more peptide molecules in the UC pellet than in the lysate. The largest fold difference in peptides with inaccessible Aβ#K28 was observed in AD patient sample A10 in which the percentage of peptide molecules with accessible Aβ#K28 increased from 89 % in the lysate to 99 % in the UC pellet. Thus, the proportion of peptide molecules in which Aβ#K28 is inaccessible for chemical modification changed > 10-fold, from % to 1 % in sample A10. For all samples, fold changes measured after UC correlated with the initial accessibility in lysate or remained almost unaltered when the number of peptide molecules with accessible Aβ#K28 was ≥ ~50 % in the lysate: AD-dLBD patient B7 showed Aβ#K28 lysine accessibility for chemical modification in 48 % of peptide molecules in Lysate and 51 % of peptide molecules in UC, and control B2 in 7 % and 4 % of peptide molecules in lysate and UC pellet, respectively. These differences show that UC can increase the proportion of Aβpeptide molecules in which Aβ#K28 was accessible for chemical dimethylation.

## Discussion

Here, we demonstrated the feasibility and versatility of covalent protein painting, CPP, to measure changes in chemical reactivity of lysine sites across a complete proteome in HEK293T cells *in vivo*. Mild heat shock unfolded or disrupted protein-protein interactions in < 20 % of protein molecules. This increase in the number of protein molecules with lysine sites accessible for chemical modification most likely reflects increased entropy in the proteome at higher temperatures. Lysine sites in three proteins, PDCD11, GSR, and GLMN switched from predominantly inaccessible to predominantly accessible (> 90 %) which suggests that this change was non-random, and thus might identify these proteins as molecular thermostats with a non-linear response to heat shock. The molecular pathways associated with these proteins are ribosome assembly (PDCD11), oxidative stress response (GSR), and protein translation (GLMN). PDCD11 supports maturation of ribosomal subunits 40S and 80S ^21^ and processing of 47S rRNA ^22^ which transiently subsides during prolonged heat shock ^23^. GLMN binds to RBX1 and prevents E2 ligase recruitment and therewith Cul1 E3 ligase-mediated ubiquitinylation of substrates ^24^. However, GLMN#K507 does not map to the interaction surface of GLMN with RBX1 ^24^ indicating that heat shock disrupts a protein-protein interaction or protein conformation that is not further characterized.

Utilizing CPP as a conformational diagnostic tool we found that Aβ#K28 in amyloid-β, TUBB#K147 in TUBB, and SDHB#K137 in SDHB were less accessible for chemical modification in patients with neurodegenerative disease than in controls. While the difference for TUBB#K147 was small (0.6 %), it might still be of biological relevance because microtubules directly support neuronal function. SDHB is part of the oxidative respiration chain in mitochondria which is a key metabolic process that fails in aging neurons. A 2-fold increase in molecules with altered accessibility to lysine site SDHB#K137 might reflect a previously unidentified alteration in SDHB protein structure or protein-protein interaction in AD. Previous work showed that the dehydrogenase activity of SDHB is blocked by amyloid-β peptide ^25^.

Misfolding and aggregation of amyloid-β peptide is a key molecular signature of AD that is intensely studied. In brief, a number of non-imaging techniques coupled to mass spectrometry were used to determine the amyloid-β misfold *in vitro* such as hydroxyl radical protein footprinting ^26^ and fast photochemical induction of hydroxyl radicals” (FPOP) ^27^, which oxidizes amino acid moieties on the surface of a limited number of proteins ^28^. Hydrogen-deuterium exchange coupled with mass spectrometry (HDX-MS) ^29^ revealed surface exposed hydrogen atoms of fibrillar amyloid-β with high spatial resolution *in vitro* ^30^ and reductive alkylation of amyloid-β protein fibrils in combination with peptide based-mass spectrometry ^31^ or limited proteolysis followed by mass spectrometry ^32^ also elucidated the structural constraints of lysine residues in *in vitro*-assembled amyloid-β fibrils. The structure of amyloid-β fibrils was first revealed in AD patient brain samples with light and later electron microscopy ^33,34^, and x-ray diffraction showed that amyloid-β fibrils consist of amyloid-β peptides that fold into two anti-parallel β-strands that associate in a short β-sheet secondary structure or “pleated sheet” configuration in fibrils ^35^. Misfolded amyloid-β molecules stack perpendicular to the planar surface of the pleated sheet either directly or in a staggering mode. These protofilamentous oligomers display a tertiary “cross-β“ conformation and continue to grow into amyloid-β fibers. Nuclear magnetic resonance (NMR) ^36^ and cryo-electron microscopy (cryo-EM) ^37^ visualized the position and interactions of individual amino acid side chains in the core of *in vitro*-purified amyloid-β fibrils in a single conformation.

In the most common proposed model for the amyloid-β misfold, lysine site K28 forms an intramolecular salt bridge with aspartate D23 which stabilizes the hairpin loop which connects the two β-strands ^38^. Amyloid-β fibers can also associate with an alternative number of laterally neighboring fibers which then influence the surface accessibility of lysine K28 ^39^. Assuming that Aβ#K28 is inaccessible for chemical modification in amyloid-β fibers, CPP quantified the relative proportion of fibrillar amyloid-β in AD, and a potentially higher proportion of fibrillar amyloid-β in AD-dLBD patient brain samples. Surprisingly, almost all amyloid-β was fibrillar in a control who was asymptomatic for AD. CPP showed that following ultracentrifugation amyloid-β aggregates displayed fewer molecules with inaccessible Aβ#K28 than in the initial lysate, suggesting that purifying fibrils might alter one or several different fibrillar amyloid-β conformers. *In vitro* outgrowth assays of amyloid-β fibers seeded with AD patient-derived brain material recently highlighted differences between clinical AD subtypes and the heterogeneity of amyloid-β conformers that can coexist ^40^. Furthermore, recent cryo-EM data suggested that Aβ#K28 can be solvent accessible in distinct strains of amyloid-β fibrils ^41–43^. In addition, denaturation assays revealed up to three different states of amyloid-β aggregation in Alzheimer disease brain samples with up to 4-fold (Aβ_1-40_) or 20-fold (Aβ_1-42_) more aggregated amyloid-β than soluble amyloid-β^44^.

In conclusion, CPP quantifies the proportion of protein molecules in which a lysine site is accessible for chemical dimethylation in a proteome. With CPP, we determined the contribution of fibrillar amyloid-β with inaccessible Aβ#K28 in AD and AD-dLBD patient brain samples and revealed that SDHB and TUBB might be conformationally altered upon neurodegeneration.

## Acknowledgements

We thank Claire Delahunty for reading the manuscript. We are thankful to Robin Park for support in mass spectrometric data analysis with IP2 (Integrated Proteomics) and Daniel McClatchy for many informal discussions. We thank Ivy Trinh and Jeffery Metcalf from the Rissman laboratory for technical assistance and dissecting brain tissues for this study.

## Author contributions

C.B., S.P. and J.R.Y. designed the research. M.M. performed the GAPDH experiments; S.P. and C.B. performed the HEK293T experiments; C.B. performed the AD experiments and J.D. measured the AD samples on the mass spectrometer. S.M.B. and C.B. conceived and S.M.B. implemented the protein residue-specific quantification and SoPaX in PCQ. R.R. and J.R.Y. provided materials and funding. C.B. wrote the manuscript and prepared the figures with help from all authors. All authors read and approved the manuscript.

## Materials and Methods

### Chemical Dimethylation of GAPDH

A Michael addition reaction was used to dimethylate primary amines. Chemical dimethylation allows the use of different combinations of carbon ^12^C, and hydrogen H and carbon ^13^C and Deuterium D in the isotope labels. CPP takes advantage of two successive independent dimethylation reactions to allow for the incorporation of two different isotope-defined dimethyl groups. The specific isotope combination in the first and the second dimethylation step in each of the experiments are listed in Extended data table 5.

Dimethylation of GAPDH was performed with recombinantly expressed, highly purified human GAPDH (LifeTechnologies) dissolved in 2 mM HEPES (pH 7.4). In this first step, ε-amines of lysine residues were labeled with isotope-defined reagents (H, ^12^C, “light”) on native proteins. Formaldehyde was in 10-fold molar excess over lysine residues present in the reaction mixture. Specifically, 1.7 μl H_2_O, 2.0 μl HEPES buffer pH 7.0 (1 M), 5 μl of GAPDH protein (1 μg/μl), 1.7 μl formaldehyde (2 % v:v, Sigma), and 0.6 μl NaBH_3_CN (160 mM, Sigma) were mixed (10 μl final) in a small reaction vial, and dimethylation was allowed to proceed for 5 min on ice. Following incubation, the reaction was quenched by the addition of ammonium bicarbonate in molar excess (0.5 μL of 0.3 M NH_4_HCO_3_).

### Native Gel Electrophoresis

Following the initial dimethylation, samples were prepared for native gel electrophoresis by the addition of loading buffer (4 × NativePage Sample Buffer, Thermo Fisher Scientific) and 1 μg of the GAPDH protein was loaded per lane on a native 4 % to 16 % Bis-Tris gel (NATIVE-PAGE, Thermo). Protein complexes were separated at 15 V/cm in a buffer cooled electrophoresis chamber (Thermo). Gels were fixed in an aqueous solution of 40 % MeOH/10 % acetic acid, microwaved for 45 s and agitated for 30 min at 24 °C. This step was repeated. Gels were subsequently stained (0.02 % Coomassie Blue in 30 % MeOH/10 % acetic acid, BioRad) for 30 min at 24 °C and washed with 8 % acetic acid (30 min at 24 °C). Electrophoretic separation of bovine serum albumin (BSA, Sigma) in an additional sample well facilitated the interpretation of the GAPDH mobility patterns because a GAPDH homo-dimer (71.8 kD) is 5.3 kD heavier than BSA (66.5 kD), and the GAPDH tetramer (145 kD) is more than twice as heavy as BSA.

### Cell culture and heat shock

HEK293T cells were grown under standard conditions (37 °C, 5 % CO_2_) in Dulbecco’s modified Eagle’s medium containing 25 mM Glucose and supplemented with 1 mM Sodium pyruvate, 2 mM Glutamax, 10 % FBS and, 1 % Penicillin/Streptomycin (GIBCO). Following heat shock (15 min, 42 °C, 5 % CO_2_) cells were immediately labeled with isotope defined reagents (2 % formaldehyde, 0.3 M sodium cyanoborohydride, in 1 × Dulbecco’s phosphate buffered saline, pH 7.3) for 15 min at 0 °C. Addition of ammonium bicarbonate (1% final w:v) quenched dimethylation of lysine sites (15 min, 0 °C), cells and cell fragments collected and sonicated for 3 min in a water bath sonicator. A methanol-chloroform precipitation according to ^45^ separated proteins from the initial labeling reagents. Precipitated proteins were resolubilized by sonication (1 h) in 1 % Rapigest (Waters), 0.1 M 4-(2-hydroxyethyl)-1-piperazineethanesulfonic acid (HEPES, Gibco), pH 7.5 and heat denatured (95 °C, 10 min). Disulfide bonds were reduced with 5 mM tris-(2-carboxyethyl)phosphine hydrochloride (TCEP, 20 min, 37 °C) and sulfhydryl moieties were alkylated in 10 mM chloroacetamide (30 min, 24 C).

### Human postmortem brain tissues

100 mg of fresh frozen human postmortem frontal cortex from neuropathologically confirmed AD and cognitively normal control cases was obtained from the Neuropathology/Brain Bank of the Shiley-Marcos Alzheimer’s Disease Research Center of the University of California, San Diego.

### Purification of the insoluble brain fraction

Purification of the insoluble fraction in brain tissue samples was performed as previously described ^19^. In brief, tissue was homogenized in 1 ml tissue lysis buffer (10 % (w:v) sucrose, 10 mM HEPES pH 7.0, 800 mM NaCl, 5 mM Ethylenediaminetetraacetic acid (EDTA), 1 mM ethylene glycol-bis(β-aminoethyl ether)-N,N,N’,N’-tetraacetic acid (EGTA), 1 × protease inhibitors Complete EDTA-free (Roche), 1 × phosphatase inhibitors (Pierce)) with a small pistil, vigorously mixed (30 s), sonicated (30 s), and tissue debris removed by centrifugation (18,000 × g, 30 min, 4 °C). The cleared tissue lysate supernatant was brought to 1 % N-auroylsarcosine (v:v), vigorously mixed (30 min, 24 °C), centrifuged (18,000 × g, 30 min, 4 °C). Protein aggregates were precipitated from the second supernatant by ultracentrifugation (100,000 × g, 1 h, 4 °C) and the protein pellet was isotope labeled and re-solubilized in one step (2 % formaldehyde, 0.3 M sodium cyanoborohydride, in 100 mM Hepes pH 7.0, 10 μl final volume, vigorously mixing, 15 min, 24 °C). The dimethylation reaction was quenched with ammonium bicarbonate (1% final w:v, 5 min, 24 °C). Proteins were denatured (8 M guanidinium chloride, 10 mM TCEP) for 1 h at 37 °C and free sulfhydryl moieties alkylated (20 mM iodoacetamide, 30 min, 24 °C).

### Enzymatic digestion and second labeling step

Only brain samples were diluted to 1 M guanidinium chloride in 0.1 mM HEPES, pH 8.0, 0.02 % Rapigest. All samples were heat denatured (5 min, 95 °C), and proteins were digested with the endoprotease Chymotrypsin at either 5 μg/ml (w:v, brain samples, 16 h, 37 °C) or at a 1: 100 ratio of protease: protein (w:w, GAPDH and HEK293T samples, 16 h, 30 °C).

Rapigest was inactivated by acidification (1 % v:v, 37 °C, 1 h) and the insoluble precipitate removed by centrifugation (18,000 × g, 15 min, 4 °C) in brain-derived or GAPDH samples. Peptides were desalted by C18 reversed phase purification (C18-tips, Thermo Fisher Scientific) to remove residual reagents from the first labeling step. While still bound to the resin newly exposed primary amines on peptides were dimethylated with isotope-defined reagents (2 % formaldehyde, 0.3 M sodium cyanoborohydride, in 100 mM HEPES, pH 7.0, occasional mixing, 15 min, 24 °C) as previously described ^46^. Peptides were eluted with 80 % acetonitrile, 0.01 % trifluoroacetic acid. The eluted samples were evaporated almost to dryness by centrifugation under vacuum, and peptides were resolubilized in liquid chromatography buffer A (5 % acetonitrile, 0.1 % formic acid).

For proteins that were methanol-chloroform precipitated, peptides were directly labeled with isotope defined reagents (2 % formaldehyde, 0.3 M sodium cyanoborohydride, in 100 mM HEPES, pH 7.0, occasional mixing, 1 h, 24 °C). Rapigest was inactivated by acidification (1 % v:v, 37 °C, 1 h), and samples reduced to near dryness *in vacuo* as described above, and finally resolubilized in liquid chromatography buffer A (5 % acetonitrile, 0.1 % formic acid).

### Mass spectrometry

In all experiments, peptides were electrospray ionized at a nano-spray tip of ~0.1 μm i.d. at 1.5 kV. Full scan (400 to 1800 m/z) spectra were acquired on an Orbitrap mass spectrometer (Thermo Fisher Scientific) at a resolution 60,000. Fragment ion spectra of > 1000 ion counts were acquired in data dependent mode for the top 20 highest intense selected ions (z = 2 or higher) with collision-induced dissociation (CID) at 35% collisional energy and recorded in the linear ion trap detector of the mass spectrometer. To avoid sampling only the most abundant peaks, dynamic exclusion with an exclusion list of 500, repeat time of 60 s and asymmetric exclusion window of –0.51 Da and +1.50 Da was used throughout all experiments.

In each experiment samples were chromatographically separated using different methods and mass spectra acquired at different Orbitrap mass spectrometers. Specifically, 250 ng of GAPDH peptides were loaded onto a 300 mm reversed phase chromatographic column with 100 μm inner diameter packed with 100 Å reversed phase resin (Aqua 3, 10 Å pore size, Phenomenex). A linear chromatographic gradient of 100 % buffer A (5% Acetonitrile, 0.1 % formic acid) to 60 % buffer B (80 % acetonitrile, 0.1 % formic acid) was applied over 1.5 h to elute peptides. Mass spectra were acquired with an Orbitrap Fusion mass spectrometer (Thermo Fisher Scientific).

For the heat shock experiment, 50 μg of the CPP labeled HEK293T proteome was loaded onto a MudPIT column ^17^ and analyzed by nano-ESI LC/LC-MS/MS on a VelosPro Orbitrap mass spectrometer. The MudPIT column was placed in line with a quaternary Agilent 1200 high pressure liquid chromatography HPLC pump and peptides were separated by reversed phase liquid chromatography in 10 sequential steps, each following an initial elution of peptides from the strong cation exchange column with buffer C (500 mM ammonium acetate, 5 % acetonitrile, 0.1 % formic acid) in buffer A in incrementally progressive concentrations (0 %, 10 %, 20 %, 30 %, 40 %, 50 %, 60 %, 70 %, 80 %, and 90 %) as described previously ^17,47^.

For patient samples, 2 μg of brain sample derived peptides were loaded onto evotip C18 tips according to the manfacturer’s protocol (EVOSEP, Denmark). Peptides were eluted from the evotip with an EVOSEP HPLC system (EVOSEP, Denmark) and separated by reversed phase chromatography on a 15 cm ReproSil C18 column (3 μm, 120 Å, id 100 μm, PepSep, Denmark) with a 45 min gradient of increasing Acetonitrile concentration with 0.1% formic acid according to manufacturer’s recommendations. Following chromatographic separation, peptides were transferred into an Orbitrap Lumos mass spectrometer by electrospray ionization (nanoEasy, Thermo Fisher Scientific). The top 25 precursor peaks were picked for collision-induced fragmentation.

### Data analysis

Following data acquisition, raw data was pre-processed and converted into ASCII file format with *RawConverter* ^48^ set to monoisotopic peak detection. Converted files were uploaded in *IP2* (Integrated Proteomics) and searched with *ProLuCID* ^49^ for the presence of spectra that matched a theoretical peptide fragment ion spectrum based on amino acid sequences listed in the UniProt database for the human proteome release v 2016.4. Amino acid sequences in the database were either digested *in silico* assuming either no endoproteolytic enzyme specificity (HEK293T cells) or minimally requiring that either the N- or C-terminus of the peptide was generated by chymotryptic cleavage (GAPDH and brain samples). A 50 ppm precursor mass tolerance window was set for peptide candidate selection, carboxyamidomethylation (*m* = 57.021464 Da) of cysteine, and dimethylation (*m* = 28.0313 Da) of N-termini. Peptides including lysine residues labeled “light” or “heavy” (+8.0442 Da) were searched separately as static modifications. Results were filtered with DTASelect v 2.1.4 to a spectrum false discovery rate (FDR) of 0.1 % or less and requiring at least one Chymotrypsin specific cleavage of either peptide N- or C-terminus and a precursor mass tolerance of Δ ≤ ±10 ppm. Subsequently, relative peptide abundances were quantified based on peptide elution profiles deduced from MS survey spectra with Census ^15^ in IP2 (Integrated Proteomics) or based on fragment ion counting in case isotopically labeled peptides were isobaric ^13^. Ratio values for each lysine residue were calculated with the SoPaX algorithm that is part of ProteinClusterQuant ^50^ (PCQ, https://github.com/proteomicsyates/ProteinClusterQuant,). Data presentations were assembled in Excel (Microsoft) or in Prism (GraphPad) to determine the FDR of lysine sites in two sample comparisons according to the modified statistical approach originally proposed by Benjamini and Hochberg ^51^. Panther ^52^ determined the Gene ontology enrichment of protein groups.

Crystal structures of proteins were downloaded from the RCSB Protein Data Bank PDB (https://www.rcsb.org/pdb/home/home.do) and visualized in Jmol (v 14.19.1). All non-protein molecules and hydrogen atoms were removed. Based on the van der Waals spheres of individual atoms the command *isosurface* in Jmol determined the solvent accessible surface (SASA) at the reactive ε-amine of each lysine residue with standard parameter settings (probe radius 1.2 Å and 2 points per Å resolution). Euclidian distances of atoms were determined with the function *distance dependent contacts of one residue with polar residues* that is available in the WHAT IF web interface (http://swift.cmbi.ru.nl/servers/html/index.html) or with the ProteinAssessibilityCalculator (PAC, https://github.com/proteomicsyates/ProteinAccessibilityCalculator).

## Extended data

### Extended data figures

**Extended data figure 1:**
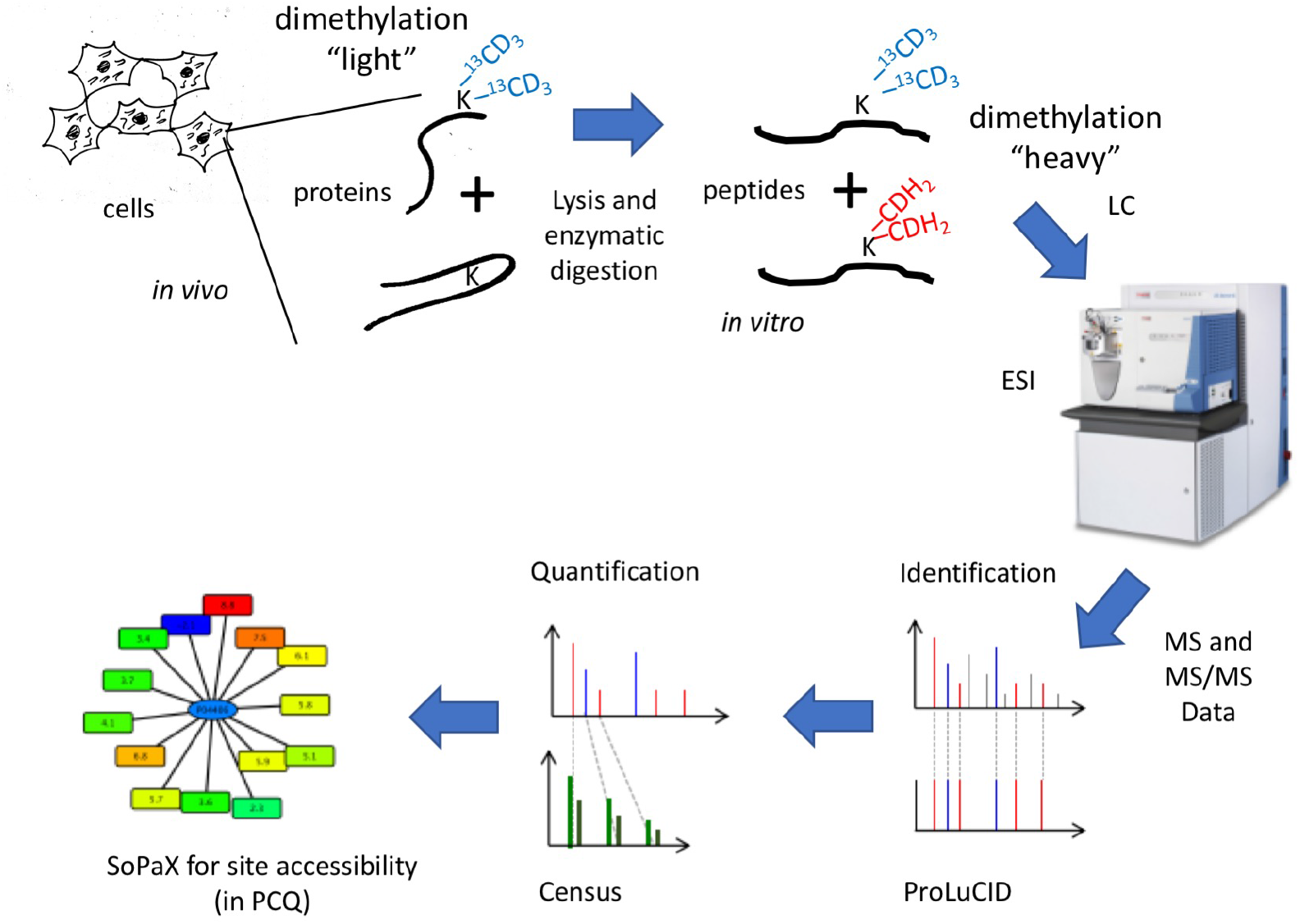
Schematic of the CPP method. CPP includes *in vivo* chemical methylation of lysine residues at the surface of proteins and its detection with mass spectrometry. The flow chart shows each labeling step and the resultant relative measurement of surface accessibility for chemical modification based on mass spectrometric quantification of isotope labeled peptides. Proteins are labeled with isotope defined reagents at solvent exposed lysine residues (K) with two methyl moieties (^13^CD_3_, heavy) *in vivo*. Protein are then digested in peptides with the endoproteinase Chymotrypsin, and all newly accessible primary amines labeled with two methyl moieties (CH_3_, light). Peptides are separated by liquid chromatography (LC) and transferred into gas phase by electrospray ionization (nano ESI). High mass resolution (Orbitrap) mass spectra (MS) and fragment ion mass spectra (MS/MS) are acquired. Peptides are identified with a database search using ProLuCID and quantified with Census. The “surfaces of all protein complexes” (SoPaX) algorithm within ProteinClusterQuant (PCQ) determines and compares the relative surface accessibility of lysine residues.

**Extended data figure 2:**
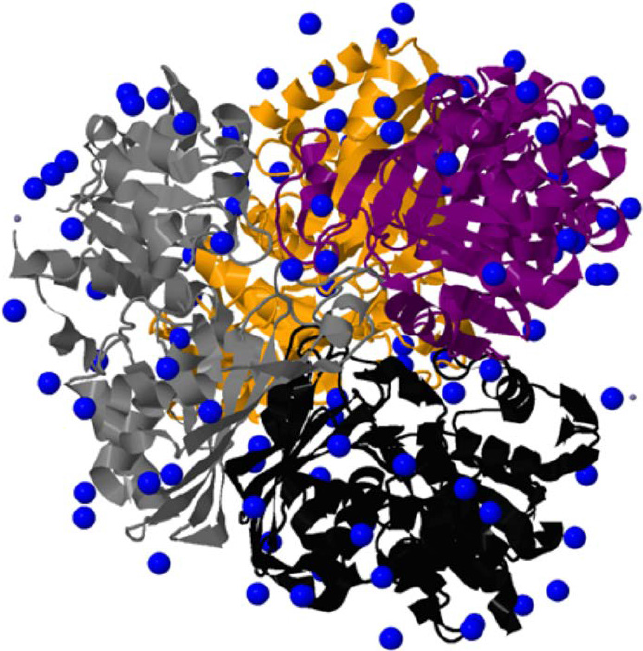
Tetrameric structure of GPADH with the ε-amine of all lysine residues highlighted with blue spheres. Blue dots indicate the location of each lysine ε-amine relative to the ribbon fold of GAPDH subunits A (yellow), B (purple), C (grey), and D (black) in the GAPDH homo-tetramer (PDB: 4wnc ^14^).

**Extended data figure 3:**
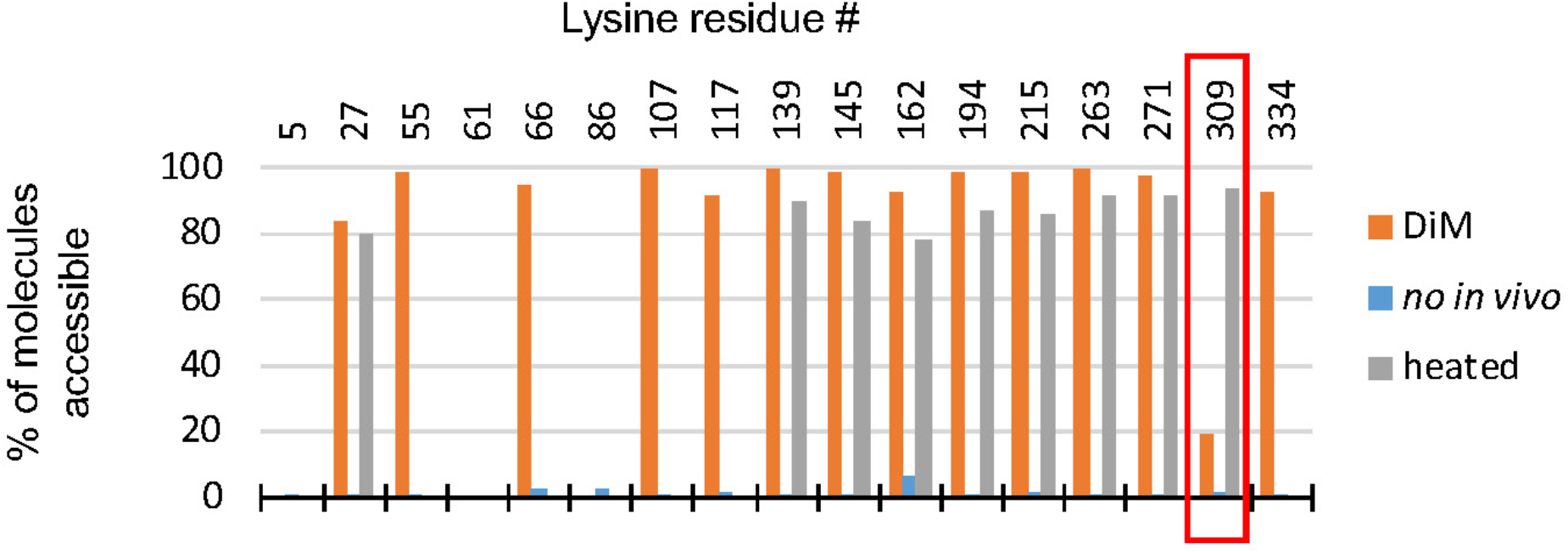
Relative accessibility of lysine residues for chemical modification expressed in percent of molecules. The bar graph shows the same set of measurements as in Figure 1D with the ratio value converted to a percentage of the total amount of lysine residues that were available during the initial labeling step, which is the percentage of GAPDH protein molecules in which the lysine residue was accessible for chemical modification. Orange bars represent relative surface accessibilities determined by standard labeling dimethylation conditions (“DiM”). In addition, GAPDH was heat denatured and subsequently labeled (grey, “heated”), or the initial labeling step was ommitted and only peptides were labeled following endo-proteolytic digestion of GAPDH (blue, “no light labeling”). Measurements for lysine residue GAPDH#K309 are highlighted with a red box.

**Extended data figure 4:**
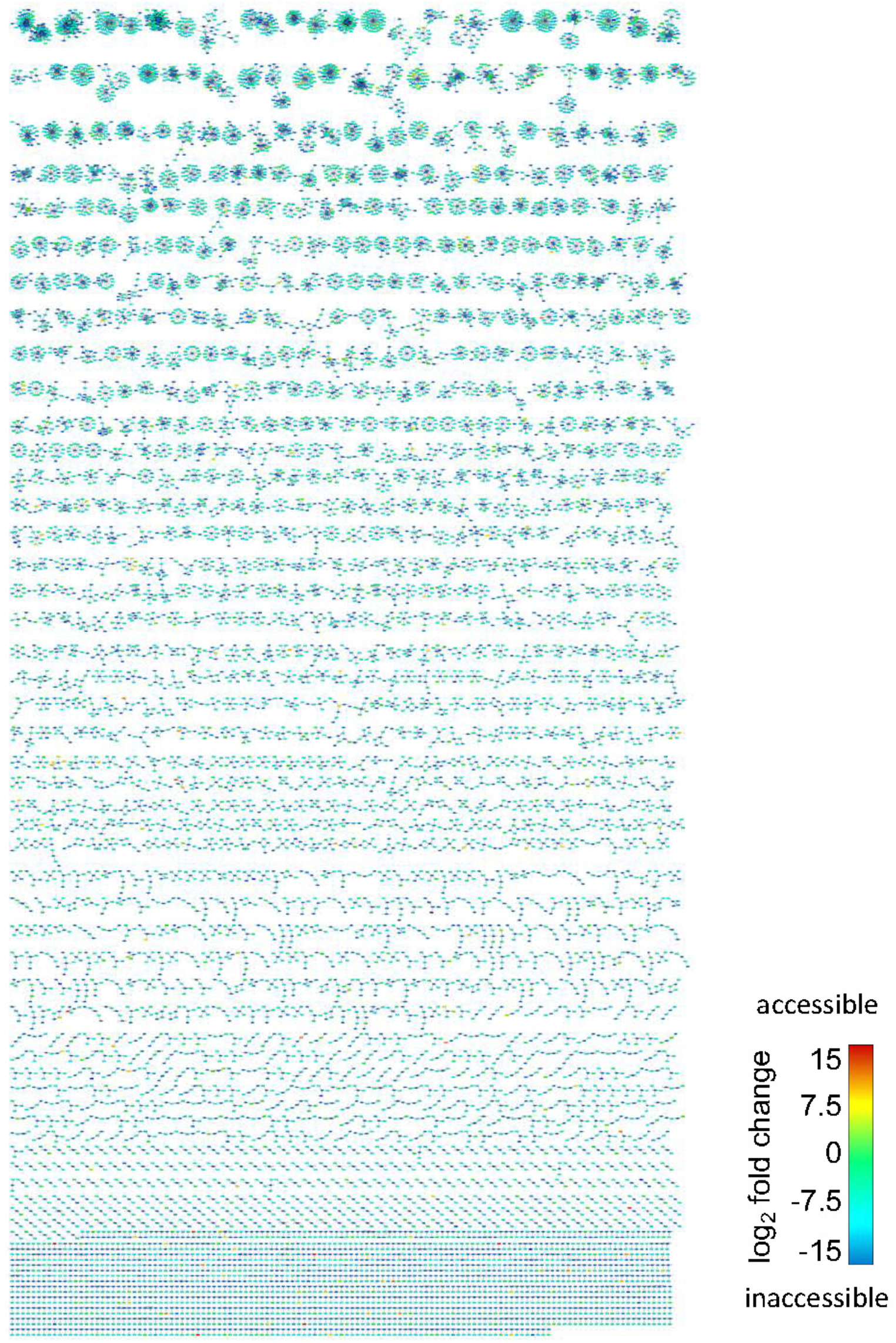
Protein-peptide representation of all lysine residues identified and quantified in HEK293T cells. The bipartite network shows all lysine-harboring peptides (rectangles) to protein (ellipses) relationships as edges that were identified with CPP in HEK293T cells. The color of each peptide node reflects the relative molar accessibility of each lysine residue according to the color scale indicted on the lower right. The predominant color turquoise in the network graph shows that most lysine residues are accessible in the majority of protein molecules.

**Extended data figure 5:**
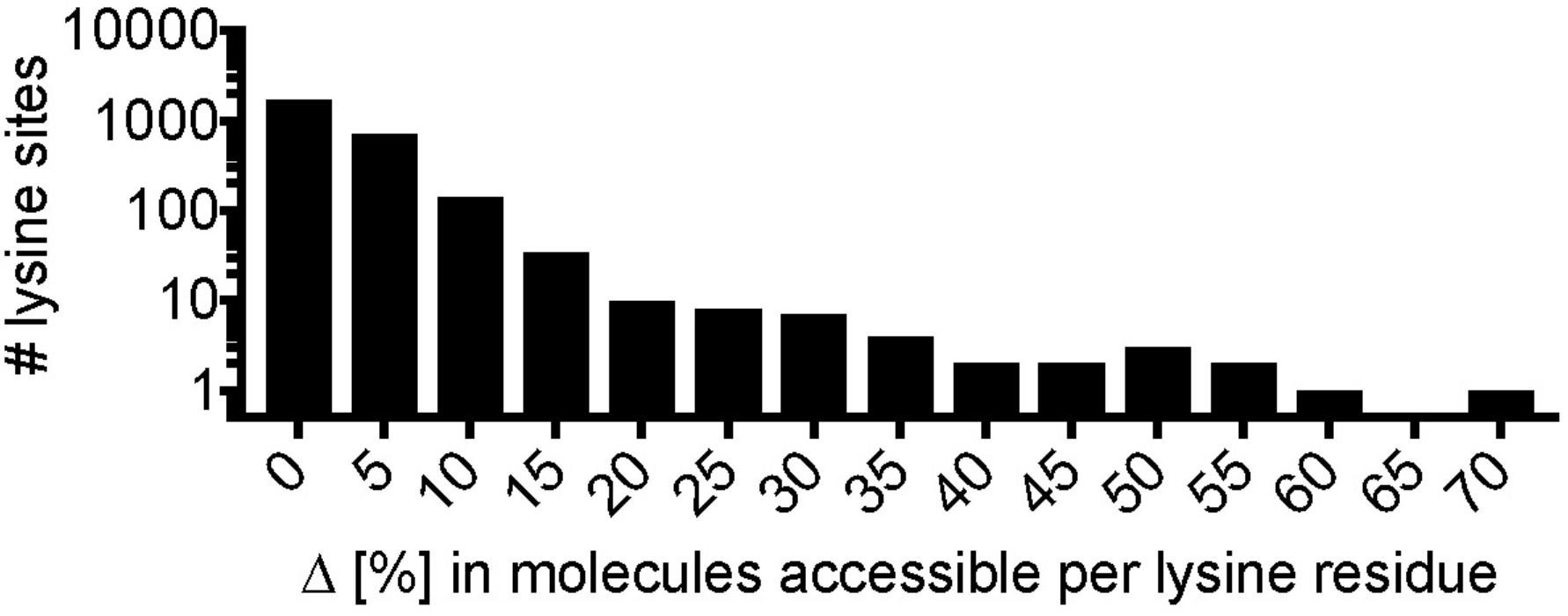
The frequency plot shows the number of lysine residues (y-axis) that alter in surface accessibility per binned percent of protein molecules (percent bin is 5, x-axis).

### Extended data network

**Extended data Network 1**: **The protein-peptide network of control HEK293T cells shown**. The bipartite network in Extended data figure 4 is available on NDEx with the following URL: http://www.ndexbio.org/#/network/dc5d06ab-fa49-11e7-adc1-0ac135e8bacf?accesskey=cbac0a4309d1dc09f5c7dab3be90923e39d7a98616bf64f5ba58feb95f5f88a7

### Extended data tables

**Extended data table 1: Surface accessibility measurements with CPP for lysine residues in HEK293T cells using isobaric methyl moieties**.

**Extended data table 2: Surface accessibility measurements with CPP for lysine residues in heat shock and control HEK293T cells**.

**Extended data table 3: Significantly different sites in HEK293T cells following heat shock**.

**Extended data table 4: Patient diagnosis of human brain samples**. Abbreviations: PM, *postmortem*; BLES, Blessed Orientation-Memory-Concentration Test; MMSE, mini mental state exam; DRS, dementia rating scale.

**Extended data table 5: Isotope combinations selected for labeling in CPP**. The list denotes the experiment and the isotope combination chosen for the first and second labeling step in CPP.

**Data Availability**: Mass spectrometric raw data, search engine result files, and quantification result files can be accessed in Massive or ProteomeXchange (MassIVE MSV000083031, ProteomeXchange PXD011351).

