## Extended data table 5 for "Covalent Protein Painting Reveals Structural Changes in the Proteome in Alzheimer Disease"

| Nnumber | Experiment | First label | Second label | Comment |
| --- | --- | --- | --- | --- |
| 1 | in vitro GAPDH | CH3 | 13CD3 | |
| 2 | HEK cells isobaric isotopologue labeled | 13CH3 | CDH2 | results of experiment number 2- 5 summed up for analysis |
| 3 | HEK cells isobaric isotopologue labeled | CDH2 | 13CH3 | results of experiment number 2- 5 summed up for analysis |
| 4 | HEK cells isobaric isotopologue labeled | 13CD2H | CD3 | results of experiment number 2- 5 summed up for analysis |
| 5 | HEK cells isobaric isotopologue labeled | CD3 | 13CD2H | results of experiment number 2- 5 summed up for analysis |
| 6 | HEK cells upon heat shock exposure | 13D3 | CH3 | |
| 7 | 20 patient sampels | 13CD3 | CDH2 | |
