## Extended data table 4 for "Covalent Protein Painting Reveals Structural Changes in the Proteome in Alzheimer Disease"

| Category | Sample # | AGE [yr] | SEX | PM [h] | BLES | MMSE | DRS | Diagnosis |
|---|---|---|---|---|---|---|---|---|
| A | 1 | 93 | F | 18 | -1 | 30 | 142 | Normal |
| A | 2 | 63 | F | 8 | 0 | 30 | 144 | Normal |
| A | 3 | 102 | F | 9 | 2 | 27 | 134 | Normal |
| A | 4 | 82 | F | 15 | 26 | 1 | 9 | AD |
| A | 5 | 83 | F | 72 | -1 | 29 | 140 | Normal |
| A | 6 | 87 | F | 7 | 3 | 11 | 43 | AD-dLBD |
| A | 7 | 94 | F | 24 | 25 | 13 | 77 | AD |
| A | 8 | 97 | F | 12 | 2 | 26 | 127 | Normal |
| A | 9 | 83 | F | 8 | 33 | 1 | 11 | AD-dLBD |
| A | 10 | 93 | F | 10 | 8 | 22 | 120 | AD |
| B | 1 | 94 | M | 12 | -1 | 30 | 141 | Normal |
| B | 2 | 77 | F | 12 | -1 | -1 | -1 | Normal |
| B | 3 | 84 | M | 36 | 0 | 30 | 143 | Normal |
| B | 4 | 84 | M | 8 | 1 | 28 | 141 | Normal |
| B | 5 | 94 | M | 19 | 2 | 20 | 120 | dLBD |
| B | 6 | 84 | M | 12 | 15 | 21 | 101 | AD-dLBD |
| B | 7 | 84 | M | 6 | 31 | 12 | 81 | AD-dLBD |
| B | 8 | 77 | F | 6 | 12 | 19 | 108 | AD-dLBD |
| B | 9 | 80 | F | 12 | 0 | 29 | 136 | Normal |
| B | 10 | 80 | F | -1 | 13 | 18 | 120 | AD-dLBD |
