## Extended data table 2 for "Covalent Protein Painting Reveals Structural Changes in the Proteome in Alzheimer Disease"

| SEQUENCE | AREA_AVER | AREA_RATIC | AREA_RATIC | AREA_AVER | AREA_RATIC | AREA_RATIC | control % ac | HS % accessi | DELTA |
|---|---|---|---|---|---|---|---|---|---|
| TTSKDGTAGIPNLQLY | 1.28907385 | 3.69164743 | 2 | -6.226412 | 0.16633049 | 2 | 29.0 | 98.7 | 69.6 |
| TIDDGIFEVKATAGDTHLGGEDFDNRLVNHF | 1.40473183 | 5.17609806 | 2 | -2.4983211 | 0.19137649 | 4 | 27.4 | 85.0 | 57.5 |
| IDQEELNKTKPIW | 2.43944767 | 0.26838301 | 4 | -1.3169063 | 5.25290043 | 2 | 15.6 | 71.4 | 55.8 |
| KIVSSSDVGHDEY | 0.10100801 | 3.0058419 | 2 | -4.8498459 | 0.50607419 | 2 | 48.3 | 96.6 | 48.4 |
| VSDRKNIDLLESFVSTF | 0.51315491 | 2.78661068 | 2 | -3.0030574 | 0.317108 | 2 | 41.2 | 88.9 | 47.7 |
| WEGTAVIDGEFKEL | -0.0449402 | 8.49551543 | 2 | -4.0620815 | 0.72807176 | 2 | 50.8 | 94.4 | 43.6 |
| NASNNELVRTKT | 1.58265641 | 6.06679581 | 3 | -1.1026993 | 8.07184957 | 2 | 25.0 | 68.2 | 43.2 |
| VEWIPNNVKTAVCDIPPRGL | -0.1811808 | 4.25652322 | 2 | -3.784727 | 0.27752536 | 2 | 53.1 | 93.2 | 40.1 |
| EELGKQIQQY | -0.4400343 | 3.19306883 | 3 | -4.8997979 | 1.38985087 | 2 | 57.6 | 96.8 | 39.2 |
| MVTEALKPY | -0.6686592 | 7.59985189 | 3 | -5.164139 | 1.30325349 | 3 | 61.4 | 97.3 | 35.9 |
| VIGRGGENVKAINQQTGAF | -0.8039391 | 6.2743697 | 2 | -6.0916372 | 1.44689788 | 3 | 63.6 | 98.6 | 35.0 |
| KVEYTPYEEGLHSVDVTYDGSPVPSSPF | -0.0331732 | 3.55330592 | 2 | -2.2984289 | 0.12066965 | 2 | 50.6 | 83.1 | 32.5 |
| NSKVAVGQEQAF | -0.871338 | 3.72092958 | 3 | -4.9350487 | 2.17449566 | 2 | 64.7 | 96.8 | 32.2 |
| KAPRNDLSPASSGNAVY | -1.0767099 | 4.75866646 | 2 | -8.9971121 | 1.36990936 | 2 | 67.8 | 99.8 | 32.0 |
| ADLLLEKETL | -1.1482552 | 2.29369352 | 2 | -7.7198577 | 3.17621978 | 2 | 68.9 | 99.5 | 30.6 |
| DIDVAKVNTL | -1.1038565 | 4.54655682 | 2 | -5.7183101 | 1.35594793 | 3 | 68.2 | 98.1 | 29.9 |
| TGLAAAIAGAKL | -0.6405179 | 3.42078011 | 2 | -3.2795547 | 0.53749552 | 4 | 60.9 | 90.7 | 29.7 |
| VLVLYDEIKKY | -0.7662212 | 0.66569179 | 8 | -3.5035969 | 0.58971566 | 5 | 63.0 | 91.9 | 28.9 |
| EVCKEKDFSPEAL | -1.1596957 | 3.16423179 | 4 | -4.7173302 | 0.89791586 | 2 | 69.1 | 96.3 | 27.3 |
| GGVPKTTGEGTSL | -1.2583098 | 4.22805515 | 3 | -4.8130635 | 1.06236926 | 4 | 70.5 | 96.6 | 26.0 |
| SIGQAGEFDYSGSQAIKAL | -0.1663909 | 4.15750009 | 2 | -1.8787709 | 4.91068885 | 4 | 52.9 | 78.6 | 25.7 |
| IVAAGVGEFEAGISKNGQ | -1.376736 | 2.58309969 | 4 | -5.0650229 | 1.73470855 | 3 | 72.2 | 97.1 | 24.9 |
| LAGPPLVKAATGEEVSAEDLGGADLH | 3.54445979 | 0.08780716 | 2 | 1.24764694 | 3.30094936 | 2 | 7.9 | 29.6 | 21.7 |
| LEALDCILPPTRPTDKPL | -1.0968425 | 1.79079045 | 2 | -2.9121356 | 0.39642054 | 2 | 68.1 | 88.3 | 20.1 |
| ESALDQLKQF | -1.5791823 | 5.43497276 | 2 | -4.2372214 | 0.94763077 | 4 | 74.9 | 95.0 | 20.0 |
| EQTKVIADNVKDW | -1.9806002 | 3.51838108 | 4 | -6.9523647 | 2.04747202 | 4 | 79.8 | 99.2 | 19.4 |
| LADLVDSCKPGDEIELTGIY | -1.5361753 | 2.59898015 | 4 | -3.8865366 | 0.70691561 | 4 | 74.4 | 93.7 | 19.3 |
| YFDYKEQLPESAY | -2.0039669 | 3.49714162 | 2 | -6.794674 | 0.97700254 | 2 | 80.0 | 99.1 | 19.1 |
| QFADSKGDVGLGLVKEGL | 0.1282804 | 3.88549847 | 2 | -0.9257148 | 4.65807543 | 2 | 47.8 | 65.5 | 17.7 |
| ALKSMTEAEQQQLIDDHF | -1.4091221 | 2.37826288 | 4 | -3.157928 | 0.06454526 | 2 | 72.6 | 89.9 | 17.3 |
| CQPTGGKARLTEGCSF | -1.2377085 | 1.48284894 | 3 | -2.8037511 | 0.1361896 | 3 | 70.2 | 87.5 | 17.3 |
| LAAELSKEGHQVALL | -2.1148076 | 2.81987357 | 3 | -6.0118393 | 1.88714104 | 3 | 81.2 | 98.5 | 17.2 |
| IHAVANNQDKLGFEDGSVL | -1.9711849 | 1.80948334 | 4 | -4.9560198 | 1.58003508 | 3 | 79.7 | 96.9 | 17.2 |
| KDNVSSPGGATIH | -1.9678141 | 1.88763286 | 4 | -4.8861971 | 1.26559178 | 4 | 79.6 | 96.7 | 17.1 |
| EATDARRAFPCWDEPAIKATF | -0.051085 | 0.47289625 | 3 | -1.0483699 | 0.34343391 | 3 | 50.9 | 67.4 | 16.5 |
| SIDSPDSLENIPEKW | -2.0970112 | 2.91355031 | 4 | -5.232119 | 1.21935498 | 4 | 81.1 | 97.4 | 16.4 |
| SFVKSQETECTY | -1.8604688 | 1.28202238 | 2 | -4.1642784 | 0.049746 | 2 | 78.4 | 94.7 | 16.3 |
| NLINKLNNDDNVDGLL | -1.9452258 | 0.33512736 | 2 | -4.3758188 | 0.38570387 | 2 | 79.4 | 95.4 | 16.0 |
| LFDKPVSPL | -2.0407443 | 3.34882811 | 8 | -4.5016402 | 1.70307273 | 5 | 80.4 | 95.8 | 15.3 |
| IGKVVVQVLAEEPEAVL | 2.27728554 | 1.37621563 | 5 | 1.07322341 | 0.41205065 | 3 | 17.1 | 32.2 | 15.1 |
| EVAPISDIIAIKSPDTF | -2.1864231 | 3.15731439 | 4 | -4.8584433 | 1.32969675 | 3 | 82.0 | 96.7 | 14.7 |
| KVTQDELKEVFEDAAEIRL | -2.380649 | 0.27657561 | 2 | -6.0911061 | 0.69396029 | 4 | 83.9 | 98.6 | 14.7 |
| AQTVSPAEKW | -2.4153691 | 0.12161097 | 2 | -6.2457269 | 1.34647153 | 3 | 84.2 | 98.7 | 14.5 |
| KQLDDLKVELSQL | -2.429579 | 0.27547307 | 3 | -6.0447004 | 1.44427237 | 8 | 84.3 | 98.5 | 14.2 |
| QASLDLGTDKF | -2.3546098 | 0.4927429 | 2 | -5.3261526 | 1.076084 | 2 | 83.6 | 97.6 | 13.9 |
| SGTIHAGQPVKVLGENY | -2.4440584 | 3.92487118 | 3 | -5.9037443 | 0.24129423 | 2 | 84.5 | 98.4 | 13.9 |
| YFDENPYFENKVL | -2.4037299 | 2.25708304 | 3 | -5.5979727 | 0.7823242 | 4 | 84.1 | 98.0 | 13.9 |
| IIIGDTGVGKSCL | 0.01801437 | 0.35590827 | 2 | -0.7850635 | 0.44397408 | 4 | 49.7 | 63.3 | 13.6 |
| KYDDAERRFY | -2.4996042 | 0.21026102 | 2 | -6.0679706 | 1.74097762 | 2 | 85.0 | 98.5 | 13.6 |
| NSPGGGGGSDYNYESKFNY | -2.6491275 | 4.69806735 | 3 | -8.1940286 | 2.11933442 | 4 | 86.3 | 99.7 | 13.4 |
| KVEEIAASKC | -2.6629326 | 0.82892016 | 2 | -7.5428667 | 0.67733091 | 3 | 86.4 | 99.5 | 13.1 |
| RVSGQPQSVTASSDKAFEDWLNDDLGSY | -1.9614823 | 0.04608352 | 2 | -3.6526793 | 2.52813651 | 2 | 79.6 | 92.6 | 13.1 |
| TISQATAAGDVIKAAY | -2.1542347 | 3.04707591 | 5 | -4.1445154 | 1.52350427 | 5 | 81.7 | 94.6 | 13.0 |
| IFVNALIENPTFDSQTKENM | -1.9994732 | 0.08029612 | 2 | -3.7215729 | 0.35045393 | 4 | 80.0 | 93.0 | 13.0 |
| NISFPATGCQKL | -2.311099 | 1.14992501 | 5 | -4.6205763 | 1.13889561 | 5 | 83.2 | 96.1 | 12.9 |
| QMNFETAKSRVTQ | -2.2545592 | 0.07952279 | 2 | -4.359634 | 0.23550498 | 2 | 82.7 | 95.4 | 12.7 |
| KLESDYEILERFPGAY | -2.08203 | 1.77764703 | 6 | -3.8417355 | 0.1353411 | 2 | 80.9 | 93.5 | 12.6 |
| HHEGGVDVGDVDAKAQ | -2.7206738 | 2.7179628 | 5 | -7.1343558 | 2.45701102 | 3 | 86.8 | 99.3 | 12.5 |
| LGVGDPKIGAAIQEEL | -2.5091071 | 0.66626881 | 3 | -5.2942596 | 0.87702816 | 2 | 85.1 | 97.5 | 12.5 |
| ELNFMNVKF | -2.7524028 | 0.66224598 | 2 | -7.3331601 | 2.84405027 | 3 | 87.1 | 99.4 | 12.3 |
| MVGSYGPRAEEYEFLTPVEEAPKGM | -1.2567157 | 0.89095841 | 3 | -2.2647927 | 0.17090224 | 2 | 70.5 | 82.8 | 12.3 |
| AGEKILGPF | -2.5065648 | 4.23277424 | 4 | -5.1116765 | 1.5981213 | 4 | 85.0 | 97.2 | 12.2 |
| GVIDVDTGKSTL | -2.7159055 | 2.33382496 | 2 | -6.4468011 | 3.14001355 | 2 | 86.8 | 98.9 | 12.1 |
| TNKGTEDFIVESLDASF | -2.6689079 | 4.15195678 | 3 | -5.9415381 | 2.53368438 | 2 | 86.4 | 98.4 | 12.0 |
| EYTLGEEKFTF | -1.4316847 | 1.39444941 | 4 | -2.483828 | 1.89097502 | 3 | 73.0 | 84.8 | 11.9 |
| AKNDLAVVDVRIGM(15.9949) | -2.2398069 | 1.54413756 | 4 | -4.0706337 | 0.22839528 | 4 | 82.5 | 94.4 | 11.9 |
| FLEDDDKLEQIRKDY | -2.7016209 | 0.3349787 | 3 | -6.0438803 | 1.32943338 | 2 | 86.7 | 98.5 | 11.8 |
| LLGEDGAGKTSL | -2.0111159 | 0.35306997 | 2 | -3.5141737 | 0.18518477 | 2 | 80.1 | 92.0 | 11.8 |
| VIIEKPEEQWW | -2.6467625 | 1.18180071 | 5 | -5.6578439 | 1.49645779 | 3 | 86.2 | 98.1 | 11.8 |
| VQSKDDVIVTASNF | -2.8703832 | 0.59030133 | 3 | -8.5375636 | 2.47375087 | 3 | 88.0 | 99.7 | 11.8 |
| ALTGDEVKKICMQ | -2.708071 | 0.60903409 | 3 | -5.8865046 | 0.77853671 | 6 | 86.7 | 98.3 | 11.6 |

| | | | | | | | | | |
|---|---|---|---|---|---|---|---|---|---|
| ISVEEVHDDGTPTSKTF | -2.8735438 | 2.82163089 | 3 | -7.6379249 | 0.90460984 | 4 | 88.0 | 99.5 | 11.5 |
| AFIQDPDGYWIEILNPNKMATL | -1.7125746 | 1.44395257 | 2 | -2.8489198 | 0.80736902 | 2 | 76.6 | 87.8 | 11.2 |
| VAINKDPEAPIFQVADY | -2.3676464 | 0.21907188 | 3 | -4.1790442 | 0.9354029 | 3 | 83.8 | 94.8 | 11.0 |
| LYFDALECLPEDKEVLTEDKCLQ | -2.500583 | 2.22951091 | 4 | -4.5563392 | 0.3347692 | 2 | 85.0 | 95.9 | 10.9 |
| RAIKQVYEEEYGSSLEDDVVGDTSGYY | -1.4425821 | 0.44636654 | 5 | -2.3839159 | 1.54072282 | 3 | 73.1 | 83.9 | 10.8 |
| TLKGGAAVDPDSGLEH | -2.6147519 | 0.41593568 | 4 | -4.8995513 | 0.73401927 | 2 | 86.0 | 96.8 | 10.8 |
| LVDASKAMFESQSEDELTPF | -1.4422584 | 1.09753065 | 2 | -2.3745231 | 0.12635212 | 2 | 73.1 | 83.8 | 10.7 |
| VDKNFINNPLAQADW | -2.8460038 | 3.34907284 | 4 | -6.0370055 | 0.66351513 | 3 | 87.8 | 98.5 | 10.7 |
| VNRTLSEAGKSTSIQSF | -2.9774325 | 0.33115199 | 9 | -7.4210409 | 4.40762488 | 3 | 88.7 | 99.4 | 10.7 |
| VLENPGTIKVGDPVYLLGQ | -3.0195155 | 1.51633335 | 2 | -7.9754754 | 1.76808668 | 3 | 89.0 | 99.6 | 10.6 |
| KASPGNLSQFEDILF | -2.425585 | 4.32720326 | 3 | -4.2137737 | 0.32053823 | 3 | 84.3 | 94.9 | 10.6 |
| VAKEARNVTMETE | -2.7504651 | 1.76946665 | 3 | -5.3634804 | 0.42734732 | 2 | 87.1 | 97.6 | 10.6 |
| SLFDVLVEKY | -2.0785778 | 0.66523646 | 4 | -3.4122465 | 2.64594334 | 2 | 80.9 | 91.4 | 10.6 |
| SEVKPAGPTVEQQGEM | -2.8018862 | 0.38981655 | 2 | -5.5985186 | 1.67439871 | 3 | 87.5 | 98.0 | 10.5 |
| VSISDLLVPKDLGTESQIF | -1.8981113 | 1.31960996 | 3 | -3.0645443 | 0.10850798 | 2 | 78.8 | 89.3 | 10.5 |
| QSQPDLIGDKLVGGLL | -2.2539484 | 0.26563633 | 3 | -3.7415935 | 1.7488067 | 4 | 82.7 | 93.0 | 10.4 |
| LAVDYENVRPDIVLLGKAL | -0.3383358 | 0.0723873 | 4 | -0.9696444 | 0.28078242 | 2 | 55.8 | 66.2 | 10.4 |
| GHLDDPASQEIERGKSYL | -2.4392763 | 0.85576487 | 4 | -4.1815986 | 1.18367849 | 2 | 84.4 | 94.8 | 10.3 |
| GNIAVEENVDLKHTGAVL | -2.9652191 | 1.09415296 | 3 | -6.6030173 | 0.7393613 | 2 | 88.6 | 99.0 | 10.3 |
| LDFKVVEGSF | -2.9292238 | 0.50857401 | 2 | -6.2078614 | 1.94143066 | 5 | 88.4 | 98.7 | 10.3 |
| GHASDRIIALDGDTKNSTF | 1.39808174 | 0.19431175 | 18 | 0.72250982 | 0.6358131 | 8 | 27.5 | 37.7 | 10.2 |
| EGLNVVKTGRVM | -3.0113082 | 0.47710865 | 4 | -6.9025903 | 1.12281365 | 3 | 89.0 | 99.2 | 10.2 |
| ADLVDSCKPGDEIELTGIY | -2.0489292 | 0.04216802 | 2 | -3.2885258 | 0.19705826 | 2 | 80.5 | 90.7 | 10.2 |
| SLLLETHKNDIPSSY | -2.9600561 | 1.62394913 | 4 | -6.2482984 | 1.47314377 | 2 | 88.6 | 98.7 | 10.1 |
| SGETAKGDYPLEAVRM | -2.7848537 | 0.11292191 | 2 | -5.2216802 | 0.96036039 | 4 | 87.3 | 97.4 | 10.1 |
| TWQGLIVPDNPPYDKGAF | -2.4356092 | 0.22898551 | 5 | -4.0764658 | 2.31353941 | 4 | 84.4 | 94.4 | 10.0 |
| GNNCVFAPADVTSEKDVQTAL | -1.3896257 | 0.13093441 | 2 | -2.2242805 | 0.32670672 | 2 | 72.4 | 82.4 | 10.0 |
| HFTKDIVDAGLAGDTNLY | -2.8492859 | 0.7044671 | 3 | -5.4565561 | 1.47075237 | 3 | 87.8 | 97.8 | 10.0 |
| MATEVAADALGEEWKGY | -2.1092902 | 0.32102909 | 5 | -3.3608638 | 0.90289603 | 2 | 81.2 | 91.1 | 9.9 |
| KDEDGDLITIFDSSDL | -2.8121851 | 0.30466547 | 2 | -5.2712608 | 0.96190313 | 2 | 87.5 | 97.5 | 9.9 |
| IAENQDSLGSKVKDTF | -2.8495415 | 0.92147787 | 3 | -5.4383116 | 1.12203906 | 3 | 87.8 | 97.7 | 9.9 |
| VGDLSPEITTEDIKSAF | -2.4351746 | 0.50966013 | 2 | -4.052688 | 0.78764972 | 2 | 84.4 | 94.3 | 9.9 |
| ALDTNQEERDKGKTVEVGRAY | -2.8661191 | 0.21182111 | 2 | -5.4566272 | 2.60554782 | 2 | 87.9 | 97.8 | 9.8 |
| IINFEEGREVKPTVQ | -3.0299935 | 0.68459976 | 2 | -6.4667396 | 0.59168863 | 2 | 89.1 | 98.9 | 9.8 |
| TGTLIVVPDVSKL | -3.0605833 | 5.00133263 | 3 | -6.7530023 | 2.69152358 | 4 | 89.3 | 99.1 | 9.8 |
| IGDAAKNQLTSNPENTVF | -3.0607947 | 0.57996017 | 4 | -6.5608805 | 2.62442866 | 7 | 89.3 | 99.0 | 9.7 |
| MPQNPCIIATKTPSSDVL | 0.35958161 | 0.22747246 | 8 | -0.1991829 | 0.21379355 | 9 | 43.8 | 53.4 | 9.6 |
| SGVVDQTKDGLISFQEF | 0.93443044 | 0.81882488 | 4 | 0.34918291 | 0.06319208 | 2 | 34.4 | 44.0 | 9.6 |
| QTSKELPQGTSGQLW | -2.9402992 | 1.08412106 | 2 | -5.6841239 | 0.69955813 | 4 | 88.5 | 98.1 | 9.6 |
| TWEPPSVTSGKIIEY | -3.038012 | 0.74642075 | 3 | -6.3125608 | 3.19202762 | 3 | 89.1 | 98.8 | 9.6 |
| VTGVHEEATEEDIHDKFAEYGEIKNIHL | -3.127567 | 0.01542386 | 2 | -7.2149632 | 1.53709867 | 3 | 89.7 | 99.3 | 9.6 |
| NDSQRQATKDAGQISGLNVL | -2.8933891 | 0.89229913 | 7 | -5.4311183 | 1.34189557 | 3 | 88.1 | 97.7 | 9.6 |
| EKSYELPDGQVITIGNERF | -1.1970658 | 0.65357586 | 4 | -1.9308168 | 0.26107835 | 5 | 69.6 | 79.2 | 9.6 |
| LRELISNASDALDKIRY | -2.7090723 | 2.7846059 | 4 | -4.6922983 | 0.51982845 | 4 | 86.7 | 96.3 | 9.5 |
| VALNPDFKPPADY | -3.1304561 | 0.79562469 | 3 | -7.0895522 | 0.71493762 | 3 | 89.8 | 99.3 | 9.5 |
| LGCVDIKDLPVSEQQERAF | -2.0489617 | 0.27947223 | 3 | -3.1765301 | 0.493513 | 2 | 80.5 | 90.0 | 9.5 |
| VADVKNLYPSSSPY | -2.9118055 | 1.01144251 | 2 | -5.4343003 | 0.82432117 | 2 | 88.3 | 97.7 | 9.5 |
| KLEILTNLANEANISTL | -2.8164644 | 0.76858262 | 2 | -5.0152413 | 1.21793411 | 6 | 87.6 | 97.0 | 9.4 |
| GGASHAKGIVL | -3.13383 | 0.43029745 | 3 | -6.9550256 | 2.13532454 | 5 | 89.8 | 99.2 | 9.4 |
| VNNKIISLDLPVAEVY | -2.6868569 | 0.58998214 | 3 | -4.5622296 | 0.3629652 | 2 | 86.6 | 95.9 | 9.4 |
| VIAKMDATANDVPSDRY | -2.7470786 | 0.45403924 | 3 | -4.7409907 | 1.45363031 | 2 | 87.0 | 96.4 | 9.4 |
| NTTSAVTVKSAIR | -3.047577 | 1.30366395 | 3 | -6.0923914 | 1.07814203 | 2 | 89.2 | 98.6 | 9.3 |
| MTDEVNDPSLTIKSIGHQW | -2.9711966 | 0.67609455 | 4 | -5.6364272 | 0.80036783 | 2 | 88.7 | 98.0 | 9.3 |
| VGNLPHDIDENELKEF | -2.9949428 | 0.40149552 | 3 | -5.7509135 | 0.8402582 | 4 | 88.9 | 98.2 | 9.3 |
| NLSFDEINKAF | 0.10105461 | 2.67888022 | 3 | -0.4399483 | 4.30908033 | 3 | 48.2 | 57.6 | 9.3 |
| SSATTPELLLKTF | -2.9277156 | 2.31109101 | 4 | -5.3363714 | 0.35188741 | 2 | 88.4 | 97.6 | 9.2 |
| QGKLPELQGVETELCY | -1.8700788 | 0.91860212 | 5 | -2.8354663 | 0.1559631 | 2 | 78.5 | 87.7 | 9.2 |
| DFPNIVIKGSEL | -3.0455983 | 1.0365059 | 4 | -5.769544 | 1.68664491 | 2 | 89.2 | 98.2 | 9.0 |
| CFASDITCSFPANGKF | -2.2551703 | 0.21498846 | 4 | -3.448554 | 0.90060136 | 3 | 82.7 | 91.6 | 8.9 |
| TATCPQSELDAETVKSILAEY | -2.3118807 | 0.63284095 | 3 | -3.5527415 | 0.52279755 | 4 | 83.2 | 92.1 | 8.9 |
| SLVDAMNGKEGVVECSF | -1.2203867 | 0.60858724 | 3 | -1.9009995 | 0.0911414 | 2 | 70.0 | 78.9 | 8.9 |
| VAEKLNIPW | -3.1675787 | 1.630865 | 2 | -6.4647904 | 3.02060969 | 3 | 90.0 | 98.9 | 8.9 |
| RTDDYLDQPCLETVNRIKLY | -1.9026324 | 0.55542517 | 4 | -2.8428088 | 0.04440369 | 3 | 78.9 | 87.8 | 8.9 |
| IDELDAIGTKRF | -2.6381265 | 0.45162582 | 3 | -4.2415987 | 1.03837173 | 3 | 86.2 | 95.0 | 8.8 |
| DFPKLAETY | -2.6343587 | 1.83864974 | 4 | -4.1774458 | 0.82818926 | 3 | 86.1 | 94.8 | 8.6 |
| VNLGIEPPKGVLL | -3.1468841 | 1.41132211 | 2 | -5.9694057 | 3.66335928 | 2 | 89.9 | 98.4 | 8.6 |
| GLQSKVENF | -3.1861169 | 1.19807125 | 4 | -6.1181677 | 0.54847932 | 3 | 90.1 | 98.6 | 8.5 |
| DLENGKQIKSIDAGPVDAW | -2.5570997 | 0.52704678 | 3 | -3.9533733 | 3.30923942 | 2 | 85.5 | 93.9 | 8.5 |
| REDSVKPGAHL | -2.2038676 | 1.41644691 | 3 | -3.2687476 | 3.61162782 | 2 | 82.2 | 90.6 | 8.4 |
| CQAGDFTNHNGTGGKSIY | -0.4742889 | 1.11362735 | 2 | -0.991659 | 0.11665947 | 3 | 58.1 | 66.5 | 8.4 |
| SAPKTDMDNQIVVSDY | -3.2602149 | 0.89373112 | 2 | -6.434699 | 1.17393496 | 3 | 90.5 | 98.9 | 8.3 |
| GIVTNWDDMEKIW | -3.1085589 | 1.27740502 | 2 | -5.5330093 | 0.65216569 | 4 | 89.6 | 97.9 | 8.3 |

| | | | | | | | | | |
|---|---|---|---|---|---|---|---|---|---|
| GGVDEKGPQLF | -2.7086486 | 4.19245456 | 4 | -4.2312407 | 0.8094213 | 3 | 86.7 | 94.9 | 8.2 |
| KTIISYIDEQFERY | -2.7163535 | 0.52797722 | 3 | -4.228645 | 0.74153455 | 3 | 86.8 | 94.9 | 8.1 |
| LGDVISIQPCPDVKY | -3.1676556 | 1.0733744 | 2 | -5.7121219 | 1.45192009 | 3 | 90.0 | 98.1 | 8.1 |
| ANDPDADRLAVAEKQDSGEW | -2.0609116 | 0.96603836 | 2 | -2.9882025 | 0.02387867 | 2 | 80.7 | 88.8 | 8.1 |
| IANKPEFDHLAEY | -3.2913294 | 0.52164912 | 5 | -6.4395034 | 1.00103112 | 3 | 90.7 | 98.9 | 8.1 |
| KYITSLNEDSTVH | -3.0969649 | 1.05286962 | 2 | -5.3675451 | 0.7115739 | 2 | 89.5 | 97.6 | 8.1 |
| GIGRSFEEAFQKAL | 0.32700267 | 0.15097375 | 3 | -0.1417494 | 0.0137204 | 3 | 44.4 | 52.5 | 8.1 |
| LKDVIATDKEDVAF | -3.367257 | 1.05278177 | 5 | -7.037188 | 1.78898293 | 3 | 91.2 | 99.2 | 8.1 |
| KLRPSGDDVELIGEEEDVSVY | -2.4725344 | 0.34501169 | 3 | -3.6862051 | 0.14236238 | 2 | 84.7 | 92.8 | 8.1 |
| GETNPADSKPGTIRGDF | -2.8265145 | 0.05784262 | 4 | -4.4710239 | 0.31596187 | 4 | 87.6 | 95.7 | 8.0 |
| LLKDVNEDPGEDVALL | -3.1942016 | 0.58826663 | 2 | -5.7578093 | 0.43042584 | 3 | 90.2 | 98.2 | 8.0 |
| GGNVLGPKSVA | -2.9853432 | 1.04295678 | 3 | -4.9247993 | 0.83813806 | 3 | 88.8 | 96.8 | 8.0 |
| EAAITPETKLVW | -3.319744 | 0.84749904 | 4 | -6.4174312 | 2.08303418 | 3 | 90.9 | 98.8 | 7.9 |
| SFVKSQETECTYF | -2.87993 | 0.03634354 | 2 | -4.5768412 | 0.26655914 | 2 | 88.0 | 96.0 | 7.9 |
| KYITSLNEDSTVHGF | -3.4261314 | 0.10281273 | 2 | -7.3435862 | 0.24134581 | 2 | 91.5 | 99.4 | 7.9 |
| VAPTANLDQKDKQF | -3.3558647 | 2.16260835 | 4 | -6.591707 | 1.90729245 | 7 | 91.1 | 99.0 | 7.9 |
| IQENLELVEKGF | -3.0816997 | 1.03423035 | 3 | -5.1725513 | 1.69478448 | 5 | 89.4 | 97.3 | 7.9 |
| VFIPEDDPLFPPIEKF | -2.701016 | 0.43757034 | 2 | -4.1128485 | 0.66554927 | 2 | 86.7 | 94.5 | 7.9 |
| EETSGVSVGDPVLRTGKPL | -3.095846 | 0.68636492 | 7 | -5.2055397 | 2.56237223 | 4 | 89.5 | 97.4 | 7.8 |
| TKDIDIHEVRIGW | -2.9280785 | 1.20184084 | 6 | -4.6592626 | 1.42076765 | 3 | 88.4 | 96.2 | 7.8 |
| QKENGTVTAANASTLNDGAAAL | -2.5194762 | 0.7610699 | 4 | -3.7128769 | 0.27764383 | 2 | 85.1 | 92.9 | 7.8 |
| NQVKTIAQGNLSNTDVQ | -2.9918822 | 0.17351201 | 4 | -4.8229626 | 0.92594941 | 4 | 88.8 | 96.6 | 7.8 |
| SRKEEQEVQATL | -2.927756 | 1.02884945 | 2 | -4.6332553 | 2.47346887 | 3 | 88.4 | 96.1 | 7.7 |
| IYDPIKTAQGSLSL | -1.3040783 | 3.94473605 | 4 | -1.9034626 | 2.90573984 | 4 | 71.2 | 78.9 | 7.7 |
| NGDNM(15.9949)LEPSANMPWFKGW | -2.475836 | 0.53266429 | 6 | -3.6212056 | 0.23899294 | 4 | 84.8 | 92.5 | 7.7 |
| NAIKLEEKGVY | -3.1515438 | 0.45061287 | 4 | -5.3457989 | 1.26003108 | 2 | 89.9 | 97.6 | 7.7 |
| RGTLDPVEKAL | -3.236482 | 0.81875288 | 2 | -5.6739721 | 1.28291357 | 6 | 90.4 | 98.1 | 7.7 |
| QDRISGASEKDIVHSGLAY | -3.0048676 | 1.46682215 | 13 | -4.825156 | 1.64742165 | 3 | 88.9 | 96.6 | 7.7 |
| SDCKIQNGAQGIRF | -3.5124041 | 1.67092311 | 4 | -7.8674132 | 2.15702235 | 3 | 91.9 | 99.6 | 7.6 |
| IANTGMDTDKIKIF | -3.0263759 | 0.39639503 | 4 | -4.8642908 | 0.51289109 | 5 | 89.1 | 96.7 | 7.6 |
| KDVLEVGELAKLAY | -3.2958181 | 3.76660128 | 3 | -5.8954672 | 0.57369743 | 4 | 90.8 | 98.3 | 7.6 |
| TQTVCLDDTTVKF | -3.35914 | 0.95453933 | 2 | -6.2327318 | 1.34441356 | 2 | 91.1 | 98.7 | 7.6 |
| VTKVSVGEF | -3.0277251 | 0.32700339 | 3 | -4.8420334 | 2.25946587 | 3 | 89.1 | 96.6 | 7.6 |
| VANFDENDPKTF | -3.0386482 | 0.97861237 | 2 | -4.8673012 | 1.22163033 | 3 | 89.2 | 96.7 | 7.5 |
| SLARALEEAMEQKAELERL | -1.980412 | 0.3737814 | 3 | -2.7788479 | 0.03295556 | 3 | 79.8 | 87.3 | 7.5 |
| TLKLEDTENW | -3.2653815 | 2.64644047 | 4 | -5.6557571 | 0.71207505 | 5 | 90.6 | 98.1 | 7.5 |
| KQGLNGVPILSEEELSLLDEFY | -2.56861 | 0.43739814 | 2 | -3.736628 | 0.34136375 | 2 | 85.6 | 93.0 | 7.4 |
| VLEDLGDGQKANDDIIVNW | -2.0979381 | 0.65183713 | 2 | -2.9442514 | 0.73980689 | 2 | 81.1 | 88.5 | 7.4 |
| ACAVVCIQKADIF | -3.0617244 | 0.35233359 | 2 | -4.8683273 | 0.34430629 | 2 | 89.3 | 96.7 | 7.4 |
| CFGPDGTGPNILTDITKGVQY | -2.0491822 | 1.94047866 | 8 | -2.8632283 | 0.30406593 | 8 | 80.5 | 87.9 | 7.4 |
| ELIVQKLLETTDRPDGHQN | -2.1993208 | 0.58040421 | 2 | -3.0853741 | 0.13770764 | 3 | 82.1 | 89.5 | 7.3 |
| SSAEVVVPRDQTPDENDQVVVKITGHFY | -1.551003 | 0.51809558 | 2 | -2.1759157 | 0.05438837 | 2 | 74.6 | 81.9 | 7.3 |
| DAGAGIALNDHFVKLISWY | -1.0633535 | 1.27721037 | 6 | -1.5792388 | 0.7659676 | 9 | 67.6 | 74.9 | 7.3 |
| VASLAEPDFVVTDFAKF | 2.00316811 | 0.36248067 | 9 | 1.41862907 | 0.78749568 | 6 | 20.0 | 27.2 | 7.3 |
| DIQVPNFPADETKGF | -2.9509489 | 0.26582072 | 3 | -4.5014183 | 0.67786225 | 2 | 88.5 | 95.8 | 7.2 |
| HLELDISDSKIRY | -3.3332347 | 5.61491446 | 2 | -5.7669016 | 0.00499933 | 2 | 91.0 | 98.2 | 7.2 |
| NAIVIKETKDWDAW | -3.4401883 | 0.43737665 | 3 | -6.3443644 | 2.95620182 | 2 | 91.6 | 98.8 | 7.2 |
| AAALETFTEGQKLDSADANF | -2.2250432 | 0.43467875 | 2 | -3.1021605 | 0.47538328 | 2 | 82.4 | 89.6 | 7.2 |
| DSGFGGGAGVETGGKL | -3.1342319 | 0.05104896 | 2 | -4.997165 | 0.8771592 | 2 | 89.8 | 97.0 | 7.2 |
| VATDQTERIVEPPENIQEKIAF | -2.5839062 | 0.66105032 | 2 | -3.7080594 | 0.41025643 | 2 | 85.7 | 92.9 | 7.2 |
| VALDFEQEMATAASSSSLEKSY | -2.3234366 | 0.12047436 | 3 | -3.2523995 | 0.50123403 | 2 | 83.3 | 90.5 | 7.2 |
| NTDNTLGTEISWENKLAEGL | -2.5433734 | 0.37212999 | 3 | -3.6233462 | 0.40660203 | 3 | 85.4 | 92.5 | 7.1 |
| IATDVASRGLDVEDVKF | -3.3108623 | 1.99824144 | 4 | -5.5984551 | 1.6662824 | 4 | 90.8 | 98.0 | 7.1 |
| GDVGKGCAQAL | -3.250874 | 0.41462209 | 2 | -5.3350488 | 1.26273456 | 4 | 90.5 | 97.6 | 7.1 |
| SLEQALPPEPKEENAEPVSKL | -3.4013108 | 0.95321783 | 2 | -5.9808881 | 0.84777169 | 2 | 91.4 | 98.4 | 7.1 |
| EKSYELPDGQVITIGNERF | -1.5511057 | 0.95760221 | 2 | -2.1515357 | 0.2241279 | 4 | 74.6 | 81.6 | 7.1 |
| GEDALANVSIEKPIHQGPDAAVTGH | -2.5857941 | 0.51330174 | 2 | -3.6711625 | 0.21467749 | 2 | 85.7 | 92.7 | 7.0 |
| NGKEPSRGINPDEAVAY | -3.6427936 | 0.89869217 | 4 | -7.9134054 | 1.78807128 | 3 | 92.6 | 99.6 | 7.0 |
| KDEDGDLITIFDSSDLSF | -3.2890231 | 0.8285745 | 4 | -5.4154967 | 0.26779343 | 5 | 90.7 | 97.7 | 7.0 |
| KTDLRFQSAAIGALQEASEAY | -1.2872688 | 0.52722703 | 7 | -1.8186809 | 0.35532896 | 2 | 70.9 | 77.9 | 7.0 |
| VLHCQGTEEEKILY | -3.3207202 | 2.26713978 | 3 | -5.5255586 | 0.06397195 | 3 | 90.9 | 97.9 | 7.0 |
| SINAEEVVVGDLVEVKGGDRIPADL | -2.6794395 | 0.62926536 | 6 | -3.8315639 | 1.05894809 | 2 | 86.5 | 93.4 | 6.9 |
| NVEFDDSQDKAVL | -3.508344 | 0.78017935 | 3 | -6.4270494 | 0.77142152 | 3 | 91.9 | 98.9 | 6.9 |
| QDLKDFMRQAGEVTY | -3.2448963 | 1.04515382 | 2 | -5.213548 | 1.78369338 | 3 | 90.5 | 97.4 | 6.9 |
| ALELLFDQLHEGAKAL | -1.8829675 | 0.4394684 | 4 | -2.5692493 | 0.90676317 | 3 | 78.7 | 85.6 | 6.9 |
| NNPKVQASL | -3.6780384 | 0.39398934 | 5 | -8.1710275 | 0.33667987 | 3 | 92.8 | 99.7 | 6.9 |
| KVGHVTERTDASSASSF | -3.6754915 | 0.9127627 | 3 | -8.0089805 | 2.76733841 | 2 | 92.7 | 99.6 | 6.9 |
| QLDNNFEVKSL | -3.5001911 | 3.75926148 | 5 | -6.284992 | 0.51803959 | 4 | 91.9 | 98.7 | 6.9 |
| SALVDGKSINAGGH | -3.6339949 | 0.59837396 | 4 | -7.3072553 | 0.84638082 | 2 | 92.5 | 99.4 | 6.8 |
| SLVFDPVQKTLL | -3.7168374 | 0.45705235 | 2 | -8.632406 | 1.88568167 | 2 | 92.9 | 99.7 | 6.8 |
| GATKADFDNTVAIHPTSSEELVTLR | -1.6921383 | 0.09897388 | 3 | -2.3047132 | 0.64694494 | 5 | 76.4 | 83.2 | 6.8 |
| CNDQSTGDIKVIGGDDLSTL | -2.5832758 | 0.28466657 | 3 | -3.6200593 | 0.66902507 | 4 | 85.7 | 92.5 | 6.8 |

| | | | | | | | | | |
|---|---|---|---|---|---|---|---|---|---|
| AESVEKAIEEKKY | -3.4681378 | 4.01149843 | 5 | -6.0269936 | 1.60898562 | 3 | 91.7 | 98.5 | 6.8 |
| IKPGAIVIDCGINY | -3.1213239 | 1.74656198 | 4 | -4.7717852 | 0.89564062 | 5 | 89.7 | 96.5 | 6.8 |
| NGLKEEDKEPLIELF | -3.2492021 | 0.75832929 | 4 | -5.1407934 | 0.75265742 | 2 | 90.5 | 97.2 | 6.8 |
| SVASASVIEKQNLEGDHSDAPMETETSF | -1.7192069 | 0.42727479 | 2 | -2.3352215 | 0.5155175 | 2 | 76.7 | 83.5 | 6.8 |
| IKNMITGTSQADCAVL | -3.3910997 | 1.81705252 | 2 | -5.6487931 | 1.34883532 | 3 | 91.3 | 98.0 | 6.7 |
| GKDFNDIRQDFLPW | -3.1394831 | 0.23526209 | 3 | -4.8092171 | 1.24566064 | 2 | 89.8 | 96.6 | 6.7 |
| LSDYEVCKEGDVLTPEQARVL | -2.1751978 | 0.38854054 | 3 | -2.9604014 | 0.06565378 | 2 | 81.9 | 88.6 | 6.7 |
| MVDKDGDVTVTNDGATIL | -3.4120208 | 0.4600855 | 2 | -5.7199271 | 0.98455857 | 2 | 91.4 | 98.1 | 6.7 |
| IQHAKEDETRY | -3.3136865 | 0.97288032 | 2 | -5.338133 | 1.4188251 | 3 | 90.9 | 97.6 | 6.7 |
| AAAAAAEPPFYKDVW | -3.1192658 | 1.13383558 | 4 | -4.7378146 | 0.92322232 | 4 | 89.7 | 96.4 | 6.7 |
| FNPNDDGKEEPPTTLLW | -3.3496213 | 0.03712276 | 2 | -5.454819 | 1.27918872 | 2 | 91.1 | 97.8 | 6.7 |
| VIANDVDNKRCY | -3.4940373 | 0.5217629 | 5 | -6.0866522 | 1.90989736 | 3 | 91.8 | 98.5 | 6.7 |
| LISASSKDTGQLY | -3.2067766 | 1.65067405 | 3 | -4.9779125 | 0.1474332 | 2 | 90.2 | 96.9 | 6.7 |
| DLENLPASKDSIVH | -3.312325 | 1.59481378 | 2 | -5.315104 | 1.030312 | 3 | 90.9 | 97.5 | 6.7 |
| VWNTHADFADECPKPELL | -2.6908207 | 1.46321496 | 2 | -3.7910004 | 0.34112428 | 2 | 86.6 | 93.3 | 6.7 |
| SASGELGNGNIKL | -3.5974147 | 0.43261746 | 3 | -6.6646331 | 0.62955348 | 2 | 92.4 | 99.0 | 6.7 |
| KIPVIENLGATLDQFDAIDF | -2.1917183 | 0.87206743 | 4 | -2.9714433 | 0.82945045 | 5 | 82.0 | 88.7 | 6.7 |
| TEALGIDPNNIKTNAKLY | -3.5014448 | 1.53623792 | 4 | -6.0687098 | 2.2214606 | 3 | 91.9 | 98.5 | 6.6 |
| TASLLIDVITVFGELTDENVKH | -1.7224114 | 0.60717177 | 2 | -2.3273625 | 0.23972714 | 2 | 76.7 | 83.4 | 6.6 |
| TLDNKLLQTDDEEEAGLL | -3.6397752 | 1.55492861 | 2 | -6.9777092 | 2.70323089 | 3 | 92.6 | 99.2 | 6.6 |
| SVNDPPDVLDRQKCL | -3.7208734 | 0.87105633 | 5 | -7.8628142 | 1.82936305 | 4 | 93.0 | 99.6 | 6.6 |
| AKATGATQQDANASSLL | -3.6609825 | 1.94721984 | 6 | -7.107975 | 2.51196575 | 3 | 92.7 | 99.3 | 6.6 |
| DVVRKESESCDCLQGF | -2.3176287 | 0.65209457 | 2 | -3.1518615 | 0.02574964 | 2 | 83.3 | 89.9 | 6.6 |
| CLRLAEDAPNFDGPAAEGQPGQKQSTTF | -2.0905264 | 0.38253882 | 3 | -2.8178341 | 0.61592747 | 4 | 81.0 | 87.6 | 6.6 |
| ETLLSQNQGGKTF | -2.3867909 | 0.28303362 | 2 | -3.2578275 | 3.58724979 | 4 | 83.9 | 90.5 | 6.6 |
| LLKVNQIGSVTE | -1.3435359 | 0.241822 | 7 | -1.8529594 | 0.90432456 | 6 | 71.7 | 78.3 | 6.6 |
| EIQDIYENSWTKL | -3.5120014 | 1.33730915 | 3 | -6.0627153 | 1.0684674 | 7 | 91.9 | 98.5 | 6.6 |
| KGGAAVDPDSGLEHSAH | -3.0224654 | 1.25032431 | 6 | -4.4463153 | 1.59018117 | 4 | 89.0 | 95.6 | 6.6 |
| VSKTGEAETITSHY | -3.6352416 | 1.39551698 | 2 | -6.7907762 | 2.81118033 | 3 | 92.6 | 99.1 | 6.6 |
| TVEKADNFEYSDPVDGSISRNQGL | -2.5574953 | 0.37886959 | 3 | -3.5295068 | 0.24420583 | 2 | 85.5 | 92.0 | 6.6 |
| KAADLNGDLTATREEF | -3.7637358 | 1.07854293 | 3 | -8.1990178 | 1.55209579 | 3 | 93.1 | 99.7 | 6.5 |
| FYPVVKEQSQPCADNAVL | -2.6561651 | 0.36558624 | 5 | -3.6933551 | 0.95710218 | 3 | 86.3 | 92.8 | 6.5 |
| IAKEEMIHNLQ | -3.4054442 | 0.86066086 | 4 | -5.5337584 | 1.23548496 | 4 | 91.4 | 97.9 | 6.5 |
| FVEWIPNNVKTAVCDIPPRGL | -2.7968685 | 0.93765869 | 2 | -3.9493762 | 0.04536891 | 3 | 87.4 | 93.9 | 6.5 |
| LKEGTDSSQGIPQLVSNISACQVIAEAVRTTL | -1.2738729 | 2.51222171 | 5 | -1.7608387 | 0.31213029 | 4 | 70.7 | 77.2 | 6.5 |
| YPDGGDQETAKTGKF | -3.5760943 | 0.41237478 | 3 | -6.263098 | 2.27205065 | 3 | 92.3 | 98.7 | 6.5 |
| VSNWDEATKRSLL | -3.554379 | 0.81449963 | 5 | -6.070501 | 1.69770508 | 3 | 92.2 | 98.5 | 6.4 |
| GIHEEMLKDEVRTL | -3.1602061 | 0.72924021 | 6 | -4.7018718 | 0.70328737 | 4 | 89.9 | 96.3 | 6.4 |
| DSIVMDPKKDVLIEF | -3.6882599 | 1.24411401 | 2 | -6.8741095 | 1.36336055 | 2 | 92.8 | 99.2 | 6.4 |
| VGGFSYADVLGSAKGW | -3.5269919 | 1.9497562 | 5 | -5.8867718 | 0.74338413 | 2 | 92.0 | 98.3 | 6.3 |
| SKEDIERMVQEAEKY | -2.6813285 | 0.90207174 | 2 | -3.6914973 | 0.31545248 | 3 | 86.5 | 92.8 | 6.3 |
| SDPFVEAEKSNL | -3.4734245 | 0.34675355 | 2 | -5.6336264 | 0.35326474 | 2 | 91.7 | 98.0 | 6.3 |
| KQVVESAYEVIKL | -3.7840439 | 5.36724563 | 4 | -7.6564418 | 2.10094598 | 2 | 93.2 | 99.5 | 6.3 |
| AIQTQQSKF | -3.7886723 | 1.104068 | 3 | -7.6904755 | 1.13168746 | 2 | 93.3 | 99.5 | 6.3 |
| NAAFLESSAKENQTAVDVF | -3.4797019 | 0.09840155 | 2 | -5.6410942 | 1.28233684 | 4 | 91.8 | 98.0 | 6.3 |
| DAGAGIALNDHFVKLISWYDNEFGY | -1.153386 | 1.29637894 | 12 | -1.6040838 | 0.90770099 | 12 | 69.0 | 75.2 | 6.3 |
| LSVAKGSDEPPVFLEIHY | -3.7742219 | 0.60529454 | 3 | -7.4977052 | 2.39230775 | 3 | 93.2 | 99.4 | 6.3 |
| INIVVIGHVDSGKSTT | -3.3835298 | 0.95668524 | 3 | -5.2946348 | 1.81559903 | 4 | 91.3 | 97.5 | 6.3 |
| GHASDRIIALDGDTKNSTF | 1.30233629 | 0.17522309 | 18 | 0.88652608 | 0.71397099 | 13 | 28.8 | 35.1 | 6.3 |
| VIKVPDNYGDEIAIEL | -3.2753597 | 0.98727573 | 2 | -4.9594773 | 0.72573486 | 2 | 90.6 | 96.9 | 6.2 |
| GYPNLKSVNEL | -3.49874 | 0.44098192 | 2 | -5.6911701 | 1.33447216 | 4 | 91.9 | 98.1 | 6.2 |
| SYAVENAKDIIACGF | -3.1593302 | 1.13313127 | 3 | -4.646493 | 1.504073 | 5 | 89.9 | 96.2 | 6.2 |
| DASKVTASGPGLSSY | -3.4285835 | 1.12254128 | 4 | -5.4221934 | 0.94442707 | 2 | 91.5 | 97.7 | 6.2 |
| SIAKAGIICQL | -2.1670874 | 0.4805088 | 3 | -2.8747874 | 1.30968893 | 2 | 81.8 | 88.0 | 6.2 |
| GSTSTICSDKTGTL | -3.8400808 | 1.43165131 | 3 | -8.2960019 | 1.64407378 | 3 | 93.5 | 99.7 | 6.2 |
| EILQSVDDAAIVIKNTKEPPLSL | -3.0939311 | 0.84459489 | 2 | -4.4759391 | 1.57952538 | 2 | 89.5 | 95.7 | 6.2 |
| QKRGTGGVDTAAVGGVF | -3.3111334 | 1.09055013 | 4 | -5.0273377 | 1.12920711 | 5 | 90.8 | 97.0 | 6.2 |
| ELDHKNAQAQQEF | -3.7425452 | 0.12240801 | 2 | -6.9959538 | 2.46993993 | 4 | 93.0 | 99.2 | 6.2 |
| LAEVPTMAVEKVLVY | -3.413436 | 0.40492934 | 3 | -5.3326788 | 1.18476349 | 5 | 91.4 | 97.6 | 6.2 |
| VKLISWYDNEFGY | 0.2212411 | 1.68078846 | 3 | -0.1340971 | 1.08530405 | 8 | 46.2 | 52.3 | 6.1 |
| EFLDKLDVVRSFL | -3.0956153 | 1.16720382 | 2 | -4.4623797 | 1.76167659 | 3 | 89.5 | 95.7 | 6.1 |
| GFEIIETKKNYY | -3.7108913 | 0.8418087 | 3 | -6.6479747 | 2.88901978 | 3 | 92.9 | 99.0 | 6.1 |
| KGSDFDCELRLL | -3.3508719 | 0.55118501 | 5 | -5.1009484 | 2.37442118 | 2 | 91.1 | 97.2 | 6.1 |
| SSKIIGINGDFF | -3.4628093 | 1.1200761 | 6 | -5.4500911 | 1.11442978 | 8 | 91.7 | 97.8 | 6.1 |
| VIEFTEQTAPKIF | -3.4352337 | 3.02395696 | 6 | -5.3466195 | 0.62289052 | 2 | 91.5 | 97.6 | 6.1 |
| VLQEKPIESFNEYLF | -3.2466483 | 0.32302228 | 4 | -4.7921523 | 1.61063087 | 2 | 90.5 | 96.5 | 6.0 |
| KYDGSTIVPGEQGAEY | -3.8667064 | 1.55703765 | 3 | -8.071042 | 1.73908033 | 3 | 93.6 | 99.6 | 6.0 |
| QKAAEEVEAKF | -3.5525124 | 1.34386303 | 5 | -5.7575066 | 1.6067048 | 4 | 92.1 | 98.2 | 6.0 |
| NDSQRQATKDAGTIAGL | -3.7212565 | 1.5380677 | 4 | -6.6100279 | 0.53223721 | 4 | 93.0 | 99.0 | 6.0 |
| NSGKVDIVAINDPFIDLNY | -2.7665184 | 0.16585852 | 9 | -3.7812289 | 2.34045547 | 9 | 87.2 | 93.2 | 6.0 |
| LITPVLQAGKAEVTGY | -3.7838753 | 1.50502585 | 2 | -7.0743626 | 2.291182 | 4 | 93.2 | 99.3 | 6.0 |
| SDCKIQNGTSGIRF | -3.8010428 | 1.2551153 | 2 | -7.2157019 | 3.88920374 | 2 | 93.3 | 99.3 | 6.0 |

| | | | | | | | | | |
|---|---|---|---|---|---|---|---|---|---|
| VSQDPKDQLLLGPTY | -3.207122 | 1.05264359 | 4 | -4.673716 | 0.91356188 | 6 | 90.2 | 96.2 | 6.0 |
| VQYPVEHPDKF | -3.0676746 | 1.33357918 | 2 | -4.3526626 | 0.67503732 | 3 | 89.3 | 95.3 | 6.0 |
| VDDGLISLQVKQKGADF | -3.6092301 | 1.41102067 | 4 | -5.9503253 | 1.7284154 | 3 | 92.4 | 98.4 | 6.0 |
| CLNDDDETEVLKEDIIQGF | -2.02345 | 0.72337756 | 8 | -2.6471827 | 0.6181273 | 5 | 80.3 | 86.2 | 6.0 |
| VLTKENFDEVVNDADIILVEF | -2.5247778 | 0.10834283 | 3 | -3.3660474 | 0.24098377 | 3 | 85.2 | 91.2 | 6.0 |
| NKLDAENAIVH | -2.9891306 | 0.32559303 | 3 | -4.18129 | 1.55948819 | 4 | 88.8 | 94.8 | 6.0 |
| SKQRAECLEELGCLVESY | -2.2439175 | 0.09264999 | 2 | -2.9468278 | 0.08288277 | 2 | 82.6 | 88.5 | 6.0 |
| SGFPFEKGSVQY | -3.2864942 | 0.98955822 | 4 | -4.8427176 | 1.50300196 | 4 | 90.7 | 96.6 | 5.9 |
| LINVKQENTQTEQ | -3.33569 | 0.98389132 | 5 | -4.9669867 | 1.05672975 | 3 | 91.0 | 96.9 | 5.9 |
| IISTDPAHNISDAFDQKF | -3.4520972 | 1.62110486 | 5 | -5.307418 | 1.06253119 | 3 | 91.6 | 97.5 | 5.9 |
| DEYGSKKSDYIKLY | -3.7344633 | 1.92398541 | 2 | -6.5175709 | 1.11803966 | 2 | 93.0 | 98.9 | 5.9 |
| ILLFSKDEDISL | -3.3883644 | 1.52391983 | 2 | -5.111328 | 0.80581047 | 2 | 91.3 | 97.2 | 5.9 |
| SKVVPPLDEDGRSLL | -3.4555905 | 0.98188159 | 4 | -5.3102655 | 2.51415286 | 2 | 91.6 | 97.5 | 5.9 |
| VDATEDPWKTNY | -3.6050345 | 0.89006195 | 4 | -5.8511221 | 2.26925747 | 3 | 92.4 | 98.3 | 5.9 |
| VELQKEEAQKL | -3.9791775 | 1.77560007 | 3 | -10.375074 | 0.36385818 | 2 | 94.0 | 99.9 | 5.9 |
| LTFSPSEVKSL | -3.1521065 | 0.91589019 | 5 | -4.499762 | 0.63544371 | 4 | 89.9 | 95.8 | 5.9 |
| GAVTNVKVIRDFNTN | -3.7249185 | 1.38883492 | 8 | -6.4213902 | 1.27355295 | 2 | 93.0 | 98.8 | 5.9 |
| AIIKFPLTTESAM | -3.6678738 | 1.13604084 | 4 | -6.0956356 | 1.1264142 | 8 | 92.7 | 98.6 | 5.9 |
| TEKITPLEIEVLEETVQTMDTS | -2.8333451 | 0.65163938 | 4 | -3.8515602 | 1.4683265 | 2 | 87.7 | 93.5 | 5.8 |
| ISQESFDVDETDSGAGLKW | -2.1834622 | 0.40275324 | 2 | -2.8443879 | 0.19655242 | 5 | 82.0 | 87.8 | 5.8 |
| SDYVGSGPPKGTGL | -3.4668173 | 1.03407604 | 7 | -5.2934658 | 0.54404578 | 2 | 91.7 | 97.5 | 5.8 |
| VLGANNQETVKY | -2.6027449 | 2.17400518 | 2 | -3.459978 | 0.00550063 | 2 | 85.9 | 91.7 | 5.8 |
| QQFKDSDLSHVQN | -3.7011279 | 0.7167702 | 2 | -6.1876778 | 1.48530506 | 3 | 92.9 | 98.6 | 5.8 |
| AQNVLSKADVIQATGDAICIF | -2.3972627 | 0.83262908 | 7 | -3.1430999 | 0.11754929 | 3 | 84.0 | 89.8 | 5.8 |
| GPRAEEYEFLTPVEEAPKGM | -2.4071843 | 0.16087813 | 4 | -3.1567478 | 0.63844618 | 3 | 84.1 | 89.9 | 5.8 |
| NIRKPNEGADGQW | -3.4825728 | 1.49332776 | 10 | -5.3144459 | 0.93973691 | 5 | 91.8 | 97.5 | 5.8 |
| VFTKEDLTEIRDML | -3.3243939 | 1.23196096 | 4 | -4.861903 | 1.0261814 | 3 | 90.9 | 96.7 | 5.8 |
| QLDNNFEVKSLIFDQSGTY | -3.0855668 | 0.83299135 | 6 | -4.2980832 | 0.41120303 | 3 | 89.5 | 95.2 | 5.7 |
| VGNLPPDITEEEMRKLF | -2.4812591 | 0.02358216 | 2 | -3.2521408 | 1.02139461 | 2 | 84.8 | 90.5 | 5.7 |
| TRQVVDCQLADVNNIGKY | -2.1474131 | 0.51691238 | 2 | -2.7745469 | 0.48081576 | 2 | 81.6 | 87.2 | 5.7 |
| ILDKIIEEDDAYDFSTDYV | -1.9629228 | 0.46549587 | 5 | -2.5273297 | 0.08684269 | 4 | 79.6 | 85.2 | 5.6 |
| DAGAGIALNDHFVKLISWY | -1.166425 | 1.40513202 | 6 | -1.5696107 | 0.7780357 | 9 | 69.2 | 74.8 | 5.6 |
| SGPAGPILSLNPQEDVEFQKEVAQVR | -2.2950795 | 0.27791015 | 2 | -2.9690308 | 0.38307649 | 2 | 83.1 | 88.7 | 5.6 |
| IGANPLAVDLLEKML | -3.7576117 | 0.76330203 | 4 | -6.2650019 | 1.17508557 | 2 | 93.1 | 98.7 | 5.6 |
| APNILENKEGLELL | -3.5147061 | 0.99138145 | 3 | -5.3121944 | 0.84995057 | 4 | 92.0 | 97.5 | 5.6 |
| GLLFQESPEQKNW | -3.7337435 | 0.66182987 | 4 | -6.1253176 | 2.70210344 | 2 | 93.0 | 98.6 | 5.6 |
| SGETAKGDYPLEAVRM | -3.5011724 | 0.66504585 | 2 | -5.2639144 | 1.53128476 | 4 | 91.9 | 97.5 | 5.6 |
| VEEVSTGQECGVVLDKTCF | -2.4598732 | 0.31098744 | 4 | -3.1995476 | 0.22719862 | 3 | 84.6 | 90.2 | 5.6 |
| KMSVQPTVSLGGF | -2.9053364 | 4.07755949 | 5 | -3.9142814 | 2.44299219 | 4 | 88.2 | 93.8 | 5.6 |
| ESVAAKNLQEAEEW | -3.8831875 | 2.54353497 | 4 | -6.9226393 | 0.4688603 | 3 | 93.7 | 99.2 | 5.5 |
| ILLWDVDGGLTQIDKY | -3.4995907 | 0.2577713 | 2 | -5.2291704 | 1.81550365 | 2 | 91.9 | 97.4 | 5.5 |
| DTTLGGRKFDEVLVNHF | -3.424572 | 1.53693275 | 6 | -5.0092019 | 1.19697089 | 2 | 91.5 | 97.0 | 5.5 |
| RCGESGHLAKDCDLQEDACY | -3.141617 | 0.64783785 | 3 | -4.3504564 | 0.89328825 | 2 | 89.8 | 95.3 | 5.5 |
| LEGKDQLGGWFQSSLL | -3.7140267 | 0.82440594 | 6 | -5.9585787 | 1.22069141 | 2 | 92.9 | 98.4 | 5.5 |
| AALGVAVDGGKDSL | -3.6721302 | 0.70837154 | 4 | -5.779365 | 1.85345431 | 4 | 92.7 | 98.2 | 5.5 |
| LSINPQKDETLETEKAQY | -3.6363922 | 0.73415031 | 7 | -5.6286883 | 1.84557409 | 3 | 92.6 | 98.0 | 5.5 |
| TLTATGGVQSTASSKNASCY | -2.8350766 | 0.38147686 | 2 | -3.7692219 | 1.40959689 | 3 | 87.7 | 93.2 | 5.5 |
| VVTAEGLRPVKEVGMY | -3.4380003 | 1.1415296 | 4 | -5.0175965 | 1.30962903 | 5 | 91.6 | 97.0 | 5.5 |
| RITESEEVVSREVSGIKAAY | -2.6037305 | 0.52032014 | 4 | -3.3925825 | 0.08065114 | 2 | 85.9 | 91.3 | 5.4 |
| DQVVKTIGLREVW | -3.8360199 | 0.49991137 | 4 | -6.4679798 | 2.04998423 | 4 | 93.5 | 98.9 | 5.4 |
| KGLVYETSVLDPDEGIRF | -2.9275442 | 0.21262939 | 6 | -3.9175389 | 1.02971975 | 2 | 88.4 | 93.8 | 5.4 |
| ALAICSQCSDISTKQAAF | -2.5703577 | 0.81736483 | 5 | -3.3369659 | 0.93888864 | 3 | 85.6 | 91.0 | 5.4 |
| KIPVIENLGATLDQF | -3.4735598 | 2.41878632 | 2 | -5.0865612 | 0.06480905 | 2 | 91.7 | 97.1 | 5.4 |
| KLLSDFLDSEVSEL | -3.7310068 | 1.55216316 | 4 | -5.9323565 | 1.89141725 | 2 | 93.0 | 98.4 | 5.4 |
| DLTAKELTEEKESAF | -3.2742494 | 0.36898971 | 2 | -4.5921994 | 0.13941017 | 2 | 90.6 | 96.0 | 5.4 |
| VIAKMDATANDVPSDRY | -3.7679666 | 0.65546288 | 3 | -6.0777883 | 0.58633585 | 3 | 93.2 | 98.5 | 5.4 |
| AWNNEVKQGL | -3.4188618 | 1.68505671 | 3 | -4.9299273 | 1.48028269 | 2 | 91.4 | 96.8 | 5.4 |
| GSGDQEAWQKGVLF | -2.8642259 | 0.81760843 | 3 | -3.7976878 | 1.29671687 | 3 | 87.9 | 93.3 | 5.4 |
| SRKEEQEVQATLESEEVDLNAGLHGNW | -1.7322567 | 0.02304732 | 2 | -2.2086444 | 0.1424656 | 3 | 76.9 | 82.2 | 5.3 |
| VAPGNAGTACSEKISNTAISISDHTALAQF | -1.1299334 | 0.576003 | 4 | -1.5077218 | 0.23287679 | 7 | 68.6 | 74.0 | 5.3 |
| AKGHYTEGAELVDSVL | -3.3967144 | 1.19090312 | 5 | -4.8518656 | 1.29650959 | 5 | 91.3 | 96.7 | 5.3 |
| SKTPELNLDQF | -3.8628084 | 0.59612869 | 2 | -6.4547669 | 0.21703288 | 2 | 93.6 | 98.9 | 5.3 |
| RAEAVESAQAGDKCDF | -3.9399344 | 0.77789807 | 3 | -6.9247219 | 1.03048431 | 4 | 93.9 | 99.2 | 5.3 |
| GGTSELSSEGTQHSYSEEEKY | -3.9670172 | 1.04154606 | 2 | -7.116675 | 4.02924893 | 2 | 94.0 | 99.3 | 5.3 |
| DLSDKSINPLGGF | -2.7545135 | 2.35482465 | 3 | -3.5992414 | 0.09911577 | 2 | 87.1 | 92.4 | 5.3 |
| VLRNMVDPKDIDDDLEGEVTEECGKF | -3.9308083 | 0.64408055 | 2 | -6.809786 | 2.77781121 | 3 | 93.8 | 99.1 | 5.3 |
| KFDTLCDLYDTL | -3.7071563 | 1.02945725 | 3 | -5.7109235 | 2.15173505 | 2 | 92.9 | 98.1 | 5.2 |
| IVVIGHVDSGKSTTT | -3.496822 | 1.0971366 | 3 | -5.0524163 | 0.00101557 | 2 | 91.9 | 97.1 | 5.2 |
| ESLQKERVEAGDVIY | -3.7390083 | 0.43151367 | 4 | -5.8015179 | 1.25664614 | 5 | 93.0 | 98.2 | 5.2 |
| TEKTPISEHAVF | -3.7202994 | 1.74487134 | 2 | -5.7169625 | 0.20196173 | 2 | 92.9 | 98.1 | 5.2 |
| TVNFLEAKEGDLHRIEIPF | -2.0547685 | 0.48075594 | 2 | -2.5934362 | 0.31782484 | 2 | 80.6 | 85.8 | 5.2 |
| LLGSTAEEKAIVQQW | -3.3213015 | 0.70404409 | 3 | -4.6187322 | 0.75600789 | 3 | 90.9 | 96.1 | 5.2 |

| | | | | | | | | | |
|---|---|---|---|---|---|---|---|---|---|
| LAEVACGDDRKQTIDNSQGAYQEAF | -2.9705102 | 0.75206726 | 4 | -3.9343778 | 0.63840756 | 3 | 88.7 | 93.9 | 5.2 |
| KGTVGEPTYDAEFQHF | -3.7060671 | 1.04210573 | 7 | -5.6527304 | 1.56367796 | 5 | 92.9 | 98.1 | 5.2 |
| ALDSPKGCGTVL | -3.7262422 | 0.94760683 | 5 | -5.7176703 | 1.49972069 | 8 | 93.0 | 98.1 | 5.2 |
| LANLTQSQIALNEKLVNL | -4.092158 | 2.51908714 | 5 | -8.0232432 | 0.82624088 | 5 | 94.5 | 99.6 | 5.2 |
| HFKVDNDENEHQLSL | -3.423017 | 1.49518131 | 6 | -4.8387096 | 1.84879205 | 3 | 91.5 | 96.6 | 5.2 |
| AKIINNTENLVREL | -3.5028627 | 0.65467671 | 3 | -5.0290781 | 1.8024162 | 2 | 91.9 | 97.0 | 5.1 |
| SFVDKDLLEPGCSVL | -3.8222051 | 0.8918653 | 3 | -6.0574236 | 0.50900943 | 2 | 93.4 | 98.5 | 5.1 |
| SIGKIGGAQNRSY | -3.6449871 | 1.0553585 | 7 | -5.417764 | 0.54022224 | 4 | 92.6 | 97.7 | 5.1 |
| VDIAKSQDAEVGDGTTSVTLL | -3.2121956 | 0.4635245 | 4 | -4.3664176 | 1.1927632 | 2 | 90.3 | 95.4 | 5.1 |
| ADPVSAQHAKL | -3.7785487 | 1.16906613 | 5 | -5.873141 | 1.44879944 | 5 | 93.2 | 98.3 | 5.1 |
| AVEIVGGATRIPAVKE | -3.907358 | 0.57796305 | 4 | -6.4373136 | 2.0530578 | 6 | 93.8 | 98.9 | 5.1 |
| VSNIDGTHIAKTL | -3.1016368 | 2.70432167 | 7 | -4.1450843 | 0.54050963 | 6 | 89.6 | 94.7 | 5.1 |
| IGGLSFETTDDSLREHFEKW | -1.9426696 | 0.29883755 | 3 | -2.4391078 | 0.44393949 | 2 | 79.4 | 84.4 | 5.1 |
| SAAILEYLTAEVLELAGNASKDL | -2.3203779 | 0.03530811 | 2 | -2.9271748 | 0.09354533 | 2 | 83.3 | 88.4 | 5.1 |
| QLAVEAESEQKW | -3.1656713 | 0.14862467 | 2 | -4.2571224 | 0.54004712 | 3 | 90.0 | 95.0 | 5.1 |
| APINANAIKAEC | -4.0038579 | 0.83293542 | 2 | -6.9279345 | 2.03439139 | 2 | 94.1 | 99.2 | 5.1 |
| ASLLDYDTVAGADKF | -1.4869612 | 0.06238304 | 2 | -1.8888055 | 0.3344261 | 6 | 73.7 | 78.7 | 5.0 |
| QNKEHVIEAL | -3.5424119 | 1.01572194 | 2 | -5.0779854 | 2.41138486 | 2 | 92.1 | 97.1 | 5.0 |
| SGWYDADLSPAGHEEAKRGGQAL | -2.6766357 | 0.4526062 | 3 | -3.4275569 | 0.23743048 | 2 | 86.5 | 91.5 | 5.0 |
| ALELLFDQLHEGAKAL | -1.993788 | 0.4051099 | 6 | -2.4953592 | 0.78829165 | 4 | 79.9 | 84.9 | 5.0 |
| KHGDEIITSTTSNY | -4.1676888 | 0.04454825 | 2 | -8.4916521 | 6.09615633 | 2 | 94.7 | 99.7 | 5.0 |
| GLVFDDVVGIVEIINSKDVKVQ | -3.9235998 | 0.09331285 | 3 | -6.3414658 | 0.64967592 | 3 | 93.8 | 98.8 | 5.0 |
| VYSKDQLQTF | -4.1714407 | 1.65911415 | 2 | -8.3455031 | 1.4058128 | 3 | 94.7 | 99.7 | 5.0 |
| GQISEVVVVKDRETQ | -3.8900127 | 1.63168817 | 5 | -6.1694297 | 2.59642287 | 5 | 93.7 | 98.6 | 4.9 |
| KQLNDLWDQIEQAHL | -3.8682346 | 1.58689314 | 2 | -6.0754569 | 2.11338897 | 3 | 93.6 | 98.5 | 4.9 |
| NQLKPGLQY | -4.0820282 | 0.89911587 | 6 | -7.2953358 | 2.51043532 | 2 | 94.4 | 99.4 | 4.9 |
| TIVKDSEADRF | -3.6264349 | 0.5479821 | 2 | -5.2483267 | 0.89575335 | 2 | 92.5 | 97.4 | 4.9 |
| TLKGGAAVDPDSGLEHSAH | -3.7956352 | 1.12371332 | 9 | -5.7726483 | 3.11789386 | 5 | 93.3 | 98.2 | 4.9 |
| EIENNPTVKASGY | -3.9595171 | 0.41065055 | 2 | -6.4624871 | 2.32385191 | 4 | 94.0 | 98.9 | 4.9 |
| FVEWIPNNVKTAVC | -4.0894242 | 1.32189049 | 3 | -7.294783 | 1.63036592 | 6 | 94.5 | 99.4 | 4.9 |
| KGILGYTEHQVVSSDFNSDTHSSTF | -2.9997144 | 0.5693151 | 6 | -3.9124041 | 0.54717001 | 2 | 88.9 | 93.8 | 4.9 |
| EAGISKNGQTREHALL | -3.7070684 | 1.1246575 | 6 | -5.4485879 | 1.49377582 | 4 | 92.9 | 97.8 | 4.9 |
| TGFIVEADTPGIQIGRKEL | -0.779044 | 0.29990881 | 2 | -1.0905987 | 1.37049339 | 6 | 63.2 | 68.0 | 4.9 |
| IAEPLEKGLAEDIENEVVQITW | -2.558817 | 1.15366996 | 3 | -3.2280641 | 0.08495324 | 2 | 85.5 | 90.4 | 4.9 |
| QGIAAKVGTGEPCCDW | -2.7398558 | 0.25276667 | 3 | -3.4912923 | 0.23347088 | 5 | 87.0 | 91.8 | 4.9 |
| SATLGLVDIVKGTNSY | -3.6778272 | 0.79977122 | 3 | -5.3431932 | 1.83076907 | 3 | 92.8 | 97.6 | 4.8 |
| DVNKPGCEVDDLKGGVAGGSIL | -4.1213356 | 1.32184349 | 6 | -7.3745061 | 2.57119279 | 6 | 94.6 | 99.4 | 4.8 |
| NDELQFLEKINKNCW | -3.7019126 | 0.78140086 | 5 | -5.4066967 | 0.20787986 | 3 | 92.9 | 97.7 | 4.8 |
| AAAGGVSHDELLDLAKF | -3.1401246 | 0.32810767 | 2 | -4.1417702 | 0.31104605 | 2 | 89.8 | 94.6 | 4.8 |
| VLDKLGDDEVRTDL | -2.8484746 | 0.10157308 | 2 | -3.6520497 | 0.13455832 | 2 | 87.8 | 92.6 | 4.8 |
| HTVTTTDDPVIRKL | -4.0430982 | 1.25164467 | 6 | -6.7715479 | 2.76666533 | 3 | 94.3 | 99.1 | 4.8 |
| SIDISSKQVENAGAIGPSRF | -4.2430095 | 1.01377115 | 3 | -8.9309988 | 2.06957101 | 4 | 95.0 | 99.8 | 4.8 |
| EQWGKLTDCVVM | -3.5729449 | 0.54887007 | 3 | -5.0348684 | 2.13728251 | 6 | 92.2 | 97.0 | 4.8 |
| APVISAEKAYHEQLSVAEITNACF | -2.9563501 | 0.48755236 | 4 | -3.8162544 | 0.53917488 | 3 | 88.6 | 93.4 | 4.8 |
| KLSGGDDDAAGQFFPEAAQVAY | -3.0716839 | 0.82024988 | 4 | -4.0078588 | 0.38224143 | 3 | 89.4 | 94.1 | 4.8 |
| GLVDIVKGTNSYY | -3.4950568 | 1.09972164 | 7 | -4.8388343 | 1.13552141 | 6 | 91.9 | 96.6 | 4.8 |
| LDIEHADGKRYF | -3.7862945 | 0.66043444 | 3 | -5.6202118 | 1.42352753 | 3 | 93.2 | 98.0 | 4.8 |
| AGKQLEDGRTL | -3.6666775 | 1.84176119 | 15 | -5.2646967 | 1.0834428 | 9 | 92.7 | 97.5 | 4.8 |
| LIGDAAKNQLTSNPENTVF | -3.7345977 | 0.7264069 | 2 | -5.452939 | 1.74522634 | 3 | 93.0 | 97.8 | 4.8 |
| QQVIEKTKSL | -3.913204 | 1.77009457 | 4 | -6.0637602 | 3.43421574 | 3 | 93.8 | 98.5 | 4.8 |
| GGGVVTIERSKS | -3.0924108 | 1.2031253 | 4 | -4.0335391 | 1.20300134 | 4 | 89.5 | 94.2 | 4.7 |
| SMMDVDHQIAKL | -3.4374916 | 0.87192629 | 3 | -4.6927232 | 1.32177579 | 2 | 91.5 | 96.3 | 4.7 |
| EVKSENLGQGDKPDYF | -4.172913 | 0.42888018 | 2 | -7.5318913 | 1.58034073 | 3 | 94.7 | 99.5 | 4.7 |
| FITTVKTAW | -4.0452247 | 1.68057576 | 9 | -6.6184695 | 2.58600265 | 6 | 94.3 | 99.0 | 4.7 |
| NLGTIETKSASVAPF | -4.1583416 | 0.10669518 | 2 | -7.3714651 | 3.66892132 | 2 | 94.7 | 99.4 | 4.7 |
| QVRDIENLKDASSF | -4.2747398 | 0.3744551 | 2 | -8.8805974 | 2.23346176 | 2 | 95.1 | 99.8 | 4.7 |
| IVSVDETIKNPR | -4.285801 | 1.3669868 | 3 | -9.1133483 | 1.20552653 | 2 | 95.1 | 99.8 | 4.7 |
| FLSSGLIDKVDNF | -3.7874924 | 1.47555803 | 8 | -5.57257 | 1.14109445 | 6 | 93.2 | 97.9 | 4.7 |
| KTTIPEEEEEEEAAGVVVEEELF | -2.9533524 | 0.16930991 | 7 | -3.7878359 | 0.4534689 | 4 | 88.6 | 93.2 | 4.7 |
| TKDIDIHEVRIGW | -4.1030085 | 1.62857786 | 4 | -6.904034 | 0.01221738 | 2 | 94.5 | 99.2 | 4.7 |
| IGGLNTETNEKALEAVF | -3.8981311 | 1.12053944 | 3 | -5.9177764 | 1.49797288 | 5 | 93.7 | 98.4 | 4.7 |
| SMDLRTKSTGGAPTF | -3.2886398 | 0.70697969 | 8 | -4.3655209 | 1.31709352 | 5 | 90.7 | 95.4 | 4.7 |
| TIAVASLGKGVACNPACF | -3.6785622 | 1.02362936 | 5 | -5.2336849 | 4.20702533 | 3 | 92.8 | 97.4 | 4.7 |
| SLSHNPEQKGVPTGF | -3.6313877 | 0.43060613 | 4 | -5.1040233 | 0.57501017 | 3 | 92.5 | 97.2 | 4.6 |
| RYLSEVASGDNKQTTVSNSQQAY | -2.8181098 | 0.56894189 | 2 | -3.5653151 | 0.02982688 | 2 | 87.6 | 92.2 | 4.6 |
| QAGQCGNQIGAKF | -3.4195051 | 1.01218034 | 16 | -4.6135306 | 0.81438065 | 12 | 91.5 | 96.1 | 4.6 |
| KEGTDSSQGIPQLVSNISACQVIAEAVRTTL | -2.3734582 | 0.45463583 | 6 | -2.9356452 | 0.37215733 | 4 | 83.8 | 88.4 | 4.6 |
| SAMVSMVTKDNPGVVTCLDEARHGFESGDFVSF | -1.4017288 | 0.08450334 | 2 | -1.7538409 | 0.0422804 | 2 | 72.5 | 77.1 | 4.6 |
| GNVAGDSKNDPPMEAAGF | -3.4787831 | 0.92603469 | 3 | -4.7129054 | 1.72098246 | 4 | 91.8 | 96.3 | 4.6 |
| QQQKTSVPVLDAELF | -3.6107118 | 1.22793962 | 4 | -5.0085147 | 1.82196581 | 3 | 92.4 | 97.0 | 4.6 |
| SAALNAPIEKTDFGIF | -3.2258739 | 0.84057192 | 4 | -4.2162281 | 0.16176093 | 2 | 90.3 | 94.9 | 4.6 |
| IVFDGDVDPEWVENLNSVLDDNKLL | -3.3927021 | 0.16118994 | 2 | -4.530575 | 0.5281142 | 2 | 91.3 | 95.9 | 4.5 |

| | | | | | | | | | |
|---|---|---|---|---|---|---|---|---|---|
| LPKDTSPGSAYQEGGGLY | -3.9062791 | 1.04667005 | 10 | -5.8461209 | 2.33197597 | 8 | 93.7 | 98.3 | 4.5 |
| DLSELPKFEKNFY | -4.0511609 | 2.21242992 | 5 | -6.4166052 | 1.37669116 | 3 | 94.3 | 98.8 | 4.5 |
| VRNLANTVTEEILEKAF | -4.0153606 | 1.28862906 | 4 | -6.2511738 | 0.03263481 | 2 | 94.2 | 98.7 | 4.5 |
| EQQVPVNQVFGQDEMIDVIGVTKGKGY | -3.4971683 | 0.74302487 | 2 | -4.7395725 | 0.74198509 | 2 | 91.9 | 96.4 | 4.5 |
| FVEKNPTIVNFPITNVDLREY | 3.86030935 | 0.9187715 | 2 | 3.02205374 | 1.17159062 | 2 | 6.4 | 11.0 | 4.5 |
| VILANNCPALRKSEIEYY | -4.1653009 | 0.47538554 | 3 | -7.0164382 | 4.17100516 | 2 | 94.7 | 99.2 | 4.5 |
| VSGACDASAKLW | -3.7200774 | 0.60287092 | 2 | -5.2597332 | 1.05330821 | 5 | 92.9 | 97.5 | 4.5 |
| TGEISPGMIKDCGATW | -1.9696766 | 0.21089483 | 3 | -2.409776 | 0.28648553 | 4 | 79.7 | 84.2 | 4.5 |
| EILTPNSIPKGF | -4.0050347 | 2.16768019 | 6 | -6.1671082 | 0.61240622 | 3 | 94.1 | 98.6 | 4.5 |
| GIKGGAAGGGY | -4.1436776 | 0.025604 | 2 | -6.8368867 | 2.26674216 | 2 | 94.6 | 99.1 | 4.5 |
| YLTRVEVTEFEDIKSGY | -2.6447516 | 0.37746091 | 5 | -3.2860814 | 1.97782271 | 4 | 86.2 | 90.7 | 4.5 |
| DSGFGGGAGVETGGKLL | -3.8940881 | 0.90457643 | 6 | -5.7478203 | 1.0964139 | 3 | 93.7 | 98.2 | 4.5 |
| GIGGLDKEFSDIF | -3.918358 | 0.67967092 | 3 | -5.8257584 | 0.67531615 | 3 | 93.8 | 98.3 | 4.5 |
| GSRQIILEKEETEEL | -3.9276067 | 0.22292644 | 4 | -5.856675 | 0.5234078 | 3 | 93.8 | 98.3 | 4.5 |
| IMIQQTEITCPKVNQF | -2.9203801 | 0.74315962 | 3 | -3.6857306 | 1.02769933 | 2 | 88.3 | 92.8 | 4.5 |
| LIANATNPESKVF | -3.8049299 | 1.02645544 | 9 | -5.4554772 | 1.72310325 | 8 | 93.3 | 97.8 | 4.4 |
| SDCKIQNGAQGIRF | -4.2549775 | 1.7021099 | 3 | -7.5272555 | 2.09541216 | 6 | 95.0 | 99.5 | 4.4 |
| KFVDGLMIHSGDPVNY | -3.4775726 | 2.91964118 | 4 | -4.6600663 | 0.86306748 | 2 | 91.8 | 96.2 | 4.4 |
| SVAKGSDEPPVFLEIHY | -3.3168982 | 0.49485622 | 6 | -4.3363284 | 1.58332752 | 3 | 90.9 | 95.3 | 4.4 |
| CYETAEKITRTPQASTY | -2.4369443 | 0.5280879 | 2 | -2.9890635 | 0.16634104 | 2 | 84.4 | 88.8 | 4.4 |
| VAPEKADIIVSELL | -4.3241356 | 1.03473176 | 4 | -8.128441 | 2.59839583 | 2 | 95.2 | 99.6 | 4.4 |
| LASSDPLAQIAEDKPYAELW | -2.5773356 | 0.58131277 | 4 | -3.1749295 | 0.43076616 | 3 | 85.6 | 90.0 | 4.4 |
| NISNGGPAPEAITDKIF | -3.8002439 | 2.01394385 | 2 | -5.3800366 | 2.04987759 | 7 | 93.3 | 97.7 | 4.4 |
| KVIGIECSSISDY | -4.2478534 | 0.55799321 | 2 | -7.2597248 | 2.62798548 | 3 | 95.0 | 99.4 | 4.4 |
| TAIVDKIGF | -4.3884284 | 2.31267853 | 2 | -8.9166821 | 1.81709837 | 3 | 95.4 | 99.8 | 4.4 |
| YEKRMATEVAADALGEEWKGY | -3.7916726 | 2.23630423 | 4 | -5.3545501 | 0.92656099 | 2 | 93.3 | 97.6 | 4.3 |
| NEGLWEIENNPGVKF | -3.1169042 | 1.11588369 | 2 | -3.9726599 | 0.53461968 | 3 | 89.7 | 94.0 | 4.3 |
| GFDSISSVKFNPIETF | -3.8080972 | 0.9076459 | 7 | -5.3959699 | 1.35853636 | 3 | 93.3 | 97.7 | 4.3 |
| TEALGIDPNNIKTNAKLY | -4.1067396 | 0.45202635 | 4 | -6.4203507 | 2.84874103 | 2 | 94.5 | 98.8 | 4.3 |
| KVTQDELKEVFEDAAEIR | -3.9254515 | 0.1489798 | 2 | -5.722559 | 1.75309169 | 2 | 93.8 | 98.1 | 4.3 |
| EVLEGEVEKEAL | -3.6275675 | 0.6759925 | 2 | -4.9286001 | 1.11103423 | 5 | 92.5 | 96.8 | 4.3 |
| GYVDFESAEDLEKALEL | -3.9053031 | 2.89740323 | 10 | -5.6425703 | 4.13893006 | 9 | 93.7 | 98.0 | 4.3 |
| TNDWEDHLAVKHF | -3.8667035 | 0.93315543 | 6 | -5.5143345 | 0.36775929 | 2 | 93.6 | 97.9 | 4.3 |
| KTILPAAAQDVY | -4.362647 | 4.68222186 | 5 | -8.0847162 | 1.12713361 | 2 | 95.4 | 99.6 | 4.3 |
| DLIANIVHDGKPSEGSY | -4.2580551 | 1.17995165 | 5 | -7.1455211 | 2.6186608 | 4 | 95.0 | 99.3 | 4.3 |
| CGGDIYVPEDPKLKDGY | -3.4685602 | 1.2558472 | 6 | -4.57531 | 1.17716336 | 3 | 91.7 | 96.0 | 4.3 |
| STEIKKTEVLMENF | -3.5202394 | 0.12844969 | 3 | -4.6722282 | 1.36384941 | 4 | 92.0 | 96.2 | 4.2 |
| MTEPIDEYCVQQLKEF | -2.1811177 | 0.34314337 | 4 | -2.6399696 | 0.26385022 | 2 | 81.9 | 86.2 | 4.2 |
| LNKMTEAQEDGQSTSELIGQF | -2.7939352 | 0.27071639 | 5 | -3.4519837 | 0.04308888 | 2 | 87.4 | 91.6 | 4.2 |
| SWESSKDPAEQQGKGVAL | -4.4248827 | 1.45723397 | 5 | -8.6781281 | 1.82102081 | 2 | 95.6 | 99.8 | 4.2 |
| AQTEGINISEEALNHLGEIGTKTTL | -3.3435188 | 0.44749444 | 5 | -4.3195497 | 0.0676133 | 2 | 91.0 | 95.2 | 4.2 |
| AIIDPGDSDIIRSMPEQTGEK | -2.1578708 | 0.33128784 | 4 | -2.6040507 | 0.48370577 | 2 | 81.7 | 85.9 | 4.2 |
| QAQIQEQHVQIDVDVSKPDLTAALRDVRQQY | -2.709374 | 0.32456138 | 2 | -3.3224671 | 0.08529069 | 2 | 86.7 | 90.9 | 4.2 |
| ASISGSSASSTSSTPEVKPL | -4.2213114 | 0.03615195 | 2 | -6.7503846 | 2.1814665 | 4 | 94.9 | 99.1 | 4.2 |
| TDCFNCLPIAAIVDEKIF | -2.609671 | 0.22705647 | 5 | -3.1832558 | 0.15222354 | 3 | 85.9 | 90.1 | 4.2 |
| VYTIDDLIDKLSTIVN | -3.2022906 | 0.54429458 | 3 | -4.0640456 | 0.64461743 | 2 | 90.2 | 94.4 | 4.2 |
| IKTEDPDLPAFY | -4.2916714 | 0.88230703 | 5 | -7.1344017 | 0.12842966 | 3 | 95.1 | 99.3 | 4.2 |
| TLCKEGCEAIVDTGTSL | -3.537358 | 0.42280203 | 3 | -4.6673917 | 0.98359709 | 2 | 92.1 | 96.2 | 4.1 |
| SADVDIKELAVETKNF | -3.1568038 | 0.42832989 | 3 | -3.9823719 | 0.7056246 | 2 | 89.9 | 94.0 | 4.1 |
| DLSDKSINPLGGFVHY | -3.7766873 | 0.9671761 | 6 | -5.1857903 | 1.45249896 | 4 | 93.2 | 97.3 | 4.1 |
| IVKNIDDGTSDRPY | -2.9118221 | 0.91396307 | 2 | -3.6031688 | 1.58447652 | 2 | 88.3 | 92.4 | 4.1 |
| MVGLDAAGKTTILY | -0.8297507 | 0.5031166 | 9 | -1.0954618 | 0.44921664 | 9 | 64.0 | 68.1 | 4.1 |
| FLTDSNNIKEVL | -4.2325513 | 1.32679068 | 5 | -6.7362661 | 0.54807155 | 2 | 94.9 | 99.1 | 4.1 |
| LVSVLSKADF | -3.9786888 | 2.21209839 | 2 | -5.7336433 | 1.42696666 | 4 | 94.0 | 98.2 | 4.1 |
| VDIAKSQDAEVGDGTTSVTL | -4.2832536 | 1.80604637 | 5 | -7.0076422 | 2.02865246 | 7 | 95.1 | 99.2 | 4.1 |
| SLQESGLKVNQPASF | -3.4798054 | 0.95393078 | 10 | -4.5410799 | 2.19535619 | 7 | 91.8 | 95.9 | 4.1 |
| KLITEDVQGKN | -4.1693341 | 1.58923745 | 11 | -6.3930801 | 1.14117626 | 7 | 94.7 | 98.8 | 4.1 |
| KGFDQTINLILDESHERVF | -3.1473706 | 0.72173188 | 3 | -3.9523858 | 0.616127 | 2 | 89.9 | 93.9 | 4.1 |
| QNIQDGSLDLNAAESGVQHKPSAPQ | -2.9600903 | 0.65669968 | 7 | -3.6631654 | 0.54545721 | 3 | 88.6 | 92.7 | 4.1 |
| KTDLRFQSAAIGALQEASEAY | -1.4606659 | 0.28140457 | 8 | -1.7772693 | 0.18390578 | 4 | 73.4 | 77.4 | 4.1 |
| FAGDTIPKSPF | -3.5784557 | 0.65366655 | 9 | -4.710924 | 0.06918397 | 2 | 92.3 | 96.3 | 4.0 |
| KDLFDPIIEDRHGGY | -4.2502764 | 0.95177675 | 7 | -6.6856921 | 1.08647457 | 6 | 95.0 | 99.0 | 4.0 |
| DLSKIGEEQSAEDAEDGPPELLF | -3.2840065 | 0.6305597 | 7 | -4.162556 | 1.29006412 | 7 | 90.7 | 94.7 | 4.0 |
| LASQKTDAEATDTEATET | -3.5564347 | 1.71574474 | 2 | -4.653298 | 0.91712971 | 3 | 92.2 | 96.2 | 4.0 |
| SLLRGGSDDSSKDPIDVNYEKL | -4.0201374 | 1.27675434 | 8 | -5.7733401 | 0.72335051 | 3 | 94.2 | 98.2 | 4.0 |
| AVNDFELARADFQKVL | -4.3057608 | 1.31244169 | 7 | -6.9333383 | 1.66143304 | 4 | 95.2 | 99.2 | 4.0 |
| LDAGLARTTTGNKVF | -2.969037 | 0.54926491 | 11 | -3.6604837 | 0.16274881 | 2 | 88.7 | 92.7 | 4.0 |
| FGAFGEIENIELPMDTKTNERRGF | -1.7164964 | 0.30363449 | 3 | -2.0597759 | 0.28728779 | 2 | 76.7 | 80.7 | 4.0 |
| GILADATEQVGQHKDAY | -4.2149053 | 1.32543121 | 6 | -6.4509356 | 1.90164484 | 7 | 94.9 | 98.9 | 4.0 |
| GEVGKTTGIPIHVF | -3.5791355 | 1.21576845 | 3 | -4.6833248 | 1.18953131 | 3 | 92.3 | 96.3 | 4.0 |
| AWKGETDEEYLW | -4.1861501 | 1.72298236 | 3 | -6.3156501 | 0.75871986 | 3 | 94.8 | 98.8 | 4.0 |
| SAEIIYHLFDAFTKY | -3.8873048 | 0.79815832 | 3 | -5.3653414 | 0.33159136 | 2 | 93.7 | 97.6 | 4.0 |

| | | | | | | | | | |
|---|---|---|---|---|---|---|---|---|---|
| IAAPSGSAADKVVIEACDELGIILAHTNL | -1.7439179 | 0.2700189 | 9 | -2.0888919 | 0.34313706 | 4 | 77.0 | 81.0 | 4.0 |
| KRILEDQEENPLPAAL | -3.6994734 | 1.68825466 | 2 | -4.919576 | 1.54796372 | 4 | 92.9 | 96.8 | 3.9 |
| EPLEPVKDTDIQGFL | -4.1500036 | 0.5891407 | 4 | -6.1533657 | 1.54885316 | 4 | 94.7 | 98.6 | 3.9 |
| GCAADVEAVQTGLDLLEILRQEKGGSRGEEVGEL | -2.0645329 | 0.71991853 | 6 | -2.4628389 | 0.30086282 | 2 | 80.7 | 84.6 | 3.9 |
| VVHDGFEGLAKGQVQEVGWHDVAGW | -3.0114644 | 0.11995153 | 2 | -3.7109619 | 0.4366998 | 2 | 89.0 | 92.9 | 3.9 |
| FKGPELLVDYQMY | -3.6190158 | 0.00802178 | 2 | -4.7454234 | 0.40931963 | 4 | 92.5 | 96.4 | 3.9 |
| TLKGGAAVDPDSGLEHSAH | -3.4121781 | 0.9976581 | 5 | -4.3559109 | 1.53782577 | 3 | 91.4 | 95.3 | 3.9 |
| TSKIPALAVEMPGSADISGL | -1.9371553 | 0.4874466 | 4 | -2.3106418 | 0.12711064 | 5 | 79.3 | 83.2 | 3.9 |
| STSISKQETELSPEMISSGSW | -2.7299865 | 0.64805855 | 2 | -3.3075675 | 0.38213152 | 4 | 86.9 | 90.8 | 3.9 |
| KASTNIQDTEGNTPL | -4.1107885 | 0.5870838 | 6 | -5.985033 | 1.22024318 | 3 | 94.5 | 98.4 | 3.9 |
| LEYESSFSNEEAQKAQAL | -4.2413479 | 1.53484991 | 3 | -6.4724946 | 1.94538456 | 2 | 95.0 | 98.9 | 3.9 |
| VPEVSALDQEIIEVDPDTKEML | -3.2657742 | 0.16397741 | 2 | -4.0996067 | 0.52666756 | 5 | 90.6 | 94.5 | 3.9 |
| DLTSKPPGTTEWE | -4.0877772 | 1.55460656 | 2 | -5.8958606 | 1.86168764 | 4 | 94.4 | 98.3 | 3.9 |
| MNVAEVDKVTGRF | -4.0438197 | 0.84126184 | 4 | -5.754462 | 1.07296453 | 5 | 94.3 | 98.2 | 3.9 |
| QAQKDELILEGNDIELVSNSAAL | -2.3081702 | 0.77927189 | 3 | -2.7537548 | 2.39140501 | 2 | 83.2 | 87.1 | 3.9 |
| KETIDNNSVSSPL | -3.3710032 | 0.290947 | 2 | -4.270282 | 1.21991796 | 2 | 91.2 | 95.1 | 3.9 |
| QKGQETSTNPIASIF | -3.6645737 | 0.76934162 | 3 | -4.8178677 | 1.58549822 | 2 | 92.7 | 96.6 | 3.9 |
| AAEIPNIKPDILIIESTY | -4.2276706 | 1.97946539 | 4 | -6.3786895 | 1.06289701 | 5 | 94.9 | 98.8 | 3.9 |
| SVIKIFPDLSSNDMLLF | -2.270728 | 0.50675034 | 2 | -2.7042509 | 0.24077933 | 2 | 82.8 | 86.7 | 3.9 |
| TSIKQLSSEELEQF | -3.7032956 | 0.62801912 | 3 | -4.878961 | 0.73170203 | 4 | 92.9 | 96.7 | 3.8 |
| VFGDENGTVSLVDTKSTSCVL | -3.0071988 | 0.34609236 | 5 | -3.6830942 | 0.39904013 | 5 | 88.9 | 92.8 | 3.8 |
| IDDHFLFDKPVSPL | -3.9745215 | 0.60894828 | 3 | -5.5133095 | 1.27901549 | 5 | 94.0 | 97.9 | 3.8 |
| AKVDQALHTQTDADPAEEY | -4.1095538 | 0.67503317 | 4 | -5.9006029 | 1.08181358 | 5 | 94.5 | 98.4 | 3.8 |
| IVMAADAEPLEIILHLPLLCEDKNVPY | -2.8061751 | 0.54254486 | 3 | -3.3945754 | 0.1622545 | 2 | 87.5 | 91.3 | 3.8 |
| SATLDAGKF | -4.1030666 | 0.76734385 | 2 | -5.875582 | 1.23237551 | 2 | 94.5 | 98.3 | 3.8 |
| ALNNFDKQIIVDPLSF | -3.9319413 | 2.20717093 | 2 | -5.3923216 | 1.28883047 | 4 | 93.9 | 97.7 | 3.8 |
| FDENPYFENKVL | -3.9638672 | 0.25846153 | 2 | -5.4690085 | 1.07334757 | 3 | 94.0 | 97.8 | 3.8 |
| AMIIDKLEEDISSSM | -4.2430075 | 0.81634725 | 4 | -6.3579372 | 1.74247024 | 2 | 95.0 | 98.8 | 3.8 |
| TAIVDKIGFY | -4.4770809 | 1.86777346 | 4 | -7.6680641 | 1.90773856 | 5 | 95.7 | 99.5 | 3.8 |
| ISVEEVHDDGTPTSKTF | -4.1409766 | 1.49953391 | 3 | -5.9832421 | 2.97796623 | 3 | 94.6 | 98.4 | 3.8 |
| TLNDLISHQDILSTIQKF | -2.347925 | 0.2394939 | 5 | -2.7927843 | 0.41406481 | 3 | 83.6 | 87.4 | 3.8 |
| SPTHEQQLISGSLDNIVKLW | -0.6041744 | 0.48934885 | 3 | -0.8379117 | 0.71699629 | 2 | 60.3 | 64.1 | 3.8 |
| AMEAVKQGSATVGL | -4.2364074 | 0.90866511 | 4 | -6.3084401 | 1.9512452 | 4 | 95.0 | 98.8 | 3.8 |
| SNVNLKEVNY | -3.9155658 | 0.72670033 | 2 | -5.3308122 | 1.28698231 | 3 | 93.8 | 97.6 | 3.8 |
| TKDIDIHEVRIGW | -4.0610453 | 1.58495321 | 3 | -5.718159 | 0.14648276 | 2 | 94.3 | 98.1 | 3.8 |
| KLVGPEGFVVTEAGF | -2.4810378 | 1.11844945 | 6 | -2.957149 | 0.52322991 | 4 | 84.8 | 88.6 | 3.8 |
| CAKSREAIDSPVSF | -4.1252629 | 0.11924106 | 3 | -5.9077145 | 1.22676656 | 2 | 94.6 | 98.4 | 3.8 |
| KMWDPHNDPNAQGDAF | -4.2799948 | 0.74220836 | 9 | -6.4429496 | 1.96256107 | 5 | 95.1 | 98.9 | 3.8 |
| DTPFLPKPLFF | -4.1575321 | 0.14933337 | 2 | -5.9909293 | 2.37050009 | 2 | 94.7 | 98.5 | 3.8 |
| EYTLGEEKF | -4.3301151 | 1.48847558 | 2 | -6.6606181 | 0.48061152 | 2 | 95.3 | 99.0 | 3.8 |
| RSGQAKETIPLQETSLY | -4.2163042 | 1.958533 | 4 | -6.1925284 | 0.18626585 | 2 | 94.9 | 98.7 | 3.8 |
| KSENLDNPIQTVSLGQSLF | -2.4392695 | 0.28973301 | 5 | -2.8997938 | 0.43854526 | 3 | 84.4 | 88.2 | 3.8 |
| SGRVAVEEVDEEGKF | -3.6169433 | 0.98832602 | 4 | -4.6626603 | 1.58004056 | 2 | 92.5 | 96.2 | 3.7 |
| VYDDANKKWVPAGGSTGF | -4.5243375 | 0.53269833 | 4 | -7.8191851 | 1.96838436 | 3 | 95.8 | 99.6 | 3.7 |
| GKRPAEDMEEEQAF | -4.2666783 | 0.83062233 | 3 | -6.3382593 | 0.82193744 | 5 | 95.1 | 98.8 | 3.7 |
| AIEAIKLGSTAIGIQTSEGVCL | -3.1090511 | 0.27509768 | 3 | -3.8061515 | 0.51335469 | 4 | 89.6 | 93.3 | 3.7 |
| DVDGGLTQIDKY | -4.3944161 | 1.34460846 | 2 | -6.9085841 | 4.20464912 | 2 | 95.5 | 99.2 | 3.7 |
| FSDDPNVTKTL | -4.306131 | 1.25997889 | 2 | -6.4879001 | 0.79124965 | 2 | 95.2 | 98.9 | 3.7 |
| TAKELTEEKESAFEFLSSA | -3.745947 | 0.62678266 | 3 | -4.9044662 | 0.73756742 | 2 | 93.1 | 96.8 | 3.7 |
| NSGKVDIVAINDPFIDLNYM(15.9949)VY | -2.8191018 | 0.54825605 | 3 | -3.3899994 | 0.68717781 | 6 | 87.6 | 91.3 | 3.7 |
| ASVPDGFLSELTQQLAQATGKPPQY | -3.6584186 | 0.7314506 | 6 | -4.7257302 | 0.35292127 | 2 | 92.7 | 96.4 | 3.7 |
| KEDVGDEGEEEKEFISY | -3.9877838 | 1.80157509 | 3 | -5.4514289 | 1.76689641 | 3 | 94.1 | 97.8 | 3.7 |
| MQASEDLLKEHY | -4.3766341 | 2.01617559 | 2 | -6.7765503 | 1.43124465 | 4 | 95.4 | 99.1 | 3.7 |
| VRTNGKEPELLEPIPY | -3.5274491 | 0.26498634 | 2 | -4.4784249 | 1.64541355 | 4 | 92.0 | 95.7 | 3.7 |
| LGVQDKLVQTPL | -4.5185133 | 0.98724479 | 3 | -7.6528144 | 1.14401008 | 3 | 95.8 | 99.5 | 3.7 |
| VVQEPGDYEVSVKFNEEHIPDSPF | -3.4959823 | 0.38967078 | 3 | -4.4216891 | 0.38899647 | 2 | 91.9 | 95.5 | 3.7 |
| VWNTHADFADECPKPELL | -2.6161581 | 0.43104521 | 2 | -3.1163186 | 0.17150592 | 2 | 86.0 | 89.7 | 3.7 |
| SLCEKGDAMIMEETGKIF | -4.089288 | 0.92998934 | 5 | -5.7128625 | 2.6001392 | 5 | 94.5 | 98.1 | 3.7 |
| WKNPEQVDLY | -3.8578706 | 0.39539536 | 2 | -5.1310918 | 1.32165366 | 2 | 93.5 | 97.2 | 3.7 |
| AIDKSLTPVTL | -4.2115406 | 0.5648616 | 3 | -6.0712319 | 1.54838097 | 5 | 94.9 | 98.5 | 3.7 |
| MVELSKSQDDEIGDGTTGVVVL | -3.068546 | 0.56380215 | 2 | -3.731831 | 1.07094707 | 4 | 89.3 | 93.0 | 3.7 |
| AADESTGSIAKRLQ | -3.8495479 | 2.32645047 | 2 | -5.0968653 | 1.63315545 | 6 | 93.5 | 97.2 | 3.6 |
| LASPEYVNLPINGNGKQ | -4.2959526 | 3.38074113 | 8 | -6.3623162 | 1.40666005 | 2 | 95.2 | 98.8 | 3.6 |
| SVKSVPVSAGGEGETSPY | -4.2848607 | 0.7227698 | 3 | -6.3079831 | 0.68996118 | 2 | 95.1 | 98.8 | 3.6 |
| ILIVKWGDESVPGSPF | -3.5023349 | 1.24480517 | 4 | -4.415711 | 0.45051765 | 2 | 91.9 | 95.5 | 3.6 |
| TTTPRPVIVEPLEQLDDEDGLPEKL | -2.5758495 | 0.8336369 | 9 | -3.0562902 | 0.31906149 | 7 | 85.6 | 89.3 | 3.6 |
| TDKVVIGM(15.9949)DVAASEF | -3.2835661 | 0.29224276 | 2 | -4.0481821 | 0.62932049 | 3 | 90.7 | 94.3 | 3.6 |
| SNSKQTAISQPASGNTF | -4.4262791 | 0.28790405 | 2 | -6.8751793 | 0.97460462 | 3 | 95.6 | 99.2 | 3.6 |
| KGNFTLPEVAECFDEITY | -3.0536655 | 0.52593105 | 7 | -3.6973094 | 0.10119148 | 2 | 89.3 | 92.8 | 3.6 |
| TKDFSALESQLQDTQELLQEENRQKL | -4.2715743 | 1.76670356 | 5 | -6.1988963 | 0.7839249 | 2 | 95.1 | 98.7 | 3.6 |
| KQLNDLWDQIEQAHL | -3.8467845 | 0.06207696 | 2 | -5.0521867 | 0.2464826 | 2 | 93.5 | 97.1 | 3.6 |
| KAEIEEKGSDGAASSY | -4.6663549 | 3.39720822 | 2 | -8.8418205 | 1.58952479 | 2 | 96.2 | 99.8 | 3.6 |

| | | | | | | | | | |
|---|---|---|---|---|---|---|---|---|---|
| SALESQLQDTQELLQEENRQKL | -3.0468038 | 0.61436326 | 5 | -3.6820967 | 0.26323397 | 4 | 89.2 | 92.8 | 3.6 |
| IKEQIVGPLESPVEDSLWPY | -3.1168459 | 1.50016201 | 2 | -3.7828223 | 0.34669001 | 2 | 89.7 | 93.2 | 3.6 |
| NDSQRQATKDAGQISGL | -3.8082106 | 1.34847069 | 3 | -4.9584028 | 0.83874151 | 3 | 93.3 | 96.9 | 3.5 |
| KGQILMPNIGY | -4.3867773 | 0.54926947 | 2 | -6.5992378 | 1.44790655 | 3 | 95.4 | 99.0 | 3.5 |
| IDVKPLGVIW | -1.6595111 | 1.50466171 | 2 | -1.9551034 | 1.32588149 | 4 | 76.0 | 79.5 | 3.5 |
| LVYDQELLGPSDKSQAALQ | -2.8130199 | 0.53154237 | 3 | -3.3515301 | 0.04695792 | 2 | 87.5 | 91.1 | 3.5 |
| QRDGEDQTQDTELVETRPAGDGTFQKW | -3.1193338 | 0.44138836 | 2 | -3.7777992 | 0.18565317 | 3 | 89.7 | 93.2 | 3.5 |
| NVDIRNHQNLLDSLEQYVKGDLLEGANAY | -2.4675773 | 0.37726539 | 4 | -2.9027515 | 0.4739204 | 2 | 84.7 | 88.2 | 3.5 |
| DDSEPVQKVL | -4.6169434 | 1.30887028 | 3 | -7.9620116 | 1.53496824 | 2 | 96.1 | 99.6 | 3.5 |
| HMFTKEELEEVIKDI | -4.1944011 | 0.92210537 | 2 | -5.883075 | 0.35759765 | 2 | 94.8 | 98.3 | 3.5 |
| ELDSNNEKGLF | -4.13981 | 2.67759574 | 2 | -5.7242912 | 1.25162863 | 4 | 94.6 | 98.1 | 3.5 |
| QKSGVISGGASDL | -3.6463954 | 0.60011068 | 3 | -4.6289029 | 1.11382066 | 4 | 92.6 | 96.1 | 3.5 |
| LGAPGAISDLEEDLGRLGKTVDALTGW | -2.5913528 | 0.71967039 | 2 | -3.0563802 | 0.5581202 | 2 | 85.8 | 89.3 | 3.5 |
| VDISSNTAGKTLF | -4.0738243 | 1.34772188 | 4 | -5.5356177 | 1.39371416 | 5 | 94.4 | 97.9 | 3.5 |
| AGRLKPDEGGEVPVLNVSY | -3.5581427 | 0.86558698 | 3 | -4.4654302 | 0.52710685 | 2 | 92.2 | 95.7 | 3.5 |
| KGVGIISEGNETVEDIAARL | -3.7535111 | 0.79681573 | 5 | -4.8143593 | 2.04573492 | 2 | 93.1 | 96.6 | 3.5 |
| ILSNSEDKSIRVW | -3.8431789 | 0.25784334 | 4 | -4.9915107 | 0.7059594 | 2 | 93.5 | 97.0 | 3.5 |
| TTNLTEEEEKSKSLAKL | -4.0939487 | 4.42465665 | 4 | -5.5541828 | 1.16529015 | 3 | 94.5 | 97.9 | 3.4 |
| LDFKVTEGSFVY | -4.4716464 | 1.34167611 | 6 | -6.8287935 | 1.88473109 | 3 | 95.7 | 99.1 | 3.4 |
| AQQVQQVADDYGKCSRL | -2.8593718 | 0.37856194 | 4 | -3.3963251 | 0.14653713 | 4 | 87.9 | 91.3 | 3.4 |
| VRQSQIQVFEDGADTTSPETPDSSASKVL | -1.8483716 | 0.2033041 | 5 | -2.1579452 | 0.09138697 | 2 | 78.3 | 81.7 | 3.4 |
| KQDDGTGPEKIAH | -4.4565002 | 3.28356782 | 2 | -6.7288438 | 0.67267811 | 2 | 95.6 | 99.1 | 3.4 |
| TLQDKAQVADVVVSRW | -3.7326669 | 0.99292019 | 8 | -4.7509548 | 1.05657675 | 5 | 93.0 | 96.4 | 3.4 |
| ASGGCDNLIKLW | -0.9591651 | 0.25147184 | 3 | -1.1847398 | 0.57831196 | 4 | 66.0 | 69.4 | 3.4 |
| TAKNAVITVPAYF | -3.6506376 | 1.43588149 | 6 | -4.5940192 | 1.05923738 | 3 | 92.6 | 96.0 | 3.4 |
| GVTIDFDTVNKTPHTATL | -4.4854904 | 0.43543042 | 5 | -6.8091795 | 0.05033099 | 2 | 95.7 | 99.1 | 3.4 |
| KYEPDSANPDALQCPIVL | -3.4384791 | 0.16990647 | 3 | -4.2299784 | 0.4992224 | 5 | 91.6 | 94.9 | 3.4 |
| SLEMSKTEALTQAF | -4.2511819 | 1.45772438 | 5 | -5.9332018 | 1.0096266 | 3 | 95.0 | 98.4 | 3.4 |
| ISPFHDIPIYADKDVF | -4.1324555 | 0.9387614 | 2 | -5.5990878 | 1.14329105 | 2 | 94.6 | 98.0 | 3.4 |
| LFKDPCAAPNEGF | -3.9814715 | 0.85800408 | 5 | -5.2372438 | 3.59296563 | 6 | 94.0 | 97.4 | 3.4 |
| QTIMEKSETELKDTY | -4.2908611 | 0.57560316 | 2 | -6.0422388 | 1.47015573 | 3 | 95.1 | 98.5 | 3.4 |
| TLVEGENDEQSSTDQASAIKTKNVF | -4.3399795 | 0.4492218 | 2 | -6.1985542 | 2.58252895 | 2 | 95.3 | 98.7 | 3.4 |
| AIGDILEDKVELTPVAIQAGRL | -1.9870221 | 0.33999241 | 3 | -2.3089133 | 0.1030759 | 2 | 79.9 | 83.2 | 3.4 |
| TLTQAIDKF | -4.6911531 | 1.6670005 | 5 | -8.051138 | 2.65540575 | 4 | 96.3 | 99.6 | 3.4 |
| FTEAGLKELSEY | -4.0298125 | 0.81016172 | 3 | -5.3319614 | 0.43483878 | 3 | 94.2 | 97.6 | 3.3 |
| FGCIGIDKFGEIL | -3.7736852 | 1.17621468 | 6 | -4.7951866 | 0.76818736 | 3 | 93.2 | 96.5 | 3.3 |
| SILGMTPGFGDKTF | -3.0172165 | 0.51786608 | 3 | -3.5904545 | 0.76462157 | 2 | 89.0 | 92.3 | 3.3 |
| NAAKVPADTEVVCAPPTAYIDF | -2.5124114 | 0.26143442 | 5 | -2.9316258 | 0.08267838 | 3 | 85.1 | 88.4 | 3.3 |
| ALTGDEVKKICM | -3.4628408 | 0.18925865 | 2 | -4.2506488 | 0.23792869 | 2 | 91.7 | 95.0 | 3.3 |
| ELINQLDGFDPRGNIKVL | -4.2618924 | 0.53101771 | 2 | -5.9132223 | 1.64916888 | 4 | 95.0 | 98.4 | 3.3 |
| EALPELKLVIDQIDNGFF | -2.3947065 | 0.25258514 | 6 | -2.7859697 | 0.35291745 | 2 | 84.0 | 87.3 | 3.3 |
| HFKVDNDENEHQL | -3.7157164 | 1.35506192 | 3 | -4.676701 | 1.72652709 | 4 | 92.9 | 96.2 | 3.3 |
| SPKPGGDFGTAFPEIPVEF | -3.3996369 | 0.3520199 | 2 | -4.1429143 | 0.96346649 | 2 | 91.3 | 94.6 | 3.3 |
| EKRMATEVAADALGEEWKGY | -4.3898649 | 1.74963528 | 4 | -6.2978332 | 0.92180047 | 2 | 95.4 | 98.7 | 3.3 |
| IIDYYFEVVQKL | -3.9561085 | 1.71416538 | 4 | -5.1364667 | 1.29958315 | 9 | 93.9 | 97.2 | 3.3 |
| SDPFVEAEKSNLAY | -3.8499514 | 1.60692109 | 2 | -4.9170668 | 1.06282656 | 3 | 93.5 | 96.8 | 3.3 |
| IKPIFTDESYLELY | -3.6815964 | 0.76805006 | 6 | -4.6010799 | 1.11576136 | 4 | 92.8 | 96.0 | 3.3 |
| STISEKVIFF | 0.23377965 | 0.38495652 | 8 | 0.04459228 | 0.065301 | 3 | 46.0 | 49.2 | 3.3 |
| EIVTDKSQEGSQF | -4.0519823 | 1.49563849 | 3 | -5.3352981 | 2.42284437 | 3 | 94.3 | 97.6 | 3.3 |
| VGVDQFLVKTGTITTF | -4.3360523 | 1.00946648 | 6 | -6.0832657 | 1.81833939 | 3 | 95.3 | 98.5 | 3.3 |
| ALETIDINKDPY | -4.5268478 | 0.80617563 | 2 | -6.7859289 | 1.49413541 | 4 | 95.8 | 99.1 | 3.3 |
| SVCTNEDPKDRPSAAHIVEALETDV | -2.2904913 | 0.11309998 | 4 | -2.6533915 | 0.44897567 | 2 | 83.0 | 86.3 | 3.3 |
| VLKSAVEAERL | -4.7299904 | 1.06334694 | 8 | -8.0365186 | 2.46120229 | 3 | 96.4 | 99.6 | 3.3 |
| HIAASAGRDEIVKAL | -3.9013876 | 1.74304577 | 2 | -5.0035063 | 1.68048406 | 2 | 93.7 | 97.0 | 3.2 |
| IVSVDETIKNPRSTV | -4.6301257 | 0.95013784 | 5 | -7.2597343 | 1.70267965 | 4 | 96.1 | 99.4 | 3.2 |
| AFGDVDGVGINAKLQHPL | -4.3254446 | 0.6794849 | 2 | -6.0191945 | 2.15196308 | 2 | 95.2 | 98.5 | 3.2 |
| LAEKYEWDVAEARKIW | -4.4718925 | 6.49775825 | 3 | -6.5002205 | 1.63791305 | 4 | 95.7 | 98.9 | 3.2 |
| RAKTGGAYGEDLGADY | -4.3787362 | 1.29724913 | 6 | -6.1696261 | 0.95177193 | 4 | 95.4 | 98.6 | 3.2 |
| NATDLKDLSSHQLNEFLAQTL | -2.3200987 | 0.26913848 | 4 | -2.6834253 | 0.274275 | 2 | 83.3 | 86.5 | 3.2 |
| NLSFDEINKAFELM | -4.5383258 | 0.56224027 | 2 | -6.7412719 | 0.25472572 | 2 | 95.9 | 99.1 | 3.2 |
| SLLYNESLQVKTQLDEARGLL | -3.1774539 | 0.19799103 | 3 | -3.7871809 | 0.09320121 | 2 | 90.0 | 93.2 | 3.2 |
| ALKSMTEAEQQQLIDDHF | -2.6840912 | 0.06985812 | 3 | -3.1264739 | 0.21333452 | 2 | 86.5 | 89.7 | 3.2 |
| NASNNELVRTKTL | -4.1792455 | 0.8123472 | 5 | -5.5817501 | 0.84255374 | 5 | 94.8 | 98.0 | 3.2 |
| GSGGQKFPPLGGGGGIGY | -3.8818349 | 1.2717631 | 2 | -4.9315882 | 0.38963962 | 3 | 93.6 | 96.8 | 3.2 |
| VVRELKEQAQQAGLDDEVPHGQESDLF | -2.8929125 | 0.37932666 | 2 | -3.393698 | 0.45521075 | 2 | 88.1 | 91.3 | 3.2 |
| IKIIENAENEYQTAISENY | -1.9787563 | 0.44289555 | 3 | -2.2814742 | 0.29027274 | 3 | 79.8 | 82.9 | 3.2 |
| SLRKEEEQATETQPIVY | -4.5975756 | 0.68876945 | 2 | -6.9631473 | 2.61945529 | 3 | 96.0 | 99.2 | 3.2 |
| SHDGNTDEPEWVKTEREQFVEF | -2.7595371 | 0.03232888 | 2 | -3.2171337 | 1.23268876 | 2 | 87.1 | 90.3 | 3.2 |
| GLTFTEKWNTDNTLGTEITVEDQLARGL | -3.2023398 | 0.22052538 | 9 | -3.8119914 | 0.81424746 | 2 | 90.2 | 93.4 | 3.2 |
| IAEIEKQTKEQSQL | -4.763675 | 1.93186947 | 3 | -7.9683436 | 1.79456219 | 3 | 96.4 | 99.6 | 3.2 |
| VKLEAEGVPEVSEKY | -4.6026896 | 1.86806701 | 5 | -6.9470339 | 0.01888036 | 2 | 96.0 | 99.2 | 3.1 |
| TVTSLQETGLKVNQPASF | -4.82999 | 1.65127262 | 3 | -8.6233749 | 1.89845354 | 2 | 96.6 | 99.7 | 3.1 |

| | | | | | | | | | |
|---|---|---|---|---|---|---|---|---|---|
| DFPNIVIKGSELQLPF | -4.2521489 | 2.0197757 | 4 | -5.7341087 | 0.66463819 | 4 | 95.0 | 98.2 | 3.1 |
| SKSLGNVIDPLDVIY | -3.3504296 | 0.80805879 | 4 | -4.0246108 | 2.74141197 | 2 | 91.1 | 94.2 | 3.1 |
| ISGLPTIDIDDLKSNTEY | -3.7581443 | 2.41023178 | 2 | -4.6824834 | 0.14003621 | 2 | 93.1 | 96.3 | 3.1 |
| ANNCPALRKSEIEYY | -4.7097289 | 3.05475405 | 8 | -7.50039 | 1.41589335 | 7 | 96.3 | 99.5 | 3.1 |
| IISTDPAHNISDAFDQKF | -4.018043 | 0.89726401 | 6 | -5.1790241 | 1.31735235 | 3 | 94.2 | 97.3 | 3.1 |
| GPDGTGPNILTDITKGVQY | -4.5882535 | 2.01516138 | 5 | -6.805552 | 2.12271743 | 7 | 96.0 | 99.1 | 3.1 |
| SDYVGSGPPKGTGLH | -4.0684783 | 1.58636388 | 2 | -5.2676339 | 1.21875706 | 3 | 94.4 | 97.5 | 3.1 |
| HTERKTEPATGFIDGDLIESFL | -3.0194532 | 0.03598092 | 2 | -3.5456699 | 0.43357043 | 2 | 89.0 | 92.1 | 3.1 |
| ENYVADIEVDGKQVEL | -4.804238 | 1.08335144 | 3 | -8.0777256 | 1.40041439 | 4 | 96.5 | 99.6 | 3.1 |
| KYGDESSNSLPGHSVAL | -4.7412722 | 1.34069627 | 2 | -7.5864908 | 2.291828 | 3 | 96.4 | 99.5 | 3.1 |
| IQVDIGGGQTKTFAPEEISAM | -3.5591275 | 0.67684453 | 6 | -4.3297696 | 0.22438072 | 2 | 92.2 | 95.3 | 3.1 |
| SDYNIQKESTLHL | -3.9987743 | 1.80831813 | 3 | -5.1140873 | 0.48097754 | 2 | 94.1 | 97.2 | 3.1 |
| EIQKELLDY | -4.113773 | 0.43281005 | 2 | -5.3581308 | 2.97889034 | 4 | 94.5 | 97.6 | 3.1 |
| LFLGSSGIGKTEL | -4.3406435 | 2.58930732 | 5 | -5.9196611 | 1.40405554 | 3 | 95.3 | 98.4 | 3.1 |
| SLLRGGSDDSSKDPIDVNYEKL | -4.2942837 | 1.30158318 | 2 | -5.7835151 | 0.47545202 | 3 | 95.2 | 98.2 | 3.1 |
| RPGVACSVSQAQKDELILEGNDIELVSNSAAL | -2.1218069 | 0.2540835 | 4 | -2.4332116 | 0.31498437 | 3 | 81.3 | 84.4 | 3.1 |
| CFLEYEDHKTAAQ | -4.8595541 | 0.87203427 | 4 | -8.5242486 | 1.6097793 | 7 | 96.7 | 99.7 | 3.1 |
| CLDNGAKSVVL | -3.7300987 | 0.67467726 | 3 | -4.6035073 | 1.02700852 | 6 | 93.0 | 96.0 | 3.1 |
| KQLASEDISHITPTQGF | -4.5792639 | 1.96327443 | 5 | -6.6888654 | 2.03553655 | 4 | 96.0 | 99.0 | 3.1 |
| IEDAERDGTLKPGDTIIEPTSGNTGIGL | -2.8539874 | 0.21401503 | 2 | -3.3210314 | 0.1266432 | 2 | 87.8 | 90.9 | 3.1 |
| KGLDVDSLVIEHIQVN | -4.3495351 | 1.3506429 | 11 | -5.921595 | 1.4768002 | 9 | 95.3 | 98.4 | 3.1 |
| VVDVDKNIDINDVTPNCRVAL | -2.1972264 | 0.59555254 | 4 | -2.5191946 | 0.32200764 | 5 | 82.1 | 85.1 | 3.0 |
| SSPSEEVKSAASY | -4.7563931 | 1.67797581 | 4 | -7.5497165 | 2.11815749 | 3 | 96.4 | 99.5 | 3.0 |
| LMEEDEDAYKKQF | -4.5348937 | 0.44645432 | 2 | -6.4898301 | 1.2820537 | 6 | 95.9 | 98.9 | 3.0 |
| TALSETQDPEEVSVTVKAF | -4.5322438 | 1.46491861 | 4 | -6.4784959 | 1.36554004 | 4 | 95.9 | 98.9 | 3.0 |
| KGFDPLLNLVLDGTIEY | -4.1979091 | 1.31142563 | 4 | -5.5149598 | 0.41718754 | 4 | 94.8 | 97.9 | 3.0 |
| SDKLDFLEGDQKPL | -4.4446583 | 1.2122811 | 2 | -6.170273 | 1.28133088 | 2 | 95.6 | 98.6 | 3.0 |
| SLKVDVEALENSAGATY | -3.7218756 | 0.54613593 | 2 | -4.5757458 | 0.30520403 | 3 | 93.0 | 96.0 | 3.0 |
| LKDTPLYTDNTANGIALL | -4.1152281 | 1.53445202 | 2 | -5.3241714 | 1.00313451 | 3 | 94.5 | 97.6 | 3.0 |
| M(15.9949)GDKSENVQDLLLLDVAPL | -3.0003067 | 0.04470546 | 3 | -3.5058716 | 1.33407193 | 3 | 88.9 | 91.9 | 3.0 |
| TVSSLQESGLKVNQPASF | -4.4603938 | 2.72524403 | 5 | -6.2141869 | 1.11300803 | 4 | 95.7 | 98.7 | 3.0 |
| AIGLINEALDEGDAQKTL | -4.5153492 | 1.28904598 | 4 | -6.3812686 | 0.84451088 | 3 | 95.8 | 98.8 | 3.0 |
| QGLIVPDNPPYDKGAF | -4.4556293 | 1.19067499 | 5 | -6.1861256 | 1.5105419 | 4 | 95.6 | 98.6 | 3.0 |
| ALAIVEALNGKEVAAQ | -4.5477019 | 1.42561221 | 4 | -6.4842297 | 2.41570931 | 4 | 95.9 | 98.9 | 3.0 |
| NALLKVY | -4.7995081 | 2.71080814 | 2 | -7.6918743 | 3.21579433 | 2 | 96.5 | 99.5 | 3.0 |
| GEKRVSISEGDDKIEY | -4.6889136 | 0.50702819 | 4 | -7.0282948 | 0.19280085 | 2 | 96.3 | 99.2 | 3.0 |
| IMELINNVAKAHGGY | -4.2394872 | 1.012979 | 5 | -5.5694149 | 1.78211389 | 3 | 95.0 | 97.9 | 3.0 |
| MGTELNGKTL | -4.6523571 | 0.67100954 | 2 | -6.8399929 | 0.64596272 | 2 | 96.2 | 99.1 | 3.0 |
| LSKIEELQLIVNDKSQNL | -4.774408 | 1.19927066 | 3 | -7.4504 | 0.58605954 | 2 | 96.5 | 99.4 | 3.0 |
| GGLQKVFEHDSVEL | -4.5042145 | 1.6101225 | 5 | -6.2788574 | 1.3545912 | 3 | 95.8 | 98.7 | 2.9 |
| SQVLANGLDNKLREDLERL | -2.6861904 | 0.12441041 | 4 | -3.0916069 | 0.01956599 | 2 | 86.6 | 89.5 | 2.9 |
| LRSVGDGETVEFDVVEGEKGAEAANVTGPDGVP | -2.2432214 | 0.2596379 | 4 | -2.5602276 | 0.18080845 | 2 | 82.6 | 85.5 | 2.9 |
| VLEFHFEPNEYFTNEVLTKTY | -1.2709429 | 0.58838193 | 13 | -1.4822641 | 0.37669677 | 5 | 70.7 | 73.6 | 2.9 |
| RMIEDAERDGTLKPGDTIIEPTSGNTGIGL | -1.8884741 | 0.25836435 | 3 | -2.1557699 | 0.08949667 | 5 | 78.7 | 81.7 | 2.9 |
| AAQDVAQRCKELGITAL | -4.2703601 | 1.30313136 | 11 | -5.620313 | 1.13596396 | 8 | 95.1 | 98.0 | 2.9 |
| RGLGDCLVKIY | -4.6935468 | 1.53429569 | 6 | -6.9778145 | 0.60077446 | 2 | 96.3 | 99.2 | 2.9 |
| LEGKIDYGEY | -3.719582 | 0.71162749 | 3 | -4.5360085 | 0.69773536 | 2 | 92.9 | 95.9 | 2.9 |
| VSKPVSWADETDDLEGDVSTTW | -2.6148787 | 0.23797475 | 3 | -2.9998649 | 0.09277383 | 4 | 86.0 | 88.9 | 2.9 |
| GACGLSEEKAVEKFGEENIEVY | -4.7027397 | 1.08390246 | 4 | -6.9943768 | 2.67733128 | 3 | 96.3 | 99.2 | 2.9 |
| KVTQDELKEVF | -3.3733159 | 1.31674048 | 4 | -3.9981677 | 0.65028645 | 2 | 91.2 | 94.1 | 2.9 |
| IVSGADDRQVKIW | -3.4179814 | 4.54177438 | 2 | -4.0626093 | 0.36336276 | 3 | 91.4 | 94.4 | 2.9 |
| ALELLFDQLHEGAKALDVGSGSGIL | -1.625887 | 0.23783839 | 3 | -1.8629532 | 0.67153149 | 3 | 75.5 | 78.4 | 2.9 |
| GQAVKNDQCYDDIRVSRVTW | -3.0416204 | 0.10294013 | 2 | -3.5378066 | 0.45052264 | 3 | 89.2 | 92.1 | 2.9 |
| VLTGDRISAQDAKQAGL | -4.8052248 | 0.81086784 | 4 | -7.4788077 | 2.54729707 | 6 | 96.5 | 99.4 | 2.9 |
| KVVGDVAYDEAKERASF | -4.4276244 | 1.00089438 | 3 | -5.9871247 | 1.06806645 | 2 | 95.6 | 98.4 | 2.9 |
| LACPVSKEDGPSVF | -2.6417286 | 0.42397296 | 3 | -3.0271915 | 0.18486983 | 2 | 86.2 | 89.1 | 2.9 |
| KDAGVIAGLNVL | -4.5947701 | 0.82953838 | 5 | -6.5013447 | 3.08698264 | 4 | 96.0 | 98.9 | 2.9 |
| TQTQAVEGEIDLAKL | -3.8836936 | 1.89814151 | 3 | -4.8004366 | 1.15560819 | 3 | 93.7 | 96.5 | 2.9 |
| VQNDTLLQVKGTGANGSF | -4.9415398 | 1.32601306 | 4 | -8.5228843 | 1.27430889 | 3 | 96.8 | 99.7 | 2.9 |
| LAKAIANECQANF | -4.6070334 | 1.55122839 | 13 | -6.5419908 | 1.62595902 | 10 | 96.1 | 98.9 | 2.9 |
| AADVGKAGAEREF | -3.838854 | 1.21543855 | 7 | -4.7213395 | 1.59666047 | 7 | 93.5 | 96.3 | 2.9 |
| SGWYDADLSPAGHEEAKRGGQ | -2.9933483 | 0.36849371 | 10 | -3.4697468 | 0.16352089 | 3 | 88.8 | 91.7 | 2.9 |
| LFTKDIALVQQLF | -4.442395 | 0.57138913 | 2 | -6.0161445 | 1.32774847 | 2 | 95.6 | 98.5 | 2.9 |
| EKRMATEVAADALGEEW | -1.8153121 | 0.32902843 | 3 | -2.0681006 | 0.16673387 | 2 | 77.9 | 80.7 | 2.9 |
| TIDDLIDKLSTIVN | -4.9524312 | 1.77146439 | 5 | -8.5142651 | 2.05275805 | 2 | 96.9 | 99.7 | 2.9 |
| DATYETKESKKEDLVF | -4.8570567 | 1.66459232 | 2 | -7.693109 | 0.06965394 | 2 | 96.7 | 99.5 | 2.9 |
| GILADATEQVGQHKDAY | -4.6824926 | 0.65477612 | 6 | -6.7894638 | 2.42972087 | 5 | 96.3 | 99.1 | 2.9 |
| NCEEPTEKLPFPIIDDRNRELAILL | -1.7129633 | 0.41734037 | 4 | -1.9534035 | 0.37596451 | 3 | 76.6 | 79.5 | 2.9 |
| SLSLNQQPAAPECKVL | -4.2079702 | 0.68251039 | 6 | -5.41749 | 1.54731714 | 2 | 94.9 | 97.7 | 2.8 |
| RFEDVVNQSSPKNCTVY | -3.0675753 | 0.54993473 | 10 | -3.557477 | 1.20690617 | 2 | 89.3 | 92.2 | 2.8 |
| EVISKSIAPSIF | -4.8406375 | 0.97938678 | 3 | -7.4703173 | 1.66380087 | 3 | 96.6 | 99.4 | 2.8 |
| EIVFEDPKIPGEKQFAY | -4.4719429 | 2.00387845 | 4 | -6.0326958 | 1.40234299 | 3 | 95.7 | 98.5 | 2.8 |

| | | | | | | | | | |
|---|---|---|---|---|---|---|---|---|---|
| TVAKDPIVNVW | -4.4219235 | 1.38635073 | 8 | -5.8951999 | 1.61963307 | 6 | 95.5 | 98.3 | 2.8 |
| CRTKIDCDNLEQY | -4.6568868 | 0.58147414 | 5 | -6.6123602 | 0.14062211 | 2 | 96.2 | 99.0 | 2.8 |
| IIYNEDNGIIKAF | -4.77837 | 1.22404017 | 5 | -7.0997481 | 0.73906759 | 4 | 96.5 | 99.3 | 2.8 |
| AINEAVESGDVGKTL | -4.5809821 | 1.1369914 | 3 | -6.3337704 | 2.23117701 | 3 | 96.0 | 98.8 | 2.8 |
| FDNQIHEADTTENQSGVSFDKTSATW | -3.3230364 | 0.10072422 | 2 | -3.8943273 | 0.05205594 | 4 | 90.9 | 93.7 | 2.8 |
| APVNVTTEVKSVEM | -4.6391549 | 0.80855132 | 7 | -6.523049 | 0.77144628 | 2 | 96.1 | 98.9 | 2.8 |
| ELQASRVSSDVIDQKVY | -4.6953332 | 0.96220611 | 4 | -6.7211429 | 0.44526572 | 2 | 96.3 | 99.1 | 2.8 |
| DFEIETKQGTQY | -4.5939277 | 1.46326225 | 5 | -6.360368 | 1.23816962 | 4 | 96.0 | 98.8 | 2.8 |
| QLAIRNDEELNKL | -4.2979296 | 0.16630397 | 4 | -5.564803 | 1.01304382 | 3 | 95.2 | 97.9 | 2.8 |
| TIEDGIFEVKSTAGDTHLGGEDFDNRM(15.9949) | -2.4860144 | 0.14797461 | 8 | -2.8232155 | 0.40839292 | 6 | 84.9 | 87.6 | 2.8 |
| FKGTAVVNGEFKDLSLDDF | -4.2661746 | 0.50575663 | 2 | -5.4909843 | 0.70328629 | 4 | 95.1 | 97.8 | 2.8 |
| INTSDIILVGLRDYQDNKADVIL | -3.0832949 | 0.64624112 | 5 | -3.5633377 | 0.29622605 | 2 | 89.4 | 92.2 | 2.8 |
| VRNLATTVTEEILEKSFSEF | -2.6353453 | 0.21822674 | 6 | -2.9993619 | 0.28224317 | 6 | 86.1 | 88.9 | 2.7 |
| FGGFGEVESIELPMDNKTN | -2.6687755 | 0.24455878 | 5 | -3.0393523 | 0.27099029 | 5 | 86.4 | 89.2 | 2.7 |
| SNTLNFQISEVEPKYY | -4.2345772 | 1.2149512 | 3 | -5.4034285 | 1.09069043 | 4 | 95.0 | 97.7 | 2.7 |
| KAISEAQESVTKTTNY | -4.8567222 | 0.88584384 | 3 | -7.3693913 | 0.49280459 | 2 | 96.7 | 99.4 | 2.7 |
| VLQEITSQPKRQY | -4.3682503 | 0.64517335 | 7 | -5.7014407 | 1.22139643 | 5 | 95.4 | 98.1 | 2.7 |
| SEIISSISDVKF | -2.0488671 | 1.13566615 | 5 | -2.315233 | 0.31660077 | 8 | 80.5 | 83.3 | 2.7 |
| KAISAHFDDSSASSL | -4.0282245 | 0.39234837 | 4 | -4.9912988 | 1.36035586 | 2 | 94.2 | 97.0 | 2.7 |
| SAVAATKSPIIF | -4.4714896 | 2.12311224 | 6 | -5.9462708 | 1.38042416 | 5 | 95.7 | 98.4 | 2.7 |
| SNPKFAEHGETL | -4.4285167 | 1.1246694 | 2 | -5.8349703 | 1.34308953 | 5 | 95.6 | 98.3 | 2.7 |
| VAKEIQTTTGNQQVL | -4.4700611 | 1.18041218 | 4 | -5.9413245 | 2.80151555 | 4 | 95.7 | 98.4 | 2.7 |
| TGKLEDGTEFDSSLPQNQPF | -3.5092807 | 0.01800464 | 2 | -4.1419495 | 0.57064048 | 4 | 91.9 | 94.6 | 2.7 |
| KLDEDEDEDDADLSKY | -5.0553572 | 0.85999464 | 2 | -8.9082298 | 1.52469057 | 3 | 97.1 | 99.8 | 2.7 |
| FLQVKEGILNDDIYCPPETAVL | -3.3779691 | 0.51783339 | 6 | -3.9530846 | 0.4808969 | 3 | 91.2 | 93.9 | 2.7 |
| SQVLANGLDNKLREDLERL | -2.6974296 | 0.15730495 | 4 | -3.0687958 | 0.35716724 | 2 | 86.6 | 89.4 | 2.7 |
| AGETLSVNDPPDVLDRQKCL | -3.0059681 | 0.00827687 | 2 | -3.4532074 | 0.28189659 | 2 | 88.9 | 91.6 | 2.7 |
| GKWSTDDVQINDISLQDY | -3.2483529 | 0.17541995 | 7 | -3.7726756 | 0.56902326 | 6 | 90.5 | 93.2 | 2.7 |
| SGGTTMYPGIADRMQKEITAL | -2.987149 | 0.2110883 | 5 | -3.4257427 | 0.69637297 | 4 | 88.8 | 91.5 | 2.7 |
| KVDNDENEHQLSL | -4.299875 | 1.43636715 | 5 | -5.5109473 | 1.88777676 | 6 | 95.2 | 97.9 | 2.7 |
| RLSEMTEAEQQQLIDDHFLFDKPVSPL | -2.0692139 | 0.22866603 | 3 | -2.3322003 | 0.01289415 | 2 | 80.8 | 83.4 | 2.7 |
| GPTAEAAQLESSKRF | -4.5063038 | 0.78932934 | 10 | -5.9952981 | 1.74566129 | 7 | 95.8 | 98.5 | 2.7 |
| IKTAEELMNF | -3.7931031 | 0.35053918 | 2 | -4.5636322 | 1.24517699 | 2 | 93.3 | 95.9 | 2.7 |
| GASKLVPVGY | -4.7222472 | 1.33071687 | 8 | -6.659813 | 2.63450346 | 3 | 96.3 | 99.0 | 2.7 |
| SYSATEETLQEVFEKATF | -3.0610496 | 0.48803081 | 2 | -3.5159571 | 0.41606071 | 2 | 89.3 | 92.0 | 2.7 |
| VVDNGSGMCKAGF | -4.9116905 | 1.13461211 | 6 | -7.4835628 | 1.65179145 | 7 | 96.8 | 99.4 | 2.7 |
| EGERAMTKDNNLLGRF | -4.8889271 | 0.69622723 | 7 | -7.3547753 | 0.65120187 | 3 | 96.7 | 99.4 | 2.7 |
| VANNVTLPAGEQRKDVY | -4.8513544 | 1.68946657 | 12 | -7.1621875 | 1.81660682 | 5 | 96.7 | 99.3 | 2.7 |
| TGGKILGFFF | -4.8407536 | 0.46315332 | 2 | -7.110892 | 3.31655808 | 5 | 96.6 | 99.3 | 2.7 |
| QERDPSKIKWGDAGAEY | -4.9204941 | 3.07506016 | 11 | -7.5152625 | 2.70021536 | 4 | 96.8 | 99.5 | 2.7 |
| SNMNVIDRKPYPDDENLVEVKF | -4.8707803 | 0.70122117 | 3 | -7.2369224 | 2.38099846 | 3 | 96.7 | 99.3 | 2.6 |
| APVISAEKAYHEQLSVAEITNACFEPANQM | -2.3660076 | 0.1903009 | 3 | -2.6658297 | 0.34622059 | 3 | 83.8 | 86.4 | 2.6 |
| FDENPYFENKVLSKEF | -3.705745 | 0.9309418 | 5 | -4.4080454 | 1.08093971 | 3 | 92.9 | 95.5 | 2.6 |
| ANAAQKFPF | -4.7778595 | 1.72562197 | 4 | -6.7831401 | 0.65388386 | 2 | 96.5 | 99.1 | 2.6 |
| VGGLSFDTNEQSLEQVFSKY | -3.0636177 | 0.78333658 | 4 | -3.510534 | 0.28035327 | 2 | 89.3 | 91.9 | 2.6 |
| IGGLPNYLNDDQVKEL | -3.4722317 | 0.58301743 | 2 | -4.0613563 | 0.3203853 | 3 | 91.7 | 94.3 | 2.6 |
| TVKNLSPVVSNELLEQAF | -4.6156453 | 1.46165099 | 9 | -6.2174129 | 1.66817432 | 5 | 96.1 | 98.7 | 2.6 |
| DQTADQTPGATPKKL | -5.0947544 | 2.01715526 | 10 | -8.6392553 | 2.22545762 | 7 | 97.2 | 99.7 | 2.6 |
| DLTKYPDANPNPNEQ | -4.467798 | 1.31734652 | 3 | -5.8265259 | 0.97845404 | 4 | 95.7 | 98.3 | 2.6 |
| IVAAGVGEFEAGISKNGQTREHALLAY | -3.2379278 | 0.9613133 | 2 | -3.7331431 | 0.08218693 | 2 | 90.4 | 93.0 | 2.6 |
| QLALKDCEECIQLEPTFIKGY | -4.4841244 | 0.3306296 | 3 | -5.863197 | 0.93925706 | 3 | 95.7 | 98.3 | 2.6 |
| EVKSTNGDTFLGGEDFDQALL | -3.2019217 | 0.39903951 | 7 | -3.6843284 | 0.77043027 | 8 | 90.2 | 92.8 | 2.6 |
| SQVITKSEMIEY | -3.2184512 | 0.80832889 | 4 | -3.7056424 | 1.801701 | 4 | 90.3 | 92.9 | 2.6 |
| EAENKHVIDFF | -3.3941915 | 1.30631284 | 2 | -3.9424758 | 0.31819617 | 2 | 91.3 | 93.9 | 2.6 |
| FKGTAVVNGEFKDLSLDDF | -4.5842639 | 0.69585926 | 2 | -6.1133966 | 0.42978587 | 3 | 96.0 | 98.6 | 2.6 |
| QQNYQNSESGEKNEGSESAPEGQAQQRRPY | -2.4297573 | 0.17701785 | 3 | -2.7307238 | 0.39621493 | 2 | 84.3 | 86.9 | 2.6 |
| DEILEASDGIMVARGDLGIEIPAEKVF | -3.1553734 | 0.58726191 | 11 | -3.6175756 | 0.51908122 | 8 | 89.9 | 92.5 | 2.6 |
| RQQEQQVPILEKF | -3.9905935 | 1.3805688 | 3 | -4.845599 | 0.75521983 | 2 | 94.1 | 96.6 | 2.6 |
| GIWEPLAVKLQTY | -3.605378 | 0.59801503 | 3 | -4.2368181 | 0.65203157 | 2 | 92.4 | 95.0 | 2.6 |
| GVTIDFDTVNKTPHTA | -3.6434147 | 1.03513332 | 4 | -4.2926199 | 0.40204112 | 2 | 92.6 | 95.1 | 2.6 |
| HGEGDQEPGLEPGDIIIVLDQKDHAVF | -2.9707539 | 0.65341644 | 6 | -3.3798133 | 1.22951251 | 5 | 88.7 | 91.2 | 2.5 |
| AIRNDEELNKLL | -4.451194 | 0.49323899 | 5 | -5.7502902 | 3.38820946 | 2 | 95.6 | 98.2 | 2.5 |
| KFVDGLM(15.9949)IHSGDPVNYY | -3.5052127 | 0.51353621 | 6 | -4.0844169 | 0.08581144 | 2 | 91.9 | 94.4 | 2.5 |
| TGEGQKLGSTAPQVL | -3.7823124 | 0.28610139 | 2 | -4.4924371 | 1.75785077 | 4 | 93.2 | 95.7 | 2.5 |
| EVFEKDPIKY | -4.835318 | 1.32821965 | 5 | -6.835838 | 1.65993896 | 6 | 96.6 | 99.1 | 2.5 |
| ALLGDLTKACF | -4.4864124 | 1.04599955 | 11 | -5.8060667 | 2.14815206 | 7 | 95.7 | 98.2 | 2.5 |
| EIKKIGDEYFTF | -5.0753484 | 0.90617723 | 7 | -8.0861424 | 2.72036027 | 2 | 97.1 | 99.6 | 2.5 |
| KMYPIDFEKDDDSNF | -4.079849 | 0.92608629 | 6 | -4.9803749 | 1.02023906 | 5 | 94.4 | 96.9 | 2.5 |
| FINTSKGQKCEFQDAY | -4.9952391 | 0.18369464 | 3 | -7.5568255 | 2.08832021 | 3 | 97.0 | 99.5 | 2.5 |
| SKQILEILNSTGVEYETFDILEDEEVRQGL | -1.9198866 | 0.15210297 | 2 | -2.1493567 | 0.28038538 | 2 | 79.1 | 81.6 | 2.5 |
| EAKGPVTDVAY | -4.3400464 | 0.25429337 | 2 | -5.4736312 | 1.00224815 | 2 | 95.3 | 97.8 | 2.5 |
| FDENPYFENKVLSKEF | -4.1501335 | 1.01989241 | 3 | -5.102796 | 0.18045823 | 4 | 94.7 | 97.2 | 2.5 |

| | | | | | | | | | |
|---|---|---|---|---|---|---|---|---|---|
| SSSGLLEWESKSDALETLGFLNHY | -3.3548243 | 0.37342014 | 8 | -3.8685576 | 0.04708255 | 2 | 91.1 | 93.6 | 2.5 |
| LSVAKGSDEPPVFLEIHY | -3.7629217 | 0.66582344 | 5 | -4.453678 | 0.70992817 | 5 | 93.1 | 95.6 | 2.5 |
| SVDDIVKGINSSNVENQLQATQ | -2.8050685 | 0.17809904 | 4 | -3.1659054 | 0.09494565 | 3 | 87.5 | 90.0 | 2.5 |
| GDLKVGDQVW | -4.89443 | 1.2929314 | 3 | -7.0046122 | 1.63599952 | 3 | 96.7 | 99.2 | 2.5 |
| KGTYFPTWEGLF | -4.4044371 | 0.88905256 | 4 | -5.5914636 | 0.94205495 | 2 | 95.5 | 98.0 | 2.5 |
| LGELKDKNEQAFEEVFQNANF | -4.6795993 | 1.52383168 | 2 | -6.2707141 | 1.77257225 | 4 | 96.2 | 98.7 | 2.5 |
| FNGKEPSRGINPDEAVAY | -5.1653883 | 0.57744417 | 6 | -8.7065167 | 1.78087328 | 2 | 97.3 | 99.8 | 2.5 |
| VELQKEEAQKLL | -4.2747229 | 1.07945213 | 4 | -5.3208886 | 1.26315876 | 3 | 95.1 | 97.6 | 2.5 |
| IFAGKQLEDGRTLSDY | -5.0822196 | 0.84999891 | 9 | -7.9730341 | 2.81817438 | 2 | 97.1 | 99.6 | 2.5 |
| NKNSNIPGSSANTSTPTVSSY | -2.8157695 | 0.3077065 | 7 | -3.1752922 | 0.54622777 | 3 | 87.6 | 90.0 | 2.5 |
| IVAAGVGEFEAGISKNGQTREH | -3.1007279 | 0.7711864 | 3 | -3.5292042 | 1.06974564 | 5 | 89.6 | 92.0 | 2.5 |
| NSGKVDIVAINDPFIDLN | -3.6571163 | 0.4347615 | 3 | -4.2855862 | 0.51536001 | 2 | 92.7 | 95.1 | 2.5 |
| DQLALDSPKGCGTVLL | -5.0539157 | 2.32108642 | 3 | -7.7359307 | 1.11565871 | 3 | 97.1 | 99.5 | 2.5 |
| ANAGKDTNGSQFF | -4.2982358 | 1.93279105 | 5 | -5.3563525 | 1.25457749 | 5 | 95.2 | 97.6 | 2.5 |
| LASPEYVNLPINGNGKQ | -4.3284674 | 2.07088347 | 10 | -5.4167564 | 1.73690587 | 6 | 95.3 | 97.7 | 2.5 |
| KFWEVISDEHGIDPTGTY | -3.8266324 | 0.66203436 | 5 | -4.5366741 | 0.20847197 | 2 | 93.4 | 95.9 | 2.5 |
| DVVHVKDANGNSF | -4.3360386 | 1.06595115 | 4 | -5.4291563 | 1.29442206 | 8 | 95.3 | 97.7 | 2.4 |
| QIATVTEKGVEIEGPLSTETNWDIAHMISGFE | -2.2826451 | 0.28266243 | 4 | -2.548293 | 0.43207204 | 2 | 83.0 | 85.4 | 2.4 |
| GATLEIVTDKSQEGSQF | -4.1361928 | 1.0177072 | 6 | -5.0482317 | 1.08850949 | 7 | 94.6 | 97.1 | 2.4 |
| LVKAGQAVDDFIEKLVPLLDTGDIIIDGGNSEY | -5.1616401 | 1.26622913 | 3 | -8.4859734 | 0.59853389 | 2 | 97.3 | 99.7 | 2.4 |
| SVPCILGQNGISDLVKVTL | -2.3889913 | 0.44658049 | 4 | -2.6678846 | 0.3827924 | 3 | 84.0 | 86.4 | 2.4 |
| DVPTASVTEIQEKW | -4.4555553 | 1.05024223 | 4 | -5.6691341 | 1.83837813 | 6 | 95.6 | 98.1 | 2.4 |
| FLINGPEIMSKL | -4.6822213 | 0.22301672 | 4 | -6.2235984 | 1.04197262 | 2 | 96.3 | 98.7 | 2.4 |
| ANGLDNKLREDLERL | -4.7576684 | 1.14208767 | 2 | -6.4423898 | 0.94434957 | 4 | 96.4 | 98.9 | 2.4 |
| THVKIFDPQNPDENE | -3.6749251 | 0.73967303 | 6 | -4.2990438 | 0.86794346 | 3 | 92.7 | 95.2 | 2.4 |
| TTTPRPVIVEPLEQLDDEDGLPEKL | -3.0260644 | 0.18646374 | 2 | -3.4265439 | 0.12682526 | 2 | 89.1 | 91.5 | 2.4 |
| ISKLQDEFENRTRNVDVY | -3.9850478 | 0.06169865 | 2 | -4.7778588 | 0.40289134 | 2 | 94.1 | 96.5 | 2.4 |
| LAEFATGNDRKEAAENSLVAY | -3.5742406 | 0.63080701 | 6 | -4.1524153 | 0.93831297 | 3 | 92.3 | 94.7 | 2.4 |
| KQVLGQMVIDEELLGDGHSY | -3.1337986 | 0.32167887 | 6 | -3.5607158 | 1.05196618 | 4 | 89.8 | 92.2 | 2.4 |
| NYEGSPIKVTLATL | -4.0279499 | 0.99551602 | 8 | -4.8439674 | 1.20183573 | 6 | 94.2 | 96.6 | 2.4 |
| IEVDDERKLRTF | -4.7592999 | 0.71134618 | 4 | -6.4209561 | 1.5786167 | 4 | 96.4 | 98.8 | 2.4 |
| GLVDIVKGTNSY | -4.2321677 | 1.13992642 | 4 | -5.2002317 | 1.31876255 | 4 | 94.9 | 97.4 | 2.4 |
| VLIPDLKWNQQQLDDLY | -2.6597323 | 0.55092793 | 3 | -2.9783673 | 0.66532572 | 2 | 86.3 | 88.7 | 2.4 |
| SVDRKVSQPIEGHAASF | -4.6428359 | 1.46566089 | 7 | -6.0901363 | 1.61499821 | 3 | 96.2 | 98.6 | 2.4 |
| AEHISDECKRRFY | -5.1377084 | 2.49917271 | 3 | -8.0976875 | 4.22622309 | 2 | 97.2 | 99.6 | 2.4 |
| GEGKSTTTIGLVQALGAHLY | -3.7485209 | 1.59584759 | 3 | -4.3981213 | 0.84661997 | 2 | 93.1 | 95.5 | 2.4 |
| KASAETVDPASLWEY | -4.4787965 | 1.93889946 | 3 | -5.6911961 | 1.34570951 | 5 | 95.7 | 98.1 | 2.4 |
| INLPEDTETAKSLPDTY | -4.7873836 | 1.05894147 | 2 | -6.4887861 | 2.73190549 | 4 | 96.5 | 98.9 | 2.4 |
| VCKSESVPPVTDW | -5.1600994 | 0.891661 | 8 | -8.2499789 | 2.09776816 | 4 | 97.3 | 99.7 | 2.4 |
| AAQDVAQRCKELGITAL | -4.7946715 | 0.97544859 | 7 | -6.5110863 | 1.3525987 | 5 | 96.5 | 98.9 | 2.4 |
| VLVGDGGTGKTTF | -2.9939153 | 0.39497576 | 9 | -3.3800193 | 0.38158504 | 8 | 88.8 | 91.2 | 2.4 |
| APVISAEKAYHEQLSVAEITNACFEPANQ | -2.5283614 | 0.37362884 | 2 | -2.8220506 | 0.32071448 | 2 | 85.2 | 87.6 | 2.4 |
| KPESEELTAERITEF | -4.7290121 | 0.19799606 | 2 | -6.3033711 | 2.21663404 | 5 | 96.4 | 98.7 | 2.4 |
| QIATVTEKGVEIEGPLSTETNW | -2.6396278 | 0.12058966 | 2 | -2.9512467 | 0.44496894 | 3 | 86.2 | 88.6 | 2.4 |
| VGGIKEDTEEHHLRDYF | -4.4212722 | 1.39458014 | 4 | -5.5500106 | 1.12001367 | 3 | 95.5 | 97.9 | 2.4 |
| QHGKVEIIANDQGNRTTPSYVAF | -2.8501779 | 0.14303283 | 2 | -3.2006268 | 0.11364303 | 3 | 87.8 | 90.2 | 2.4 |
| AQAVEGPSENVRKL | -4.239414 | 2.2128129 | 10 | -5.1928214 | 1.55362138 | 4 | 95.0 | 97.3 | 2.4 |
| VVTGSVDQTVKVW | -3.6056107 | 0.77747449 | 3 | -4.1805403 | 0.90687387 | 4 | 92.4 | 94.8 | 2.4 |
| ALQLAEKLGSLVENNERVF | -3.5826465 | 0.17246599 | 3 | -4.1475773 | 0.3192586 | 2 | 92.3 | 94.7 | 2.4 |
| NSGKVDIVAINDPFIDLNY | -2.4913622 | 0.46879074 | 6 | -2.7747511 | 0.26891061 | 4 | 84.9 | 87.3 | 2.3 |
| KAENNSEVGASGY | -4.5072689 | 1.33200945 | 3 | -5.7138508 | 1.44230921 | 4 | 95.8 | 98.1 | 2.3 |
| NEGLWEIDNNPKVKF | -5.0810951 | 1.77894136 | 5 | -7.5555098 | 1.30889845 | 4 | 97.1 | 99.5 | 2.3 |
| EVTVPAQNTGLGPEKTSFF | -4.0013525 | 0.59165371 | 4 | -4.7681978 | 2.93697699 | 4 | 94.1 | 96.5 | 2.3 |
| GADDIELLPEAQHKAEVY | -4.5457313 | 1.41734286 | 5 | -5.7874073 | 1.00014391 | 5 | 95.9 | 98.2 | 2.3 |
| FQKLENDQIESL | -5.0086443 | 0.99657799 | 3 | -7.1734834 | 0.22632038 | 2 | 97.0 | 99.3 | 2.3 |
| GAIAPCEVTVPAQNTGLGPEKTSFF | -2.9465919 | 0.27333868 | 2 | -3.3095833 | 0.06138561 | 2 | 88.5 | 90.8 | 2.3 |
| IIHGSDSVESAEKEIGLW | -3.9719829 | 2.16263919 | 5 | -4.713631 | 0.6405714 | 4 | 94.0 | 96.3 | 2.3 |
| KDSAYPEELSRVTASGFPVIL | -2.5696827 | 0.21174247 | 4 | -2.8612962 | 0.871056 | 3 | 85.6 | 87.9 | 2.3 |
| KTDSSPNQARAQAAL | -4.2284435 | 1.24228427 | 11 | -5.1456311 | 1.02048506 | 3 | 94.9 | 97.3 | 2.3 |
| LSGGQSEEEASINLNAINKCPL | -3.655971 | 0.34196139 | 5 | -4.2371049 | 0.13264729 | 2 | 92.6 | 95.0 | 2.3 |
| ENLGIQPPKGVL | -4.8325724 | 1.20525548 | 7 | -6.5146401 | 1.93209207 | 5 | 96.6 | 98.9 | 2.3 |
| SVAEASKNETGGGEGIEVL | -4.404599 | 1.55099296 | 3 | -5.4707935 | 1.1271874 | 4 | 95.5 | 97.8 | 2.3 |
| AALEKEAQTSF | -4.7069484 | 1.63602955 | 3 | -6.1491244 | 1.94219818 | 4 | 96.3 | 98.6 | 2.3 |
| QHGKVEIIANDQGNRTTPSY | -2.4300285 | 0.12938112 | 2 | -2.6978493 | 0.23998113 | 2 | 84.3 | 86.6 | 2.3 |
| SFDLGKGEVIKAW | -5.0533021 | 0.37256267 | 2 | -7.3106051 | 1.47979051 | 2 | 97.1 | 99.4 | 2.3 |
| YNEATGGKYVPRAIL | -4.8926529 | 1.63085418 | 14 | -6.6813705 | 1.99112786 | 8 | 96.7 | 99.0 | 2.3 |
| AIGSDGLCCQSREVKEW | -2.5978927 | 0.31274934 | 3 | -2.8889679 | 0.10149927 | 3 | 85.8 | 88.1 | 2.3 |
| TLGGQKCSVIRDSLLQDGEF | -2.7168913 | 0.44904418 | 7 | -3.0280068 | 0.62333467 | 4 | 86.8 | 89.1 | 2.3 |
| SACSACLGPAAPAAANSSGDGGAAGDGTVVDCI | -2.6118957 | 0.4076898 | 3 | -2.9051334 | 0.28767412 | 2 | 85.9 | 88.2 | 2.3 |
| QAGQCGNQIGAKFW | -4.4418914 | 1.70433669 | 10 | -5.5255488 | 1.04414347 | 8 | 95.6 | 97.9 | 2.3 |
| GACDGGTGTKVRQGTL | -5.2204141 | 2.75347713 | 5 | -8.190334 | 2.51086594 | 2 | 97.4 | 99.7 | 2.3 |
| SSFYPDGGDQETAKTGKF | -5.0430973 | 0.25095173 | 2 | -7.197848 | 0.98162832 | 2 | 97.1 | 99.3 | 2.3 |

| | | | | | | | | | |
|---|---|---|---|---|---|---|---|---|---|
| VTAKDAERAINTL | -3.8989432 | 2.39897283 | 4 | -4.5769569 | 1.31065787 | 2 | 93.7 | 96.0 | 2.3 |
| ENLREIGNLLHPSVPISNDEDVDNKVERIW | -2.3342682 | 0.20471499 | 5 | -2.5841557 | 0.22086345 | 3 | 83.5 | 85.7 | 2.3 |
| TVLVDNYVKEEEGTGVVHQAPY | -2.7687782 | 0.1594566 | 4 | -3.0850056 | 0.21864843 | 4 | 87.2 | 89.5 | 2.3 |
| EVSENGNLVVSGKVY | -4.2421435 | 0.73822932 | 2 | -5.1302091 | 1.35530866 | 4 | 95.0 | 97.2 | 2.2 |
| ENKREFEDAFPADF | -4.2954274 | 1.0009915 | 2 | -5.2256693 | 0.0041989 | 2 | 95.2 | 97.4 | 2.2 |
| VGVNLPQKAGGF | -5.0150258 | 1.58926304 | 5 | -7.0238187 | 2.71597711 | 2 | 97.0 | 99.2 | 2.2 |
| LSDLTHQISKDYGVY | -4.8588219 | 0.48658652 | 4 | -6.4980215 | 0.64277024 | 4 | 96.7 | 98.9 | 2.2 |
| DMAENVVIVASQKRPL | -4.0453807 | 1.33564444 | 3 | -4.7956629 | 1.64970714 | 2 | 94.3 | 96.5 | 2.2 |
| SVSQAQKDELILEGNDIELVSNSAAL | -3.3922262 | 0.35516415 | 4 | -3.8543101 | 0.81518113 | 3 | 91.3 | 93.5 | 2.2 |
| AAVTEVIRSQGGKETETEYF | -2.3313535 | 0.35629194 | 2 | -2.5771932 | 0.30831664 | 3 | 83.4 | 85.6 | 2.2 |
| NAKDLEEALEMGVDW | -4.1855887 | 0.0948226 | 2 | -5.0168848 | 0.14553342 | 2 | 94.8 | 97.0 | 2.2 |
| LLIPVDNGTLTDFLEKEEEEA | -3.374268 | 0.60161128 | 2 | -3.825893 | 0.37872827 | 2 | 91.2 | 93.4 | 2.2 |
| ITGETKDQVANSAFVERL | -4.5317373 | 2.24130553 | 3 | -5.6616075 | 1.50220121 | 3 | 95.9 | 98.1 | 2.2 |
| DALYDEDVVKEDAFY | -3.4680486 | 0.14998758 | 3 | -3.9485758 | 0.93181954 | 5 | 91.7 | 93.9 | 2.2 |
| TVLVDNYVKEEEGTGVVHQAPY | -2.9088075 | 0.38959959 | 3 | -3.2431639 | 0.0735885 | 2 | 88.2 | 90.4 | 2.2 |
| KIDIIPNPQERTL | -4.8157566 | 1.56736925 | 4 | -6.3242571 | 0.5490605 | 8 | 96.6 | 98.8 | 2.2 |
| NCETEDYGEKFDENDVITCF | -3.0175691 | 0.51128251 | 3 | -3.3739162 | 0.38073291 | 2 | 89.0 | 91.2 | 2.2 |
| AAQELQAKLAEIGAPIQGNREELVERLQSY | -2.4154365 | 0.42058514 | 2 | -2.6679396 | 0.2516873 | 2 | 84.2 | 86.4 | 2.2 |
| IGDAAKNQVALNPQNTVF | -4.8977195 | 1.39510859 | 12 | -6.5496395 | 2.06663797 | 6 | 96.8 | 98.9 | 2.2 |
| EIMDGAPVKGESIPIRLF | -2.1701298 | 0.13221412 | 3 | -2.3932507 | 0.38207041 | 3 | 81.8 | 84.0 | 2.2 |
| TLSGADPEGCFPVILGHEGAGIVESVGEGVTKL | -2.2504344 | 0.29429474 | 6 | -2.4824371 | 0.33051904 | 2 | 82.6 | 84.8 | 2.2 |
| IVVIGHVDSGKSTT | -5.0582736 | 3.1850336 | 2 | -7.0820827 | 2.69479239 | 2 | 97.1 | 99.3 | 2.2 |
| KQMGEQRELQEGTY | -5.2397112 | 1.15761749 | 7 | -7.958504 | 1.23712975 | 6 | 97.4 | 99.6 | 2.2 |
| GTVIQEVKTSNVAREAALGAGF | -2.3939275 | 0.26541412 | 5 | -2.6419607 | 0.37471848 | 2 | 84.0 | 86.2 | 2.2 |
| QVRDIENLKDASSF | -5.0887806 | 1.43551197 | 9 | -7.1851486 | 2.27884235 | 3 | 97.1 | 99.3 | 2.2 |
| AFIEFASFEDAKEALN | -3.7964543 | 2.54002641 | 9 | -4.3913393 | 0.20623975 | 2 | 93.3 | 95.5 | 2.2 |
| SWPNDKDPVVVPFPTMTF | -2.6226015 | 0.0040837 | 2 | -2.9012411 | 0.83522598 | 2 | 86.0 | 88.2 | 2.2 |
| TFGAEVVAKF | -4.2623851 | 1.66256583 | 6 | -5.1195075 | 3.94528339 | 6 | 95.0 | 97.2 | 2.2 |
| MDRGEGGTTNPHIFPEGSEPKVY | -3.5685898 | 0.4298496 | 3 | -4.0704826 | 0.34136549 | 2 | 92.2 | 94.4 | 2.2 |
| TDKVVIGMDVAASEF | -4.8221147 | 1.60889065 | 4 | -6.2890682 | 1.32730263 | 5 | 96.6 | 98.7 | 2.2 |
| LAIQSSQAGGELKSSTPAQL | -4.7603914 | 1.61047713 | 3 | -6.1310652 | 0.9307333 | 3 | 96.4 | 98.6 | 2.2 |
| KDGRAGVVANDAGDRVTPAVVAY | -2.7409863 | 0.07558481 | 3 | -3.0367355 | 0.75759589 | 2 | 87.0 | 89.1 | 2.1 |
| NIQKESTL | -4.5497721 | 1.07402809 | 3 | -5.6494428 | 1.18926073 | 6 | 95.9 | 98.0 | 2.1 |
| NDSQRQATKDAGVIAGLNVL | -4.9191108 | 1.67120006 | 15 | -6.5354581 | 0.9099895 | 8 | 96.8 | 98.9 | 2.1 |
| KLEDTENWLYEDGEDQPKQVY | -4.9033722 | 1.22732832 | 2 | -6.4887879 | 0.40054007 | 2 | 96.8 | 98.9 | 2.1 |
| KLDYFGEEAFLTQSSQLY | -3.0526185 | 0.3565048 | 5 | -3.4049542 | 1.01552142 | 3 | 89.2 | 91.4 | 2.1 |
| IGNLNESVTPADLEKVF | -4.6996049 | 1.75317114 | 5 | -5.9607731 | 1.09833827 | 4 | 96.3 | 98.4 | 2.1 |
| LAADKDGNVTCEREVPGPDCRF | -3.0663717 | 0.20240756 | 3 | -3.4200285 | 0.06244806 | 2 | 89.3 | 91.5 | 2.1 |
| KSSPEPVALTESETEY | -4.6888112 | 1.52853957 | 6 | -5.9290937 | 1.04569655 | 8 | 96.3 | 98.4 | 2.1 |
| LQEFYQDDELGKKQF | -4.8029573 | 0.56485871 | 2 | -6.1966743 | 2.09516386 | 6 | 96.5 | 98.7 | 2.1 |
| AFIEFASFEDAKEAL | -4.1668086 | 3.0305586 | 12 | -4.9357884 | 2.23994714 | 8 | 94.7 | 96.8 | 2.1 |
| AKATGATQQDANASSLLDIY | -4.8713882 | 1.23447292 | 5 | -6.3479982 | 1.29751897 | 3 | 96.7 | 98.8 | 2.1 |
| YPLEIDYGQDEEAVKKL | -4.9556934 | 0.53236081 | 6 | -6.5831589 | 0.97989373 | 5 | 96.9 | 99.0 | 2.1 |
| YVSNIDGTHIAKTL | -4.8339089 | 1.07378966 | 5 | -6.2456719 | 1.36776372 | 8 | 96.6 | 98.7 | 2.1 |
| TGFIVEADTPGIQIGRKEL | -1.2349867 | 0.8883069 | 6 | -1.3819257 | 1.23077539 | 5 | 70.2 | 72.3 | 2.1 |
| ALGAGDLFNVNDNSEYVETIIAKCIDHY | -3.4906601 | 0.66928303 | 8 | -3.9484221 | 0.33116289 | 2 | 91.8 | 93.9 | 2.1 |
| KVDVEALENSAGATY | -4.2843994 | 1.14124621 | 4 | -5.1152623 | 0.71789985 | 4 | 95.1 | 97.2 | 2.1 |
| QIENFVPVLVEEAFHGKFY | -2.5460409 | 0.23174002 | 3 | -2.8016339 | 0.43915428 | 2 | 85.4 | 87.5 | 2.1 |
| FDKRDNSDFDLLTVSETANEPPQDEGNSF | -4.4883525 | 1.00150517 | 5 | -5.4802274 | 1.74252994 | 3 | 95.7 | 97.8 | 2.1 |
| RSCGSSTPDEFPTDIPGTKGNF | -2.8810897 | 0.29977973 | 10 | -3.1896073 | 0.03313278 | 2 | 88.0 | 90.1 | 2.1 |
| QAKDVEGSTSPQIGDKVEF | -5.4255564 | 0.85761017 | 7 | -8.9600439 | 1.42245562 | 5 | 97.7 | 99.8 | 2.1 |
| QLVDTARTQKTAY | -4.9255482 | 1.00438412 | 4 | -6.4732595 | 1.83894516 | 2 | 96.8 | 98.9 | 2.1 |
| IFVKDVPNSQL | -1.2538393 | 0.61450502 | 13 | -1.3998089 | 0.20827045 | 3 | 70.5 | 72.5 | 2.1 |
| ISEAQAIKADLAAVEAKVN | -5.3049883 | 1.33671889 | 4 | -7.9322469 | 1.98866529 | 4 | 97.5 | 99.6 | 2.1 |
| GKDATNVGDEGGFAPN | -1.4894928 | 0.99829402 | 6 | -1.6470387 | 1.96341469 | 3 | 73.7 | 75.8 | 2.1 |
| TIDDGIFEVKATAGDTHLGGEDFDNRLVNHF | -2.5646567 | 0.12321456 | 3 | -2.820496 | 0.20122384 | 5 | 85.5 | 87.6 | 2.1 |
| NYEGSPIKVTL | -4.9221433 | 1.0240431 | 4 | -6.4426108 | 2.0007035 | 4 | 96.8 | 98.9 | 2.1 |
| MDVISIDKTGENF | -4.7955805 | 1.32844816 | 6 | -6.1179235 | 1.37666909 | 4 | 96.5 | 98.6 | 2.1 |
| LTTFPRPVTVEPMDQLDDEEGLPEKL | -1.6720333 | 0.09693522 | 2 | -1.8401592 | 0.21465766 | 4 | 76.1 | 78.2 | 2.1 |
| TIDNGVFEVVATNGDTHLGGEDFDQRVMEHFIK | -2.6401057 | 0.28958399 | 2 | -2.9058316 | 0.35358707 | 2 | 86.2 | 88.2 | 2.1 |
| GFIERGDVVKEIFF | -4.2230716 | 1.51761931 | 2 | -4.999367 | 0.86413051 | 4 | 94.9 | 97.0 | 2.1 |
| AVNDFELARADFQKVL | -3.9094651 | 0.52937133 | 3 | -4.5149302 | 1.06674665 | 2 | 93.8 | 95.8 | 2.0 |
| LIKTVETRDGQVINETSQHHDDLE | -3.2932919 | 0.56474323 | 14 | -3.686122 | 0.26781633 | 7 | 90.7 | 92.8 | 2.0 |
| VSHDSTVSVADASKSVQVSTL | -2.4617173 | 0.43067745 | 4 | -2.702375 | 0.45315034 | 2 | 84.6 | 86.7 | 2.0 |
| DIPLVKNWY | -4.4313924 | 0.65202995 | 7 | -5.3534245 | 1.4595047 | 5 | 95.6 | 97.6 | 2.0 |
| IEKIQPSGGTNINEALL | -5.3570449 | 4.07370262 | 3 | -8.1923269 | 1.42156317 | 6 | 97.6 | 99.7 | 2.0 |
| KVFLENVIRDAVTY | -5.0890208 | 3.17135368 | 4 | -6.9283293 | 1.6554784 | 6 | 97.1 | 99.2 | 2.0 |
| NTYPIKLFY | -5.2140849 | 1.53384215 | 6 | -7.3996355 | 1.31593593 | 3 | 97.4 | 99.4 | 2.0 |
| HDNANGGQNGTVQEIMIPAGKAGL | -3.579591 | 1.32179812 | 2 | -4.0522432 | 0.10691155 | 2 | 92.3 | 94.3 | 2.0 |
| LFLEPNPEDPLNKEAAEVL | -2.4693732 | 0.18319556 | 4 | -2.7089856 | 0.40604974 | 6 | 84.7 | 86.7 | 2.0 |
| QKALDLDSSCKEAADGY | -4.9879526 | 0.75082556 | 2 | -6.5836404 | 2.83356391 | 2 | 96.9 | 99.0 | 2.0 |
| QASDLPASASLPAKADLLL | -4.8401171 | 0.46038857 | 2 | -6.1899648 | 0.90271515 | 2 | 96.6 | 98.6 | 2.0 |

| | | | | | | | | | |
|---|---|---|---|---|---|---|---|---|---|
| GFIERGDVVKEIF | -4.8324605 | 2.18358383 | 3 | -6.168153 | 1.61040782 | 5 | 96.6 | 98.6 | 2.0 |
| LANIEQQHGNSGRNSESESNKVAAETQSPSLF | -2.0677051 | 0.24010644 | 4 | -2.2627624 | 0.10139812 | 2 | 80.7 | 82.8 | 2.0 |
| SAQLREAATQAQQTLGSTIDKATGILLY | -2.9316021 | 0.52475204 | 5 | -3.239577 | 0.2136116 | 2 | 88.4 | 90.4 | 2.0 |
| SGAQIKIANPVEGSTDRQVTITGSAASISLAQY | -2.2729505 | 0.33457681 | 7 | -2.4874066 | 0.18268385 | 5 | 82.9 | 84.9 | 2.0 |
| KGMQELGVHPDQETY | -4.1555607 | 0.56542321 | 2 | -4.8702393 | 0.38293676 | 2 | 94.7 | 96.7 | 2.0 |
| KLEDTENWLYEDGEDQPKQVY | -4.7982549 | 1.63842376 | 2 | -6.0730542 | 0.63527948 | 3 | 96.5 | 98.5 | 2.0 |
| CVDADWDTKHVL | -3.792613 | 0.52809638 | 3 | -4.3333964 | 4.21907816 | 6 | 93.3 | 95.3 | 2.0 |
| CAATGPSIKIW | -4.4457299 | 1.38079347 | 5 | -5.356059 | 0.48035091 | 3 | 95.6 | 97.6 | 2.0 |
| SIDISSKQVENAGAIGPSRF | -3.7707883 | 1.00302701 | 2 | -4.3004288 | 0.46606421 | 5 | 93.2 | 95.2 | 2.0 |
| GAILNLVPLAESVVKL | -1.8779061 | 1.50506788 | 6 | -2.0553605 | 1.963905 | 3 | 78.6 | 80.6 | 2.0 |
| KAVTELNEPLSNEDRNLL | -5.2304563 | 0.95021239 | 7 | -7.3637805 | 0.53976836 | 2 | 97.4 | 99.4 | 2.0 |
| KTIEEVVGRANNSTY | -4.1871759 | 1.10855288 | 7 | -4.9128926 | 0.38705389 | 5 | 94.8 | 96.8 | 2.0 |
| AVHQEGNKNVGLDIEAEVPAVKDMLEAGILDTY | -4.7880048 | 1.37715449 | 2 | -6.0340926 | 1.08638896 | 2 | 96.5 | 98.5 | 2.0 |
| AADAEPLEIILHLPLLCEDKNVPY | -2.5659504 | 0.52083262 | 2 | -2.8123342 | 0.61545283 | 5 | 85.6 | 87.5 | 2.0 |
| FDKRDNSDFDLL | -4.8105556 | 0.68881358 | 4 | -6.0805349 | 0.14692215 | 2 | 96.6 | 98.5 | 2.0 |
| SLPKDFWEQEVRDIRSY | -2.8177517 | 0.60248505 | 3 | -3.1007632 | 0.04436659 | 2 | 87.6 | 89.6 | 2.0 |
| GMTPSKGVLF | -5.3021775 | 0.67456652 | 4 | -7.658926 | 2.91505537 | 4 | 97.5 | 99.5 | 2.0 |
| KIEGVYARDETEFY | -5.2778229 | 1.61523885 | 6 | -7.5420163 | 0.44700479 | 2 | 97.5 | 99.5 | 2.0 |
| AGIDSSSPEVKGY | -4.8943829 | 1.95224189 | 3 | -6.2757347 | 0.13505881 | 2 | 96.7 | 98.7 | 2.0 |
| VGGIKEDTEEHHLRDYFEQY | -2.9868292 | 0.69727128 | 2 | -3.2971034 | 0.28574714 | 2 | 88.8 | 90.8 | 2.0 |
| VGGLNPEATEEKIREY | -5.0609268 | 2.02906045 | 8 | -6.7057178 | 1.94739902 | 7 | 97.1 | 99.1 | 2.0 |
| TEAPLNPKANRE | -4.6198979 | 0.79673971 | 4 | -5.6532184 | 1.24421322 | 5 | 96.1 | 98.1 | 2.0 |
| GVPIIVQASQAEKN | -4.586105 | 0.63449747 | 5 | -5.5857792 | 1.34504221 | 3 | 96.0 | 98.0 | 2.0 |
| ISDKDASIVGFFDDSFSEAHSEF | -3.2537069 | 0.36555717 | 9 | -3.6175727 | 0.46260823 | 3 | 90.5 | 92.5 | 2.0 |
| TLDVTTGQRKY | -5.3424446 | 1.43259155 | 4 | -7.789013 | 1.44301716 | 3 | 97.6 | 99.5 | 2.0 |
| MIEIAGVKLLY | -4.2618694 | 0.35691087 | 2 | -5.0145543 | 1.10118385 | 5 | 95.0 | 97.0 | 2.0 |
| GSRSQKELPTEPPYTAY | -5.1655142 | 0.60470571 | 9 | -7.0170727 | 1.90464049 | 7 | 97.3 | 99.2 | 1.9 |
| AFVTFDDHDSVDKIVIQKY | -5.1371687 | 3.24150558 | 9 | -6.9198173 | 1.56676039 | 11 | 97.2 | 99.2 | 1.9 |
| KNSQGEEVAQRSTVF | -5.2287615 | 0.89427403 | 11 | -7.2445305 | 2.77594635 | 4 | 97.4 | 99.3 | 1.9 |
| VYPPDYNPEGKVT | -4.9217572 | 1.75764013 | 3 | -6.298278 | 1.71832028 | 6 | 96.8 | 98.7 | 1.9 |
| FVRNLATTVTEEILEKSFSEF | -2.7044857 | 0.05528805 | 3 | -2.9635471 | 0.00938981 | 2 | 86.7 | 88.6 | 1.9 |
| INIVVIGHVDSGKSTT | -4.9041625 | 1.71395661 | 4 | -6.2447829 | 0.97806443 | 5 | 96.8 | 98.7 | 1.9 |
| NMNEEEVEKRF | -3.9578348 | 0.53335134 | 2 | -4.541206 | 1.19979536 | 3 | 94.0 | 95.9 | 1.9 |
| LNAQESAKSARL | -4.3986312 | 1.79564461 | 3 | -5.2257568 | 0.82904336 | 2 | 95.5 | 97.4 | 1.9 |
| VQSLKEIVINVPEQSAVTL | -3.0315018 | 0.25843288 | 9 | -3.3420463 | 0.26908814 | 4 | 89.1 | 91.0 | 1.9 |
| TFPRPVTVEPM(15.9949)DQLDDEEGLPEKL | -2.5064755 | 0.69468715 | 2 | -2.7365908 | 0.17269348 | 3 | 85.0 | 87.0 | 1.9 |
| KTIGYDPIISPEVSASF | -5.0151244 | 2.90491694 | 8 | -6.5140923 | 1.50444455 | 5 | 97.0 | 98.9 | 1.9 |
| TSEIGTKQITQSAL | -5.085285 | 1.54018658 | 3 | -6.7010425 | 1.38290383 | 2 | 97.1 | 99.0 | 1.9 |
| QLELSKVREEFKEL | -4.8266017 | 2.22883513 | 2 | -6.0366977 | 0.35977586 | 2 | 96.6 | 98.5 | 1.9 |
| IIGADTVMNESEDLFSLIETHEAKPL | -2.0626957 | 0.17069704 | 3 | -2.2458403 | 0.20197421 | 2 | 80.7 | 82.6 | 1.9 |
| SPDGSKIAAGSADRF | -5.4822799 | 2.22839807 | 8 | -8.4371874 | 2.05649817 | 4 | 97.8 | 99.7 | 1.9 |
| GILSDDVETDTVAPGENLKIRL | -3.0272203 | 0.28693261 | 6 | -3.3329929 | 0.48122932 | 3 | 89.1 | 91.0 | 1.9 |
| ADAHKERTNEGVIEF | -5.3791693 | 0.56614356 | 5 | -7.7919535 | 2.14024498 | 4 | 97.7 | 99.6 | 1.9 |
| SEAREDMAALEKDYEEVGVDSVEGEGEEEGEEY | -2.3542206 | 0.13945754 | 7 | -2.5640706 | 0.18973143 | 3 | 83.6 | 85.5 | 1.9 |
| LTVKDNQVVQLHPSTVL | -5.4006092 | 1.57865005 | 2 | -7.8664943 | 3.18314576 | 3 | 97.7 | 99.6 | 1.9 |
| IIEELNVRKVTL | -5.0608776 | 1.64611501 | 2 | -6.5818009 | 1.98715189 | 3 | 97.1 | 99.0 | 1.9 |
| IAQLEEELEEEQGNTELINDRLKKANL | -4.4851254 | 1.11678063 | 2 | -5.3461879 | 0.19887318 | 2 | 95.7 | 97.6 | 1.9 |
| KLADDVDLEQVANETHGHVGADLAAL | -2.5215313 | 0.34001062 | 3 | -2.7478059 | 0.29192042 | 4 | 85.2 | 87.0 | 1.9 |
| VAASSDIGDPLRELGSECGIEFDEEKTAVIDHHNY | -1.7310943 | 0.09487036 | 5 | -1.8876299 | 0.11228106 | 2 | 76.9 | 78.7 | 1.9 |
| DPVPVGVTKVIL | -5.1778388 | 6.52553653 | 4 | -6.9280948 | 1.06030939 | 4 | 97.3 | 99.2 | 1.9 |
| CIEVTPQSKIAW | -4.8508933 | 0.69363179 | 5 | -6.0608951 | 0.97802166 | 4 | 96.7 | 98.5 | 1.9 |
| EALELRDNDKTRY | -4.6880412 | 1.44852862 | 5 | -5.7181319 | 0.75243315 | 7 | 96.3 | 98.1 | 1.9 |
| AKSSVGETVYGGSDELSDDITQQQLLPGVKDPNL | -5.3344432 | 0.48923142 | 2 | -7.5023291 | 0.71756984 | 2 | 97.6 | 99.5 | 1.9 |
| GEGKSTTTIGLVQALGAHLY | -4.3822497 | 0.66963256 | 6 | -5.1671086 | 1.470103 | 2 | 95.4 | 97.3 | 1.9 |
| AEIAKAELDDTPM | -5.0818965 | 0.34354061 | 2 | -6.6289869 | 1.68762331 | 3 | 97.1 | 99.0 | 1.9 |
| RSTLEPVEKAL | -4.4501673 | 1.50483087 | 2 | -5.2791396 | 1.2073914 | 2 | 95.6 | 97.5 | 1.9 |
| SADIKDSKAYF | -5.1895251 | 1.94221608 | 8 | -6.9459778 | 2.09054196 | 5 | 97.3 | 99.2 | 1.9 |
| KFIDTTSKF | -5.5688211 | 1.01985627 | 2 | -8.935553 | 1.9379063 | 2 | 97.9 | 99.8 | 1.9 |
| QAKLAEQAERY | -5.2968782 | 1.07737844 | 3 | -7.3202651 | 1.81194949 | 5 | 97.5 | 99.4 | 1.9 |
| GAEVKAEPVEVVAPR | -4.8998714 | 2.1336561 | 7 | -6.1548031 | 1.28631275 | 3 | 96.8 | 98.6 | 1.9 |
| VTRTQGTKIASDGL | -4.7219318 | 1.27086272 | 7 | -5.7741925 | 0.63196385 | 4 | 96.3 | 98.2 | 1.9 |
| IAQQTDTSDPEKVVSAFL | -5.1140001 | 1.12179459 | 3 | -6.7039068 | 2.25279581 | 4 | 97.2 | 99.0 | 1.9 |
| ELQASRVSSDVIDQKVY | -5.6166952 | 1.6720667 | 3 | -9.4053319 | 0.65222654 | 4 | 98.0 | 99.9 | 1.9 |
| KLITEDVQGKNCL | -5.142001 | 1.93094596 | 8 | -6.7589669 | 1.33195636 | 5 | 97.2 | 99.1 | 1.8 |
| KTVTISDHGTVTY | -5.1061528 | 1.41279508 | 6 | -6.6547624 | 1.52969577 | 3 | 97.2 | 99.0 | 1.8 |
| MIEQSGPPSKEIL | -3.8434115 | 0.53521023 | 12 | -4.3493651 | 0.30394236 | 5 | 93.5 | 95.3 | 1.8 |
| MTEDNKDLIQGKDLL | -4.1013483 | 1.07530354 | 5 | -4.7142841 | 1.61561384 | 3 | 94.5 | 96.3 | 1.8 |
| QAVAQRDPIIAKQLF | -5.2222041 | 2.13630528 | 3 | -7.0033434 | 2.22684059 | 5 | 97.4 | 99.2 | 1.8 |
| QAVAQRDPIIAKQLF | -4.2164043 | 1.64855313 | 3 | -4.8860567 | 0.18346 | 2 | 94.9 | 96.7 | 1.8 |
| YFPVKNVIDGDLCEQF | -3.4176458 | 0.46154345 | 7 | -3.7926443 | 1.66865683 | 2 | 91.4 | 93.3 | 1.8 |
| SSSLEKSYELPDGQVITIGNERF | -2.9096961 | 0.3168078 | 3 | -3.1830639 | 0.05910404 | 2 | 88.3 | 90.1 | 1.8 |
| ESVAAKNLQEAEEWY | -5.0798168 | 1.69754488 | 4 | -6.5603617 | 2.86303454 | 6 | 97.1 | 99.0 | 1.8 |

| | | | | | | | | | |
|---|---|---|---|---|---|---|---|---|---|
| KMVVESAYEVIKL | -5.5084078 | 0.87271163 | 3 | -8.2153833 | 2.47544087 | 2 | 97.9 | 99.7 | 1.8 |
| TQAPGNPVLAVQINQDKNF | -5.5853427 | 1.57482721 | 6 | -8.7807021 | 1.67595924 | 2 | 98.0 | 99.8 | 1.8 |
| SKFGEVVDCTL | -5.4239891 | 2.16152626 | 4 | -7.74306 | 3.28921544 | 4 | 97.7 | 99.5 | 1.8 |
| SATLGLVDIVKGTNSYY | -4.1388929 | 1.03653791 | 11 | -4.7592953 | 1.3649549 | 6 | 94.6 | 96.4 | 1.8 |
| HGLDEEAEQKLF | -4.4258878 | 0.8355577 | 2 | -5.207289 | 0.57121591 | 5 | 95.6 | 97.4 | 1.8 |
| LLGMLGAESAKL | -4.6672294 | 1.81077052 | 3 | -5.6306548 | 1.69471568 | 5 | 96.2 | 98.0 | 1.8 |
| LLQYIQEEIPDYRNAVIVAKSPA | -3.8415012 | 0.82137494 | 4 | -4.3376263 | 0.37982016 | 3 | 93.5 | 95.3 | 1.8 |
| AQDVRKESPLLF | -4.4505851 | 0.45595894 | 5 | -5.2441664 | 0.88486416 | 2 | 95.6 | 97.4 | 1.8 |
| NIPLVSDPKRTIAQDY | -4.7994066 | 2.1761506 | 2 | -5.8840126 | 3.07189757 | 4 | 96.5 | 98.3 | 1.8 |
| HNEAQVNPERKNL | -5.4059965 | 1.13758559 | 6 | -7.6044192 | 1.47343626 | 6 | 97.7 | 99.5 | 1.8 |
| LWKDLTLDQAY | -4.4835793 | 1.118121 | 6 | -5.2934584 | 1.33980527 | 4 | 95.7 | 97.5 | 1.8 |
| SLLATEDKEAL | -5.4318969 | 1.5630308 | 5 | -7.7138519 | 2.42014497 | 3 | 97.7 | 99.5 | 1.8 |
| EVAWEVANKVGGIY | -4.8004692 | 2.37026306 | 2 | -5.8755216 | 1.53422752 | 2 | 96.5 | 98.3 | 1.8 |
| SAELAKVINDGLF | -4.9647886 | 1.84654792 | 5 | -6.2308172 | 2.05185508 | 4 | 96.9 | 98.7 | 1.8 |
| VIVGDGACGKTCL | -4.1298648 | 1.14624378 | 4 | -4.7360469 | 0.88631074 | 3 | 94.6 | 96.4 | 1.8 |
| GTGEDVKVIL | -4.8008384 | 1.45999035 | 2 | -5.8730254 | 1.79465045 | 2 | 96.5 | 98.3 | 1.8 |
| CFLEYEDHKSAAQ | -5.6649312 | 0.88686847 | 3 | -9.2876261 | 1.30030299 | 4 | 98.1 | 99.8 | 1.8 |
| SAKNLRDIDEVSSLL | -4.6681268 | 0.88602952 | 5 | -5.6060913 | 1.66648353 | 4 | 96.2 | 98.0 | 1.8 |
| VEKLTEVSISSDAFFPF | -4.0996896 | 0.09843035 | 2 | -4.6856429 | 0.63396875 | 5 | 94.5 | 96.3 | 1.8 |
| TGINQTGDQSLPSKPSSTL | -4.9126907 | 1.42911732 | 8 | -6.0943159 | 2.66859913 | 4 | 96.8 | 98.6 | 1.8 |
| LVKEAGLNVTTSHSPAAPGEQGFGECL | -3.3848076 | 0.42350594 | 2 | -3.7388387 | 0.22778981 | 2 | 91.3 | 93.0 | 1.8 |
| ASSSSLEKSYELPDGQVITIGNERF | -3.0779836 | 0.22358087 | 9 | -3.3693662 | 0.05734363 | 3 | 89.4 | 91.2 | 1.8 |
| DLSKIGEEQSPEDAEDGPPELLF | -3.4987548 | 0.62569787 | 5 | -3.8787613 | 0.70326361 | 6 | 91.9 | 93.6 | 1.8 |
| ELSENDLNFIKQSKDGAGF | -5.3509754 | 1.53988578 | 6 | -7.2991718 | 1.45841545 | 3 | 97.6 | 99.4 | 1.8 |
| TNHHFYDESKPF | -5.5876789 | 1.33850502 | 6 | -8.4555421 | 1.07732099 | 2 | 98.0 | 99.7 | 1.8 |
| LINTEGAIKLADF | -3.8840024 | 1.26831552 | 4 | -4.3769935 | 1.25628898 | 3 | 93.7 | 95.4 | 1.8 |
| VRNLATTVTEEILEKSF | -4.5260036 | 1.77899544 | 4 | -5.3361066 | 1.88234563 | 2 | 95.8 | 97.6 | 1.7 |
| KNVTELNEPLSNEERNLL | -2.9163391 | 0.30285761 | 2 | -3.1759364 | 0.16854358 | 3 | 88.3 | 90.0 | 1.7 |
| CTGQIKTGAPC | -1.2217942 | 0.44347655 | 12 | -1.3423478 | 0.50132129 | 6 | 70.0 | 71.7 | 1.7 |
| ENERGELANEVKVL | -5.1334028 | 0.8308426 | 2 | -6.5640261 | 1.3852578 | 2 | 97.2 | 99.0 | 1.7 |
| LAVDYENVRPDIVLLGKAL | -0.7867236 | 0.77429174 | 7 | -0.8946481 | 0.21107655 | 5 | 63.3 | 65.0 | 1.7 |
| ATIKEQNGDVKEAASILQELQVETY | -4.6938576 | 1.54186273 | 6 | -5.6134071 | 1.34031598 | 2 | 96.3 | 98.0 | 1.7 |
| DFTGAVEDISKIPEQSVL | -4.5676104 | 0.48144031 | 4 | -5.3865011 | 0.65677058 | 7 | 96.0 | 97.7 | 1.7 |
| KVPSTEAEALASSLM | -5.3371479 | 0.56433977 | 2 | -7.1426654 | 1.51033624 | 5 | 97.6 | 99.3 | 1.7 |
| ELISETGGSHDKRF | -4.6749834 | 1.54037344 | 5 | -5.5730321 | 2.08960713 | 7 | 96.2 | 97.9 | 1.7 |
| NQKIRDGWQVEEADDWLRY | -3.1784017 | 0.50828289 | 3 | -3.4770956 | 0.12693582 | 3 | 90.1 | 91.8 | 1.7 |
| QEFTDHLVKTH | -4.4853489 | 0.16138039 | 2 | -5.2438735 | 1.057567 | 4 | 95.7 | 97.4 | 1.7 |
| KTSEQLDQPISACCF | -2.9418896 | 0.09692766 | 5 | -3.1998587 | 0.13524016 | 4 | 88.5 | 90.2 | 1.7 |
| KVFSDEVQQQAQL | -5.2301194 | 1.08723309 | 7 | -6.7907345 | 1.70177139 | 9 | 97.4 | 99.1 | 1.7 |
| EALDTILPPTRPTDKPL | -3.3369292 | 2.23553231 | 5 | -3.6649395 | 0.89914731 | 2 | 91.0 | 92.7 | 1.7 |
| AFVTFDDHDSVDKIVIQ | -3.709058 | 0.76620031 | 3 | -4.1279168 | 0.36703449 | 2 | 92.9 | 94.6 | 1.7 |
| VTGVHEEATEEDIHDKFAEYGEIKNIHL | -5.5305002 | 1.40619763 | 2 | -7.8680081 | 0.80144426 | 2 | 97.9 | 99.6 | 1.7 |
| FEAIQKGTGARPY | -5.4958235 | 0.71639887 | 9 | -7.6755366 | 2.18814935 | 5 | 97.8 | 99.5 | 1.7 |
| ISNSSDALDKIRY | -4.6179733 | 2.50431316 | 4 | -5.452347 | 1.1383311 | 5 | 96.1 | 97.8 | 1.7 |
| VGKSEEQLKEEGIEY | -5.4538301 | 1.59011198 | 7 | -7.4934834 | 2.18930217 | 4 | 97.8 | 99.4 | 1.7 |
| SSEARVRDVVTKY | -4.4208479 | 0.97854928 | 2 | -5.1270139 | 1.20557163 | 3 | 95.5 | 97.2 | 1.7 |
| SDKLFEMVLGPAAY | -4.8802381 | 0.55446642 | 6 | -5.9371144 | 0.04437447 | 2 | 96.7 | 98.4 | 1.7 |
| YFDPKQTDVLQQL | -4.8037914 | 1.57177268 | 4 | -5.7854521 | 1.56762543 | 4 | 96.5 | 98.2 | 1.7 |
| VFLGLDNAGKTTLL | 0.37214775 | 0.26484438 | 3 | 0.27427686 | 1.37218959 | 3 | 43.6 | 45.3 | 1.7 |
| GPAAPAAANSSGDGGAAGDGTVVDCPVCKQQ( | -2.8350726 | 0.15482531 | 6 | -3.0735906 | 0.43603846 | 3 | 87.7 | 89.4 | 1.7 |
| KATAAGVKQTESTSF | -5.6075407 | 2.21728842 | 9 | -8.2023373 | 1.93388104 | 7 | 98.0 | 99.7 | 1.7 |
| SSAQADFNQLAELDRQIKSF | -2.3755307 | 0.19865161 | 5 | -2.5612534 | 0.4651225 | 2 | 83.8 | 85.5 | 1.7 |
| NIGKLAQF | -4.9027444 | 1.08514991 | 2 | -5.9722279 | 2.68604455 | 2 | 96.8 | 98.4 | 1.7 |
| NSGKVDIVAINDPFIDLNYM(15.9949)VY | -2.9458975 | 0.36110866 | 9 | -3.1987731 | 0.39257085 | 9 | 88.5 | 90.2 | 1.7 |
| DLSKIGEEQSAEDAEDGPPELL | -3.3989923 | 0.55063844 | 4 | -3.7330927 | 0.53838083 | 2 | 91.3 | 93.0 | 1.7 |
| AVTKDGRVF | -4.428489 | 0.83280104 | 5 | -5.1283075 | 1.91350883 | 3 | 95.6 | 97.2 | 1.7 |
| TKDEPNSTPEKTEQFY | -5.5289462 | 0.5976707 | 3 | -7.7369251 | 1.94826442 | 3 | 97.9 | 99.5 | 1.7 |
| NAGGSVKDRISL | -0.3013431 | 1.45175699 | 4 | -0.3981703 | 0.89799035 | 4 | 55.2 | 56.9 | 1.7 |
| GDPVVQSDMKHWPF | -3.594814 | 0.54721921 | 9 | -3.9718453 | 2.39334628 | 2 | 92.4 | 94.0 | 1.7 |
| TLLLGQDENSVIKSF | -4.554809 | 0.51671945 | 3 | -5.3275711 | 1.42230059 | 4 | 95.9 | 97.6 | 1.7 |
| YPEDVAEELIQDITQKLF | -3.304748 | 0.95696268 | 2 | -3.6141122 | 0.45381357 | 2 | 90.8 | 92.4 | 1.6 |
| APADVTSEKDVQTAL | -4.0567095 | 0.47329551 | 2 | -4.5731863 | 0.5370166 | 4 | 94.3 | 96.0 | 1.6 |
| ATASHRNTEEEGLKY | -5.5517334 | 1.09213512 | 7 | -7.7776094 | 1.37330013 | 5 | 97.9 | 99.5 | 1.6 |
| LIKTVETRDGQVINETSQHHDDLE | -3.4717278 | 0.27591645 | 6 | -3.8140419 | 0.30420356 | 2 | 91.7 | 93.4 | 1.6 |
| NTLGVCKY | -5.2323847 | 2.1930313 | 2 | -6.6830025 | 1.27491135 | 2 | 97.4 | 99.0 | 1.6 |
| IAGKVAVVAGY | -4.9071823 | 1.21466267 | 3 | -5.9443314 | 0.05213296 | 2 | 96.8 | 98.4 | 1.6 |
| IGGLPNYLNDDQVKELL | -5.4951298 | 1.07145291 | 3 | -7.5097323 | 2.67218016 | 3 | 97.8 | 99.5 | 1.6 |
| KVGSAADIPINISETDLSLL | -4.5064982 | 0.99322761 | 8 | -5.2317561 | 1.16843139 | 5 | 95.8 | 97.4 | 1.6 |
| LDGIYVSEKGTVQQADE | -5.2040849 | 3.34539105 | 3 | -6.6004282 | 2.54558867 | 4 | 97.4 | 99.0 | 1.6 |
| YNEATGGKY | -4.8661862 | 1.22677848 | 3 | -5.8537794 | 1.33506679 | 3 | 96.7 | 98.3 | 1.6 |
| SWLVKIPSEQEQL | -4.537444 | 0.97624952 | 3 | -5.2746668 | 1.56433935 | 2 | 95.9 | 97.5 | 1.6 |
| AANTKGICF | -5.5232453 | 2.03889337 | 8 | -7.5775423 | 1.59944814 | 4 | 97.9 | 99.5 | 1.6 |

| | | | | | | | | | |
|---|---|---|---|---|---|---|---|---|---|
| STQDHAAAAIAKAGIPVYAW | -4.3697384 | 0.98350311 | 4 | -5.0113762 | 0.7861909 | 2 | 95.4 | 97.0 | 1.6 |
| NVLYEKEGEFVAQF | -4.5015539 | 1.11010348 | 8 | -5.2150706 | 1.40689167 | 9 | 95.8 | 97.4 | 1.6 |
| GYPNLKSVNELIY | -5.2135164 | 2.62078269 | 4 | -6.6014442 | 1.95352885 | 7 | 97.4 | 99.0 | 1.6 |
| HVDISLTDFIQKY | -4.5188257 | 0.93476477 | 9 | -5.2414248 | 1.52498135 | 3 | 95.8 | 97.4 | 1.6 |
| MIEMDGTENKSKF | -5.4036481 | 1.58255386 | 9 | -7.1351936 | 1.86805934 | 6 | 97.7 | 99.3 | 1.6 |
| ESSFSNEEAQKAQAL | -5.4587019 | 3.34014414 | 4 | -7.3201299 | 2.32730681 | 2 | 97.8 | 99.4 | 1.6 |
| EVKATAGDTHLGGEDFDNRLVNHFVEEF | -2.3317825 | 0.06884605 | 5 | -2.5055859 | 0.28568939 | 5 | 83.4 | 85.0 | 1.6 |
| DLGGGTFDISILEIQKGVF | -4.1201423 | 1.16722246 | 9 | -4.6458631 | 1.17932856 | 5 | 94.6 | 96.2 | 1.6 |
| IAIASTTVETTDPEKEVEPALELLEPIDQKF | -4.9839836 | 1.92913446 | 4 | -6.0701548 | 1.38009092 | 3 | 96.9 | 98.5 | 1.6 |
| VKELSEAL | -5.7833446 | 0.92699882 | 4 | -9.0553782 | 2.04458196 | 2 | 98.2 | 99.8 | 1.6 |
| ASICQQNGIVPIVEPEILPDGDHDLKRCQY | -2.7351517 | 0.63797011 | 7 | -2.9490388 | 0.72399868 | 4 | 86.9 | 88.5 | 1.6 |
| EGSRTQEEIVAKVREVSQPDWTPPPEVTL | -2.9061334 | 0.30238713 | 4 | -3.1417372 | 0.1389494 | 3 | 88.2 | 89.8 | 1.6 |
| GLVKELLSNVDEGIY | -3.9225151 | 1.40223531 | 4 | -4.3747562 | 0.63155069 | 2 | 93.8 | 95.4 | 1.6 |
| GLPTSDEQKKQEIL | -5.4438863 | 0.78361869 | 6 | -7.2385175 | 1.47073275 | 6 | 97.8 | 99.3 | 1.6 |
| AIGDILEDKVELTPVAIQAGRLL | -2.004624 | 0.22047533 | 4 | -2.1522595 | 0.08273668 | 2 | 80.1 | 81.6 | 1.6 |
| NEATGGKYVPRAVL | -5.0884786 | 0.60281818 | 4 | -6.2773439 | 1.16322113 | 2 | 97.1 | 98.7 | 1.6 |
| DVVRKEAESCDCL | -5.3422804 | 0.00827704 | 2 | -6.9103685 | 1.7235181 | 4 | 97.6 | 99.2 | 1.6 |
| VEVQLLESKTY | -5.0582965 | 1.01425537 | 4 | -6.2077758 | 1.11357066 | 4 | 97.1 | 98.7 | 1.6 |
| KDLVKVIQQESY | -4.9530363 | 1.17233767 | 4 | -5.9885773 | 2.84597351 | 4 | 96.9 | 98.4 | 1.6 |
| GAFVKPAVVTVGDFPEEDYGLDEI | -3.4710044 | 0.32192535 | 6 | -3.80031 | 0.19932788 | 6 | 91.7 | 93.3 | 1.6 |
| NLVGEKLQWF | -4.9163581 | 1.12830531 | 4 | -5.9130426 | 2.32973844 | 4 | 96.8 | 98.4 | 1.6 |
| SLRTVSLGAGAKDEL | -5.0566945 | 3.23311763 | 2 | -6.1953642 | 1.2130703 | 2 | 97.1 | 98.7 | 1.6 |
| VIKDLVPDLSNF | 2.31860849 | 0.17776595 | 12 | 2.16189793 | 0.26678892 | 6 | 16.7 | 18.3 | 1.6 |
| GDRYFDPANGKF | -5.2599063 | 0.33767582 | 3 | -6.6626991 | 2.13723466 | 4 | 97.5 | 99.0 | 1.6 |
| CKASAELALGENSEVL | -4.0500791 | 0.07636456 | 2 | -4.5370278 | 0.56220329 | 2 | 94.3 | 95.9 | 1.6 |
| LVLQEILESEEKGDPNKPSGF | -4.6720823 | 0.22321782 | 2 | -5.4634642 | 1.53182893 | 3 | 96.2 | 97.8 | 1.6 |
| IAIASTTVETTDPEKEVEPALELLEPIDQKF | -5.2731384 | 2.37054739 | 5 | -6.6849609 | 0.91917818 | 3 | 97.5 | 99.0 | 1.6 |
| NTHADFADECPKPELL | -5.0121363 | 1.06549811 | 5 | -6.0854353 | 1.40760477 | 5 | 97.0 | 98.5 | 1.6 |
| TIMFGPDKCGEDY | -5.1701468 | 1.13615047 | 4 | -6.4261895 | 0.41987146 | 3 | 97.3 | 98.9 | 1.6 |
| SGNPIKVSF | -5.2569105 | 1.74153411 | 9 | -6.6293833 | 1.43912993 | 4 | 97.5 | 99.0 | 1.5 |
| SDSFGGDAQADEGQARKSQL | -5.4246313 | 0.36510329 | 4 | -7.0860883 | 0.61156555 | 3 | 97.7 | 99.3 | 1.5 |
| CQLILDPIFKVF | -5.3690811 | 0.06913131 | 2 | -6.9130098 | 1.2469621 | 4 | 97.6 | 99.2 | 1.5 |
| VKGLSEDTTEETLKESFDGSVR | -5.1922182 | 1.80833334 | 7 | -6.4548384 | 1.14224164 | 4 | 97.3 | 98.9 | 1.5 |
| APVISAEKAY | -4.6249986 | 1.14634305 | 4 | -5.3687119 | 1.14186353 | 8 | 96.1 | 97.6 | 1.5 |
| IVAAGVGEFEAGISKN | -4.0498853 | 0.90713136 | 9 | -4.5226205 | 1.37408775 | 8 | 94.3 | 95.8 | 1.5 |
| KPVFVTEITDDLHFY | -5.2055134 | 0.14107253 | 4 | -6.4679464 | 0.48852422 | 3 | 97.4 | 98.9 | 1.5 |
| MGLDCGPESSKKY | -5.8959323 | 2.40346968 | 8 | -9.5782853 | 0.69524075 | 5 | 98.3 | 99.9 | 1.5 |
| IYPAQNPELLNKL | -4.5182743 | 0.66742103 | 3 | -5.1926451 | 1.01020748 | 3 | 95.8 | 97.3 | 1.5 |
| YVPPEQAKVVASVH | -3.8763093 | 0.80477077 | 5 | -4.2919854 | 3.48731769 | 2 | 93.6 | 95.1 | 1.5 |
| VKGLSEDTTEETLKESF | -5.4189979 | 2.08046487 | 6 | -7.0179053 | 2.24934691 | 7 | 97.7 | 99.2 | 1.5 |
| ELLEGKVLPGVDALSNI | -5.6179116 | 1.56293907 | 6 | -7.7009779 | 1.53805725 | 4 | 98.0 | 99.5 | 1.5 |
| SSIEREEYGKLF | -4.9683713 | 0.87101116 | 3 | -5.9618563 | 0.16324016 | 2 | 96.9 | 98.4 | 1.5 |
| EFHLPLSPEELLKSGGVNQY | -3.2024528 | 0.48595775 | 2 | -3.468057 | 0.45312456 | 2 | 90.2 | 91.7 | 1.5 |
| TLQDKAQVADVVVSRW | -4.7380909 | 0.97590844 | 4 | -5.541032 | 0.2148208 | 3 | 96.4 | 97.9 | 1.5 |
| DQLALDSPKGCGTVL | -4.5065 | 1.34829727 | 5 | -5.1671065 | 2.51336486 | 5 | 95.8 | 97.3 | 1.5 |
| TKSPYQEF | -5.4067599 | 0.42594687 | 3 | -6.9581604 | 0.73849816 | 2 | 97.7 | 99.2 | 1.5 |
| SLNSKVVSQY | -5.1891395 | 1.01728416 | 11 | -6.4076705 | 1.7800756 | 7 | 97.3 | 98.8 | 1.5 |
| SPNSSNPIIVSCGWDKL | -3.0037661 | 0.63544821 | 7 | -3.2381101 | 0.27808111 | 5 | 88.9 | 90.4 | 1.5 |
| AIRNDEELNKL | -5.4287585 | 3.9796521 | 8 | -7.0165269 | 2.14356521 | 6 | 97.7 | 99.2 | 1.5 |
| QLDPTASISAKVNNSSL | -5.6124659 | 1.25617129 | 6 | -7.629258 | 1.15973621 | 3 | 98.0 | 99.5 | 1.5 |
| VGDLHPDVTEAMLYEKF | -3.1879086 | 0.16103739 | 4 | -3.4491104 | 0.12904439 | 2 | 90.1 | 91.6 | 1.5 |
| KAVQAQGGESQQEAQRL | -5.4348106 | 2.8181567 | 5 | -7.0252311 | 4.20591685 | 3 | 97.7 | 99.2 | 1.5 |
| AQDCPLRTPEDFVFPLLPNEELREKY | -2.2729964 | 0.61401574 | 2 | -2.4305804 | 0.64765455 | 2 | 82.9 | 84.4 | 1.5 |
| NVNDNSEYVETIIAKCIDHY | -3.8807631 | 0.20817507 | 3 | -4.2905914 | 0.56858466 | 2 | 93.6 | 95.1 | 1.5 |
| QQIDSVANADIINAAKKF | -5.6905058 | 1.36688711 | 9 | -7.9432783 | 1.78771513 | 5 | 98.1 | 99.6 | 1.5 |
| TVGKEPFPTIY | -5.1200151 | 1.96604841 | 10 | -6.2425558 | 1.15311563 | 3 | 97.2 | 98.7 | 1.5 |
| TSFIGAIAIGDLVKSTL | -5.5706469 | 1.99200392 | 4 | -7.4331852 | 1.20128062 | 7 | 97.9 | 99.4 | 1.5 |
| VWVPSDKSGFEPASL | -4.6307897 | 1.35305005 | 9 | -5.3481246 | 0.81421667 | 5 | 96.1 | 97.6 | 1.5 |
| LSSAVDHGSDEVKF | -5.2288413 | 0.44517953 | 3 | -6.4652303 | 0.68980056 | 4 | 97.4 | 98.9 | 1.5 |
| AVSEGTKAVTKY | -5.8891223 | 1.38296843 | 16 | -9.0945298 | 1.46482994 | 3 | 98.3 | 99.8 | 1.5 |
| KDM(15.9949)AIATGGAVFGEEGLTL | -2.0681327 | 0.20723256 | 5 | -2.2093443 | 0.11811461 | 2 | 80.7 | 82.2 | 1.5 |
| FVRNLANTVTEEILEKAF | -2.6118376 | 0.25992144 | 4 | -2.796088 | 0.24717227 | 2 | 85.9 | 87.4 | 1.5 |
| HFEPNEYFTNEVLTKTY | -0.8017159 | 0.54532121 | 7 | -0.8942724 | 0.15470259 | 3 | 63.5 | 65.0 | 1.5 |
| LTDSNNIKEVL | -5.7003018 | 0.37320725 | 3 | -7.8904923 | 1.9887594 | 2 | 98.1 | 99.6 | 1.5 |
| AKFFEDYGLF | -5.4275632 | 0.97310652 | 8 | -6.9460415 | 2.60974804 | 3 | 97.7 | 99.2 | 1.5 |
| QQIELDLKATQAL | -3.7865865 | 2.19675848 | 2 | -4.1605871 | 0.60760718 | 4 | 93.2 | 94.7 | 1.5 |
| AIIDPGDSDIIRSMPEQTGEK | -2.4856371 | 0.24707594 | 5 | -2.6561764 | 0.26437771 | 4 | 84.9 | 86.3 | 1.5 |
| GKESQAKDVIEEYF | -5.6001776 | 2.71407948 | 3 | -7.4647996 | 3.00547736 | 3 | 98.0 | 99.4 | 1.5 |
| IAEDENGKIVGY | -5.7019226 | 0.51098842 | 4 | -7.8565487 | 1.48867484 | 2 | 98.1 | 99.6 | 1.5 |
| GFPVDLTGLIAEEKGLVVDMDGFEEERKLAQL | -5.4846058 | 0.19383234 | 2 | -7.0834722 | 0.79757238 | 5 | 97.8 | 99.3 | 1.5 |
| AMIGLGCGEGGEIIVTDMDTIEKSNL | -2.7011068 | 0.31605125 | 3 | -2.8915018 | 0.7188814 | 4 | 86.7 | 88.1 | 1.5 |
| GFINRNDTKEDVF | -5.186998 | 0.97714369 | 6 | -6.3390217 | 0.34520289 | 2 | 97.3 | 98.8 | 1.5 |

| | | | | | | | | | |
|---|---|---|---|---|---|---|---|---|---|
| GLDKSEDKVIAVY | -5.5810956 | 1.52700364 | 8 | -7.3807626 | 1.76991215 | 4 | 98.0 | 99.4 | 1.4 |
| NFKTSLW | -4.5222723 | 1.05449876 | 4 | -5.1594132 | 1.83009951 | 3 | 95.8 | 97.3 | 1.4 |
| LLGSTAEEKAIVQQWLEY | -3.4841707 | 0.90143843 | 7 | -3.7867529 | 0.41363095 | 2 | 91.8 | 93.2 | 1.4 |
| LNTVTVSPDGSLCASGGKDGQAMLW | -3.4037955 | 0.32009504 | 6 | -3.6910082 | 0.36750855 | 3 | 91.4 | 92.8 | 1.4 |
| ASGSPDNIKQW | -3.8844017 | 0.57595396 | 2 | -4.2797215 | 0.16606062 | 2 | 93.7 | 95.1 | 1.4 |
| VCQKDSAATTDEERQHLQEVGLF | -3.5369375 | 0.23475984 | 2 | -3.849221 | 0.4271345 | 3 | 92.1 | 93.5 | 1.4 |
| AKYGEPGEVF | -5.8570134 | 1.28419115 | 4 | -8.620531 | 0.4578849 | 2 | 98.3 | 99.7 | 1.4 |
| KVFDFELSSQDMTTL | -4.1010266 | 0.7845791 | 2 | -4.5606327 | 0.32928927 | 3 | 94.5 | 95.9 | 1.4 |
| NVDLIPKFL | 3.11165122 | 0.23546664 | 9 | 2.90108519 | 0.5173282 | 11 | 10.4 | 11.8 | 1.4 |
| VAGDSKNDPPMEAAGF | -4.6034234 | 0.3058604 | 2 | -5.2770736 | 1.11511237 | 4 | 96.0 | 97.5 | 1.4 |
| NIIGGLDEEGKGAVY | -4.7850559 | 1.22363776 | 3 | -5.5653438 | 1.21770156 | 5 | 96.5 | 97.9 | 1.4 |
| LYEPQVNSPKDFTEGTLF | -3.3097623 | 0.650323 | 3 | -3.5767548 | 0.04285082 | 3 | 90.8 | 92.3 | 1.4 |
| IMESGAKGCEVVVSGKL | -5.8607625 | 1.82699616 | 5 | -8.5601253 | 1.12494408 | 7 | 98.3 | 99.7 | 1.4 |
| SDNQLQEGKNVIGL | -4.8385408 | 0.57420921 | 4 | -5.6517148 | 1.12156983 | 4 | 96.6 | 98.0 | 1.4 |
| FVLDEIDAALDNTNIGKVANY | -3.4034891 | 0.45442379 | 4 | -3.6855015 | 0.50490786 | 5 | 91.4 | 92.8 | 1.4 |
| KNVTELNEPLSNEERNLL | -2.8540988 | 0.25991018 | 4 | -3.056521 | 0.12782509 | 3 | 87.9 | 89.3 | 1.4 |
| ELLDYIVNEPPPKLPSGVF | -2.2354152 | 0.32215594 | 4 | -2.3818703 | 0.29616303 | 4 | 82.5 | 83.9 | 1.4 |
| YGQLPKFQDGDLTLY | -5.344503 | 1.09612233 | 8 | -6.6535404 | 1.81896045 | 4 | 97.6 | 99.0 | 1.4 |
| SQVLANGLDNKLREDLERL | -2.932711 | 0.03263541 | 2 | -3.1434246 | 0.24639674 | 4 | 88.4 | 89.8 | 1.4 |
| FVMGVNHEKY | -4.7869979 | 1.34688435 | 10 | -5.5545807 | 1.03007431 | 6 | 96.5 | 97.9 | 1.4 |
| ADLSEAANRNNDALRQAKQESTEY | -4.0020291 | 0.24832391 | 2 | -4.4200997 | 0.23838109 | 2 | 94.1 | 95.5 | 1.4 |
| SDPQGVPYPEAKIGRF | -4.8349119 | 1.62673817 | 3 | -5.6281924 | 1.40140149 | 4 | 96.6 | 98.0 | 1.4 |
| CYVEFDEVDSLKEALTY | -2.7496651 | 0.28906372 | 4 | -2.9373945 | 0.95905268 | 8 | 87.1 | 88.5 | 1.4 |
| TAAEVVVPRDQTPDENDQVIVKIIGHFY | -2.5596517 | 0.26819667 | 4 | -2.7282418 | 0.37417829 | 6 | 85.5 | 86.9 | 1.4 |
| TGLGPEKTSFF | -5.7839364 | 1.32938852 | 5 | -7.9826097 | 0.48284874 | 3 | 98.2 | 99.6 | 1.4 |
| TVEKMSAEINEIIRVL | -5.3363099 | 0.30549231 | 2 | -6.5877634 | 1.84005852 | 7 | 97.6 | 99.0 | 1.4 |
| KAEGSEIRLAKVDATEESDLAQQY | -5.9068163 | 2.97457338 | 4 | -8.6028433 | 1.30923894 | 7 | 98.4 | 99.7 | 1.4 |
| KAENGQFEEFGQTFGENDVIGCF | -3.9916904 | 0.63631056 | 3 | -4.3935052 | 0.67784749 | 2 | 94.1 | 95.5 | 1.4 |
| STDDVQINDISLQDYIAVKEKY | -4.5809147 | 1.59914224 | 11 | -5.1997776 | 0.78641095 | 11 | 96.0 | 97.4 | 1.4 |
| SVETDYTFPLAEKVKAF | -5.3374378 | 1.52088967 | 5 | -6.5506176 | 0.5448666 | 5 | 97.6 | 98.9 | 1.4 |
| LMEEDEDAYKKQFSQY | -4.4883303 | 1.12770853 | 4 | -5.0603003 | 3.20267091 | 4 | 95.7 | 97.1 | 1.4 |
| LSVPCILGQNGISDLVKVTL | -2.6971092 | 0.29660541 | 4 | -2.8731269 | 0.29008934 | 3 | 86.6 | 88.0 | 1.4 |
| LRFDDTNPEKEEAKFF | -5.4360447 | 3.35549865 | 2 | -6.7704504 | 1.42312313 | 2 | 97.7 | 99.1 | 1.4 |
| TFDLTKYPDANPNPNEQ | -5.3312694 | 0.59041674 | 4 | -6.5224049 | 0.15929291 | 3 | 97.6 | 98.9 | 1.3 |
| VIPNTLAVNAAQDSTDLVAKL | -3.4847703 | 0.29490397 | 5 | -3.7643367 | 0.2206157 | 5 | 91.8 | 93.1 | 1.3 |
| AVDVLVSSGEGKAKDAGQRTTIY | -6.0339779 | 1.92297223 | 3 | -9.2877124 | 0.95893848 | 2 | 98.5 | 99.8 | 1.3 |
| GLPVGAVINCADNTGAKNLY | -2.6731543 | 0.24758099 | 8 | -2.845883 | 0.36121532 | 3 | 86.4 | 87.8 | 1.3 |
| LSDLTHQISKDYGVY | -5.0754928 | 1.40028242 | 2 | -6.0002331 | 1.68344866 | 3 | 97.1 | 98.5 | 1.3 |
| ALQEQPVVNAVIDDTTKEVVY | -2.6607724 | 0.55671408 | 4 | -2.8321598 | 0.29376475 | 3 | 86.3 | 87.7 | 1.3 |
| AENSGVKANEVISKLY | -4.4398807 | 1.95398494 | 7 | -4.9826231 | 3.60380328 | 9 | 95.6 | 96.9 | 1.3 |
| KTNEAQAIETARAW | -5.5300887 | 1.80842961 | 7 | -6.9803971 | 2.12952231 | 7 | 97.9 | 99.2 | 1.3 |
| TKVTDETSGCSLTCAQF | -2.8377265 | 0.30480217 | 2 | -3.025208 | 0.10852653 | 3 | 87.7 | 89.1 | 1.3 |
| AKIAEIEAEMARTQ | -4.3852863 | 1.0049778 | 2 | -4.9003459 | 1.64934465 | 2 | 95.4 | 96.8 | 1.3 |
| SPDGSKIAAGSADRFVY | -5.1741202 | 1.74188565 | 5 | -6.1669353 | 1.08434566 | 2 | 97.3 | 98.6 | 1.3 |
| VLVASKELECAEDPGSAGEAARAGHF | -2.9818963 | 0.02756212 | 2 | -3.1835519 | 0.13142749 | 2 | 88.8 | 90.1 | 1.3 |
| EVKATAGDTHLGGEDFDNRLVNHFVEEF | -2.2298415 | 0.13153737 | 6 | -2.3651573 | 0.14255707 | 4 | 82.4 | 83.7 | 1.3 |
| SLGKEDGSGDRGDGPF | -5.2761841 | 0.96829687 | 6 | -6.3604669 | 1.92676533 | 6 | 97.5 | 98.8 | 1.3 |
| TFDQLALDSPKGCGTVL | -5.4880749 | 1.7560823 | 6 | -6.8382746 | 1.26009927 | 5 | 97.8 | 99.1 | 1.3 |
| GLVSLGGKATT | -4.3545257 | 1.32850158 | 3 | -4.851713 | 1.60378325 | 2 | 95.3 | 96.7 | 1.3 |
| LTMFGEKLNGTDPEDVIRNAF | -2.8446835 | 0.47980516 | 4 | -3.0298209 | 0.53804312 | 2 | 87.8 | 89.1 | 1.3 |
| QSITPIEIDMIVGKDREGF | -2.2870136 | 0.32973584 | 3 | -2.4253088 | 0.19978312 | 3 | 83.0 | 84.3 | 1.3 |
| QSAGVAPESFEYIEAHGTGTKVGDPQELNGITRAI | -2.0078674 | 0.54316785 | 5 | -2.1294467 | 0.27740511 | 4 | 80.1 | 81.4 | 1.3 |
| TLPEVAECFDEITYVELQKEEAQKLLEQY | -5.787454 | 2.31334765 | 3 | -7.7242666 | 0.04314129 | 2 | 98.2 | 99.5 | 1.3 |
| TIANQFPLNKLTELL | -5.0559154 | 0.73428639 | 3 | -5.932589 | 0.96548086 | 2 | 97.1 | 98.4 | 1.3 |
| DLSEKLEQSEAQLGRGSF | -4.1120533 | 0.57310367 | 3 | -4.5251144 | 0.43338424 | 2 | 94.5 | 95.8 | 1.3 |
| IVKKSEDEEDAAKDTY | -5.8856994 | 1.49452757 | 3 | -8.1000399 | 3.23156409 | 3 | 98.3 | 99.6 | 1.3 |
| NGDNMLEPSANMPWFKGW | -3.4435715 | 0.2544352 | 8 | -3.7052519 | 0.33501182 | 4 | 91.6 | 92.9 | 1.3 |
| DLAGRDLTDYLMKILTERGY | -3.7913163 | 0.3070764 | 3 | -4.1196746 | 0.13846304 | 5 | 93.3 | 94.6 | 1.3 |
| DIAQVNLKYLL | -5.8344733 | 1.33103198 | 8 | -7.8602594 | 1.85651253 | 3 | 98.3 | 99.6 | 1.3 |
| AGDIANQLATDAVQILGGNGFNTEYPVEKL | -0.6416279 | 0.40930371 | 7 | -0.7204934 | 0.5770126 | 3 | 60.9 | 62.2 | 1.3 |
| SVIAHVDHGKSTL | -6.109845 | 0.97245089 | 3 | -9.5377894 | 0.60527612 | 2 | 98.6 | 99.9 | 1.3 |
| LLSKGLQVGGCEPEPQVC | -4.386094 | 0.66671784 | 7 | -4.8826441 | 1.22214042 | 6 | 95.4 | 96.7 | 1.3 |
| KVDNDENEHQL | -5.2109011 | 2.68544087 | 2 | -6.1955783 | 2.31256573 | 5 | 97.4 | 98.7 | 1.3 |
| KTIISYIDEQFERY | -5.3295566 | 0.56856042 | 2 | -6.4334614 | 0.28736719 | 2 | 97.6 | 98.9 | 1.3 |
| LLTGGQDKLL | -5.7455316 | 0.95926598 | 4 | -7.4962703 | 0.88289004 | 4 | 98.2 | 99.4 | 1.3 |
| TAVHDAILEDLVFPSEIVGKR | -5.6634076 | 3.47101716 | 2 | -7.2417666 | 2.80603951 | 2 | 98.1 | 99.3 | 1.3 |
| QLLQEAGIKTAF | -4.2941664 | 1.9422805 | 4 | -4.7512737 | 1.08032464 | 4 | 95.2 | 96.4 | 1.3 |
| LNEIKDSVVAGF | -5.8832987 | 2.52101361 | 8 | -7.9743103 | 1.82093377 | 9 | 98.3 | 99.6 | 1.3 |
| TIEDGIFEVKSTAGDTHLGGEDFDNRM(15.9949) | -2.509063 | 0.1459706 | 6 | -2.6585503 | 0.05905703 | 2 | 85.1 | 86.3 | 1.3 |
| AEPVPGIKAEPDESNARY | -5.2026329 | 0.60995214 | 4 | -6.1577246 | 3.38169574 | 2 | 97.4 | 98.6 | 1.3 |
| LTQTVCLDDTTVKF | -5.5727171 | 1.1776998 | 3 | -6.9603842 | 0.38230276 | 4 | 97.9 | 99.2 | 1.3 |
| VLPEKLGVDLDAQTW | -4.306477 | 0.95242661 | 4 | -4.7637217 | 0.8691493 | 3 | 95.2 | 96.4 | 1.3 |

| | | | | | | | | | |
|---|---|---|---|---|---|---|---|---|---|
| SAGLQASDLPASASLPAKADLL | -4.1675536 | 1.83263795 | 3 | -4.5798089 | 0.54494856 | 3 | 94.7 | 96.0 | 1.3 |
| LSLLSDAEKLEQVLQW | -4.8335059 | 0.02638034 | 3 | -5.5207768 | 1.62949324 | 2 | 96.6 | 97.9 | 1.3 |
| SIFTEKDEILSDVASRLW | -3.3020377 | 0.54633856 | 2 | -3.5336104 | 0.43912368 | 5 | 90.8 | 92.1 | 1.3 |
| VDDSIVRGNTISPIIKLL | -5.9613392 | 0.56052462 | 3 | -8.2669698 | 1.96772981 | 4 | 98.4 | 99.7 | 1.3 |
| IFLVDASKAMF | -4.0403747 | 0.13334953 | 2 | -4.4144199 | 2.21189152 | 2 | 94.3 | 95.5 | 1.2 |
| ILTSSDDMLIKLW | -4.8040972 | 1.67637776 | 4 | -5.4692762 | 1.71263385 | 6 | 96.5 | 97.8 | 1.2 |
| GNVGTVEWADPIEDPDPEVMAKVKVLF | -4.6352366 | 0.72610835 | 2 | -5.2155035 | 1.52950877 | 3 | 96.1 | 97.4 | 1.2 |
| KSFIKDYPVVSIEDPFDQDDWGAW | -5.8785144 | 1.88562027 | 4 | -7.8554528 | 1.22146032 | 6 | 98.3 | 99.6 | 1.2 |
| ELLEKEILFY | -5.8380084 | 1.16785668 | 4 | -7.70314 | 1.11190386 | 6 | 98.3 | 99.5 | 1.2 |
| NFVKGVVDSDDLPLNVSRETL | -4.4145141 | 0.42549565 | 4 | -4.8998908 | 0.5225218 | 4 | 95.5 | 96.8 | 1.2 |
| AENSGVKANEVISKLY | -4.9797991 | 1.66882618 | 4 | -5.7421107 | 1.27964887 | 6 | 96.9 | 98.2 | 1.2 |
| LGSLVDEFKEL | -5.8289006 | 2.10778548 | 3 | -7.6613235 | 1.48131824 | 2 | 98.3 | 99.5 | 1.2 |
| GTEKDKVNCSF | -5.9449558 | 2.27603847 | 2 | -8.0937899 | 0.29991206 | 2 | 98.4 | 99.6 | 1.2 |
| CIQALPEFDGKRF | -5.6344315 | 0.53595532 | 4 | -7.0603409 | 1.00239201 | 4 | 98.0 | 99.3 | 1.2 |
| QAEWDDYVPKLY | -5.7836404 | 1.10867139 | 5 | -7.486226 | 1.06749269 | 8 | 98.2 | 99.4 | 1.2 |
| STASDNQPTVTIKVY | -5.4819502 | 1.52300671 | 7 | -6.6864387 | 0.7385719 | 7 | 97.8 | 99.0 | 1.2 |
| KVFPGSTTEDY | -4.759583 | 0.98411202 | 2 | -5.3833924 | 1.49401613 | 5 | 96.4 | 97.7 | 1.2 |
| VSDENDAKILNDVQDRFEVNISELPDEIDISSY | -2.5947663 | 0.12427479 | 10 | -2.7442308 | 0.23505897 | 8 | 85.8 | 87.0 | 1.2 |
| SGAQIKIANPVEGSSGRQVT | -4.8494083 | 0.32709305 | 2 | -5.5166411 | 2.78441405 | 2 | 96.6 | 97.9 | 1.2 |
| TWFTDHSDAGADELGEVIKDDIWPNPLQY | -3.8169254 | 0.25197829 | 12 | -4.1276831 | 0.25782205 | 3 | 93.4 | 94.6 | 1.2 |
| QMKNPNGPYPY | -5.9596491 | 0.80586804 | 5 | -8.0808984 | 2.66563116 | 2 | 98.4 | 99.6 | 1.2 |
| AKVDATAETDLAKRF | -6.0451432 | 2.64718379 | 3 | -8.4840621 | 1.55599621 | 6 | 98.5 | 99.7 | 1.2 |
| KELAPYDENWFY | -5.4192819 | 1.22696304 | 4 | -6.5296373 | 1.66000389 | 5 | 97.7 | 98.9 | 1.2 |
| VFTKEDLTEIRDMLL | -4.9666506 | 0.2066677 | 2 | -5.7003694 | 0.87694439 | 4 | 96.9 | 98.1 | 1.2 |
| SATM(15.9949)PSDVLEVTKKF | -4.3894797 | 3.27784386 | 2 | -4.854086 | 1.32470573 | 4 | 95.4 | 96.7 | 1.2 |
| QVKSGTIFDNF | -5.591827 | 1.85536456 | 9 | -6.9155542 | 1.77149151 | 8 | 98.0 | 99.2 | 1.2 |
| ESLTDPSKL | -6.0611089 | 0.90954377 | 2 | -8.5499267 | 2.2631407 | 2 | 98.5 | 99.7 | 1.2 |
| AIAAKEKDIQEESTF | -5.8998901 | 0.81311357 | 6 | -7.8277742 | 1.20367808 | 2 | 98.4 | 99.6 | 1.2 |
| TLNVLEDLGDGQKANDDIIVNW | -2.7169049 | 0.22737772 | 4 | -2.8753941 | 0.22126119 | 3 | 86.8 | 88.0 | 1.2 |
| SLGVNDILEIKFF | -5.327556 | 1.61352983 | 6 | -6.330561 | 0.96044328 | 7 | 97.6 | 98.8 | 1.2 |
| AIRNDEELNKLL | -5.7958934 | 1.53527873 | 6 | -7.45648 | 1.33953538 | 5 | 98.2 | 99.4 | 1.2 |
| EISDIVIKDSNTDVGAKNY | -5.4261932 | 1.42803056 | 2 | -6.5288822 | 1.15398514 | 3 | 97.7 | 98.9 | 1.2 |
| CVQQLKEF | -5.7275482 | 1.36133043 | 12 | -7.2470023 | 0.07588921 | 4 | 98.1 | 99.3 | 1.2 |
| SGIESSSPEVKGYW | -4.7553141 | 1.56740416 | 4 | -5.3616854 | 1.09406997 | 5 | 96.4 | 97.6 | 1.2 |
| FLINGPEIMSKLAGESESNL | -3.0370965 | 0.20292947 | 3 | -3.2246338 | 0.32415459 | 4 | 89.1 | 90.3 | 1.2 |
| QQLKDETGELSSADEKRY | -5.6539774 | 2.10443113 | 2 | -7.045021 | 1.06965828 | 4 | 98.1 | 99.2 | 1.2 |
| SSSQDPPEKSIPICTL | -5.3761111 | 1.6131227 | 6 | -6.4130272 | 1.14769767 | 4 | 97.6 | 98.8 | 1.2 |
| IVFGEAKIEDLSQQAQLAAAEKF | -5.5236231 | 1.21101807 | 6 | -6.7205374 | 1.67893662 | 3 | 97.9 | 99.1 | 1.2 |
| YGTEKDKNSVNF | -6.0970919 | 1.03867663 | 3 | -8.6293175 | 1.99866602 | 4 | 98.6 | 99.7 | 1.2 |
| LAPKDKPSGDTAAVFEEGGDVDDLLDM(15.994[illegible] | -5.1433233 | 0.4575959 | 3 | -5.9757349 | 1.2149703 | 4 | 97.2 | 98.4 | 1.2 |
| KLVVGGLLSPEEDQSLLL | -5.3127708 | 0.79023065 | 2 | -6.2810641 | 2.70217155 | 2 | 97.5 | 98.7 | 1.2 |
| KIGDQEFDHLPALLEFY | -2.2322334 | 0.32725535 | 3 | -2.3535037 | 0.04363421 | 2 | 82.5 | 83.6 | 1.2 |
| AAMEELVDEGLVKAIGISNF | -2.4528383 | 0.21191422 | 5 | -2.5875346 | 1.08266544 | 6 | 84.6 | 85.7 | 1.2 |
| MDRGEGGTTNPHIFPEGSEPKVY | -3.2920464 | 0.31854695 | 11 | -3.5071189 | 0.11580834 | 2 | 90.7 | 91.9 | 1.2 |
| KGFYEGIMLVDGF | -4.2619391 | 0.32336933 | 3 | -4.6715843 | 0.67181195 | 2 | 95.0 | 96.2 | 1.2 |
| VSDENDAKILNDVQDRFEVNVAELPEEIDISTY | -4.2990379 | 1.58862521 | 3 | -4.7195602 | 2.75842499 | 2 | 95.2 | 96.3 | 1.2 |
| EGERPLTKDNHLLGTF | -5.4623656 | 1.08516106 | 5 | -6.5711593 | 0.65850758 | 5 | 97.8 | 99.0 | 1.2 |
| TDTERLIGDAAKNQVAMNPTNTVF | -5.8615591 | 0.4428064 | 5 | -7.5842269 | 2.12527122 | 3 | 98.3 | 99.5 | 1.2 |
| EGSPIKVTLATL | -5.0297715 | 1.18828634 | 13 | -5.7697496 | 2.80430035 | 7 | 97.0 | 98.2 | 1.2 |
| TDTGKASGNLETKY | -5.1908791 | 2.03243522 | 4 | -6.0393944 | 2.81761574 | 3 | 97.3 | 98.5 | 1.2 |
| LCGKPDVPIYDGSWVEW | -2.4626132 | 0.10971109 | 2 | -2.5962723 | 0.45007152 | 2 | 84.6 | 85.8 | 1.2 |
| LINVKQENTQTEQKL | -5.9382144 | 2.53404844 | 2 | -7.8249352 | 3.02761787 | 2 | 98.4 | 99.6 | 1.2 |
| FGAFGEIENIELPMDTKTNERRGF | -1.8847647 | 0.4460074 | 4 | -1.9870269 | 0.29623358 | 3 | 78.7 | 79.9 | 1.2 |
| SMKEVDEQMLNVQNKNSSYF | -6.1550787 | 1.1788836 | 5 | -8.8254214 | 1.84631326 | 3 | 98.6 | 99.8 | 1.2 |
| M(15.9949)VVNDAGRPKVQVEY | -5.8666439 | 0.85531228 | 3 | -7.5727856 | 1.06984822 | 5 | 98.3 | 99.5 | 1.2 |
| QDLKDHMREAGDVCY | -4.2335979 | 0.33264114 | 2 | -4.6286538 | 0.44268641 | 4 | 95.0 | 96.1 | 1.2 |
| NLVKDSATGL | -6.1582935 | 0.63992589 | 6 | -8.8296663 | 0.68661142 | 5 | 98.6 | 99.8 | 1.2 |
| HLSNIPPSVSEEDLKVLF | -5.420177 | 1.9942165 | 10 | -6.4615527 | 0.52029543 | 7 | 97.7 | 98.9 | 1.2 |
| LSVPCILGQNGISDLVKVTL | -2.7325951 | 0.32876264 | 4 | -2.8854075 | 0.29796003 | 3 | 86.9 | 88.1 | 1.2 |
| SFCPTVNLDKLW | -5.6264241 | 1.4453125 | 8 | -6.8979785 | 1.70134525 | 4 | 98.0 | 99.2 | 1.2 |
| DILGVPPGASENELKKAY | -5.4522075 | 2.37070394 | 6 | -6.5136715 | 0.24163653 | 3 | 97.8 | 98.9 | 1.2 |
| LSDSGSTGEHTKSLVTQY | -6.2418263 | 0.81486587 | 3 | -9.3409025 | 0.92748307 | 4 | 98.7 | 99.8 | 1.2 |
| FSDCKIQNGAQGIRF | -5.0143127 | 0.4434566 | 5 | -5.7239014 | 1.35895541 | 2 | 97.0 | 98.1 | 1.1 |
| DISILEIQKGVF | -5.2512974 | 0.74759887 | 6 | -6.1204291 | 1.40682328 | 7 | 97.4 | 98.6 | 1.1 |
| AGKILNDDTALKEY | -5.4413687 | 2.37644902 | 5 | -6.4780226 | 2.2349006 | 6 | 97.8 | 98.9 | 1.1 |
| VGVILDEAKGKNDGTVQ | -3.6384595 | 0.98260951 | 3 | -3.8960071 | 0.37775654 | 4 | 92.6 | 93.7 | 1.1 |
| SGAPAAESKEIVRGY | -4.7370854 | 0.97607105 | 5 | -5.2990709 | 1.63269053 | 2 | 96.4 | 97.5 | 1.1 |
| NAAKVPADTEVVCAPPTAY | -2.0788884 | 0.58044013 | 2 | -2.1873209 | 0.18942987 | 3 | 80.9 | 82.0 | 1.1 |
| VAPGNAGTACSEKISNTAISISDHTAL | -1.6781348 | 0.83004768 | 2 | -1.7698606 | 0.6742165 | 4 | 76.2 | 77.3 | 1.1 |
| KEVDEQMLNVQNKNSSYF | -4.6466203 | 0.30283494 | 4 | -5.1669284 | 2.14244039 | 2 | 96.2 | 97.3 | 1.1 |
| KLLDEVFFSEKIY | -5.5044698 | 1.49445884 | 3 | -6.5940935 | 0.17717656 | 2 | 97.8 | 99.0 | 1.1 |
| HELTQTDKALF | -4.9013612 | 1.75885968 | 4 | -5.5366042 | 0.96485041 | 6 | 96.8 | 97.9 | 1.1 |

| | | | | | | | | | |
|---|---|---|---|---|---|---|---|---|---|
| SDVQLIKTGDKVGASEATLL | -5.9703387 | 2.4210365 | 7 | -7.8124925 | 2.04987576 | 5 | 98.4 | 99.6 | 1.1 |
| ELSDNRVSGGLEVLAEKCPNLTHL | -2.3483679 | 0.30212895 | 2 | -2.4700117 | 0.2865475 | 6 | 83.6 | 84.7 | 1.1 |
| LSNLSSTSHVPEVDPGSAELQKVLQGDLVMNVY | -2.7260345 | 0.14257038 | 2 | -2.8736441 | 0.13030585 | 2 | 86.9 | 88.0 | 1.1 |
| GIGQDIQPKRDL | -5.4826706 | 1.45766368 | 6 | -6.5373845 | 1.1883346 | 7 | 97.8 | 98.9 | 1.1 |
| STQVDTVATKVKDALEFWLQAGVDGF | -5.6970175 | 0.5047917 | 4 | -7.0109121 | 0.99547989 | 2 | 98.1 | 99.2 | 1.1 |
| FLGASLKDEVL | -5.4243834 | 0.5783626 | 5 | -6.4151291 | 1.86143646 | 4 | 97.7 | 98.8 | 1.1 |
| NNDKSRDYTRPDLPSGDSQPSLDQTMAAAF | -2.1148194 | 0.17259454 | 8 | -2.2230479 | 0.07152282 | 2 | 81.2 | 82.4 | 1.1 |
| CPTVNLDKLW | -5.1540583 | 1.43051792 | 10 | -5.9281964 | 1.84026541 | 7 | 97.3 | 98.4 | 1.1 |
| VRNLGDVGEKETETEVL | -5.4645635 | 1.48671783 | 5 | -6.4906608 | 2.05776996 | 6 | 97.8 | 98.9 | 1.1 |
| TLEGIKQFF | -6.1780911 | 1.90834032 | 2 | -8.6392921 | 1.43038119 | 3 | 98.6 | 99.7 | 1.1 |
| IQEAKEIDVDAVASDGVVAAIAISEHVENAGVH | -3.460788 | 0.85742675 | 3 | -3.6849598 | 0.18229651 | 6 | 91.7 | 92.8 | 1.1 |
| NVTVTKTDKTLVL | -4.3680007 | 2.15800228 | 4 | -4.7815274 | 0.58329911 | 3 | 95.4 | 96.5 | 1.1 |
| SAEKVEIATL | -5.8297472 | 2.08377613 | 6 | -7.3268691 | 1.87512564 | 4 | 98.3 | 99.4 | 1.1 |
| FADMVSKSANY | -5.6471774 | 3.78709877 | 3 | -6.866548 | 1.03857064 | 4 | 98.0 | 99.2 | 1.1 |
| IITEETAKIRPF | -4.7337703 | 0.26996158 | 3 | -5.2752301 | 3.39496598 | 4 | 96.4 | 97.5 | 1.1 |
| ITTKNSNILEDLETL | -5.4270945 | 1.32100706 | 4 | -6.402739 | 1.34533416 | 5 | 97.7 | 98.8 | 1.1 |
| VLSLEIGKTLMEDVENSFF | -3.7353901 | 0.273232 | 3 | -4.0000159 | 0.837139 | 5 | 93.0 | 94.1 | 1.1 |
| VKPAPDETSFSEALL | -5.4128472 | 0.93775255 | 6 | -6.370056 | 0.89133544 | 4 | 97.7 | 98.8 | 1.1 |
| VCKSESVPPVTDWAWY | -4.9562316 | 1.29193983 | 9 | -5.5975982 | 0.43143765 | 4 | 96.9 | 98.0 | 1.1 |
| LDLILNDFVRQKF | -5.156781 | 0.34767758 | 2 | -5.9133486 | 0.15478856 | 2 | 97.3 | 98.4 | 1.1 |
| KPVFVTEITDDLHFY | -5.6299745 | 0.15668035 | 3 | -6.8060125 | 1.61198686 | 4 | 98.0 | 99.1 | 1.1 |
| GGAGVGKTVL | -5.8397653 | 1.42295737 | 8 | -7.3145269 | 1.72942612 | 7 | 98.3 | 99.4 | 1.1 |
| VGLTDFKPGYW | -5.4232178 | 1.26038502 | 10 | -6.3780251 | 0.99091919 | 5 | 97.7 | 98.8 | 1.1 |
| TSSGSANTETTKVTGSLETKY | -6.0212198 | 2.37871545 | 5 | -7.8634107 | 0.94654876 | 3 | 98.5 | 99.6 | 1.1 |
| SNQTVDIPENVDITLKGR | -5.4807254 | 2.98451239 | 6 | -6.4853615 | 1.89079039 | 3 | 97.8 | 98.9 | 1.1 |
| TGVLKPGMVVTF | -4.340529 | 1.49233311 | 4 | -4.7353939 | 0.37533216 | 5 | 95.3 | 96.4 | 1.1 |
| LAKLVINTEEVF | -6.2205097 | 1.04209976 | 3 | -8.6809036 | 0.3259153 | 3 | 98.7 | 99.8 | 1.1 |
| LLTTTPRPVIVEPLEQLDDEDGLPEKL | -1.7622166 | 0.25564249 | 5 | -1.8523129 | 0.30050065 | 3 | 77.2 | 78.3 | 1.1 |
| SDPQGVPYPEAKIGRF | -5.1519432 | 1.4825038 | 6 | -5.8905617 | 1.24872235 | 6 | 97.3 | 98.3 | 1.1 |
| FHSLSEKY | -6.1040815 | 1.50685627 | 7 | -8.1219607 | 1.36143137 | 5 | 98.6 | 99.6 | 1.1 |
| VGGLKGDVAEGDLIEHF | -5.5925358 | 1.73441153 | 8 | -6.6900212 | 1.69820096 | 6 | 98.0 | 99.0 | 1.1 |
| DVDFPKEQLTEEAREGIKQLL | -3.933016 | 1.26996695 | 3 | -4.2256845 | 0.6118452 | 3 | 93.9 | 94.9 | 1.1 |
| GMGNPLLDISAVVDKDFLDKY | -5.9168159 | 0.41740506 | 2 | -7.4743949 | 1.7852143 | 4 | 98.4 | 99.4 | 1.1 |
| ALSVETDYTFPLAEKVKAF | -4.223559 | 0.70941775 | 3 | -4.5802058 | 0.35257066 | 2 | 94.9 | 96.0 | 1.1 |
| SIAGSADSKPIDVSRL | -5.5344945 | 0.52800561 | 10 | -6.5651742 | 1.59268454 | 5 | 97.9 | 99.0 | 1.1 |
| KSDVLQPGAEVTTDDRAY | -5.7879116 | 1.83865922 | 2 | -7.1242011 | 2.18534301 | 4 | 98.2 | 99.3 | 1.1 |
| LAEKYEWDVAEARKIW | -6.0095626 | 0.1431589 | 2 | -7.74794 | 0.41810722 | 2 | 98.5 | 99.5 | 1.1 |
| AYVQFEDVRDAEDALHNLDRKW | -3.0171188 | 0.47549916 | 4 | -3.1812667 | 0.04906023 | 3 | 89.0 | 90.1 | 1.1 |
| ASPAHAVDAVKNADGY | -5.23136 | 0.8803129 | 13 | -5.9996852 | 1.08154251 | 8 | 97.4 | 98.5 | 1.1 |
| DLIANIVHDGKPSEGSY | -5.7494314 | 0.09738043 | 3 | -7.0078939 | 0.70609305 | 4 | 98.2 | 99.2 | 1.1 |
| ALSVETDYTFPLAEKVKAF | -4.2166876 | 0.43506494 | 4 | -4.5660396 | 0.25632649 | 3 | 94.9 | 95.9 | 1.1 |
| AEIAKVELDNMPL | -5.9087536 | 0.94619736 | 5 | -7.409132 | 0.19390764 | 2 | 98.4 | 99.4 | 1.1 |
| MRDSGSKAATDAQDANQCCTSCEDNAPATSY | -2.2677027 | 0.25953269 | 3 | -2.3769742 | 0.27346066 | 3 | 82.8 | 83.9 | 1.1 |
| VISIKANCIDSTASAEAVF | -3.2555685 | 0.32581234 | 7 | -3.4418774 | 0.43301459 | 8 | 90.5 | 91.6 | 1.1 |
| SAAIESCNKALELDSNNEKGLF | -5.8034984 | 0.89821281 | 6 | -7.1315097 | 1.4761058 | 3 | 98.2 | 99.3 | 1.1 |
| FLKDISTTL | -5.2347764 | 1.14553784 | 6 | -6.0005649 | 0.83956272 | 2 | 97.4 | 98.5 | 1.0 |
| YNEATGGKYVPR | -5.5522922 | 2.12782423 | 12 | -6.5711949 | 1.55564468 | 6 | 97.9 | 99.0 | 1.0 |
| HLNESGDPSSKSTEIKW | -6.1817595 | 2.52224782 | 13 | -8.2934744 | 2.82865474 | 8 | 98.6 | 99.7 | 1.0 |
| QSAGVAPESFEYIEAHGTGTKVGDPQEL | -3.1477188 | 0.46791886 | 2 | -3.3197752 | 0.1667521 | 3 | 89.9 | 90.9 | 1.0 |
| VGNLGNNGNKTELERAF | -6.0224635 | 1.68069848 | 2 | -7.6986494 | 0.61152613 | 3 | 98.5 | 99.5 | 1.0 |
| KSLGLSDEEIVKF | -6.0924966 | 2.33107227 | 4 | -7.9261192 | 1.32191182 | 3 | 98.6 | 99.6 | 1.0 |
| STVFKDDDDVVIGKVF | -5.2068592 | 1.79007275 | 6 | -5.9385848 | 1.95661867 | 4 | 97.4 | 98.4 | 1.0 |
| SDLKLDYLDLY | -5.5331218 | 0.51233163 | 2 | -6.5110688 | 0.82870832 | 4 | 97.9 | 98.9 | 1.0 |
| EALSVKEETKEDAEEKQ | -5.8477544 | 2.33730454 | 3 | -7.1956905 | 2.88517599 | 3 | 98.3 | 99.3 | 1.0 |
| IFAENDVVIPLKDVSLL | -5.7273197 | 1.37985833 | 4 | -6.9111361 | 2.32764739 | 4 | 98.1 | 99.2 | 1.0 |
| VVQEPGDYEVSVKF | -3.361348 | 0.44900627 | 3 | -3.5546244 | 0.56085082 | 2 | 91.1 | 92.2 | 1.0 |
| QASDLPASASLPAKADLL | -5.8308413 | 1.06530401 | 8 | -7.1402336 | 2.32366505 | 4 | 98.3 | 99.3 | 1.0 |
| SVLENGVDIVVGTPGRLDDLVSTGKL | -3.3247184 | 0.38512329 | 4 | -3.5128913 | 0.35145677 | 2 | 90.9 | 91.9 | 1.0 |
| EAGISKNGQTREHALL | -5.1574272 | 1.07567628 | 5 | -5.846624 | 2.39981764 | 2 | 97.3 | 98.3 | 1.0 |
| ALQLAEKLGSLVENNERVF | -3.964916 | 0.09438683 | 3 | -4.2462513 | 0.47297971 | 2 | 94.0 | 95.0 | 1.0 |
| GDPVVQSDM(15.9949)KHWPF | -4.2544404 | 0.92951319 | 6 | -4.59759 | 1.44797562 | 5 | 95.0 | 96.0 | 1.0 |
| IVVPDVSKL | -6.3611547 | 3.52776914 | 3 | -9.0349591 | 0.37567748 | 3 | 98.8 | 99.8 | 1.0 |
| FSEKFPTLW | -4.1427841 | 0.25769025 | 6 | -4.4593301 | 1.67060355 | 3 | 94.6 | 95.7 | 1.0 |
| KAVTEQGHELSNEERNLL | -4.4974507 | 1.10957325 | 6 | -4.9045708 | 1.09609769 | 4 | 95.8 | 96.8 | 1.0 |
| NMDVPNIKRNDF | -5.4902384 | 1.98982838 | 3 | -6.3998289 | 0.92268211 | 5 | 97.8 | 98.8 | 1.0 |
| FQFQEEGKEGENRAVIHY | -2.9281303 | 0.14842947 | 4 | -3.0751105 | 0.26428045 | 3 | 88.4 | 89.4 | 1.0 |
| IGNLDPEIDEKLLYDTF | -4.5336389 | 0.42902683 | 5 | -4.9495254 | 1.17908679 | 5 | 95.9 | 96.9 | 1.0 |
| IFLDDPQAVSDILEKLVKEDNLL | -5.0666245 | 1.40464114 | 2 | -5.6948657 | 1.56251 | 4 | 97.1 | 98.1 | 1.0 |
| FVVKAYLPVNESFGF | -5.1331064 | 0.46575818 | 3 | -5.7966802 | 1.89402128 | 3 | 97.2 | 98.2 | 1.0 |
| VISDRKELEEDFIKSEL | -4.9393866 | 1.09298115 | 4 | -5.5050756 | 4.97746935 | 2 | 96.8 | 97.8 | 1.0 |
| ELESKLNEAKEEF | -5.7574606 | 0.75505773 | 5 | -6.9253423 | 1.31473881 | 4 | 98.2 | 99.2 | 1.0 |
| KLVVGGLLSPEEDQSLL | -4.3280677 | 1.01742256 | 9 | -4.6842147 | 1.07538137 | 4 | 95.3 | 96.3 | 1.0 |

| | | | | | | | | | |
|---|---|---|---|---|---|---|---|---|---|
| TGKTITDVINIGIGGSDLGPL | -5.1791187 | 0.77847501 | 6 | -5.8642405 | 0.55430837 | 5 | 97.3 | 98.3 | 1.0 |
| TDDDKTDHLSWEW | -6.0625323 | 1.00721775 | 3 | -7.7058746 | 0.95818676 | 3 | 98.5 | 99.5 | 1.0 |
| VGGLSPDTSEEQIKEY | -5.2248885 | 1.15239527 | 5 | -5.9357197 | 2.55861708 | 7 | 97.4 | 98.4 | 1.0 |
| QEALDAAGDKLVVVDF | -5.4000735 | 0.69172245 | 6 | -6.2279367 | 1.26257967 | 8 | 97.7 | 98.7 | 1.0 |
| VVDVDKNIDINDVTPNCRVAL | -2.6714895 | 0.48401828 | 4 | -2.797644 | 0.12100777 | 2 | 86.4 | 87.4 | 1.0 |
| SVNDPPDVLDRQKCL | -5.0933148 | 0.55881628 | 3 | -5.7256289 | 1.31733806 | 4 | 97.2 | 98.1 | 1.0 |
| ICERIFYPEIEEVQALDDTERGSGGFGSTGKN | -2.3461091 | 0.0669378 | 5 | -2.4526156 | 0.27575963 | 3 | 83.6 | 84.6 | 1.0 |
| RGDLGIEIPAEKVF | -4.8744129 | 1.31437516 | 4 | -5.4031491 | 1.59571452 | 2 | 96.7 | 97.7 | 1.0 |
| SEGLWEIENNPTVKASGY | -5.4932807 | 0.96843247 | 3 | -6.3810943 | 1.51571362 | 5 | 97.8 | 98.8 | 1.0 |
| EVNFQNGIECGGAYVKLL | -3.199934 | 0.87139064 | 5 | -3.3679422 | 0.02801426 | 2 | 90.2 | 91.2 | 1.0 |
| AGKQLEDGRTLSDY | -5.5287953 | 1.00944349 | 5 | -6.4366636 | 1.15384755 | 5 | 97.9 | 98.9 | 1.0 |
| LLGFIPAKADSVVVL | -4.8549973 | 1.2827054 | 3 | -5.3673333 | 1.11295779 | 2 | 96.7 | 97.6 | 1.0 |
| LAEPVPGIKAEPDESNARY | -6.296314 | 1.7513199 | 10 | -8.4512135 | 1.37245291 | 3 | 98.7 | 99.7 | 1.0 |
| SSIKSQTIY | 5.68053914 | 1.0269181 | 3 | 5.07645615 | 0.46812434 | 8 | 1.9 | 2.9 | 1.0 |
| LGEFLHPCEDDIVCKCTTDENKVPY | -6.0290862 | 2.03965904 | 3 | -7.5095839 | 1.12610408 | 2 | 98.5 | 99.5 | 1.0 |
| ELLFKEGVM | -5.7419055 | 2.00890383 | 6 | -6.8262924 | 0.82027151 | 5 | 98.2 | 99.1 | 1.0 |
| VHGGVDASGKPQEAVY | -4.9384727 | 1.1356972 | 7 | -5.4754932 | 0.80810966 | 6 | 96.8 | 97.8 | 1.0 |
| STQVDTVATKVKDALEF | -6.3626397 | 1.54872305 | 8 | -8.6804314 | 1.28246722 | 2 | 98.8 | 99.8 | 1.0 |
| QQIDSVANADIINAAKKF | -5.8478805 | 1.48432786 | 8 | -7.0381426 | 1.30563955 | 3 | 98.3 | 99.2 | 1.0 |
| KTVFDEAIRAVL | -5.3006953 | 0.86305123 | 3 | -6.0140092 | 0.52841643 | 3 | 97.5 | 98.5 | 1.0 |
| VSDLLPPTDKELRETIALL | -2.6798156 | 0.1494281 | 2 | -2.8006723 | 0.40483847 | 2 | 86.5 | 87.4 | 0.9 |
| RTVSLGAGAKDEL | -6.3926817 | 3.08021337 | 2 | -8.7675675 | 0.71025051 | 2 | 98.8 | 99.8 | 0.9 |
| SHLVPVYDVEPLEKITDAYLDQYLW | -4.3893108 | 0.35437843 | 2 | -4.7393003 | 0.33549524 | 4 | 95.4 | 96.4 | 0.9 |
| SCVVKMPSGEF | -5.6721617 | 2.13594423 | 3 | -6.6612418 | 1.54995277 | 6 | 98.1 | 99.0 | 0.9 |
| KAADPPAENSSAPEAEQGGAE | -6.0656773 | 0.72981506 | 2 | -7.5581309 | 1.75976004 | 5 | 98.5 | 99.5 | 0.9 |
| LIAGIQHSCQDIGAKSL | -5.9831979 | 0.79450483 | 3 | -7.3366052 | 2.19749038 | 6 | 98.4 | 99.4 | 0.9 |
| LTYNDFINKELILF | -5.4575659 | 0.73712654 | 10 | -6.2598831 | 1.74423709 | 4 | 97.8 | 98.7 | 0.9 |
| NALGGWGELQNSVKTF | -5.0013302 | 0.90830757 | 8 | -5.5491507 | 1.11944319 | 6 | 97.0 | 97.9 | 0.9 |
| IVGADNVGSKQM | -5.8505482 | 0.81263926 | 6 | -7.0077823 | 1.2289169 | 7 | 98.3 | 99.2 | 0.9 |
| LAPISSSKLY | -5.3070683 | 0.83287673 | 3 | -6.006347 | 1.49124799 | 2 | 97.5 | 98.5 | 0.9 |
| ENERGELANEVKVLL | -5.4740633 | 0.9403365 | 2 | -6.2820489 | 1.9089148 | 3 | 97.8 | 98.7 | 0.9 |
| SDYNIQKESTL | -5.6069263 | 1.32240886 | 5 | -6.5180685 | 1.0679277 | 5 | 98.0 | 98.9 | 0.9 |
| VGGLKGDVAEGDLIEHF | -4.6139739 | 1.08951841 | 7 | -5.0187264 | 0.89289969 | 5 | 96.1 | 97.0 | 0.9 |
| GLSARDLDELSRYPEDKITPENLPQILL | -2.3018348 | 0.13887576 | 2 | -2.3997916 | 0.38807091 | 2 | 83.1 | 84.1 | 0.9 |
| VLAPEGSVPNKF | -4.9790381 | 0.84251794 | 9 | -5.5119789 | 1.98334023 | 2 | 96.9 | 97.9 | 0.9 |
| VSSKADLPEGVAVSGPSPAEF | -4.9838105 | 0.45232406 | 4 | -5.5162021 | 1.81881277 | 3 | 96.9 | 97.9 | 0.9 |
| IGGLSWDTSKKDLTEY | -5.2645344 | 1.93178369 | 7 | -5.9327875 | 3.02590531 | 5 | 97.5 | 98.4 | 0.9 |
| LLGLADNEAAIVQAESEETKERLF | -4.5345075 | 0.28400581 | 3 | -4.9133855 | 1.14829199 | 5 | 95.9 | 96.8 | 0.9 |
| LGGVDKHTQFW | -4.2721027 | 1.4287823 | 7 | -4.5860073 | 0.83297881 | 2 | 95.1 | 96.0 | 0.9 |
| GTLTDCVVVVNPQTKR | -5.5150741 | 1.06306797 | 3 | -6.3402102 | 1.21328108 | 2 | 97.9 | 98.8 | 0.9 |
| KVTQLTGFSDPVYAEAY | -5.7669589 | 1.62520655 | 2 | -6.8073856 | 1.82504578 | 2 | 98.2 | 99.1 | 0.9 |
| TDEEPVKKLLESRY | -5.684567 | 1.10723626 | 7 | -6.6439024 | 0.28876386 | 2 | 98.1 | 99.0 | 0.9 |
| GVTIDFDTVNKTPH | -4.7075662 | 1.31170162 | 7 | -5.1336887 | 1.32756953 | 2 | 96.3 | 97.2 | 0.9 |
| IVFDGDVDPEWVENLNSVLDDNKLL | -3.6544535 | 0.35909303 | 4 | -3.8605584 | 0.23220622 | 4 | 92.6 | 93.6 | 0.9 |
| IVFGEAKIEDLSQQAQL | -5.6566135 | 1.05832892 | 8 | -6.589458 | 1.58539512 | 10 | 98.1 | 99.0 | 0.9 |
| VLEGKELEFY | -6.2170445 | 1.4070979 | 3 | -7.9051386 | 1.75667978 | 7 | 98.7 | 99.6 | 0.9 |
| NAINKCPL | -5.495843 | 0.98539402 | 9 | -6.293653 | 1.24773406 | 7 | 97.8 | 98.7 | 0.9 |
| KDFSELEPDKF | -5.8474226 | 0.69008278 | 4 | -6.9557188 | 1.64220471 | 5 | 98.3 | 99.2 | 0.9 |
| IKTVETRDGQVINETSQHHDDLE | -2.737258 | 0.06808023 | 3 | -2.8562121 | 0.01691592 | 2 | 87.0 | 87.9 | 0.9 |
| QVINDGDKPKVQ | -5.5850936 | 6.57539648 | 2 | -6.4438602 | 3.32805577 | 5 | 98.0 | 98.9 | 0.9 |
| IADVAPSAIRENDIKSY | -4.7639438 | 1.52565962 | 6 | -5.2018267 | 1.86396079 | 7 | 96.5 | 97.4 | 0.9 |
| ELKADVVPKTAENF | -5.6003988 | 1.24166651 | 9 | -6.465888 | 1.5390139 | 6 | 98.0 | 98.9 | 0.9 |
| GTPAEEDGGGKIDLIEAGAF | -0.0013724 | 0.17521096 | 4 | -0.0532443 | 0.0942901 | 5 | 50.0 | 50.9 | 0.9 |
| AEAVTRAKQIVW | -5.364717 | 1.09775397 | 9 | -6.0653279 | 0.91638239 | 2 | 97.6 | 98.5 | 0.9 |
| YVSNIDGTHIAKTL | -4.9932178 | 1.26024353 | 7 | -5.5106269 | 1.11765253 | 8 | 97.0 | 97.9 | 0.9 |
| MREGIVTATEQEVKEDIAKL | -4.0542758 | 2.36991286 | 2 | -4.315762 | 0.79198193 | 2 | 94.3 | 95.2 | 0.9 |
| QYADPVSAQHAKL | -4.3101783 | 1.10996294 | 7 | -4.6207964 | 1.70457602 | 7 | 95.2 | 96.1 | 0.9 |
| SEKGESSGKNVTLPAVF | -6.0992182 | 2.42113907 | 5 | -7.5117997 | 1.58054575 | 5 | 98.6 | 99.5 | 0.9 |
| VALDFEQEMATAASSSSLEKSYELPDGQVITIGNE | -2.0680468 | 0.22902146 | 12 | -2.1523559 | 0.31453554 | 8 | 80.7 | 81.6 | 0.9 |
| IKTVETRDGQVINETSQHHDDLE | -2.5518687 | 0.46781583 | 12 | -2.6578734 | 0.31897884 | 3 | 85.4 | 86.3 | 0.9 |
| ASQPDVDGFLVGGASLKPEFVDIINAKQ | -5.9694583 | 2.05665928 | 4 | -7.1872692 | 1.95938834 | 3 | 98.4 | 99.3 | 0.9 |
| RLEAPDADELPKGEFDPGQDTY | -2.5108282 | 1.04553772 | 3 | -2.614161 | 0.44673607 | 3 | 85.1 | 86.0 | 0.9 |
| VGSQATNYGEDLTKNHDEL | -4.7629487 | 1.59788077 | 2 | -5.1890581 | 0.62187033 | 2 | 96.4 | 97.3 | 0.9 |
| VARGDLGIEIPAEKVFL | -5.9079578 | 1.25789747 | 3 | -7.0380498 | 0.65118175 | 2 | 98.4 | 99.2 | 0.9 |
| RIEEVPELPLVVEDKVEGY | -2.8270824 | 0.22898141 | 3 | -2.948443 | 0.15537649 | 6 | 87.6 | 88.5 | 0.9 |
| TSSGSANTETTKVTGSLETKY | -6.0113238 | 1.70524324 | 6 | -7.2676549 | 2.33922926 | 3 | 98.5 | 99.4 | 0.9 |
| MIKALELDSNLY | -5.1013649 | 1.42331888 | 2 | -5.6510623 | 1.10120444 | 2 | 97.2 | 98.0 | 0.9 |
| VAEGTRDVPIGAIICITVGKPEDIEAF | -1.4570928 | 0.22885254 | 2 | -1.5226059 | 0.3111144 | 3 | 73.3 | 74.2 | 0.9 |
| SRAKS(79.9663)PQPPVEEEDEHFDDTVVCLDTY | -2.2282906 | 0.08675109 | 6 | -2.3172459 | 0.18083934 | 8 | 82.4 | 83.3 | 0.9 |
| MVTEEDKRTL | -6.3148639 | 1.8166403 | 3 | -8.0941005 | 1.29754756 | 4 | 98.8 | 99.6 | 0.9 |
| EFHLEFLDLVKPEPVY | -2.3234256 | 0.41460268 | 4 | -2.416308 | 0.57874181 | 3 | 83.3 | 84.2 | 0.9 |
| SAAIESCNKALELDSNNEKGLF | -6.2512344 | 1.36854372 | 9 | -7.8821259 | 2.15218036 | 6 | 98.7 | 99.6 | 0.9 |

| | | | | | | | | | |
|---|---|---|---|---|---|---|---|---|---|
| IVSVDETIKNPRSTVDAPT | -6.5354852 | 0.19805378 | 3 | -8.9825399 | 1.13592866 | 4 | 98.9 | 99.8 | 0.9 |
| DATYETKESKKEDLVF | -6.1338238 | 1.44476262 | 9 | -7.5331951 | 2.40171553 | 7 | 98.6 | 99.5 | 0.9 |
| VAARGLDIPEVDLVIQSSPPKDVESY | -3.2923604 | 0.15191802 | 3 | -3.4478142 | 0.19801668 | 4 | 90.7 | 91.6 | 0.9 |
| ELINQLDGFDPRGNIKVL | -5.5551099 | 0.88789803 | 5 | -6.3404941 | 1.14921887 | 6 | 97.9 | 98.8 | 0.9 |
| QVLNADAIVVKL | -6.2897099 | 0.43965216 | 3 | -7.9545303 | 2.10280354 | 6 | 98.7 | 99.6 | 0.9 |
| AGSSRKEAESSPF | -5.5916322 | 0.52022425 | 2 | -6.3987642 | 2.72303153 | 2 | 98.0 | 98.8 | 0.9 |
| VGGLKGDVAEGDLIEHFSQF | -2.8544412 | 0.4931748 | 7 | -2.9744179 | 0.17377984 | 3 | 87.9 | 88.7 | 0.9 |
| GVTIDFDTVNKTPHTATL | -6.5029937 | 0.74202812 | 3 | -8.7482928 | 3.32961381 | 5 | 98.9 | 99.8 | 0.9 |
| AVKLGTITPDGADVY | -4.2418559 | 1.78991143 | 6 | -4.5245853 | 0.36602494 | 3 | 95.0 | 95.8 | 0.9 |
| VTASQCQQPAENKLSDLLAPISEQIKEVITF | -5.0713057 | 0.20953806 | 2 | -5.5894633 | 2.16421228 | 2 | 97.1 | 98.0 | 0.9 |
| TAASSQLKEHFAQF | -2.8414975 | 0.39800759 | 3 | -2.9585121 | 0.75767847 | 2 | 87.8 | 88.6 | 0.8 |
| ELRNRTPSDVKEL | -5.617973 | 1.65957818 | 9 | -6.4223858 | 1.39510161 | 5 | 98.0 | 98.8 | 0.8 |
| EKEAAEMGKGSF | -5.6331604 | 0.65186272 | 5 | -6.4451761 | 2.84003427 | 7 | 98.0 | 98.9 | 0.8 |
| KQIDSSPVGGETDETTVSQNY | -2.640071 | 0.04241258 | 2 | -2.7444462 | 0.35781988 | 2 | 86.2 | 87.0 | 0.8 |
| AKAIANECQANF | -5.8053261 | 1.35698724 | 4 | -6.7510803 | 1.91422833 | 4 | 98.2 | 99.1 | 0.8 |
| GFLGKIDEL | -6.0598179 | 1.26520104 | 9 | -7.2780449 | 1.31495032 | 3 | 98.5 | 99.4 | 0.8 |
| QALKDTANRL | -5.3735093 | 1.96895807 | 5 | -6.0189114 | 2.61733609 | 3 | 97.6 | 98.5 | 0.8 |
| NHFDKDHGGALGPEEF | -5.870513 | 2.99512081 | 4 | -6.8735319 | 2.18016938 | 5 | 98.3 | 99.2 | 0.8 |
| FLEFPKDSSTW | -4.7976811 | 1.68148141 | 3 | -5.2069774 | 1.52767783 | 3 | 96.5 | 97.4 | 0.8 |
| AYTLGVKQL | -5.74714 | 0.60215644 | 2 | -6.634767 | 1.1242494 | 2 | 98.2 | 99.0 | 0.8 |
| LVKIPSEQEQL | -5.7010107 | 1.22764492 | 5 | -6.5518601 | 2.22595167 | 4 | 98.1 | 98.9 | 0.8 |
| FLEPNPEDPLNKEAAEVL | -5.5815492 | 1.07679161 | 5 | -6.3459597 | 0.63444337 | 5 | 98.0 | 98.8 | 0.8 |
| CEKAIEVGRENREDY | -6.4844509 | 0.82547249 | 3 | -8.5098182 | 1.00558983 | 4 | 98.9 | 99.7 | 0.8 |
| DLSDKSINPLGGFVHYGEVTNDF | -3.3841694 | 0.19931583 | 3 | -3.5412724 | 0.25645058 | 2 | 91.3 | 92.1 | 0.8 |
| KNLPIYSEEIVEMY | -5.0552593 | 0.08206208 | 2 | -5.5500637 | 0.15173725 | 4 | 97.1 | 97.9 | 0.8 |
| ENERGELANEVKVL | -3.6877153 | 0.29279278 | 2 | -3.8767844 | 2.66199712 | 3 | 92.8 | 93.6 | 0.8 |
| EILTPNAIPKGF | -5.2775783 | 0.10560544 | 2 | -5.8663819 | 1.2307832 | 6 | 97.5 | 98.3 | 0.8 |
| GFIERGDVVKEIFF | -5.3341578 | 1.25966933 | 4 | -5.9507691 | 1.57389405 | 5 | 97.6 | 98.4 | 0.8 |
| LYPGLQALDEEYLKVDAQF | -2.6346683 | 1.16808448 | 3 | -2.7371594 | 0.00285677 | 2 | 86.1 | 87.0 | 0.8 |
| VIKDLVPDLSNFY | 2.02737971 | 0.20510084 | 8 | 1.95334379 | 0.16411958 | 4 | 19.7 | 20.5 | 0.8 |
| AFVTFDDHDTVDKIVVQKY | -5.2072425 | 1.09882533 | 4 | -5.7587339 | 1.98823121 | 2 | 97.4 | 98.2 | 0.8 |
| NVTNTAGTSLPSVDLLQKL | -4.8767206 | 1.32006399 | 5 | -5.3034532 | 1.13190216 | 2 | 96.7 | 97.5 | 0.8 |
| VKGLSEDTTEETLKESF | -4.9914143 | 1.91931393 | 4 | -5.4565709 | 1.50426808 | 5 | 97.0 | 97.8 | 0.8 |
| VDNKGFPTWL | -6.093661 | 1.544057 | 2 | -7.3165581 | 1.15240454 | 5 | 98.6 | 99.4 | 0.8 |
| VESREKQAKGDTEF | -6.1738803 | 0.21100401 | 2 | -7.4946991 | 1.72365778 | 4 | 98.6 | 99.4 | 0.8 |
| VSISDLLVPKDLGTESQIF | -2.6498239 | 0.21600747 | 6 | -2.7514312 | 0.30281934 | 7 | 86.3 | 87.1 | 0.8 |
| ESLTDPSKLDSGKELH | -6.2389359 | 0.93743334 | 5 | -7.6524129 | 0.90128336 | 3 | 98.7 | 99.5 | 0.8 |
| IAGQVLDINLAAEPKVN | -6.5992307 | 1.53416594 | 3 | -8.8968197 | 1.31867067 | 5 | 99.0 | 99.8 | 0.8 |
| AADESTGSIAKRL | -5.1408848 | 1.27415392 | 12 | -5.6555527 | 1.56933069 | 7 | 97.2 | 98.1 | 0.8 |
| TEGAELVDSVLDVVRKEAESCDCL | -2.3177725 | 0.34206629 | 5 | -2.4032649 | 0.23391237 | 2 | 83.3 | 84.1 | 0.8 |
| DLLEGKEKPVCGTTY | -6.0091108 | 1.82843924 | 5 | -7.1052425 | 2.49527476 | 3 | 98.5 | 99.3 | 0.8 |
| IIGELLKQADLY | -5.6444199 | 0.77349633 | 8 | -6.4221626 | 1.71355537 | 9 | 98.0 | 98.8 | 0.8 |
| FDEISQDTGKY | -6.3231626 | 3.6135183 | 3 | -7.8615647 | 0.88973087 | 2 | 98.8 | 99.6 | 0.8 |
| IDDHFLFDKPVSPL | -6.2495308 | 1.4459244 | 4 | -7.6580654 | 1.62318509 | 6 | 98.7 | 99.5 | 0.8 |
| SEEELKTHISKGTL | -6.0361215 | 2.25693836 | 6 | -7.1544143 | 0.5541329 | 2 | 98.5 | 99.3 | 0.8 |
| GETNPADSKPGTIRGDF | -1.9971016 | 1.13927748 | 9 | -2.0705846 | 1.89056579 | 10 | 80.0 | 80.8 | 0.8 |
| EKPLEEKGEGGEF | -6.6262747 | 2.24276166 | 4 | -8.9638283 | 0.67644749 | 4 | 99.0 | 99.8 | 0.8 |
| REGSLVINSKNQY | -5.3294397 | 1.59006512 | 4 | -5.9176584 | 0.6040023 | 3 | 97.6 | 98.4 | 0.8 |
| VLAPEGSVANKF | -5.1268499 | 1.60354587 | 9 | -5.6261313 | 1.29914446 | 8 | 97.2 | 98.0 | 0.8 |
| NQVLDQFGEKFQADGTY | -5.9357841 | 1.37754986 | 5 | -6.9355982 | 3.10493076 | 6 | 98.4 | 99.2 | 0.8 |
| KCPNPTCENMNF | -6.4227608 | 1.60259189 | 9 | -8.1242235 | 1.87786181 | 5 | 98.8 | 99.6 | 0.8 |
| FLDTLIKF | -6.5838444 | 1.17562171 | 6 | -8.7187077 | 2.38116271 | 3 | 99.0 | 99.8 | 0.8 |
| VKDGKTLNDELEIIEGM | -4.1490833 | 1.28191105 | 2 | -4.3925079 | 1.53676486 | 3 | 94.7 | 95.5 | 0.8 |
| ESEGTRESAINVAEGKKQAQIL | -6.0886791 | 1.72071773 | 4 | -7.235354 | 1.95953678 | 2 | 98.6 | 99.3 | 0.8 |
| IVAAGVGEFEAGISKNGQTREH | -2.6961233 | 0.45250194 | 3 | -2.7965406 | 0.41259411 | 3 | 86.6 | 87.4 | 0.8 |
| YELSENDLNFIKQSKDGAGF | -6.5382423 | 4.04448391 | 8 | -8.4792174 | 1.50768054 | 5 | 98.9 | 99.7 | 0.8 |
| KAINCPEDIVFPALDIL | -3.5559305 | 0.28616794 | 2 | -3.7203994 | 0.67411491 | 2 | 92.2 | 92.9 | 0.8 |
| DILVTKDNQTRSVGQY | -4.6436467 | 1.31920299 | 4 | -4.9835354 | 0.31721903 | 3 | 96.2 | 96.9 | 0.8 |
| SIKALVQNDTLL | -5.3329635 | 1.67165616 | 5 | -5.9061897 | 1.8360866 | 2 | 97.6 | 98.4 | 0.8 |
| TIDDGIFEVKATAGDTHLGGEDFDNRLVNHFVEE | -2.1431595 | 0.06472046 | 5 | -2.2190492 | 0.08789326 | 6 | 81.5 | 82.3 | 0.8 |
| CDCKLPNSKQSQDEPL | -6.7984822 | 2.13513744 | 4 | -9.79337 | 0.24383057 | 2 | 99.1 | 99.9 | 0.8 |
| QLDPTASISAKVNNSSLIGVGY | -4.0851698 | 0.55577986 | 11 | -4.3133336 | 0.6816799 | 6 | 94.4 | 95.2 | 0.8 |
| ITKEEASGSSVTAEEAKKF | -6.6502336 | 2.54340475 | 5 | -8.8543569 | 2.1777056 | 3 | 99.0 | 99.8 | 0.8 |
| AITIEAMKSDIDEVALQGIEFW | -3.3809724 | 0.43966062 | 5 | -3.5258092 | 0.01915204 | 2 | 91.2 | 92.0 | 0.8 |
| TTVEDLGSKILL | -6.8470427 | 2.71222534 | 4 | -10.091799 | 0.64085819 | 4 | 99.1 | 99.9 | 0.8 |
| VKPAVVTVGDFPEEDYGLDEI | -2.4802078 | 0.49409994 | 5 | -2.5678938 | 0.19932638 | 4 | 84.8 | 85.6 | 0.8 |
| SVLLEIQKELL | -6.2306465 | 3.14781455 | 4 | -7.488053 | 0.92561734 | 3 | 98.7 | 99.4 | 0.8 |
| LAEVAAGDDKKGIVDQSQQAY | -6.7560497 | 1.85620459 | 8 | -9.3011582 | 0.40069707 | 4 | 99.1 | 99.8 | 0.8 |
| VFTKEDLTEIRDMLL | -6.5037794 | 2.01714255 | 4 | -8.2265744 | 2.13001726 | 6 | 98.9 | 99.7 | 0.8 |
| IADLDAKSGASL | -5.620309 | 0.80819332 | 2 | -6.3159216 | 1.51877873 | 4 | 98.0 | 98.8 | 0.8 |
| TVINQTQKENLR | -6.2482953 | 3.22457205 | 3 | -7.500541 | 1.39296951 | 2 | 98.7 | 99.5 | 0.7 |
| MTISDEWDIPEKQPF | -3.1710926 | 0.34216085 | 5 | -3.2945166 | 0.15781141 | 4 | 90.0 | 90.8 | 0.7 |

| | | | | | | | | | |
|---|---|---|---|---|---|---|---|---|---|
| QKADDGRPFPQ | -5.3041345 | 0.84021536 | 5 | -5.8320952 | 1.43545044 | 5 | 97.5 | 98.3 | 0.7 |
| STVFKDDDDVVIGKVF | -5.7475483 | 1.65067056 | 5 | -6.5064145 | 1.43702514 | 3 | 98.2 | 98.9 | 0.7 |
| TLENAKARLNQY | -3.9543161 | 1.79834243 | 2 | -4.1529787 | 1.1367686 | 2 | 93.9 | 94.7 | 0.7 |
| SLLESNKDLLLTSSY | -5.6641617 | 1.03233872 | 6 | -6.3673884 | 2.20036692 | 6 | 98.1 | 98.8 | 0.7 |
| STINVGLTSIANLPKLN | -5.3336291 | 0.97997849 | 3 | -5.8678954 | 0.65209094 | 4 | 97.6 | 98.3 | 0.7 |
| IDQEELNKTKPIWTRNPDDITQEEY | -6.5135483 | 1.77305876 | 2 | -8.1557807 | 3.32523824 | 3 | 98.9 | 99.7 | 0.7 |
| DVKAPAMF | -6.8464282 | 2.38564381 | 3 | -9.5873358 | 0.58802743 | 4 | 99.1 | 99.9 | 0.7 |
| SGLDMVGVAQTGSGKTL | -2.4298475 | 1.16132537 | 3 | -2.5109538 | 0.27402595 | 5 | 84.3 | 85.1 | 0.7 |
| VGNLGNNGNKTELERAF | -5.9812707 | 1.10288003 | 14 | -6.8992217 | 1.96641466 | 9 | 98.4 | 99.2 | 0.7 |
| KLAVEALSSLDGDLAGRY | -5.8450366 | 1.49834214 | 12 | -6.6534405 | 1.72473473 | 7 | 98.3 | 99.0 | 0.7 |
| KLTTPTYGDLNHLVSATM | -2.8363929 | 0.36317041 | 8 | -2.9356951 | 0.80118434 | 5 | 87.7 | 88.4 | 0.7 |
| IIATKTPSSDVL | -1.3461154 | 0.32012759 | 4 | -1.3977153 | 0.45713682 | 4 | 71.8 | 72.5 | 0.7 |
| VTGTIGEDEDLIKW | -5.2239461 | 0.91098965 | 5 | -5.6944434 | 0.90966459 | 7 | 97.4 | 98.1 | 0.7 |
| MANGQLVKML | -6.2381346 | 0.23435297 | 2 | -7.3814697 | 2.61652026 | 3 | 98.7 | 99.4 | 0.7 |
| KATAVMPDGQF | -5.0687549 | 1.08159428 | 3 | -5.4843821 | 1.09096026 | 5 | 97.1 | 97.8 | 0.7 |
| GEAKIEDLSQQAQLAAAEKF | -5.2533292 | 1.43699114 | 6 | -5.7301216 | 1.50200296 | 5 | 97.4 | 98.2 | 0.7 |
| DDHDTVDKIVVQKY | -6.5041258 | 2.03138787 | 6 | -8.0142111 | 1.61583291 | 5 | 98.9 | 99.6 | 0.7 |
| AGKQLEDGRTLSDY | -5.6013615 | 1.1824476 | 10 | -6.2272872 | 1.08643253 | 9 | 98.0 | 98.7 | 0.7 |
| AADVGKSGTEREF | -5.2274723 | 0.64295397 | 6 | -5.6887584 | 0.3884409 | 4 | 97.4 | 98.1 | 0.7 |
| SIVNLPSNLEKETTH | -5.5151937 | 1.83622509 | 2 | -6.0938584 | 1.46059052 | 3 | 97.9 | 98.6 | 0.7 |
| NIQKESTLHLVL | -5.4929194 | 0.84081852 | 5 | -6.0603926 | 0.51489684 | 3 | 97.8 | 98.5 | 0.7 |
| EAPNQEKVSDY | -5.9381533 | 1.46849506 | 4 | -6.7680176 | 0.49405518 | 4 | 98.4 | 99.1 | 0.7 |
| KILDAVVAQEPL | -6.0004864 | 2.55180393 | 4 | -6.879052 | 1.67195924 | 5 | 98.5 | 99.2 | 0.7 |
| AKALESPERPF | -4.3937929 | 0.50500076 | 2 | -4.6436562 | 0.37217957 | 7 | 95.5 | 96.2 | 0.7 |
| KASGGLPQFGDEYDFY | -5.1320565 | 0.96489469 | 5 | -5.5565664 | 1.16297024 | 4 | 97.2 | 97.9 | 0.7 |
| EMNSEEKLEQSTIVKERGTVY | -5.4117875 | 1.42650876 | 5 | -5.9369988 | 1.48006627 | 4 | 97.7 | 98.4 | 0.7 |
| ATAAGSEDAEKKVL | -5.9498551 | 3.04337444 | 2 | -6.7698207 | 2.15529814 | 5 | 98.4 | 99.1 | 0.7 |
| GILLDQGQLNKY | -4.7170285 | 1.27333196 | 2 | -5.0246673 | 1.01169919 | 2 | 96.3 | 97.0 | 0.7 |
| AALVASKVF | -6.3640604 | 0.29226297 | 2 | -7.5798415 | 3.37423265 | 2 | 98.8 | 99.5 | 0.7 |
| IGGLPNYLNDDQVKELL | -6.6537379 | 1.20202023 | 8 | -8.3555951 | 2.65399497 | 6 | 99.0 | 99.7 | 0.7 |
| VQSEIFPLETPAFAIKEQGF | -2.6252425 | 0.51323307 | 8 | -2.7085533 | 0.44747657 | 4 | 86.1 | 86.7 | 0.7 |
| LIPNATQPESKVF | -6.2685834 | 0.5591168 | 5 | -7.3645211 | 2.42075738 | 4 | 98.7 | 99.4 | 0.7 |
| VGGIKEDTEEHHLRDYF | -5.2400851 | 1.11565877 | 4 | -5.6892778 | 1.14626534 | 2 | 97.4 | 98.1 | 0.7 |
| DQALQQAVVDDDANNAKAVVKTF | -6.3096146 | 1.59139626 | 5 | -7.4447143 | 0.31882936 | 3 | 98.8 | 99.4 | 0.7 |
| GAVAKQGFDFL | -6.2486707 | 1.93515067 | 3 | -7.3067922 | 0.88371073 | 3 | 98.7 | 99.4 | 0.7 |
| GYQLDLPKANLLF | -5.9855816 | 1.33940042 | 9 | -6.8088131 | 0.21689644 | 2 | 98.4 | 99.1 | 0.7 |
| MTEDNKDLIQGKDLL | -5.6878064 | 0.6950804 | 3 | -6.3218939 | 0.73711305 | 2 | 98.1 | 98.8 | 0.7 |
| GEKFEDENF | -5.175865 | 1.13638077 | 8 | -5.5971278 | 1.40118473 | 11 | 97.3 | 98.0 | 0.7 |
| EFLDKLDVVRSFL | -4.128307 | 1.13835075 | 2 | -4.3283556 | 1.66790788 | 4 | 94.6 | 95.3 | 0.7 |
| INENLIVNTDELGRDCLINAAKTSM | -2.3594102 | 0.37490793 | 2 | -2.4306887 | 0.09429296 | 2 | 83.7 | 84.4 | 0.7 |
| VMEVEVDGQKF | -5.9505167 | 1.56417624 | 4 | -6.736734 | 1.23558989 | 5 | 98.4 | 99.1 | 0.7 |
| NLSKEDDVRQY | -6.0806656 | 1.68440345 | 6 | -6.9628478 | 0.79696818 | 6 | 98.5 | 99.2 | 0.7 |
| DVSGYPTLKIF | -6.3739406 | 1.4002073 | 9 | -7.5460158 | 2.10376362 | 7 | 98.8 | 99.5 | 0.7 |
| EHVTSEIGAEEAEEVGVEHLLRDIKDTTVGTL | -2.8698254 | 0.59237661 | 2 | -2.9614978 | 0.43796053 | 2 | 88.0 | 88.6 | 0.7 |
| KSLVASLAEPDFVVTDF | -5.5720837 | 1.58099062 | 9 | -6.1316823 | 1.71452634 | 7 | 97.9 | 98.6 | 0.7 |
| NFGDVGLSAGLQASDLPASASLPAKADLL | -5.0355756 | 0.83690934 | 2 | -5.4045103 | 0.31717496 | 2 | 97.0 | 97.7 | 0.7 |
| EALELRDNDKTRY | -5.4416073 | 0.87348581 | 4 | -5.9451681 | 0.39474614 | 3 | 97.8 | 98.4 | 0.7 |
| RPQVVTDDDGQAPEAKDGSSF | -2.6042027 | 0.05518346 | 6 | -2.6832285 | 0.04909605 | 4 | 85.9 | 86.5 | 0.7 |
| ARVEKIPGGIIEDSCVL | -3.3665958 | 0.77867444 | 3 | -3.4870907 | 0.78311259 | 7 | 91.2 | 91.8 | 0.7 |
| SCTSHKDYPFHEEF | -7.0313836 | 1.32684669 | 5 | -9.8463909 | 0.16884781 | 2 | 99.2 | 99.9 | 0.7 |
| AVNAAQDSTDLVAKL | -6.3317486 | 0.9500564 | 6 | -7.4268797 | 0.69280834 | 4 | 98.8 | 99.4 | 0.6 |
| TGKTITDVINIGIGGSDLGPL | -5.5738673 | 1.87063636 | 11 | -6.1296135 | 0.89994703 | 12 | 97.9 | 98.6 | 0.6 |
| YGDEEKDKGLQTSQDARFY | -6.8797466 | 0.95036848 | 6 | -8.9803627 | 1.15777234 | 3 | 99.2 | 99.8 | 0.6 |
| VLIEGVDQPIVKTATITEPRGNEEAQIF | -1.5880507 | 0.53725722 | 9 | -1.6380695 | 0.10963214 | 7 | 75.0 | 75.7 | 0.6 |
| DLAGRDLTDYLMKILTERGY | -3.8635185 | 0.19246716 | 7 | -4.0252425 | 0.18750343 | 8 | 93.6 | 94.2 | 0.6 |
| FGGVDEKGPQLF | -5.4655571 | 0.6158741 | 5 | -5.9672207 | 0.88583514 | 5 | 97.8 | 98.4 | 0.6 |
| LAEVASGEKKNSVVEASEAAY | -5.5784411 | 2.52606504 | 7 | -6.116391 | 3.08993001 | 2 | 98.0 | 98.6 | 0.6 |
| VGNLPPDIRTKDIEDVFY | -2.8039958 | 0.23917019 | 6 | -2.8885853 | 0.20940888 | 7 | 87.5 | 88.1 | 0.6 |
| MREGIVTATEQEVKEDIAKLL | -6.2929675 | 0.34417259 | 4 | -7.2986575 | 0.83422168 | 2 | 98.7 | 99.4 | 0.6 |
| GTPVPLEGFEDKVFY | -5.3789117 | 0.71390781 | 3 | -5.8355902 | 1.23369418 | 6 | 97.7 | 98.3 | 0.6 |
| SAMTEEAAVAIKAMAK | -5.5632859 | 1.81004101 | 6 | -6.0911543 | 0.27319885 | 2 | 97.9 | 98.6 | 0.6 |
| DENAKTGQATVASGIPAGW | -5.4488239 | 0.85191635 | 2 | -5.930657 | 1.95225 | 4 | 97.8 | 98.4 | 0.6 |
| IIPNVVKY | -5.7328654 | 1.06624219 | 2 | -6.3385487 | 1.27644843 | 2 | 98.2 | 98.8 | 0.6 |
| CGQVFEKSPL | -5.8558387 | 0.56281065 | 8 | -6.5270044 | 1.16653008 | 8 | 98.3 | 98.9 | 0.6 |
| SDLTKQISRDY | -5.4424285 | 1.48563393 | 3 | -5.9208359 | 1.38645608 | 6 | 97.8 | 98.4 | 0.6 |
| VVFDSIESAKKF | -4.4511946 | 1.19445887 | 6 | -4.6825116 | 1.29244407 | 4 | 95.6 | 96.3 | 0.6 |
| QNVAKEGVKF | -6.1543795 | 4.11125013 | 3 | -7.0260839 | 0.70425309 | 4 | 98.6 | 99.2 | 0.6 |
| ALVFEAPNQEKVSDY | -6.4134983 | 0.93411373 | 6 | -7.5330917 | 1.92402342 | 5 | 98.8 | 99.5 | 0.6 |
| AKPNEGAIVEVALEGYY | -5.2787922 | 1.02983015 | 6 | -5.6980927 | 0.76798797 | 8 | 97.5 | 98.1 | 0.6 |
| EGERAMTKDNNLLGRF | -6.4092008 | 0.70710678 | 2 | -7.5120793 | 0.87939716 | 2 | 98.8 | 99.5 | 0.6 |
| RDLVRNSQDDYDEERKTNL | -4.2565526 | 0.71951571 | 2 | -4.4565134 | 0.87881464 | 3 | 95.0 | 95.6 | 0.6 |
| SVNSSSESLNHLLYDEFVKSVL | -3.9558679 | 0.92308031 | 2 | -4.1199171 | 0.14231203 | 2 | 93.9 | 94.6 | 0.6 |

| | | | | | | | | | |
|---|---|---|---|---|---|---|---|---|---|
| AKDAASLESQLQDTQELLQEETRQKL | -6.1081464 | 1.20021758 | 4 | -6.9259663 | 0.23650632 | 4 | 98.6 | 99.2 | 0.6 |
| TLVALPKEDPTAVACTF | -5.6945622 | 1.30031075 | 6 | -6.2630356 | 0.26627428 | 5 | 98.1 | 98.7 | 0.6 |
| TIKAFIPAIDSF | -6.7664312 | 1.57098784 | 7 | -8.3667108 | 2.54528218 | 3 | 99.1 | 99.7 | 0.6 |
| YGDEEKDKGLQTSQDARFY | -6.7586234 | 2.72256248 | 7 | -8.3210441 | 0.1466837 | 2 | 99.1 | 99.7 | 0.6 |
| SDYNIQKESTLHLVL | -5.5002361 | 0.83491627 | 10 | -5.978039 | 1.34648251 | 3 | 97.8 | 98.4 | 0.6 |
| IVTDRETGSSKGF | -5.9745442 | 1.01977366 | 4 | -6.6727142 | 0.2955789 | 2 | 98.4 | 99.0 | 0.6 |
| MIGAQTDQTVQEHLIEKY | -3.4866107 | 0.34563224 | 5 | -3.6045648 | 0.27646648 | 5 | 91.8 | 92.4 | 0.6 |
| VGLPVEEAVKGILEQGW | -5.7175475 | 2.01848589 | 4 | -6.2792596 | 0.13049094 | 3 | 98.1 | 98.7 | 0.6 |
| LLQNSVKQY | -5.9249952 | 1.75078836 | 4 | -6.5906616 | 1.34716587 | 4 | 98.4 | 99.0 | 0.6 |
| SNEEAQKAQAL | -6.4252823 | 0.82916147 | 2 | -7.4742082 | 1.16187529 | 4 | 98.8 | 99.4 | 0.6 |
| SGGTTMYPGIADRMQKEITAL | -2.8674761 | 0.24530695 | 8 | -2.9496385 | 0.36922159 | 10 | 87.9 | 88.5 | 0.6 |
| CGKETMVTSTTEPSRCEY | -3.7207318 | 0.42408956 | 4 | -3.8560771 | 0.47740429 | 3 | 92.9 | 93.5 | 0.6 |
| NKVTQLTGFSDPVYAEAY | -4.8890511 | 2.29580234 | 3 | -5.1835163 | 1.22425496 | 4 | 96.7 | 97.3 | 0.6 |
| QISKEYERF | -5.7546158 | 1.60426572 | 5 | -6.3222056 | 1.70412559 | 8 | 98.2 | 98.8 | 0.6 |
| KSLVASLAEPDFVVTDF | -6.2482446 | 0.92312277 | 3 | -7.11654 | 1.87272557 | 4 | 98.7 | 99.3 | 0.6 |
| TVKNLSPVVSNELLEQAFSQF | -3.4589168 | 0.67230951 | 8 | -3.5718485 | 0.12414382 | 2 | 91.7 | 92.2 | 0.6 |
| TIDDGIFEVKATAGDTHLGGEDFDNRLVNHFVEE | -2.1157202 | 0.0537546 | 4 | -2.1709874 | 0.03797171 | 10 | 81.3 | 81.8 | 0.6 |
| YSDPQGVPYPEAKIGRF | -6.8751844 | 2.01982966 | 5 | -8.5368381 | 0.86509258 | 5 | 99.2 | 99.7 | 0.6 |
| NTHADFADECPKPELL | -6.2840414 | 1.16498452 | 5 | -7.1641113 | 0.73640676 | 5 | 98.7 | 99.3 | 0.6 |
| EARIAQLEEELEEEQGNTELINDRLKKANL | -4.8310917 | 1.71617349 | 3 | -5.104998 | 0.73168366 | 2 | 96.6 | 97.2 | 0.6 |
| VIKDIEREDIEF | -4.0990169 | 1.49167334 | 9 | -4.2651021 | 0.69552756 | 4 | 94.5 | 95.1 | 0.6 |
| NFVKGVVDSDDLPLNVSRETL | -4.1762436 | 0.49738418 | 4 | -4.3509147 | 0.33189604 | 5 | 94.8 | 95.3 | 0.6 |
| IGGLSFETTDDSLREHFEKWGTL | -3.3046511 | 0.32504954 | 6 | -3.4058457 | 0.2636314 | 6 | 90.8 | 91.4 | 0.6 |
| NTSQSHQTASAVSKVSTN | -6.388687 | 3.30279692 | 3 | -7.3431506 | 3.70896419 | 2 | 98.8 | 99.4 | 0.6 |
| SLGIETLGGVFTKL | -5.8940647 | 1.09613619 | 5 | -6.5041805 | 0.93885086 | 5 | 98.3 | 98.9 | 0.6 |
| TLKLEDTENWLYEDGEDQPKQVY | -5.25102 | 1.46749322 | 3 | -5.614407 | 0.93948549 | 2 | 97.4 | 98.0 | 0.6 |
| EDGCKTVDLKPDWGKGY | -5.8078832 | 2.90483732 | 4 | -6.368527 | 0.12383059 | 2 | 98.2 | 98.8 | 0.6 |
| SGNNIELGTACGKY | -6.1124202 | 1.35424831 | 4 | -6.8351829 | 2.31241926 | 6 | 98.6 | 99.1 | 0.6 |
| LLDLLNATGKDSL | -6.6727921 | 1.55594502 | 8 | -7.8940092 | 1.88878966 | 9 | 99.0 | 99.6 | 0.6 |
| VLPLLEDKQPCYILF | -2.1147455 | 0.30225814 | 3 | -2.167531 | 0.32915723 | 4 | 81.2 | 81.8 | 0.6 |
| AVSQEADKCPTLEQY | -6.4094611 | 2.16573113 | 4 | -7.3395881 | 1.2991567 | 6 | 98.8 | 99.4 | 0.5 |
| IKDYPVVSIEDPFDQDDWGAW | -2.4146886 | 0.26811376 | 8 | -2.4750725 | 0.25686796 | 7 | 84.2 | 84.8 | 0.5 |
| ESLTDPSKLDSGKELH | -7.1832984 | 1.62624039 | 5 | -9.5148758 | 1.52591789 | 8 | 99.3 | 99.9 | 0.5 |
| IQVDIGGGQTKTF | -5.496213 | 1.06694161 | 7 | -5.9232939 | 1.19000368 | 6 | 97.8 | 98.4 | 0.5 |
| KTDTESELDLISRL | -5.516311 | 1.49310196 | 2 | -5.9487847 | 0.55768982 | 3 | 97.9 | 98.4 | 0.5 |
| GATLNKDATKAATAAADF | -6.6597596 | 3.6978047 | 5 | -7.8399039 | 2.59235186 | 4 | 99.0 | 99.6 | 0.5 |
| ATIVKDLVAQAPL | -5.3873225 | 2.64028995 | 2 | -5.7773461 | 1.56905136 | 2 | 97.7 | 98.2 | 0.5 |
| NEDNGIIKAF | -5.2145553 | 1.2897311 | 5 | -5.556632 | 0.84908545 | 5 | 97.4 | 97.9 | 0.5 |
| HLNESGDPSSKSTEIKW | -6.3544118 | 1.98485524 | 12 | -7.221608 | 2.29222316 | 3 | 98.8 | 99.3 | 0.5 |
| VELQKEEAQKLLEQY | -7.0920237 | 1.4283363 | 8 | -9.0627141 | 1.17552054 | 5 | 99.3 | 99.8 | 0.5 |
| AAKVPADTEVVCAPPTAY | -2.3330364 | 0.2847426 | 3 | -2.390259 | 0.30220616 | 4 | 83.4 | 84.0 | 0.5 |
| LSINPQKDETLETEKAQY | -6.9402042 | 1.59553888 | 8 | -8.5432755 | 1.58081481 | 6 | 99.2 | 99.7 | 0.5 |
| SGNNIELGTACGKYY | -6.5824648 | 2.39254569 | 2 | -7.6564354 | 1.81038661 | 5 | 99.0 | 99.5 | 0.5 |
| KAELNEFLTRELAEDGY | -5.3214206 | 1.47109572 | 10 | -5.6897209 | 1.28157685 | 4 | 97.6 | 98.1 | 0.5 |
| NDLLQNPLLVPVKVL | -6.0625465 | 0.50350915 | 2 | -6.7270332 | 2.19578259 | 5 | 98.5 | 99.1 | 0.5 |
| KDVESDSAKQF | -6.4339671 | 0.28252622 | 3 | -7.3460158 | 0.62039119 | 4 | 98.9 | 99.4 | 0.5 |
| DDHDSVDKIVIQKY | -6.692359 | 1.80690457 | 7 | -7.8632266 | 1.6286442 | 9 | 99.0 | 99.6 | 0.5 |
| FVGDEDLLEIIGNSKNVAKL | -5.2926292 | 0.93529138 | 4 | -5.6457044 | 0.89439173 | 6 | 97.5 | 98.0 | 0.5 |
| RVDKAAAAAAAL | -5.9180407 | 1.65716326 | 8 | -6.4935955 | 1.13213194 | 4 | 98.4 | 98.9 | 0.5 |
| MIGAQTDQTVQEHLIEKY | -3.5876274 | 0.94958834 | 7 | -3.6985833 | 0.09396917 | 3 | 92.3 | 92.8 | 0.5 |
| KEVDEQMLNVQNKNSSY | -6.583168 | 1.90174405 | 7 | -7.6196339 | 2.1176866 | 5 | 99.0 | 99.5 | 0.5 |
| LNETQTQEITEDIPVKTL | -5.5162503 | 0.72769866 | 3 | -5.9305732 | 1.03756398 | 5 | 97.9 | 98.4 | 0.5 |
| IAKVATAQDDITGDGTTSNVLIIGELLKQADLY | -5.8264445 | 1.81828183 | 4 | -6.35401 | 1.05349008 | 3 | 98.3 | 98.8 | 0.5 |
| DVPVVKVASGNDHLVML | -2.7420384 | 2.09663813 | 6 | -2.8097345 | 0.77481352 | 5 | 87.0 | 87.5 | 0.5 |
| VARGDLGIEIPAEKVF | -5.5690906 | 1.91221953 | 9 | -5.9969521 | 1.17436605 | 7 | 97.9 | 98.5 | 0.5 |
| VGDEAQSKRGIL | -5.9487014 | 1.5606122 | 15 | -6.5276403 | 1.44320375 | 12 | 98.4 | 98.9 | 0.5 |
| EIVFEDPKIPGEKQF | -6.6701954 | 2.09665486 | 5 | -7.7833552 | 2.31043176 | 6 | 99.0 | 99.5 | 0.5 |
| VASSSKDERQEDPYGPQTKEVNEQTHF | -6.6455143 | 0.36068436 | 2 | -7.7304182 | 2.89395077 | 2 | 99.0 | 99.5 | 0.5 |
| VVNDAGRPKVQVEY | -6.113874 | 1.10523751 | 2 | -6.777491 | 0.4420943 | 4 | 98.6 | 99.1 | 0.5 |
| TIDDGIFEVKATAGDTHLGGEDFDNRLVNHFVEE | -2.1296359 | 0.04694126 | 3 | -2.1797304 | 0.05044362 | 4 | 81.4 | 81.9 | 0.5 |
| DLKNPSDSAVHSPF | -4.7688665 | 1.02937769 | 9 | -5.0056902 | 0.94562683 | 6 | 96.5 | 97.0 | 0.5 |
| TVKVEDLTFTSPF | -6.1491768 | 0.89586792 | 5 | -6.8311915 | 1.30064467 | 4 | 98.6 | 99.1 | 0.5 |
| VAIVDPHIKVDSGY | -5.487638 | 1.14229362 | 9 | -5.8873118 | 1.50014051 | 6 | 97.8 | 98.3 | 0.5 |
| RSVGDGETVEFDVVEGEKGAEAANVTGPGGVPV | -6.4218243 | 1.2277411 | 2 | -7.288479 | 0.73504089 | 3 | 98.8 | 99.4 | 0.5 |
| VELQKEEAQKLLEQY | -6.8656411 | 1.26841443 | 6 | -8.2189532 | 2.08176117 | 4 | 99.1 | 99.7 | 0.5 |
| MIPCEKVSTLPAITL | -4.2982861 | 0.14877002 | 3 | -4.468219 | 0.60839925 | 5 | 95.2 | 95.7 | 0.5 |
| LLTMDKLW | -6.1537381 | 2.36434556 | 6 | -6.8270487 | 2.44221539 | 4 | 98.6 | 99.1 | 0.5 |
| GM(15.9949)KTIGYDPIISPEVSASF | -2.8731058 | 0.15779507 | 3 | -2.9443305 | 0.51261695 | 4 | 88.0 | 88.5 | 0.5 |
| GGARPREEVVQKEQE | -6.8005349 | 1.34249874 | 11 | -8.0399683 | 1.52364464 | 6 | 99.1 | 99.6 | 0.5 |
| TAVHDAILEDLVFPSEIVGKR | -5.6535879 | 0.85667609 | 4 | -6.0965017 | 0.73968294 | 2 | 98.1 | 98.6 | 0.5 |
| ELLNQLDGFQPNTQVKVIAATN | -5.0887932 | 0.48059222 | 5 | -5.3770491 | 1.03485006 | 2 | 97.1 | 97.7 | 0.5 |
| SGLEIVPNGITLPVDPEGKITGEAF | -3.333489 | 0.15370743 | 5 | -3.4236435 | 0.10329391 | 3 | 91.0 | 91.5 | 0.5 |

| | | | | | | | | | |
|---|---|---|---|---|---|---|---|---|---|
| IFTTIKAPL | -7.2873084 | 2.39053649 | 4 | -9.4840472 | 0.68127909 | 2 | 99.4 | 99.9 | 0.5 |
| LVHNVKELEVL | -6.4471813 | 1.2910703 | 4 | -7.2844654 | 1.71897972 | 7 | 98.9 | 99.4 | 0.5 |
| AAASIANIVKSSLGPVGLD | -6.2009329 | 1.14607775 | 8 | -6.872463 | 0.95918488 | 7 | 98.7 | 99.2 | 0.5 |
| YELSENDLNFIKQSKDGAGFL | -6.4074147 | 1.92717758 | 9 | -7.2064066 | 2.12360873 | 4 | 98.8 | 99.3 | 0.5 |
| IFDNVAKVW | -5.2616401 | 1.73458725 | 5 | -5.5789435 | 3.11264761 | 2 | 97.5 | 98.0 | 0.5 |
| EVKATAGDTHLGGEDFDNRLVNHFVEEF | -2.4116328 | 0.07499695 | 2 | -2.4654572 | 0.05745123 | 3 | 84.2 | 84.7 | 0.5 |
| GLTFTEKW | -6.3621874 | 1.08585599 | 8 | -7.1216914 | 0.49606468 | 6 | 98.8 | 99.3 | 0.5 |
| KIVSSSDVGHDEYSTQ | -6.5469035 | 0.63360609 | 2 | -7.440995 | 1.42662201 | 2 | 98.9 | 99.4 | 0.5 |
| AISILQQIELDLKATQAL | -6.0019132 | 0.82962627 | 7 | -6.5550542 | 1.248268 | 8 | 98.5 | 98.9 | 0.5 |
| IDKVRSLETENSAL | -5.7557367 | 1.00964626 | 7 | -6.2069838 | 3.53806511 | 2 | 98.2 | 98.7 | 0.5 |
| GLNFGSKEDANVF | -4.2844051 | 1.26763285 | 2 | -4.440166 | 1.03539988 | 6 | 95.1 | 95.6 | 0.5 |
| LAEPVPGIKAEPDESNARY | -6.4478475 | 1.19301142 | 7 | -7.2425188 | 2.19829816 | 5 | 98.9 | 99.3 | 0.5 |
| VFPQDLLEKGLEANNF | -5.7533124 | 0.37959632 | 6 | -6.1965028 | 0.46432775 | 2 | 98.2 | 98.7 | 0.5 |
| GGVKLEDLIVKDGLTDVY | -5.5247853 | 1.82036672 | 2 | -5.895478 | 1.55226005 | 2 | 97.9 | 98.3 | 0.5 |
| LLDLLNATGKDSLTL | -4.3896247 | 1.14816071 | 9 | -4.5550346 | 1.15541375 | 4 | 95.4 | 95.9 | 0.5 |
| VVATEKNVIAAL | -6.9334895 | 2.69409821 | 3 | -8.1946057 | 0.90544696 | 2 | 99.2 | 99.7 | 0.5 |
| IVVIGHVDSGKSTT | -6.7822232 | 0.90774957 | 8 | -7.8411245 | 1.7146826 | 5 | 99.1 | 99.6 | 0.5 |
| SLLESNKDLLL | -7.5719754 | 3.12620045 | 5 | -10.76914 | 1.13720146 | 6 | 99.5 | 99.9 | 0.5 |
| TIGTIDEIQKL | -6.7409716 | 2.15650191 | 4 | -7.7534685 | 2.4335854 | 5 | 99.1 | 99.5 | 0.5 |
| SATMPSDVLEVTKKF | -7.0652807 | 1.14820419 | 3 | -8.4644628 | 2.63191142 | 5 | 99.3 | 99.7 | 0.5 |
| QEVFDDKAKLAPGTIVEVW | -5.3611487 | 0.81868182 | 5 | -5.6745832 | 1.1319731 | 3 | 97.6 | 98.1 | 0.5 |
| IKDYPVVSIEDPFDQDDWGAW | -2.2719064 | 0.14920829 | 8 | -2.3184483 | 0.29176411 | 7 | 82.8 | 83.3 | 0.5 |
| DVSGYPTIKIL | -7.1060606 | 2.55876373 | 9 | -8.5390702 | 1.51276351 | 7 | 99.3 | 99.7 | 0.5 |
| ITGESKEQVANSAF | -6.372623 | 1.1511704 | 4 | -7.0659505 | 0.20196461 | 2 | 98.8 | 99.3 | 0.5 |
| LGVGDPKIGAAIQEELGY | -4.4439636 | 2.2730035 | 2 | -4.6064085 | 1.51153169 | 3 | 95.6 | 96.1 | 0.4 |
| DEILEASDGIM(15.9949)VARGDLGIEIPAEKVF | -3.4010621 | 0.80989202 | 4 | -3.4849271 | 0.07981666 | 2 | 91.4 | 91.8 | 0.4 |
| GQSGGAGSDSNSPGNVQPNSAPSVESHPVLEKL | -2.4388855 | 0.07399808 | 5 | -2.4886729 | 0.2049768 | 2 | 84.4 | 84.9 | 0.4 |
| GFSHLEALLDDSKELQRF | -2.5734066 | 0.45589184 | 4 | -2.6261977 | 0.51708036 | 2 | 85.6 | 86.1 | 0.4 |
| SQELPEDWDKQPVKVL | -6.3346835 | 1.59860047 | 5 | -6.9921144 | 1.09914269 | 6 | 98.8 | 99.2 | 0.4 |
| ASESKDTELAEELLQW | -5.5058234 | 0.58638428 | 5 | -5.8422266 | 1.24100623 | 5 | 97.8 | 98.3 | 0.4 |
| KTEADAEKTFEEKQGTEIDGR | -7.6294407 | 0.8149983 | 4 | -10.628554 | 1.0368434 | 2 | 99.5 | 99.9 | 0.4 |
| VLGRESLDDLTNLVVKLF | -6.1714607 | 0.46435579 | 2 | -6.7346867 | 0.66950279 | 3 | 98.6 | 99.1 | 0.4 |
| VMGVNHEKY | -6.0916288 | 1.27025679 | 5 | -6.6178482 | 1.58032598 | 3 | 98.6 | 99.0 | 0.4 |
| TEAISLCPTEKNVDLSTFY | -3.6793039 | 0.48942916 | 2 | -3.7759037 | 0.35545756 | 7 | 92.8 | 93.2 | 0.4 |
| SKLPIGDVATQY | -6.3620435 | 0.83173511 | 11 | -7.0208866 | 1.4479905 | 6 | 98.8 | 99.2 | 0.4 |
| AKDAASLESQLQDTQELLQEETRQKL | -6.3246318 | 0.53635637 | 4 | -6.962677 | 0.46849169 | 3 | 98.8 | 99.2 | 0.4 |
| KDPGLVDQLVKADSEW | -6.7206588 | 1.40678626 | 8 | -7.6276175 | 1.80804437 | 3 | 99.1 | 99.5 | 0.4 |
| IFAENDVVIPLKDVSLL | -5.5804858 | 1.58032821 | 3 | -5.9320467 | 1.14159432 | 2 | 98.0 | 98.4 | 0.4 |
| GQDEM(15.9949)IDVIGVTKGKGY | -5.4749579 | 1.33303444 | 7 | -5.7980379 | 1.61566288 | 4 | 97.8 | 98.2 | 0.4 |
| TISPLDLAKL | -6.9074171 | 2.05226254 | 10 | -7.9837436 | 1.81912973 | 6 | 99.2 | 99.6 | 0.4 |
| SVAVKCATITPDEARVEEF | -4.0775653 | 0.23164988 | 4 | -4.2001168 | 0.1382753 | 2 | 94.4 | 94.8 | 0.4 |
| RLEAPDADELPKGEFDPGQDTY | -2.5359223 | 0.17510173 | 3 | -2.586127 | 0.06033013 | 2 | 85.3 | 85.7 | 0.4 |
| ELLDYIVNEPPPKLPSGVF | -2.158795 | 0.03940131 | 2 | -2.2007128 | 0.38704065 | 3 | 81.7 | 82.1 | 0.4 |
| RLIVDEAINEDNSVVSLSQPKMDELQLF | -3.6483926 | 0.62986828 | 5 | -3.741207 | 0.31887579 | 2 | 92.6 | 93.0 | 0.4 |
| RGEPEKLGQALTEVY | -5.1872995 | 0.47265183 | 2 | -5.4453923 | 0.15986161 | 2 | 97.3 | 97.8 | 0.4 |
| QLPQGEEEIPEQIANTETTEDVKGRIY | -4.5651711 | 0.26254146 | 3 | -4.7324008 | 0.94729982 | 4 | 95.9 | 96.4 | 0.4 |
| AEGVIPDEAKAL | -6.9943582 | 1.10646982 | 2 | -8.1374675 | 2.35440833 | 2 | 99.2 | 99.6 | 0.4 |
| QLLEPFDKW | -5.1166754 | 1.39987404 | 5 | -5.359515 | 1.55808335 | 3 | 97.2 | 97.6 | 0.4 |
| GPALSVKVM | -6.7191079 | 0.17068809 | 2 | -7.5874313 | 2.77430583 | 4 | 99.1 | 99.5 | 0.4 |
| KSCAHDWVYE | -5.7351976 | 1.33197925 | 5 | -6.1151015 | 1.86204516 | 3 | 98.2 | 98.6 | 0.4 |
| NVTVTKTDKTL | -7.0878141 | 1.39286102 | 9 | -8.3281011 | 0.94628569 | 6 | 99.3 | 99.7 | 0.4 |
| FWEHFDKDGW | -6.6826621 | 0.7180346 | 8 | -7.50863 | 1.39906802 | 8 | 99.0 | 99.5 | 0.4 |
| SANDWQCKTCSNVNW | -3.3601744 | 0.59606183 | 5 | -3.4360768 | 0.28344565 | 3 | 91.1 | 91.5 | 0.4 |
| CVQQLKEFEGKTL | -6.1495945 | 1.59307261 | 4 | -6.6667751 | 1.96932287 | 6 | 98.6 | 99.0 | 0.4 |
| GADDIELLPEAQHKAEVY | -5.32728 | 0.80295973 | 5 | -5.6028125 | 1.41080793 | 3 | 97.6 | 98.0 | 0.4 |
| AEPVPGIKAEPDESNARYF | -3.0923149 | 0.52286859 | 6 | -3.1570326 | 0.47963739 | 4 | 89.5 | 89.9 | 0.4 |
| ALEKLGAVF | -6.5657811 | 1.821903 | 10 | -7.293289 | 2.77384774 | 5 | 99.0 | 99.4 | 0.4 |
| MVVNDAGRPKVQVEY | -5.2554165 | 1.66069775 | 15 | -5.514542 | 1.38852004 | 9 | 97.4 | 97.9 | 0.4 |
| VLQELDNPGAKRIL | -6.2633299 | 0.7494476 | 4 | -6.8211731 | 1.86966876 | 8 | 98.7 | 99.1 | 0.4 |
| LAPKDKPSGDTAAVFEEGGDVDDLLDMI | -5.6967834 | 0.54173666 | 3 | -6.0461027 | 1.16844465 | 5 | 98.1 | 98.5 | 0.4 |
| EVSLADLQNDEVAFRKF | -5.8875069 | 2.2260106 | 5 | -6.2898817 | 1.75879305 | 3 | 98.3 | 98.7 | 0.4 |
| KDLFDPIIEDRHGGY | -6.0769574 | 0.70690358 | 5 | -6.5425069 | 0.87720456 | 5 | 98.5 | 98.9 | 0.4 |
| RKLDPGSEETQTL | -6.2792555 | 1.38734097 | 7 | -6.8269631 | 2.12987311 | 3 | 98.7 | 99.1 | 0.4 |
| LAAAPQVGEKIAF | -5.3957757 | 0.95280757 | 2 | -5.6720324 | 0.46055439 | 2 | 97.7 | 98.1 | 0.4 |
| LDVDANGKPLGRVVL | -0.6047649 | 1.1356954 | 2 | -0.6287024 | 0.77755633 | 5 | 60.3 | 60.7 | 0.4 |
| AAPPEKDILW | -5.2600808 | 0.36503291 | 4 | -5.510289 | 1.0084567 | 3 | 97.5 | 97.9 | 0.4 |
| VVFDDSEPVQKVL | -5.6864435 | 1.33169551 | 5 | -6.0277152 | 0.46063277 | 2 | 98.1 | 98.5 | 0.4 |
| KEQFLDGDGW | -6.310992 | 2.3051963 | 5 | -6.8660765 | 2.19387455 | 6 | 98.8 | 99.2 | 0.4 |
| FICTAKTEW | -6.1120271 | 1.4009533 | 3 | -6.5801591 | 0.16306132 | 3 | 98.6 | 99.0 | 0.4 |
| KTGAAPIIDVVRSGYY | -5.4040591 | 1.43419625 | 3 | -5.6739419 | 0.7516211 | 5 | 97.7 | 98.1 | 0.4 |
| LQDKDVIAINQDPLGKQGY | -7.0315558 | 1.23681485 | 3 | -8.0523829 | 2.70595813 | 2 | 99.2 | 99.6 | 0.4 |
| DINLAAEPKVN | -7.1374832 | 1.42642235 | 2 | -8.2717033 | 1.18594329 | 4 | 99.3 | 99.7 | 0.4 |

| | | | | | | | | | |
|---|---|---|---|---|---|---|---|---|---|
| GPMEEPLVIEKDLPHYF | -2.1029404 | 0.2497834 | 2 | -2.1388083 | 0.12952665 | 2 | 81.1 | 81.5 | 0.4 |
| IVEPSRELAEQTLNNIKQF | -3.0774388 | 0.90665971 | 3 | -3.1354128 | 0.26397651 | 2 | 89.4 | 89.8 | 0.4 |
| TWRKLDPGSEETQTL | -7.2288799 | 1.05109847 | 12 | -8.436626 | 2.16255633 | 2 | 99.3 | 99.7 | 0.4 |
| ISKQEYDESGPSIVHR | -6.810495 | 1.61030938 | 10 | -7.6074424 | 0.52727737 | 2 | 99.1 | 99.5 | 0.4 |
| LAEKYEWDVAEAR | -5.817095 | 0.90228299 | 3 | -6.1644918 | 1.45765811 | 5 | 98.3 | 98.6 | 0.4 |
| AGIDSSSPEVKGYW | -4.9695575 | 0.975268 | 4 | -5.1572494 | 0.81339557 | 4 | 96.9 | 97.3 | 0.4 |
| FEYIEENKY | -6.0895679 | 1.22898116 | 2 | -6.5163149 | 2.69195691 | 2 | 98.6 | 98.9 | 0.4 |
| LAVANDEELNQLLKGVTIASGGVLPNIHPELL | -2.4494404 | 0.09374452 | 2 | -2.4901954 | 0.01914358 | 2 | 84.5 | 84.9 | 0.4 |
| KLVQDVANNTNEEAGDGTTTATVL | -4.4094286 | 0.54657671 | 3 | -4.5370162 | 0.33421478 | 3 | 95.5 | 95.9 | 0.4 |
| VLAEKAAAY | -5.535523 | 1.050405 | 3 | -5.8136152 | 1.22722232 | 5 | 97.9 | 98.3 | 0.4 |
| SVMPSPKVSDTVVEPY | -4.8918578 | 0.54265945 | 5 | -5.0673782 | 1.46504885 | 3 | 96.7 | 97.1 | 0.4 |
| TKVSASTVPTDGSSRNEETPAAPTPAGATGGSSA' | -2.1755356 | 0.15263375 | 5 | -2.2108904 | 0.21172287 | 2 | 81.9 | 82.2 | 0.4 |
| AKNDLAVVDVR | -4.7520786 | 0.36858496 | 7 | -4.9101191 | 1.41335996 | 3 | 96.4 | 96.8 | 0.4 |
| SWEGAFQHVGKAF | -5.2526328 | 1.14454944 | 5 | -5.475766 | 0.8322027 | 3 | 97.4 | 97.8 | 0.4 |
| DISNLDRLGKSEVELVQLVIDGVNY | -4.2605734 | 0.29404029 | 2 | -4.3734964 | 0.31235607 | 4 | 95.0 | 95.4 | 0.4 |
| RYLSEVASGDNKQTTVSNSQQAY | -3.62037 | 0.18842139 | 2 | -3.6958014 | 0.23036483 | 4 | 92.5 | 92.8 | 0.4 |
| LAEVATGEKRATVVESSEKAY | -7.4188094 | 0.27471504 | 3 | -8.7737041 | 1.24696875 | 2 | 99.4 | 99.8 | 0.4 |
| VTEQEKIDKL | -6.6657192 | 2.09831664 | 4 | -7.3107834 | 2.98527348 | 3 | 99.0 | 99.4 | 0.3 |
| LGKDTNGENIAESLVAEGL | -5.3322999 | 0.70710767 | 4 | -5.5611075 | 1.63637113 | 6 | 97.6 | 97.9 | 0.3 |
| KTIVDDPEGFFEQGGW | -5.6795514 | 1.06891026 | 7 | -5.9731085 | 0.95114691 | 4 | 98.1 | 98.4 | 0.3 |
| IVLKEPISVSSEQVL | -6.4437836 | 1.18031076 | 5 | -6.972778 | 1.52258594 | 8 | 98.9 | 99.2 | 0.3 |
| INAIPVASLDPIKEW | -6.2360557 | 2.67836793 | 4 | -6.676754 | 2.1578952 | 5 | 98.7 | 99.0 | 0.3 |
| GSVKDSIVQGF | -5.1662774 | 1.07780709 | 2 | -5.3653767 | 1.47034748 | 2 | 97.3 | 97.6 | 0.3 |
| RTGEEDKKINEELESQY | -7.1201372 | 1.36768228 | 4 | -8.0564956 | 1.75524107 | 3 | 99.3 | 99.6 | 0.3 |
| TVPRKSDLFQDDLYPDTAGPEAALEAEEW | -2.7990478 | 0.18997638 | 6 | -2.8439307 | 0.37733238 | 8 | 87.4 | 87.8 | 0.3 |
| DLAGRDLTDYLMKILTERGY | -3.9612114 | 0.10609227 | 2 | -4.0494824 | 0.17564272 | 6 | 94.0 | 94.3 | 0.3 |
| LAEVATGEKRATVVESSEKAY | -7.4666645 | 2.19105083 | 3 | -8.7924618 | 1.03197399 | 2 | 99.4 | 99.8 | 0.3 |
| GFVTFDDHDPVDKIVLQKY | -7.2917035 | 0.76812999 | 2 | -8.3893755 | 1.98328219 | 3 | 99.4 | 99.7 | 0.3 |
| KFENAAAGNKPEAVEVTFADFDGVLY | -5.9974571 | 0.73290787 | 4 | -6.3547307 | 0.22745085 | 2 | 98.5 | 98.8 | 0.3 |
| ASEAEKAEQINQAAGEASAVL | -4.6132175 | 0.91904097 | 4 | -4.7420565 | 1.38038484 | 6 | 96.1 | 96.4 | 0.3 |
| STSGSSNTDTGKVTGTLETKY | -7.794494 | 1.33375535 | 8 | -9.6263559 | 0.94471823 | 5 | 99.6 | 99.9 | 0.3 |
| VFEAPNQEKVSDY | -6.4788811 | 0.91776798 | 2 | -6.9773585 | 1.06066045 | 3 | 98.9 | 99.2 | 0.3 |
| IVFGEAKIEDLSQQAQL | -6.3603098 | 0.84863344 | 2 | -6.8127516 | 2.16389145 | 2 | 98.8 | 99.1 | 0.3 |
| FKDDANNDPQWSEEQLIAAKF | -4.4498467 | 1.73379007 | 4 | -4.5644085 | 0.56926981 | 3 | 95.6 | 95.9 | 0.3 |
| VGNLPADITEDEFKRLF | -6.320746 | 3.66737115 | 9 | -6.7573028 | 1.11089996 | 4 | 98.8 | 99.1 | 0.3 |
| IRSLVSDIGKEL | -4.0395147 | 0.12741072 | 2 | -4.126361 | 0.09439281 | 2 | 94.3 | 94.6 | 0.3 |
| MTKDGRISVAGVTSSNVGY | -1.2115278 | 0.3582372 | 5 | -1.2332479 | 0.4086669 | 2 | 69.8 | 70.2 | 0.3 |
| ACIGGTNVGEDIRKL | -4.8361171 | 1.59292061 | 6 | -4.9823322 | 1.10954373 | 3 | 96.6 | 96.9 | 0.3 |
| GTVEKAEIIADKQ | -5.8043478 | 0.55034383 | 4 | -6.0947755 | 1.50435567 | 6 | 98.2 | 98.6 | 0.3 |
| EALDCILPPTRPTDKPL | -2.4617047 | 0.16750497 | 5 | -2.4968793 | 0.24761042 | 5 | 84.6 | 85.0 | 0.3 |
| TTQETITNAETAKEW | -6.813889 | 0.78340124 | 4 | -7.4493478 | 1.56465761 | 6 | 99.1 | 99.4 | 0.3 |
| ACPTPKEDGLAQQQTQL | -5.8144163 | 1.34866345 | 10 | -6.1027398 | 1.79890929 | 9 | 98.3 | 98.6 | 0.3 |
| YNEATGGKYVPRAVL | -5.3949286 | 1.17613878 | 13 | -5.6073004 | 1.75479969 | 4 | 97.7 | 98.0 | 0.3 |
| ARPSSEVIKDANLY | -5.9494584 | 1.29168744 | 8 | -6.2676456 | 0.78963375 | 6 | 98.4 | 98.7 | 0.3 |
| EWDVAEARKIW | -7.5101782 | 3.47275145 | 2 | -8.721681 | 0.68883514 | 2 | 99.5 | 99.8 | 0.3 |
| KGSFSEQGINEFLREL | -6.4251768 | 1.56267614 | 2 | -6.8798006 | 0.15243533 | 2 | 98.8 | 99.2 | 0.3 |
| SSNGGVVKGF | -5.1705798 | 1.48544614 | 5 | -5.3469702 | 1.59376357 | 2 | 97.3 | 97.6 | 0.3 |
| EVSLADLQNDEVAFRKF | -6.5212626 | 0.88966856 | 6 | -6.9983748 | 1.31314293 | 3 | 98.9 | 99.2 | 0.3 |
| MLDKLTGVF | -7.7568122 | 2.04971073 | 10 | -9.2848028 | 0.75599432 | 5 | 99.5 | 99.8 | 0.3 |
| TLGVKQL | -5.4130058 | 0.54980652 | 8 | -5.6197863 | 1.64437677 | 9 | 97.7 | 98.0 | 0.3 |
| QERDPSKIKW | -4.4918292 | 2.12310565 | 2 | -4.601805 | 0.72900091 | 2 | 95.7 | 96.0 | 0.3 |
| GGDPIPKSPF | -4.4148859 | 1.56180383 | 22 | -4.518953 | 0.77962947 | 10 | 95.5 | 95.8 | 0.3 |
| GAPEVLENLEVKSGSPAVL | -6.1855945 | 1.56924087 | 5 | -6.5486175 | 2.14244745 | 4 | 98.6 | 98.9 | 0.3 |
| STSGSSNTDTGKVTGTLETKY | -6.1264624 | 1.49591096 | 5 | -6.470738 | 3.12281712 | 2 | 98.6 | 98.9 | 0.3 |
| GEVGKTTGIPIHVF | -5.6971298 | 1.60719909 | 3 | -5.9471602 | 1.06295088 | 7 | 98.1 | 98.4 | 0.3 |
| VLKEGVEY | -6.7335192 | 2.1760447 | 7 | -7.2728898 | 1.01625771 | 4 | 99.1 | 99.4 | 0.3 |
| LLEEDKPEEPTAHAF | -5.7967509 | 1.48350729 | 4 | -6.0568211 | 0.55214862 | 2 | 98.2 | 98.5 | 0.3 |
| SKGPAVGIDLGTTY | -5.5965762 | 0.46031836 | 7 | -5.8196123 | 1.02016916 | 6 | 98.0 | 98.3 | 0.3 |
| LAKAIEPPPLDAVIEAEH | -5.3596096 | 0.95906094 | 4 | -5.5474577 | 0.25174736 | 2 | 97.6 | 97.9 | 0.3 |
| IASAGADALAKVFV | -4.5016854 | 1.72668792 | 7 | -4.6061054 | 1.53622189 | 4 | 95.8 | 96.1 | 0.3 |
| VSDPKRTIAQDYGVL | -6.3251518 | 0.88223422 | 12 | -6.70327 | 0.82080124 | 6 | 98.8 | 99.0 | 0.3 |
| VNAIEKVF | -7.4879684 | 2.06367447 | 8 | -8.51426 | 0.99722316 | 4 | 99.4 | 99.7 | 0.3 |
| ALELPDKHSLAF | -3.8396033 | 0.15942127 | 3 | -3.9068272 | 0.76649875 | 2 | 93.5 | 93.7 | 0.3 |
| AIRNDEELNKL | -7.5693217 | 0.48771942 | 4 | -8.6653314 | 1.64713142 | 3 | 99.5 | 99.8 | 0.3 |
| TFIDKNGETEL | -5.04649 | 0.70742438 | 4 | -5.1939878 | 0.99841337 | 3 | 97.1 | 97.3 | 0.3 |
| QVINDGDKPKVQVSY | -7.785251 | 0.19996252 | 2 | -9.1490614 | 1.04190757 | 3 | 99.5 | 99.8 | 0.3 |
| TKSPVESTTEPPAVR | -5.5475251 | 1.56889199 | 2 | -5.7539891 | 2.14681786 | 2 | 97.9 | 98.2 | 0.3 |
| GVDVTTKEIVLADVIDNDSW | -2.6626958 | 0.17124394 | 6 | -2.6963357 | 0.83177844 | 7 | 86.4 | 86.6 | 0.3 |
| TLLLKIQEY | -6.3054404 | 0.23828194 | 2 | -6.6628161 | 0.332023 | 3 | 98.8 | 99.0 | 0.3 |
| YSDPQGVPYPEAKIGRF | -6.1101929 | 2.33097968 | 5 | -6.4176341 | 2.39258411 | 4 | 98.6 | 98.8 | 0.3 |
| KNLQTVNVDEN | -7.089772 | 1.37060108 | 7 | -7.7619098 | 2.39449327 | 6 | 99.3 | 99.5 | 0.3 |
| VSDREGSDATGDGTKEKPF | -7.6742705 | 1.87201251 | 5 | -8.8442187 | 1.69623679 | 5 | 99.5 | 99.8 | 0.3 |

| | | | | | | | | | |
|---|---|---|---|---|---|---|---|---|---|
| VRNLANTVTEEILEKAF | -6.5189818 | 1.28799437 | 2 | -6.9369812 | 1.47545281 | 2 | 98.9 | 99.2 | 0.3 |
| AISILQQIELDLKATQ | -6.6356666 | 2.22734873 | 5 | -7.0941356 | 1.21974721 | 6 | 99.0 | 99.3 | 0.3 |
| AAAGGVSHDELLDLAKF | -6.0474286 | 1.97371083 | 4 | -6.3373662 | 0.9476003 | 4 | 98.5 | 98.8 | 0.3 |
| GAFVKPAVVTVGDFPEEDYGLDEI | -3.656744 | 0.15594583 | 8 | -3.7139221 | 0.04281781 | 3 | 92.7 | 92.9 | 0.3 |
| CIQALPEFDGKRF | -7.1618099 | 0.82534086 | 3 | -7.8506222 | 1.17469152 | 5 | 99.3 | 99.6 | 0.3 |
| NQKIRDGWQVEEADDWLRY | -3.5141931 | 0.53892604 | 3 | -3.5659793 | 0.07875281 | 2 | 92.0 | 92.2 | 0.3 |
| SNFPISEETIKLL | -5.8332259 | 0.95985558 | 8 | -6.071705 | 1.02012568 | 8 | 98.3 | 98.5 | 0.3 |
| ELSENDLNFIKQSKDGAGF | -6.6757197 | 0.87516774 | 4 | -7.1265495 | 1.10776985 | 4 | 99.0 | 99.3 | 0.3 |
| GTINIVHPKLGSY | -6.4840386 | 0.79941121 | 2 | -6.86702 | 1.72993983 | 7 | 98.9 | 99.2 | 0.3 |
| NVTEQEKIDKL | -7.2323777 | 2.25852741 | 11 | -7.9406884 | 2.76753825 | 9 | 99.3 | 99.6 | 0.3 |
| KGLGTDEDTIIDIITH | -5.9212517 | 1.56621679 | 3 | -6.1700643 | 2.3083095 | 2 | 98.4 | 98.6 | 0.3 |
| LFDTKPL | -6.7080733 | 1.9802026 | 6 | -7.1603723 | 1.68557339 | 2 | 99.1 | 99.3 | 0.3 |
| TTMEKAGAHL | -5.6608011 | 1.77023606 | 7 | -5.8621039 | 1.69670856 | 6 | 98.1 | 98.3 | 0.2 |
| TFDQLALDSPKGCGTVLL | -4.1424852 | 1.03194855 | 3 | -4.2143684 | 0.1521124 | 2 | 94.6 | 94.9 | 0.2 |
| YELSENDLNFIKQSKDGAGF | -7.2006337 | 2.28488096 | 9 | -7.8566138 | 1.80706513 | 5 | 99.3 | 99.6 | 0.2 |
| SLPIKESEIIDF | -7.4370591 | 0.50742002 | 2 | -8.2457068 | 0.42121567 | 2 | 99.4 | 99.7 | 0.2 |
| QAGQCGNQIGAKFWEVISDEHGIDPTGTY | -2.9197395 | 0.3960141 | 22 | -2.9539213 | 0.44942676 | 12 | 88.3 | 88.6 | 0.2 |
| GVQGFPTIKIF | -7.783318 | 1.52544957 | 3 | -8.8890433 | 1.47213806 | 7 | 99.5 | 99.8 | 0.2 |
| EGSPIKVTL | -6.2210281 | 0.34278272 | 5 | -6.5151803 | 2.10443682 | 7 | 98.7 | 98.9 | 0.2 |
| KTEADAEKTFEEKQGTEIDGRSISLY | -7.291236 | 2.72366144 | 5 | -7.9758641 | 2.2191755 | 3 | 99.4 | 99.6 | 0.2 |
| CTKEAAQEAVKLY | -6.4237024 | 1.8806827 | 7 | -6.7619444 | 2.64718451 | 7 | 98.8 | 99.1 | 0.2 |
| KDPGLVDQLVKADSEW | -5.8472667 | 0.66531131 | 6 | -6.0670631 | 1.00588785 | 2 | 98.3 | 98.5 | 0.2 |
| IATNGESAVVQLPKNGPIYDVVW | -3.2116104 | 0.48256232 | 2 | -3.2508854 | 0.14418859 | 2 | 90.3 | 90.5 | 0.2 |
| GAVTNVKVIRDF | -4.8633473 | 1.65962996 | 5 | -4.9728117 | 1.13218987 | 3 | 96.7 | 96.9 | 0.2 |
| KAGADEERAETARL | -5.4893313 | 1.52447056 | 7 | -5.6541778 | 1.79826421 | 8 | 97.8 | 98.1 | 0.2 |
| VQGLGENVTIESVADYFKQIGIIKTN | -6.9924285 | 0.55606178 | 4 | -7.5013557 | 3.31061238 | 4 | 99.2 | 99.5 | 0.2 |
| KTEADAEKTFEEKQGTEIDGRSISLY | -5.9444387 | 1.44045171 | 6 | -6.1699258 | 2.34976732 | 4 | 98.4 | 98.6 | 0.2 |
| YLLSGAGEHLKTDL | -4.7617283 | 1.88069117 | 3 | -4.8597075 | 1.29485203 | 3 | 96.4 | 96.7 | 0.2 |
| QFTDKHGEVCPAGW | -5.278471 | 1.16328453 | 11 | -5.4174785 | 2.05653048 | 6 | 97.5 | 97.7 | 0.2 |
| DISLVVPKDRVAL | -7.4522006 | 1.8374482 | 6 | -8.1823948 | 2.37424406 | 3 | 99.4 | 99.7 | 0.2 |
| YQKADDGRPFPQ | -6.1199705 | 1.65672723 | 5 | -6.3712145 | 1.99435245 | 4 | 98.6 | 98.8 | 0.2 |
| VANLDYKVGW | -6.1463267 | 1.01137926 | 6 | -6.4002064 | 1.27895462 | 5 | 98.6 | 98.8 | 0.2 |
| ISKQEYDESGPSIVH | -7.1161246 | 0.82122125 | 10 | -7.64897 | 0.83948765 | 8 | 99.3 | 99.5 | 0.2 |
| NDSQRQATKDAGTIAGLNVL | -6.4187001 | 0.9908767 | 11 | -6.7233958 | 1.35769152 | 7 | 98.8 | 99.1 | 0.2 |
| NVAEVDKVTGRF | -4.9630032 | 1.16680913 | 7 | -5.0704289 | 0.86064079 | 8 | 96.9 | 97.1 | 0.2 |
| ALKATLVESSTSGF | -4.4247659 | 1.6729022 | 3 | -4.499888 | 0.8802276 | 2 | 95.6 | 95.8 | 0.2 |
| EALGIAQPKGVL | -7.2854051 | 0.74478611 | 4 | -7.8864001 | 1.97604163 | 7 | 99.4 | 99.6 | 0.2 |
| SVVPSPKVSDTVVEPY | -5.541178 | 1.41593685 | 15 | -5.7000721 | 1.26328291 | 12 | 97.9 | 98.1 | 0.2 |
| VLAKMEEDPDDVPHGHITSL | -4.2397784 | 0.30054181 | 2 | -4.3044088 | 0.66340659 | 4 | 95.0 | 95.2 | 0.2 |
| LGAVEEAKKEGGTVVY | -7.3594577 | 0.6534844 | 6 | -7.966283 | 2.11151854 | 7 | 99.4 | 99.6 | 0.2 |
| NLSQVCDGKVSVH | -4.7125951 | 2.30867382 | 3 | -4.7992827 | 0.95866029 | 3 | 96.3 | 96.5 | 0.2 |
| HAIIVDGHSVEELCKAF | -2.5397296 | 0.29301034 | 6 | -2.5636695 | 0.24658327 | 3 | 85.3 | 85.5 | 0.2 |
| TIEDGIFEVKSTAGDTHLGGEDFDNRMVNHF | -2.7430598 | 0.28972858 | 7 | -2.7694463 | 0.16579551 | 6 | 87.0 | 87.2 | 0.2 |
| SELAEDKENY | -7.8657702 | 1.42537739 | 3 | -8.8082135 | 1.93379657 | 5 | 99.6 | 99.8 | 0.2 |
| FLQVKEGILSDEIYCPPETAVL | -4.1573679 | 0.5809015 | 6 | -4.2167821 | 0.53935423 | 3 | 94.7 | 94.9 | 0.2 |
| KTAENATSGETLEENEAGD | -7.0030433 | 0.91033098 | 2 | -7.4455204 | 0.75156281 | 4 | 99.2 | 99.4 | 0.2 |
| LEIQKELLDY | -6.8955667 | 0.25561064 | 2 | -7.2937651 | 2.94152123 | 4 | 99.2 | 99.4 | 0.2 |
| VGGLNFNTDEQALEDHFSSFGPISEVVVVKDRET( | -2.6610176 | 0.1715624 | 4 | -2.685572 | 0.136066 | 4 | 86.3 | 86.5 | 0.2 |
| GISFPDPKML | -5.1966762 | 0.47511285 | 4 | -5.3122154 | 0.62926895 | 8 | 97.3 | 97.5 | 0.2 |
| EEIVKEVSTY | -6.6271793 | 0.99364567 | 5 | -6.9497056 | 0.95395346 | 6 | 99.0 | 99.2 | 0.2 |
| SVEKIDISPVLL | -5.9315928 | 1.83552718 | 3 | -6.124583 | 1.19275121 | 5 | 98.4 | 98.6 | 0.2 |
| KIDDVVNTR | -6.3040337 | 0.87014786 | 10 | -6.5537582 | 1.62861627 | 8 | 98.8 | 98.9 | 0.2 |
| KAADPPAENSSAPEAEQGGAE | -7.1877873 | 1.87216578 | 2 | -7.680969 | 1.48139171 | 4 | 99.3 | 99.5 | 0.2 |
| GPTAEAAQLESSKRF | -6.9443221 | 1.40416285 | 2 | -7.3443459 | 0.22170648 | 2 | 99.2 | 99.4 | 0.2 |
| SRQIKQVEDDIQQLL | -5.5078192 | 0.13052703 | 3 | -5.6462516 | 0.2355204 | 2 | 97.8 | 98.0 | 0.2 |
| AFVTFDDHDSVDKIVIQKY | -7.8666331 | 0.89690564 | 5 | -8.7295881 | 0.99193436 | 4 | 99.6 | 99.8 | 0.2 |
| TDDDKTDHLSWEW | -8.1817682 | 2.5229797 | 2 | -9.340041 | 1.04292506 | 6 | 99.7 | 99.8 | 0.2 |
| ELFADKVPKTAENF | -7.8743885 | 1.85169828 | 9 | -8.7267097 | 1.30039603 | 8 | 99.6 | 99.8 | 0.2 |
| FSWEGAFQHVGKAF | -7.2473036 | 3.84451223 | 2 | -7.7409574 | 0.02836934 | 2 | 99.3 | 99.5 | 0.2 |
| GSWIGPDHDKF | -5.6511131 | 1.96968137 | 9 | -5.7993787 | 1.55022614 | 4 | 98.0 | 98.2 | 0.2 |
| LANLTQSQIALNEKLVNL | -7.6265611 | 1.9685303 | 8 | -8.2929313 | 1.39042068 | 5 | 99.5 | 99.7 | 0.2 |
| STEIAKFL | -7.9817609 | 1.95370503 | 3 | -8.8647731 | 2.50002522 | 4 | 99.6 | 99.8 | 0.2 |
| VAAIKAAQLGF | -6.2053981 | 1.04954808 | 3 | -6.4144794 | 2.92889257 | 2 | 98.7 | 98.8 | 0.2 |
| EGLDWLSNELSKR | -7.6878086 | 2.6178526 | 5 | -8.3485065 | 2.19944601 | 4 | 99.5 | 99.7 | 0.2 |
| AISILQQIELDLKATQ | -7.0746517 | 2.36337438 | 7 | -7.4707372 | 0.75220249 | 6 | 99.3 | 99.4 | 0.2 |
| AFVTFDDHDSVDKIVIQKY | -7.2977547 | 1.5733867 | 6 | -7.7668027 | 1.69630227 | 5 | 99.4 | 99.5 | 0.2 |
| RGDVVPKDVNAAIA | -4.4484013 | 1.38693101 | 2 | -4.5095614 | 1.36361629 | 3 | 95.6 | 95.8 | 0.2 |
| SATEETLQEVFEKATF | -6.118025 | 1.69893237 | 12 | -6.3089813 | 1.14908301 | 13 | 98.6 | 98.8 | 0.2 |
| TDTERLIGDAAKNQVAMNPTNTVF | -5.6248174 | 0.94679357 | 5 | -5.758888 | 2.55485619 | 4 | 98.0 | 98.2 | 0.2 |
| EFHLPLSPEELLKSGGVNQY | -2.6002116 | 0.11796079 | 2 | -2.6208106 | 0.04448504 | 2 | 85.8 | 86.0 | 0.2 |
| IAKIPNFW | -6.6020324 | 1.06191297 | 4 | -6.8697848 | 0.52541687 | 8 | 99.0 | 99.2 | 0.2 |
| GDVQRVKILF | -6.2003529 | 2.26636461 | 8 | -6.3950158 | 1.36690572 | 5 | 98.7 | 98.8 | 0.2 |

| | | | | | | | | | |
|---|---|---|---|---|---|---|---|---|---|
| LRELISNASDALDKIRY | -6.1328864 | 1.60168584 | 3 | -6.3168216 | 2.47293683 | 3 | 98.6 | 98.8 | 0.2 |
| SAEIIYHLFDAFTKY | -5.9673465 | 3.3262815 | 4 | -6.1300508 | 0.77492967 | 3 | 98.4 | 98.6 | 0.2 |
| KEPISVSSEQVL | -5.1265845 | 0.98736776 | 3 | -5.2172916 | 1.56732224 | 6 | 97.2 | 97.4 | 0.2 |
| AEPVPGIKAEPDESNARY | -6.7206065 | 1.78495831 | 7 | -6.9987001 | 2.51748093 | 5 | 99.1 | 99.2 | 0.2 |
| NQVEIKPEMIGHY | -4.1830893 | 0.68435677 | 9 | -4.2306388 | 0.29905577 | 3 | 94.8 | 94.9 | 0.2 |
| ANFESDEVELSYAKNGQDLGVAF | -4.3376551 | 1.02377954 | 10 | -4.389591 | 0.83432589 | 12 | 95.3 | 95.4 | 0.2 |
| LYEDGEDQPKQVY | -5.9278896 | 0.14086821 | 2 | -6.0788146 | 2.15541159 | 3 | 98.4 | 98.5 | 0.2 |
| SIKALVQNDTL | -5.6944615 | 1.46591481 | 7 | -5.8209352 | 1.04308838 | 3 | 98.1 | 98.3 | 0.2 |
| IGVATAADTLVTKF | -5.5093365 | 0.66296176 | 6 | -5.6191952 | 1.14656571 | 6 | 97.9 | 98.0 | 0.2 |
| TAAVEAKQVAQQEAQRAQF | -7.6841431 | 1.61292178 | 12 | -8.2303909 | 2.34544335 | 4 | 99.5 | 99.7 | 0.2 |
| KTDEFQLHTNVNDGTEFGGSIY | -3.8151654 | 0.17999669 | 6 | -3.850025 | 0.35067938 | 3 | 93.4 | 93.5 | 0.1 |
| VALATGEKGFGY | -6.5385382 | 1.13079833 | 9 | -6.7547657 | 2.68495937 | 4 | 98.9 | 99.1 | 0.1 |
| SLVFDPVQKTL | -6.0678433 | 1.23514769 | 9 | -6.2182397 | 1.31781185 | 4 | 98.5 | 98.7 | 0.1 |
| TVINQTQKENL | -7.3772838 | 0.52505467 | 2 | -7.7657856 | 1.96207385 | 8 | 99.4 | 99.5 | 0.1 |
| AVSEGTKAVT | -8.2564513 | 2.27979846 | 6 | -9.0701396 | 1.44383971 | 9 | 99.7 | 99.8 | 0.1 |
| NEGLWEIDNNPKVKF | -5.9399223 | 0.71907765 | 7 | -6.0739382 | 0.8100843 | 4 | 98.4 | 98.5 | 0.1 |
| GPGLEKSGCIVNNLAEF | -5.3417487 | 0.37104872 | 8 | -5.429616 | 1.63338807 | 6 | 97.6 | 97.7 | 0.1 |
| SCKVVCDENGSKGY | -6.8420809 | 1.89695171 | 10 | -7.0905837 | 1.29848492 | 5 | 99.1 | 99.3 | 0.1 |
| TDTERLIGDAAKNQVALNPQNTVF | -4.8169258 | 1.20702512 | 9 | -4.872565 | 0.54172152 | 4 | 96.6 | 96.7 | 0.1 |
| KAEDEVQRERVSAKNALESY | -8.2174776 | 0.65977366 | 3 | -8.8925102 | 2.27163098 | 3 | 99.7 | 99.8 | 0.1 |
| SLPIKESEIIDFF | -7.0248682 | 1.31747469 | 11 | -7.2834798 | 1.18979242 | 7 | 99.2 | 99.4 | 0.1 |
| KNPNIVNFLDSYLVGDELF | -2.8864562 | 0.50081859 | 2 | -2.903498 | 0.13843407 | 2 | 88.1 | 88.2 | 0.1 |
| GLPIEIKVL | -5.0896465 | 2.41068356 | 2 | -5.1544299 | 3.184694 | 3 | 97.1 | 97.3 | 0.1 |
| IEEQKIVVKVL | -8.3374548 | 1.5366997 | 3 | -9.0641708 | 1.27507407 | 2 | 99.7 | 99.8 | 0.1 |
| MFQYDSTHGKF | -8.1983689 | 1.32018625 | 12 | -8.8405808 | 1.12203378 | 6 | 99.7 | 99.8 | 0.1 |
| KLAVEALSSLDGDLAGRY | -7.1856813 | 1.96331905 | 8 | -7.4655623 | 0.72818063 | 4 | 99.3 | 99.4 | 0.1 |
| VAVVIDPTRTISAGKVNL | -6.0714545 | 2.83254336 | 2 | -6.1948565 | 0.89811697 | 2 | 98.5 | 98.7 | 0.1 |
| LVKGGEDLRQDQRVEQLF | -7.5742139 | 2.10822548 | 3 | -7.936746 | 1.80157428 | 3 | 99.5 | 99.6 | 0.1 |
| AISILQQIELDLKATQAL | -6.0587978 | 1.52750951 | 11 | -6.1755772 | 1.27399889 | 9 | 98.5 | 98.6 | 0.1 |
| ALDLDEVIKVY | -6.4662268 | 0.42799157 | 2 | -6.617398 | 2.6973998 | 4 | 98.9 | 99.0 | 0.1 |
| SGLTKGDAVRDVDIIDHHDNTY | -2.5870739 | 0.30920823 | 5 | -2.6000063 | 0.29248236 | 4 | 85.7 | 85.8 | 0.1 |
| YMDYLAALAKAPF | -5.952235 | 0.64457225 | 8 | -6.0535741 | 0.77714203 | 3 | 98.4 | 98.5 | 0.1 |
| LLPGELAKHAVSEGT | -7.4975529 | 1.77338682 | 3 | -7.80782 | 2.51901149 | 4 | 99.4 | 99.6 | 0.1 |
| GIQKELQF | -6.1569803 | 1.5344127 | 4 | -6.2733915 | 0.2369894 | 2 | 98.6 | 98.7 | 0.1 |
| NDELEIIEGMKF | -5.0837735 | 3.30592397 | 4 | -5.1391252 | 0.73621118 | 5 | 97.1 | 97.2 | 0.1 |
| VSIGIVGKDLEF | -4.4869686 | 2.37713492 | 4 | -4.5228394 | 0.86514516 | 3 | 95.7 | 95.8 | 0.1 |
| SSAVDHGSDEVKF | -7.4756208 | 1.78707798 | 2 | -7.7616434 | 1.35191215 | 3 | 99.4 | 99.5 | 0.1 |
| SPKVFIEGADAETF | -5.5116934 | 1.08309509 | 7 | -5.5807503 | 1.0847207 | 4 | 97.9 | 98.0 | 0.1 |
| LRDPSASPGDAGEQAIRQILDEAGKVGEL | -3.5977821 | 0.01976119 | 2 | -3.6179911 | 0.08084621 | 5 | 92.4 | 92.5 | 0.1 |
| ELFADKVPKTAENF | -8.1289243 | 1.06920046 | 9 | -8.5950438 | 1.20769877 | 8 | 99.6 | 99.7 | 0.1 |
| RVQVQDNEGCPVEALVKDNGNGTY | -2.8073688 | 0.14263327 | 2 | -2.8202839 | 0.15777996 | 3 | 87.5 | 87.6 | 0.1 |
| DLSKIGEEQSPEDAEDGPPELLF | -3.8757701 | 0.12825592 | 2 | -3.8994924 | 0.94490223 | 2 | 93.6 | 93.7 | 0.1 |
| LIGDAAKNQVALNPQNTVF | -6.2469473 | 0.9730996 | 10 | -6.3607394 | 1.62385479 | 2 | 98.7 | 98.8 | 0.1 |
| GQDEMIDVIGVTKGKGY | -5.4049405 | 0.7117292 | 7 | -5.4685825 | 1.14353553 | 6 | 97.7 | 97.8 | 0.1 |
| TFAEKYLPALGY | -6.6425756 | 1.05476437 | 8 | -6.7911859 | 1.50579516 | 4 | 99.0 | 99.1 | 0.1 |
| FACEKNENLAANF | -4.6159204 | 1.42334379 | 2 | -4.6525356 | 0.99755418 | 3 | 96.1 | 96.2 | 0.1 |
| AAAAEIDEEPVSKAKQ | -8.7918076 | 1.22904877 | 3 | -9.5764215 | 0.81064715 | 6 | 99.8 | 99.9 | 0.1 |
| QIDQINTDLNLERSHAQKNENARQQL | -2.172282 | 0.43986153 | 3 | -2.1812856 | 0.05413565 | 3 | 81.8 | 81.9 | 0.1 |
| SASEGEEVPQDKAPSHVPF | -6.8509249 | 0.33183434 | 5 | -7.008712 | 1.79658093 | 7 | 99.1 | 99.2 | 0.1 |
| VVLDLPDLPEDKLAQSVQ | -7.180656 | 0.66734247 | 2 | -7.3772708 | 0.91559197 | 5 | 99.3 | 99.4 | 0.1 |
| IQDGKGDVTITNDGATIL | -6.5619319 | 2.08066934 | 4 | -6.687289 | 1.23577288 | 11 | 99.0 | 99.0 | 0.1 |
| TPKELGGLGML | -4.5837559 | 0.33944442 | 2 | -4.6163148 | 0.42650161 | 2 | 96.0 | 96.1 | 0.1 |
| ELHGEGSSSGKATGDETGAKVERADGYEPPVQES | -8.4778969 | 1.00593713 | 5 | -8.991297 | 1.18367378 | 2 | 99.7 | 99.8 | 0.1 |
| TGKDVNFEFPEFQL | -7.4814629 | 1.11240838 | 7 | -7.7150995 | 1.7421327 | 6 | 99.4 | 99.5 | 0.1 |
| VIDNGSGMCKAGF | -6.7756104 | 1.2778828 | 5 | -6.9147741 | 1.06570296 | 7 | 99.1 | 99.2 | 0.1 |
| GFVTFDDHDPVDKIVLQKY | -7.2258501 | 0.7466898 | 3 | -7.4142453 | 1.05243518 | 2 | 99.3 | 99.4 | 0.1 |
| VVGDSSPAVDAVVECNSKLDPTKTTLL | -5.6809145 | 1.93447022 | 4 | -5.7432388 | 1.5173575 | 4 | 98.1 | 98.2 | 0.1 |
| GGVKLEDLIVKDGLTDVY | -5.8789282 | 0.7754386 | 2 | -5.9476028 | 1.99461646 | 4 | 98.3 | 98.4 | 0.1 |
| GQCKAIF | -8.5937037 | 1.48513984 | 2 | -9.0830169 | 1.24842159 | 2 | 99.7 | 99.8 | 0.1 |
| LIEEQKIVVKVL | -9.2062894 | 2.1528166 | 4 | -10.027259 | 0.10647758 | 3 | 99.8 | 99.9 | 0.1 |
| VGGLDEKVSEPLLWELF | -6.6890287 | 1.15161426 | 7 | -6.8023267 | 0.56788116 | 3 | 99.0 | 99.1 | 0.1 |
| TASAGIQVVGDDLTVTNPKR | -5.9056333 | 1.26977691 | 9 | -5.9688137 | 1.7938822 | 4 | 98.4 | 98.4 | 0.1 |
| TSRPVETTLENNEGGQEQGPSVEGLNVPTKATL | -2.7903881 | 0.31147304 | 4 | -2.7994512 | 0.20249939 | 2 | 87.4 | 87.4 | 0.1 |
| VGGLSPDTSEEQIKEYF | -5.3437327 | 0.0919368 | 2 | -5.3843292 | 0.50118904 | 3 | 97.6 | 97.7 | 0.1 |
| MIEMDGTENKSKF | -9.1237738 | 1.03289275 | 4 | -9.7772519 | 0.32654768 | 3 | 99.8 | 99.9 | 0.1 |
| NWGTVKF | -8.8138777 | 2.30381311 | 4 | -9.304826 | 0.8881224 | 3 | 99.8 | 99.8 | 0.1 |
| KCDVDIRKDLY | -7.3026268 | 2.58093283 | 8 | -7.4575412 | 2.45131099 | 7 | 99.4 | 99.4 | 0.1 |
| DLEAEEYVPLPKGDVH | -4.4283858 | 0.28745205 | 2 | -4.4501796 | 0.22045272 | 4 | 95.6 | 95.6 | 0.1 |
| SATEETLQEVFEKATF | -8.0596741 | 1.30057944 | 4 | -8.3220844 | 0.577357 | 7 | 99.6 | 99.7 | 0.1 |
| IGGLSFETTDDSLREHFEKWGTL | -3.4215034 | 0.11536318 | 2 | -3.4329494 | 0.2561407 | 4 | 91.5 | 91.5 | 0.1 |
| ILENKEGLELL | -8.0353241 | 1.17809899 | 8 | -8.2914526 | 1.22905922 | 9 | 99.6 | 99.7 | 0.1 |
| SALLDGKNVNAGGH | -6.4886303 | 1.3592556 | 2 | -6.5711682 | 2.53694459 | 4 | 98.9 | 99.0 | 0.1 |

| | | | | | | | | | |
|---|---|---|---|---|---|---|---|---|---|
| LGASLKDEVL | -6.523223 | 0.84499516 | 4 | -6.6041804 | 1.72212995 | 4 | 98.9 | 99.0 | 0.1 |
| SATKTFVDFF | -7.6633691 | 4.16665698 | 2 | -7.839357 | 0.1284154 | 2 | 99.5 | 99.6 | 0.1 |
| SSAVDHGSDEVKF | -6.0880971 | 1.48186615 | 2 | -6.1433672 | 0.31988936 | 4 | 98.6 | 98.6 | 0.1 |
| SWEGAFQHVGKAF | -6.904712 | 2.04742569 | 5 | -7.0014243 | 1.02098944 | 5 | 99.2 | 99.2 | 0.1 |
| VAIICGSGLGGLTDKLTQAQIF | -2.7516087 | 0.35293983 | 3 | -2.7584197 | 0.19379265 | 3 | 87.1 | 87.1 | 0.1 |
| SVINQKLKDDEVAQL | -7.3847746 | 2.28312336 | 11 | -7.5074569 | 1.81795002 | 9 | 99.4 | 99.5 | 0.0 |
| VGGLSFDTNEQSLEQVFSKY | -3.4040435 | 0.38916645 | 3 | -3.4128799 | 0.12182871 | 3 | 91.4 | 91.4 | 0.0 |
| TSEIGTKQITQSALL | -7.3870603 | 3.64686645 | 2 | -7.5011557 | 2.28842684 | 4 | 99.4 | 99.5 | 0.0 |
| IQENLELVEKGFSNL | -4.8647279 | 1.89838137 | 4 | -4.8836441 | 1.11496042 | 4 | 96.7 | 96.7 | 0.0 |
| SLLITTEGKLW | -7.7578536 | 1.21372212 | 5 | -7.8765554 | 1.77602371 | 5 | 99.5 | 99.6 | 0.0 |
| KDVLEVGELAKL | -7.3546961 | 1.56389564 | 5 | -7.442802 | 2.19070959 | 7 | 99.4 | 99.4 | 0.0 |
| EIDLQKMPL | -5.8045997 | 1.2337157 | 3 | -5.831339 | 3.41863028 | 4 | 98.2 | 98.3 | 0.0 |
| AKNDLAVVDVRIGM | -5.5834949 | 1.80967183 | 2 | -5.6055159 | 1.03484632 | 5 | 98.0 | 98.0 | 0.0 |
| GALKGAVDGGL | -6.6578767 | 1.02870491 | 8 | -6.6981782 | 0.38492782 | 4 | 99.0 | 99.0 | 0.0 |
| KTVAGGAW | -7.745089 | 0.94028105 | 2 | -7.8276956 | 0.73031152 | 3 | 99.5 | 99.6 | 0.0 |
| LAPKIEDEEGS | -7.3857749 | 0.35488831 | 4 | -7.4437349 | 1.40352432 | 7 | 99.4 | 99.4 | 0.0 |
| KVIFLENY | -8.1553901 | 1.81605924 | 7 | -8.2452285 | 0.54748742 | 2 | 99.7 | 99.7 | 0.0 |
| SVDDIVKGINSSNVENQLQATQ | -2.8195047 | 0.36687405 | 3 | -2.8222596 | 0.34795618 | 2 | 87.6 | 87.6 | 0.0 |
| GFVTFDDHDPVDKIVL | -9.0473492 | 1.72391611 | 6 | -9.2117219 | 1.49754684 | 6 | 99.8 | 99.8 | 0.0 |
| KEGIPALDNFLDKL | -11.382964 | 1.37107961 | 3 | -12.448676 | 1.6418511 | 2 | 100.0 | 100.0 | 0.0 |
| SIKKDEEVL | -6.8844899 | 2.82450988 | 2 | -6.9105818 | 1.07822577 | 2 | 99.2 | 99.2 | 0.0 |
| SLDALSKEGIVAL | -6.247846 | 1.70369361 | 9 | -6.263586 | 1.86160256 | 9 | 98.7 | 98.7 | 0.0 |
| KALGFPEGLVIQAY | -4.948367 | 0.25251901 | 3 | -4.9527856 | 1.32215 | 2 | 96.9 | 96.9 | 0.0 |
| ELLDSPGKVLL | -8.8123614 | 1.77773508 | 9 | -8.8667503 | 1.89600129 | 8 | 99.8 | 99.8 | 0.0 |
| AAQTTDGKLPEVTKDVERTDGALL | -5.2341875 | 1.79315241 | 3 | -5.2377946 | 1.54354838 | 4 | 97.4 | 97.4 | 0.0 |
| ILATGSADKTVALW | -6.232296 | 1.65443112 | 10 | -6.2376674 | 0.96350686 | 8 | 98.7 | 98.7 | 0.0 |
| IKNFGEEVDDESLKELF | -3.6568857 | 1.55493382 | 6 | -3.6578322 | 0.41394634 | 2 | 92.7 | 92.7 | 0.0 |
| KNQNSWGTGEDVKVIL | -6.2993836 | 0.65620034 | 3 | -6.3020978 | 1.37799179 | 2 | 98.7 | 98.7 | 0.0 |
| TKLVVDLTDIDPDVAY | -4.7425232 | 1.53003378 | 5 | -4.7415224 | 1.12083746 | 5 | 96.4 | 96.4 | 0.0 |
| KSVTEQGAELSNEERNLL | -5.3346633 | 1.11376995 | 5 | -5.3326343 | 1.16815478 | 7 | 97.6 | 97.6 | 0.0 |
| GNFAKATF | -8.1080492 | 1.90846961 | 10 | -8.0891086 | 1.12813922 | 7 | 99.6 | 99.6 | 0.0 |
| MGDKSENVQDLLLLDVAPL | -2.6844756 | 0.37007919 | 11 | -2.6835457 | 0.3752684 | 4 | 86.5 | 86.5 | 0.0 |
| TAAVEAKQVAQQEAQRAQF | -8.1092294 | 1.91144485 | 11 | -8.0782075 | 2.10858482 | 5 | 99.6 | 99.6 | 0.0 |
| NVPVITGSKDLQNVNITL | -7.5223665 | 1.47050152 | 8 | -7.4969285 | 1.29818461 | 2 | 99.5 | 99.4 | 0.0 |
| VGGLSPDTPEEKIREY | -6.3050051 | 0.90104199 | 9 | -6.2927136 | 1.46934189 | 8 | 98.8 | 98.7 | 0.0 |
| ESKSDALETLGFLNHY | -3.9044382 | 0.63831213 | 2 | -3.9017426 | 0.24928723 | 2 | 93.7 | 93.7 | 0.0 |
| GIGQDIQPKRDLT | -5.3799545 | 0.73699076 | 3 | -5.3714312 | 1.63939367 | 5 | 97.7 | 97.6 | 0.0 |
| GEKRVSISEGDDKIEY | -6.2007179 | 1.34905756 | 5 | -6.185088 | 0.0994236 | 2 | 98.7 | 98.6 | 0.0 |
| HLKEDQTEY | -7.8396389 | 2.7964346 | 4 | -7.7830877 | 2.12545381 | 6 | 99.6 | 99.5 | 0.0 |
| KADLNIILSSPEQLELF | -6.1461503 | 1.05431262 | 7 | -6.1262183 | 1.03348781 | 6 | 98.6 | 98.6 | 0.0 |
| KEVDEQMLNVQNKNSSY | -6.2264269 | 1.9749284 | 4 | -6.2051764 | 0.35126371 | 2 | 98.7 | 98.7 | 0.0 |
| QWPKEQPFGEGVPLDGDDPEHIQW | -3.5301983 | 0.12308582 | 2 | -3.5263283 | 0.00235082 | 2 | 92.0 | 92.0 | 0.0 |
| LLQLGVPVNDKDDAGWSPL | -5.86038 | 1.10012535 | 6 | -5.8425099 | 1.25949002 | 5 | 98.3 | 98.3 | 0.0 |
| DFIPDVKGAVL | -6.0919941 | 1.24545942 | 6 | -6.0694927 | 2.67588133 | 6 | 98.6 | 98.5 | 0.0 |
| SLGKEDGSGDRGDGPF | -7.8399611 | 2.17626869 | 5 | -7.766103 | 2.11222686 | 4 | 99.6 | 99.5 | 0.0 |
| ADALLIIPKVL | -9.9072279 | 0.47420019 | 5 | -9.5780591 | 0.77545033 | 4 | 99.9 | 99.9 | 0.0 |
| VKPHEEYQEAYDECLEEVQSLPLPL | -2.9985633 | 0.2276176 | 5 | -2.9943444 | 0.21042369 | 5 | 88.9 | 88.9 | 0.0 |
| VNHPQVSALLGEEDEEALHYLTRVEVTEFEDIKSG | -1.9453083 | 0.18964985 | 4 | -1.9427197 | 0.19730409 | 3 | 79.4 | 79.4 | 0.0 |
| ELLAKPGQF | -6.5423821 | 1.60545789 | 12 | -6.4993597 | 1.65877055 | 8 | 98.9 | 98.9 | 0.0 |
| STDVSVDEVKALASLM | -5.0825649 | 0.14850773 | 2 | -5.066164 | 0.15910093 | 2 | 97.1 | 97.1 | 0.0 |
| KENNIEQIYPVNAISF | -3.7705721 | 0.86195575 | 5 | -3.7628761 | 0.04749556 | 2 | 93.2 | 93.1 | 0.0 |
| NDPSVQQDIKFLPF | -6.431006 | 1.06875254 | 9 | -6.3823021 | 1.34240177 | 5 | 98.9 | 98.8 | 0.0 |
| YDQALQQAVVDDDANNAKAVVKTF | -7.1519874 | 1.98739917 | 5 | -7.064967 | 0.62792662 | 2 | 99.3 | 99.3 | 0.0 |
| GSAGPPPTGEEDTAEKDEL | -6.1758816 | 0.66830683 | 5 | -6.1289861 | 1.00122372 | 6 | 98.6 | 98.6 | 0.0 |
| VKTRDGSDYEGW | -6.0791385 | 1.18125536 | 2 | -6.034486 | 3.30633305 | 3 | 98.5 | 98.5 | 0.0 |
| NFVKGVVDSDDLPL | -6.1637078 | 1.74635238 | 3 | -6.1132867 | 1.72809983 | 7 | 98.6 | 98.6 | 0.0 |
| GQLPKFQDGDLTLY | -6.845759 | 0.7839009 | 7 | -6.7639422 | 1.95619758 | 5 | 99.1 | 99.1 | 0.0 |
| SFEVEKELF | -6.4489477 | 1.9577478 | 5 | -6.3824231 | 1.4491136 | 6 | 98.9 | 98.8 | -0.1 |
| NFEKPFLW | -7.0192644 | 1.75600437 | 3 | -6.9178615 | 1.40993638 | 5 | 99.2 | 99.2 | -0.1 |
| AMIIDKLEEDINSSMTN | -3.7672251 | 0.86096716 | 2 | -3.7540827 | 0.24999468 | 3 | 93.2 | 93.1 | -0.1 |
| DLQETLVKIQAEHSESGQL | -3.2387858 | 0.4004174 | 4 | -3.2291022 | 0.44450892 | 3 | 90.4 | 90.4 | -0.1 |
| IQDGSQNTNVDKPL | -5.2005251 | 1.79025338 | 3 | -5.1653614 | 0.51118838 | 2 | 97.4 | 97.3 | -0.1 |
| AFIEFASFEDAKEAL | -9.1166759 | 1.20134477 | 4 | -8.62902 | 1.54063218 | 3 | 99.8 | 99.7 | -0.1 |
| SAVVAAQSNWEKVVPENLGLQEGTHEL | -3.2303332 | 0.28977005 | 4 | -3.2182377 | 0.73114928 | 3 | 90.4 | 90.3 | -0.1 |
| TIEDGIFEVKSTAGDTHLGGEDFDNRMVNHF | -2.9308511 | 0.1628965 | 6 | -2.9205734 | 0.5540229 | 7 | 88.4 | 88.3 | -0.1 |
| DDHDPVDKIVLQKY | -7.9284739 | 1.35960523 | 9 | -7.6897241 | 2.73237605 | 6 | 99.6 | 99.5 | -0.1 |
| CFNKPEDK | -7.2797958 | 0.47445626 | 2 | -7.1198079 | 0.78233541 | 2 | 99.4 | 99.3 | -0.1 |
| NDFINKELILF | -5.7029151 | 0.8304698 | 3 | -5.6458271 | 0.66230985 | 4 | 98.1 | 98.0 | -0.1 |
| GYVDFESAEDLEKALELTGL | -2.6656799 | 0.19813211 | 12 | -2.6563668 | 0.33078593 | 9 | 86.4 | 86.3 | -0.1 |
| QVINDGDKPKVQVSY | -8.4221703 | 1.69029233 | 14 | -8.0847797 | 2.07975624 | 10 | 99.7 | 99.6 | -0.1 |
| SQLKEQISDIDDAVRKL | -8.2484729 | 1.83510128 | 4 | -7.9387201 | 2.44668631 | 6 | 99.7 | 99.6 | -0.1 |
| VGGLNPEATEEKIREY | -6.4012789 | 0.86516536 | 5 | -6.3064523 | 1.13891753 | 2 | 98.8 | 98.8 | -0.1 |

| | | | | | | | | | |
|---|---|---|---|---|---|---|---|---|---|
| KNTPWPEAEAIAPQVGNDAVFL | -2.8766582 | 0.30932073 | 4 | -2.8659478 | 0.87666332 | 2 | 88.0 | 87.9 | -0.1 |
| SVDIPLDKTVVNKDVF | -8.5232781 | 1.39781721 | 9 | -8.1317398 | 2.5123514 | 6 | 99.7 | 99.6 | -0.1 |
| TLNVLEDLGDGQKANDDIIVNW | -2.9964976 | 0.04550624 | 2 | -2.9838839 | 0.39151791 | 3 | 88.9 | 88.8 | -0.1 |
| HPEQLITGKEDAANNY | -5.9755463 | 0.97406601 | 16 | -5.895354 | 1.21459224 | 13 | 98.4 | 98.3 | -0.1 |
| EALDCILPPTRPTDKPL | -2.4832554 | 0.1537463 | 5 | -2.4729379 | 0.21225762 | 6 | 84.8 | 84.7 | -0.1 |
| LIPNLALIDKQESEITY | -4.0594229 | 0.29121725 | 2 | -4.0331245 | 0.46778615 | 4 | 94.3 | 94.2 | -0.1 |
| ILGQNGISDLVKVTL | -6.1618727 | 1.21832125 | 7 | -6.0572114 | 0.55092095 | 5 | 98.6 | 98.5 | -0.1 |
| AAFEEPELLAQDIRKF | -7.0993443 | 1.71716765 | 5 | -6.9023175 | 1.13623739 | 2 | 99.3 | 99.2 | -0.1 |
| KVFLENVIRDAVTY | -7.4637322 | 1.75335527 | 3 | -7.2116285 | 1.41047322 | 4 | 99.4 | 99.3 | -0.1 |
| KAVTEQGAELSNEERNLL | -6.2670043 | 1.10466737 | 5 | -6.1490321 | 1.2664771 | 5 | 98.7 | 98.6 | -0.1 |
| VAPEDLRDDIIENAPTTHTEEYSGEEKTW | -3.158743 | 0.12747757 | 3 | -3.1413046 | 0.06506555 | 2 | 89.9 | 89.8 | -0.1 |
| QYDSTHGKF | -8.6539328 | 1.12548258 | 2 | -8.1172043 | 1.8219613 | 3 | 99.8 | 99.6 | -0.1 |
| IVDGRGKATISNDGATIL | -5.7228301 | 1.4571031 | 5 | -5.6368199 | 1.49585583 | 7 | 98.1 | 98.0 | -0.1 |
| LAEVAAGDDKKGIVDQSQQAY | -8.7207658 | 0.8794084 | 6 | -8.1575105 | 1.56618156 | 3 | 99.8 | 99.7 | -0.1 |
| AEKLIQEGKAY | -6.1589082 | 1.74021397 | 5 | -6.0433655 | 0.03834804 | 2 | 98.6 | 98.5 | -0.1 |
| LRELISNSSDALDKIRY | -7.7125499 | 0.83382828 | 6 | -7.3925709 | 1.68634388 | 8 | 99.5 | 99.4 | -0.1 |
| FWEHFDKDGW | -8.3671094 | 1.17939123 | 5 | -7.8900894 | 1.26884885 | 5 | 99.7 | 99.6 | -0.1 |
| LRELISNASDALDKIRY | -8.6823767 | 0.05447624 | 2 | -8.1003285 | 0.89490735 | 2 | 99.8 | 99.6 | -0.1 |
| IDQEELNKTKPIWTRNPDDITNEEYGEFY | -7.989826 | 0.87805628 | 9 | -7.588093 | 2.44584429 | 5 | 99.6 | 99.5 | -0.1 |
| KDAASVDKVLEL | -6.397556 | 0.74312381 | 3 | -6.2483477 | 1.55454361 | 3 | 98.8 | 98.7 | -0.1 |
| VAEYHSEPVEDEKP | -5.7646991 | 0.99964521 | 4 | -5.660664 | 1.63895324 | 3 | 98.2 | 98.1 | -0.1 |
| VGGIKEDTEEY | -6.0822986 | 0.54841865 | 2 | -5.9518267 | 1.49666272 | 4 | 98.5 | 98.4 | -0.1 |
| NVKSPITGNDLSPPVSF | -6.7616684 | 0.42152813 | 6 | -6.558529 | 1.11936044 | 5 | 99.1 | 99.0 | -0.1 |
| ASGSFDKTASVF | -4.9776598 | 1.98607161 | 2 | -4.9125537 | 0.57181048 | 3 | 96.9 | 96.8 | -0.1 |
| GAEVKAEPVEVVAPR | -6.019406 | 1.2847892 | 12 | -5.8912346 | 0.80024293 | 7 | 98.5 | 98.3 | -0.1 |
| ASEEEIGQLVKQMLDDFGPHRY | -2.9210732 | 0.32603347 | 4 | -2.9016977 | 0.29402439 | 4 | 88.3 | 88.2 | -0.1 |
| EKLEGQGDVPTPKQF | -6.0600062 | 2.11905432 | 3 | -5.9245083 | 1.49566985 | 2 | 98.5 | 98.4 | -0.1 |
| GDVQRVKILF | -9.4311355 | 1.30961677 | 6 | -8.4247451 | 1.90090788 | 4 | 99.9 | 99.7 | -0.1 |
| MFQYDSTHGKF | -6.5648075 | 1.12479781 | 10 | -6.3734999 | 0.63965771 | 3 | 99.0 | 98.8 | -0.1 |
| VDLEKDFAAEVVHPGDLKNSVEVALN | -7.2958593 | 0.49214194 | 2 | -6.990918 | 1.00600951 | 3 | 99.4 | 99.2 | -0.1 |
| QTGKDISTNYY | -5.9723991 | 1.20576776 | 2 | -5.8378947 | 1.63970629 | 4 | 98.4 | 98.3 | -0.2 |
| DFLIKGQFL | -7.6036816 | 1.72067574 | 6 | -7.2263988 | 2.02907506 | 5 | 99.5 | 99.3 | -0.2 |
| SKYGPIADVSIVY | -6.0421391 | 1.48747203 | 2 | -5.9001922 | 0.60059516 | 4 | 98.5 | 98.4 | -0.2 |
| GNKFGEPVLAGF | -5.6382958 | 1.66381457 | 5 | -5.5255731 | 1.96165535 | 2 | 98.0 | 97.9 | -0.2 |
| TASAGIQVVGDDLTVTNPKR | -6.9728981 | 1.6863894 | 8 | -6.707465 | 1.84360927 | 3 | 99.2 | 99.1 | -0.2 |
| IIYDDLSKQAVAY | -4.8148994 | 0.93510683 | 5 | -4.7466398 | 0.28507573 | 5 | 96.6 | 96.4 | -0.2 |
| VARGDLGIEIPAEKVFL | -7.5337799 | 0.96248839 | 3 | -7.1537338 | 3.97683988 | 2 | 99.5 | 99.3 | -0.2 |
| NLEDVQPHDLGKVGEVIVTKDDAMLL | -4.7734523 | 1.17247743 | 3 | -4.7062887 | 1.54442539 | 3 | 96.5 | 96.3 | -0.2 |
| CGAAIGTHEDDKY | -7.4326048 | 0.33992076 | 4 | -7.0721499 | 1.55551594 | 6 | 99.4 | 99.3 | -0.2 |
| SDYNIQKESTLHLVL | -6.7281647 | 1.07792353 | 4 | -6.4918438 | 0.71106519 | 3 | 99.1 | 98.9 | -0.2 |
| QLGDVSQKTTW | -3.1378231 | 0.71914244 | 7 | -3.1118488 | 1.09658225 | 4 | 89.8 | 89.6 | -0.2 |
| TLQPKLPITVL | -8.0544192 | 2.70307841 | 2 | -7.5201935 | 1.48538225 | 6 | 99.6 | 99.5 | -0.2 |
| VGNLGTGAGKGELERAF | -5.9194621 | 1.06206812 | 11 | -5.7714607 | 0.67938872 | 7 | 98.4 | 98.2 | -0.2 |
| SQELPEDWDKQPVKVL | -6.8542487 | 2.38604563 | 8 | -6.586833 | 1.72873043 | 3 | 99.1 | 99.0 | -0.2 |
| TTFPRPVTVEPM(15.9949)DQLDDEEGLPEKL | -1.9643578 | 0.20889831 | 4 | -1.9488268 | 0.25113593 | 5 | 79.6 | 79.4 | -0.2 |
| IVDLTVQEKEKY | -7.3781385 | 2.24329663 | 3 | -7.0020731 | 4.19132059 | 2 | 99.4 | 99.2 | -0.2 |
| SEIDIKVADPVVTF | -5.2188985 | 2.395847 | 6 | -5.119874 | 2.04338119 | 6 | 97.4 | 97.2 | -0.2 |
| AGAAVDELGKVL | -6.1697405 | 2.37311586 | 4 | -5.9882848 | 1.68769963 | 7 | 98.6 | 98.4 | -0.2 |
| ILIDEVDKIGRGY | -4.5627338 | 0.88562021 | 4 | -4.4969434 | 0.74788595 | 3 | 95.9 | 95.8 | -0.2 |
| ISKQEYDESGPSIVH | -7.3700333 | 1.19451327 | 16 | -6.9836935 | 1.34378391 | 15 | 99.4 | 99.2 | -0.2 |
| QRVDKDRSGVISDTELQQAL | -3.2614888 | 0.74328352 | 2 | -3.2306347 | 0.20581886 | 2 | 90.6 | 90.4 | -0.2 |
| ADSPSKAGAAPY | -6.737459 | 2.22710677 | 4 | -6.4680748 | 0.52836899 | 4 | 99.1 | 98.9 | -0.2 |
| LRELISNSSDALDKIRY | -7.4148892 | 0.97620663 | 9 | -6.9938513 | 1.61463016 | 7 | 99.4 | 99.2 | -0.2 |
| TPEGERLIGDAAKNQLTSNPENTVF | -5.1914715 | 0.49968944 | 6 | -5.0854693 | 0.55481367 | 5 | 97.3 | 97.1 | -0.2 |
| YLAPKIEDEEGS | -5.8357028 | 1.45915147 | 4 | -5.6659413 | 2.71332771 | 6 | 98.3 | 98.1 | -0.2 |
| CICADFEKVF | -8.3143919 | 2.48744626 | 4 | -7.568772 | 1.3783021 | 3 | 99.7 | 99.5 | -0.2 |
| STGEKGFGY | -7.5282212 | 0.8620644 | 5 | -7.0424819 | 1.48696502 | 5 | 99.5 | 99.2 | -0.2 |
| TDTERLIGDAAKNQVALNPQNTVF | -4.3027039 | 0.6910898 | 12 | -4.2359845 | 0.76115978 | 5 | 95.2 | 95.0 | -0.2 |
| ILSADFPALVVKASGL | -7.9626661 | 1.48777134 | 6 | -7.3243436 | 1.0689335 | 4 | 99.6 | 99.4 | -0.2 |
| CFITYTDEEPVKKLL | -5.5999751 | 1.39193009 | 4 | -5.4466146 | 1.51488586 | 3 | 98.0 | 97.8 | -0.2 |
| VASVHQDLSDDDIKSVF | -6.0347719 | 0.87643706 | 3 | -5.830964 | 2.20519137 | 8 | 98.5 | 98.3 | -0.2 |
| STQDHAAAAIAKAGIPVY | -4.0447878 | 1.7244627 | 10 | -3.985703 | 4.95314722 | 2 | 94.3 | 94.1 | -0.2 |
| AVDVLVSSGEGKAKDAGQRTTIY | -4.2863747 | 1.92946331 | 3 | -4.2174495 | 1.06734094 | 7 | 95.1 | 94.9 | -0.2 |
| SKVLTFY | -6.2548726 | 2.27679051 | 2 | -6.0151297 | 1.04381019 | 2 | 98.7 | 98.5 | -0.2 |
| SVDIPLDKTVVNKDVF | -8.7660264 | 1.54430723 | 8 | -7.7508686 | 1.83833328 | 9 | 99.8 | 99.5 | -0.2 |
| NNVLGEYEEYITKLF | -5.9628083 | 1.11587149 | 6 | -5.7592052 | 0.92798952 | 4 | 98.4 | 98.2 | -0.2 |
| MLGDPGTAKSQLL | -2.7859326 | 0.88468928 | 2 | -2.7554601 | 2.82548655 | 3 | 87.3 | 87.1 | -0.2 |
| NEFPEPIKL | -6.7657527 | 0.63255363 | 2 | -6.4299458 | 1.83430793 | 2 | 99.1 | 98.9 | -0.2 |
| VRLAPDYDALDVANKIGII | -3.1893807 | 0.22040249 | 5 | -3.1511139 | 0.13699687 | 3 | 90.1 | 89.9 | -0.2 |
| SGDKSENVQDLLLLDVTPL | -4.3695834 | 1.34980302 | 12 | -4.2923677 | 1.3516207 | 9 | 95.4 | 95.1 | -0.2 |
| ACDCWDAESKTSY | -7.8260486 | 3.02604317 | 2 | -7.1882119 | 0.29302286 | 3 | 99.6 | 99.3 | -0.2 |
| AEERDRAEAEAREKETKAL | -8.9326565 | 1.40230186 | 4 | -7.7999001 | 1.52169603 | 4 | 99.8 | 99.6 | -0.2 |

| | | | | | | | | | |
|---|---|---|---|---|---|---|---|---|---|
| KLLDFGSLSNL | -6.1925382 | 1.26203649 | 10 | -5.9494252 | 1.57189149 | 6 | 98.7 | 98.4 | -0.2 |
| GIKDDVF | -8.4932989 | 0.00367618 | 2 | -7.56763 | 1.36278772 | 6 | 99.7 | 99.5 | -0.2 |
| LSGLELVKQGAEARVF | -7.7543684 | 0.26359114 | 2 | -7.130425 | 2.68953246 | 2 | 99.5 | 99.3 | -0.2 |
| EETSGVSVGDPVLRTGKPL | -4.5550877 | 1.4498552 | 6 | -4.4658232 | 1.19607772 | 2 | 95.9 | 95.7 | -0.2 |
| VAIIGEQELKDGVIKL | -8.7124421 | 0.97201344 | 5 | -7.6705443 | 1.71819395 | 4 | 99.8 | 99.5 | -0.3 |
| SGLEIVPNGITLPVDPEGKITGEAF | -3.3191249 | 0.24823449 | 8 | -3.2753974 | 0.16022561 | 5 | 90.9 | 90.6 | -0.3 |
| NQVEIKPEMIGHYLGEF | -2.9410513 | 0.01041967 | 2 | -2.9053617 | 0.43322562 | 2 | 88.5 | 88.2 | -0.3 |
| LNVQVKELEANVL | -6.5929337 | 1.77036232 | 2 | -6.2691404 | 1.20962681 | 4 | 99.0 | 98.7 | -0.3 |
| FVMGVNHEKY | -7.4899834 | 2.07046572 | 7 | -6.9351075 | 0.21909167 | 3 | 99.4 | 99.2 | -0.3 |
| FKEQFLDGDGW | -4.6911161 | 0.80028487 | 5 | -4.5906677 | 1.21483952 | 7 | 96.3 | 96.0 | -0.3 |
| DISLVVPKDRVAL | -6.8229966 | 1.36991817 | 3 | -6.4442019 | 0.87211509 | 5 | 99.1 | 98.9 | -0.3 |
| DLNEGKHLY | -6.7995238 | 1.11224485 | 3 | -6.4237011 | 1.30013612 | 4 | 99.1 | 98.8 | -0.3 |
| KLESDYEILERFPGAYL | -2.5268499 | 0.13848188 | 3 | -2.497047 | 0.44896613 | 2 | 85.2 | 85.0 | -0.3 |
| TSVINQKLKDDEVAQL | -6.9728445 | 1.46910852 | 12 | -6.5508218 | 2.00120644 | 8 | 99.2 | 98.9 | -0.3 |
| HAETIKNVRTATESF | -6.262798 | 1.03674305 | 10 | -5.9873047 | 1.16145162 | 7 | 98.7 | 98.4 | -0.3 |
| NTSTGGLLLPSDTKRSQIY | -7.0577974 | 1.36649869 | 14 | -6.6117057 | 1.33284506 | 6 | 99.3 | 99.0 | -0.3 |
| IDQEELNKTKPIWTRNPDDITQEEYGEFY | -8.4919806 | 4.10101058 | 8 | -7.5108749 | 2.03632615 | 6 | 99.7 | 99.5 | -0.3 |
| LFDLKAFF | -7.5606116 | 1.82076837 | 2 | -6.9609036 | 1.2688223 | 2 | 99.5 | 99.2 | -0.3 |
| GFVTFDDHDPVDKIVL | -7.7530441 | 1.36641715 | 6 | -7.0843202 | 1.37987284 | 5 | 99.5 | 99.3 | -0.3 |
| GGLQKVF | -7.3657519 | 3.67700113 | 2 | -6.8244345 | 2.02451525 | 5 | 99.4 | 99.1 | -0.3 |
| FFKDDANNDPQWSEEQLIAAKF | -5.8834965 | 1.90358985 | 7 | -5.6608156 | 1.24550583 | 7 | 98.3 | 98.1 | -0.3 |
| TLGGQKCSVIRDSLLQDGEF | -2.7772672 | 0.42308165 | 4 | -2.7419992 | 0.71555189 | 2 | 87.3 | 87.0 | -0.3 |
| KTEQGGAHF | -5.6318654 | 0.33538226 | 2 | -5.4403619 | 1.66405703 | 3 | 98.0 | 97.7 | -0.3 |
| ALTNIASGTSEQTKAVVDGGAIPAFISLL | -4.2141635 | 0.18232768 | 4 | -4.1338882 | 0.62814889 | 3 | 94.9 | 94.6 | -0.3 |
| SQLKEQISDIDDAVRKL | -7.060834 | 1.9277189 | 6 | -6.5932346 | 2.9160559 | 5 | 99.3 | 99.0 | -0.3 |
| NAAEITDKLGLH | -6.1044836 | 1.18129648 | 8 | -5.8385126 | 2.17915823 | 4 | 98.6 | 98.3 | -0.3 |
| YFDENPYFENKVLSKEF | -4.2308254 | 0.53847837 | 5 | -4.1472595 | 0.95154982 | 2 | 94.9 | 94.7 | -0.3 |
| GYDHLILNVEWAKPSTN | -5.8467776 | 0.63593363 | 3 | -5.6169873 | 0.6394579 | 3 | 98.3 | 98.0 | -0.3 |
| LVQLQEKALF | -7.436511 | 1.65700261 | 7 | -6.836135 | 2.07473186 | 6 | 99.4 | 99.1 | -0.3 |
| KTNEAQAIETARAW | -7.2562596 | 1.88518017 | 4 | -6.708761 | 3.53553391 | 2 | 99.4 | 99.1 | -0.3 |
| GPGLETGQVGKSADFVVEAIGTEVGTLGF | -4.4811025 | 1.05988361 | 3 | -4.3790672 | 0.43931202 | 4 | 95.7 | 95.4 | -0.3 |
| TSVINQKLKDDEVAQL | -6.7810928 | 1.18823692 | 8 | -6.3578906 | 1.0077559 | 2 | 99.1 | 98.8 | -0.3 |
| ALLDQTKTLAESALQLLY | -6.907032 | 1.54989549 | 8 | -6.4510624 | 1.29305246 | 6 | 99.2 | 98.9 | -0.3 |
| LYFDALECLPEDKEVL | -3.8986938 | 0.29628677 | 3 | -3.824935 | 0.53512855 | 3 | 93.7 | 93.4 | -0.3 |
| LTRVEVTEFEDIKSGY | -5.6278198 | 1.45421262 | 13 | -5.4139167 | 1.14512512 | 10 | 98.0 | 97.7 | -0.3 |
| QSITPIEIDMIVGKDREGFF | -2.6986349 | 0.39332671 | 2 | -2.6601463 | 0.39040546 | 5 | 86.7 | 86.3 | -0.3 |
| RLLEELEEGQKGVGDGTVSW | -2.649408 | 0.07102237 | 5 | -2.6116529 | 0.15516743 | 2 | 86.3 | 85.9 | -0.3 |
| GPGLEGGVVGKSADFVVEAIGDDVGTL | -3.6159401 | 0.88322182 | 3 | -3.5519691 | 0.46603745 | 2 | 92.5 | 92.1 | -0.3 |
| AIPQPDLTKL | -4.9030351 | 1.36939222 | 3 | -4.7621903 | 1.68183742 | 2 | 96.8 | 96.4 | -0.3 |
| DGRDGELPVEDDIDLSDVELDDLGKDEL | -1.795822 | 0.24650797 | 5 | -1.7692506 | 0.01822523 | 3 | 77.6 | 77.3 | -0.3 |
| GFINRNDTKEDVF | -6.9657778 | 0.59485666 | 8 | -6.4680325 | 0.72864661 | 3 | 99.2 | 98.9 | -0.3 |
| KAASADSTTEGTPADGFTVL | -5.187556 | 0.99739538 | 2 | -5.0174613 | 1.28917584 | 5 | 97.3 | 97.0 | -0.3 |
| ATILNAGTNTDGFKEQGVTFPSGDIQEQLIR | -2.8822494 | 0.4221031 | 3 | -2.8377284 | 0.94163991 | 2 | 88.1 | 87.7 | -0.3 |
| GLTEAVKVPYPVFESNPEFLY | -2.887242 | 0.25606668 | 5 | -2.8424163 | 0.51149931 | 4 | 88.1 | 87.8 | -0.3 |
| LVVNMKGNDISSGTVL | -5.5647282 | 1.25497278 | 4 | -5.345861 | 0.33455278 | 2 | 97.9 | 97.6 | -0.3 |
| CIQVGRNIIHGSDSVESAEKEIGLW | -3.6156338 | 0.31906864 | 2 | -3.5474681 | 0.28273307 | 2 | 92.5 | 92.1 | -0.3 |
| AKVDATAETDLAKRF | -5.4878443 | 4.83852874 | 2 | -5.2752873 | 1.38016256 | 4 | 97.8 | 97.5 | -0.3 |
| QIACGISQGLADNTVIAKVNNVVW | -4.8364796 | 0.85310436 | 2 | -4.6918239 | 0.14237085 | 3 | 96.6 | 96.3 | -0.3 |
| GPISEVVVVKDRETQ | -6.52617 | 1.90117584 | 9 | -6.1094449 | 0.91542514 | 6 | 98.9 | 98.6 | -0.4 |
| LLSLDDSVDETEAVKRY | -4.7277755 | 1.50687838 | 3 | -4.5874253 | 0.85001804 | 6 | 96.4 | 96.0 | -0.4 |
| HFKVDNDENEHQLSL | -5.2710594 | 3.31210474 | 3 | -5.0749512 | 2.23104532 | 2 | 97.5 | 97.1 | -0.4 |
| LAFPSPEKLL | -5.3244016 | 1.469511 | 3 | -5.1208342 | 0.36682802 | 2 | 97.6 | 97.2 | -0.4 |
| HVDISLTDFIQKY | -7.0870577 | 2.85182526 | 4 | -6.5013346 | 2.12824059 | 3 | 99.3 | 98.9 | -0.4 |
| TFLDLNLELSKF | -4.8093487 | 1.66628292 | 2 | -4.6585187 | 1.65293136 | 3 | 96.6 | 96.2 | -0.4 |
| TIEDGIFEVKSTAGDTHLGGEDFDNRMVNHF | -2.6411494 | 0.06426497 | 2 | -2.597337 | 0.12806877 | 6 | 86.2 | 85.8 | -0.4 |
| SAMTEEAAVAIKAMAK | -8.3341727 | 1.56876581 | 8 | -7.1989717 | 1.70272553 | 9 | 99.7 | 99.3 | -0.4 |
| EVKLFPQETLF | -7.129601 | 2.45803583 | 3 | -6.5197104 | 1.28159639 | 5 | 99.3 | 98.9 | -0.4 |
| GFIERGDVVKEIF | -5.5779731 | 0.72299534 | 3 | -5.3305289 | 0.80465068 | 4 | 97.9 | 97.6 | -0.4 |
| SVAVKCATITPDEARVEEF | -4.2817124 | 0.28570335 | 3 | -4.1693356 | 1.02052296 | 3 | 95.1 | 94.7 | -0.4 |
| TDKVVIGMDVAASEFF | -6.3610313 | 1.46376254 | 8 | -5.9635272 | 1.26767606 | 12 | 98.8 | 98.4 | -0.4 |
| AAVQGPVGTDFKPL | -4.4922273 | 0.82250893 | 9 | -4.3623022 | 0.70410302 | 4 | 95.7 | 95.4 | -0.4 |
| IVAAGVGEFEAGISKNGQTREHALL | -3.1386599 | 1.13998181 | 2 | -3.0789127 | 0.35328755 | 2 | 89.8 | 89.4 | -0.4 |
| TVAENEAETKLQAILEDIQVTLF | -4.3793504 | 0.31859591 | 4 | -4.2562309 | 0.38964477 | 5 | 95.4 | 95.0 | -0.4 |
| SVIAHVDHGKSTL | -5.9914737 | 1.43348784 | 10 | -5.6625902 | 1.89500674 | 7 | 98.5 | 98.1 | -0.4 |
| KDAGTIAGLNVL | -6.5888958 | 1.62252804 | 2 | -6.1202923 | 1.22218065 | 5 | 99.0 | 98.6 | -0.4 |
| DQLLAEEKTISAKY | -6.597495 | 1.43453975 | 4 | -6.1262777 | 1.67230948 | 6 | 99.0 | 98.6 | -0.4 |
| LTRVEVTEFEDIKSGY | -5.5038125 | 0.59229324 | 9 | -5.2578494 | 1.4217986 | 10 | 97.8 | 97.5 | -0.4 |
| VQTKGTGASGSF | -6.3685183 | 1.05417955 | 16 | -5.9548928 | 1.30534295 | 13 | 98.8 | 98.4 | -0.4 |
| VMQIVINPKDNNQF | -5.2084735 | 1.66156116 | 5 | -5.0026664 | 1.49488931 | 5 | 97.4 | 97.0 | -0.4 |
| LQALEKEGSLL | -7.0219464 | 2.8805156 | 4 | -6.4182546 | 2.47854225 | 2 | 99.2 | 98.8 | -0.4 |
| YFEGIKQTF | -6.1497128 | 1.02900672 | 7 | -5.7837496 | 1.47153408 | 6 | 98.6 | 98.2 | -0.4 |
| EIMDGAPVKGESIPIRLF | -2.7083482 | 0.44220755 | 3 | -2.6595411 | 0.44142712 | 3 | 86.7 | 86.3 | -0.4 |

| | | | | | | | | | |
|---|---|---|---|---|---|---|---|---|---|
| ELQNLTAAEVVVPRDQTPDENDQVIVKIIGHFY | -3.1614848 | 0.43982659 | 2 | -3.0993595 | 0.25256119 | 4 | 89.9 | 89.6 | -0.4 |
| SSEVQFGHAGACANQASETAVAKN | -4.5365805 | 0.69099073 | 3 | -4.3964182 | 0.50421325 | 3 | 95.9 | 95.5 | -0.4 |
| NCLPIAAIVDEKIF | -5.4732079 | 0.88789901 | 7 | -5.2246871 | 0.61586264 | 6 | 97.8 | 97.4 | -0.4 |
| GEAKIEDLSQQAQL | -4.3452858 | 0.61242148 | 2 | -4.220095 | 0.56457401 | 3 | 95.3 | 94.9 | -0.4 |
| NSGKVDIVAINDPFIDLNYMVY | -3.2564118 | 0.18456047 | 13 | -3.1894082 | 0.24290623 | 8 | 90.5 | 90.1 | -0.4 |
| KAEIQVGLQPGQGEQLLASEPSWW | -3.2428559 | 0.44736302 | 3 | -3.1755685 | 0.43617561 | 3 | 90.4 | 90.0 | -0.4 |
| KATAVMPDGQFKDISLSDY | -6.5835953 | 0.7582974 | 2 | -6.0854262 | 1.78071839 | 3 | 99.0 | 98.5 | -0.4 |
| LCAATGPSIKIW | -4.9673668 | 1.18291992 | 3 | -4.7778678 | 1.42722697 | 2 | 96.9 | 96.5 | -0.4 |
| ASTCPDDEEIELAYEQVAKALK | -6.0100288 | 1.29281688 | 5 | -5.652717 | 2.99490796 | 2 | 98.5 | 98.1 | -0.4 |
| SSIGEVESAKL | -6.1364247 | 0.47368233 | 2 | -5.7508155 | 2.25996043 | 3 | 98.6 | 98.2 | -0.4 |
| KIGEHTPSAL | -4.8872614 | 1.92394603 | 2 | -4.7050162 | 1.32670042 | 2 | 96.7 | 96.3 | -0.4 |
| VAKVDCTAHSDVC | -7.3059155 | 3.37572859 | 4 | -6.5553101 | 1.69376826 | 4 | 99.4 | 98.9 | -0.4 |
| EAASAIVNLPGCSAKELAPAVSVL | -2.9717549 | 0.39665994 | 4 | -2.9115664 | 0.43354916 | 3 | 88.7 | 88.3 | -0.4 |
| KRQLEEAEEEAQRAN | -7.2126938 | 2.9299137 | 4 | -6.4942316 | 1.88085124 | 4 | 99.3 | 98.9 | -0.4 |
| GEATKQPGITF | -5.1152152 | 0.87920341 | 4 | -4.9034041 | 1.48213097 | 2 | 97.2 | 96.8 | -0.4 |
| ILSADFPALVVKASGL | -9.2034906 | 1.32033145 | 3 | -7.3711207 | 0.88789137 | 5 | 99.8 | 99.4 | -0.4 |
| TTGKIAGAGLLF | -5.3342918 | 0.23238729 | 3 | -5.0891192 | 0.16995385 | 2 | 97.6 | 97.1 | -0.4 |
| NQLLKPTLSEIELF | -5.7162751 | 0.87284704 | 8 | -5.4067894 | 0.64584882 | 6 | 98.1 | 97.7 | -0.4 |
| DNSLKIISNASCTTN | -6.3108576 | 0.50769517 | 4 | -5.8704785 | 1.35150387 | 7 | 98.8 | 98.3 | -0.4 |
| IYDPIKTAQGSL | -4.6787832 | 1.25237005 | 4 | -4.5132074 | 0.34341872 | 3 | 96.2 | 95.8 | -0.4 |
| GVTIDFDTVNKTPH | -5.9480047 | 0.45992587 | 4 | -5.5851678 | 1.61417635 | 4 | 98.4 | 98.0 | -0.4 |
| TEVLKTHGLLV | -7.7015162 | 2.40791599 | 10 | -6.7348139 | 2.28409451 | 7 | 99.5 | 99.1 | -0.5 |
| SGAQIKIANPVEGSSGRQVTITGSAASISLAQY | -2.0756557 | 0.45971079 | 2 | -2.0337688 | 0.49156513 | 3 | 80.8 | 80.4 | -0.5 |
| TIEFTEEYPNKPPTVRF | -2.8007509 | 0.10086489 | 4 | -2.7414207 | 0.29414535 | 6 | 87.4 | 87.0 | -0.5 |
| VQFEDVRDAEDALHNLDRKW | -3.6755207 | 0.82474186 | 2 | -3.5799665 | 0.10967213 | 3 | 92.7 | 92.3 | -0.5 |
| EVKSTAGDTHLGGEDFDNRMVNHF | -3.5120645 | 0.31425588 | 3 | -3.4248751 | 0.55441266 | 4 | 91.9 | 91.5 | -0.5 |
| ALKLLDEDEIRGY | -7.2106803 | 1.41589449 | 7 | -6.4495719 | 2.36268656 | 7 | 99.3 | 98.9 | -0.5 |
| QYADPVSAQHAKL | -6.2826557 | 1.4577079 | 3 | -5.8235098 | 1.58915834 | 4 | 98.7 | 98.3 | -0.5 |
| FKPEELVDY | -7.0265581 | 0.25869556 | 3 | -6.3267779 | 1.27578779 | 7 | 99.2 | 98.8 | -0.5 |
| IAAPSGSAADKVVIEACDELGIILAHTNL | -2.1433475 | 0.62692845 | 4 | -2.0981243 | 0.24297455 | 6 | 81.5 | 81.1 | -0.5 |
| SGYGRLLEVDLKNGY | -6.8169538 | 1.60944667 | 2 | -6.1847353 | 1.96111518 | 4 | 99.1 | 98.6 | -0.5 |
| FGKDISTTL | -6.5084363 | 1.0547956 | 2 | -5.9720161 | 0.159923 | 2 | 98.9 | 98.4 | -0.5 |
| IAGKCGLVPVLAENY | -6.9925241 | 0.31185046 | 2 | -6.2842123 | 0.91790979 | 8 | 99.2 | 98.7 | -0.5 |
| ELQLANPKEF | -5.5455325 | 0.82099422 | 3 | -5.2322208 | 0.89188641 | 4 | 97.9 | 97.4 | -0.5 |
| LTLTQAIDKF | -8.310609 | 1.79938746 | 3 | -6.9283737 | 2.48847246 | 6 | 99.7 | 99.2 | -0.5 |
| VATNVAARGLDIPEVDLVIQSSPPKDVESY | -2.4898035 | 0.14720339 | 6 | -2.433956 | 0.20083852 | 5 | 84.9 | 84.4 | -0.5 |
| YTGEKGQNQDY | -5.4777249 | 1.12632219 | 4 | -5.1704928 | 0.65388061 | 6 | 97.8 | 97.3 | -0.5 |
| RTVSLGAGAKDELHIVEAEAMNY | -2.8430454 | 0.15929939 | 3 | -2.7760974 | 0.35005215 | 2 | 87.8 | 87.3 | -0.5 |
| FLQVKEGILSDEIY | -4.5046471 | 1.5696821 | 2 | -4.332659 | 0.09205567 | 2 | 95.8 | 95.3 | -0.5 |
| GFVDFNSEEDAKAAKEAMEDGEIDGNKVTL | -6.4580132 | 1.00223021 | 2 | -5.9101727 | 1.13061302 | 3 | 98.9 | 98.4 | -0.5 |
| TASAGIQVVGDDLTVTNPKRIA | -3.4132041 | 0.81423934 | 15 | -3.3204938 | 0.41062985 | 10 | 91.4 | 90.9 | -0.5 |
| DLVAGKTRVTL | -5.4943394 | 1.01034479 | 2 | -5.1766594 | 1.67089148 | 2 | 97.8 | 97.3 | -0.5 |
| GPGLEGGVVGKSADFVVEAIGDDVGTLGF | -3.6455504 | 0.89592587 | 5 | -3.5393626 | 0.61602452 | 4 | 92.6 | 92.1 | -0.5 |
| DLSKIGEEQSPEDAEDGPPELL | -3.5130935 | 0.18592526 | 3 | -3.4130147 | 0.539948 | 4 | 91.9 | 91.4 | -0.5 |
| VGLPVEEAVKGILEQGW | -7.7051667 | 1.95978961 | 3 | -6.6202397 | 1.28399772 | 5 | 99.5 | 99.0 | -0.5 |
| NIIIIAEGAIDRNGKPISSSY | -2.6447892 | 0.35072019 | 5 | -2.5808941 | 0.39098195 | 4 | 86.2 | 85.7 | -0.5 |
| TTKPAIIF | -6.8193934 | 4.44966865 | 2 | -6.1188051 | 2.24696758 | 4 | 99.1 | 98.6 | -0.5 |
| VGNLPADITEDEFKRLF | -6.2322262 | 1.67602643 | 10 | -5.7221476 | 1.19022292 | 3 | 98.7 | 98.1 | -0.5 |
| YPSDGVATEKAVEL | -0.7047155 | 0.50391327 | 4 | -0.6712507 | 0.51222403 | 7 | 62.0 | 61.4 | -0.5 |
| AQIKEMVELPL | -5.584139 | 2.68335553 | 3 | -5.2325221 | 1.02807954 | 3 | 98.0 | 97.4 | -0.5 |
| VSHRSGETEDTFIADLVVGLCTGQIKTGAPC | -1.1089348 | 0.0612252 | 2 | -1.0725012 | 0.02070661 | 3 | 68.3 | 67.8 | -0.5 |
| NVEVLIDVLKELNPSLNF | -2.3855165 | 0.07034924 | 2 | -2.327161 | 0.60817903 | 2 | 83.9 | 83.4 | -0.6 |
| IGGLSFETTDDSLREHFEKW | -2.2913027 | 0.43161454 | 7 | -2.2349366 | 0.31515592 | 7 | 83.0 | 82.5 | -0.6 |
| KVTADVINAAEKL | -7.8825246 | 1.95806451 | 5 | -6.6393524 | 2.40792871 | 6 | 99.6 | 99.0 | -0.6 |
| TNDWEDHLAVKHF | -7.8912533 | 1.21500774 | 6 | -6.6393035 | 1.36360596 | 3 | 99.6 | 99.0 | -0.6 |
| KLTSDDVKEQIY | -6.0932253 | 1.48196733 | 4 | -5.5909004 | 2.87115163 | 7 | 98.6 | 98.0 | -0.6 |
| IAKVATAQDDITGDGTTSNVL | -5.8893522 | 1.32455241 | 4 | -5.4377473 | 0.87650237 | 5 | 98.3 | 97.7 | -0.6 |
| STDDVQINDISLQDYIAVKEKYAKY | -5.1034493 | 1.0904016 | 6 | -4.8184308 | 1.03119291 | 2 | 97.2 | 96.6 | -0.6 |
| SGANKEKLEATINELV | -7.2959054 | 1.84010697 | 7 | -6.3254678 | 1.46846173 | 8 | 99.4 | 98.8 | -0.6 |
| INAIPVASLDPIKEW | -4.9958807 | 0.6651723 | 6 | -4.7249939 | 0.68770652 | 6 | 97.0 | 96.4 | -0.6 |
| VAIVDPHIKVDSGY | -5.5160488 | 1.9210228 | 7 | -5.1446606 | 1.09070494 | 3 | 97.9 | 97.3 | -0.6 |
| QVSLQDKTGF | 7.02223778 | 2.39461255 | 6 | 9.40856025 | 1.80063185 | 5 | 0.8 | 0.1 | -0.6 |
| VIAKM(15.9949)DATANDVPSDRY | -7.6342087 | 2.72626689 | 6 | -6.4658297 | 1.00884834 | 3 | 99.5 | 98.9 | -0.6 |
| DFNIIRKDSLPVCPVDASGCF | -2.9896394 | 0.42988894 | 4 | -2.9008521 | 0.90487258 | 3 | 88.8 | 88.2 | -0.6 |
| KDMAIATGGAVFGEEGLTL | -2.3664131 | 0.21595777 | 7 | -2.3006093 | 0.87225809 | 7 | 83.8 | 83.1 | -0.6 |
| DGLDSGKLY | -5.666215 | 2.05850861 | 2 | -5.2489415 | 0.53012791 | 2 | 98.1 | 97.4 | -0.6 |
| EALPELKLVIDQIDNGFF | -2.96851 | 0.07065235 | 2 | -2.8797582 | 0.35530563 | 2 | 88.7 | 88.0 | -0.6 |
| ESLTDPSKLDSGKEL | -7.9019239 | 2.49678496 | 15 | -6.5568987 | 1.98014228 | 9 | 99.6 | 98.9 | -0.6 |
| NSLPAERIQEIQKAIELF | -2.359887 | 0.31668322 | 4 | -2.2936338 | 0.3161754 | 7 | 83.7 | 83.1 | -0.6 |
| MTLLTDALPLLEQKQVIF | -3.1040497 | 0.59155667 | 2 | -3.0077971 | 0.08334309 | 2 | 89.6 | 88.9 | -0.6 |
| RIDFYFDENPYFENKVL | -2.4947774 | 0.16526794 | 8 | -2.4233926 | 0.11632814 | 7 | 84.9 | 84.3 | -0.6 |
| VEEITDLPIKL | -8.7836082 | 2.19694388 | 4 | -6.8122006 | 1.98573413 | 7 | 99.8 | 99.1 | -0.7 |

| | | | | | | | | | |
|---|---|---|---|---|---|---|---|---|---|
| SVETDYTFPLAEKVKAF | -4.9762941 | 1.54193764 | 2 | -4.6874405 | 1.41534243 | 3 | 96.9 | 96.3 | -0.7 |
| MVLITTKNSNILEDLETL | -2.8289364 | 0.33403245 | 7 | -2.7429394 | 0.58225108 | 6 | 87.7 | 87.0 | -0.7 |
| TLKQFAEMY | 0.43960415 | 0.32372721 | 11 | 0.47862883 | 0.73268033 | 6 | 42.4 | 41.8 | -0.7 |
| SNVCDEEMDLAIEASEAAEQGRPPEHTSKF | -2.4779908 | 0.23875146 | 4 | -2.405499 | 0.46251957 | 4 | 84.8 | 84.1 | -0.7 |
| GYVDFESAEDLEKALELTGLKVF | -6.6371279 | 1.78432191 | 2 | -5.8867213 | 3.14390037 | 2 | 99.0 | 98.3 | -0.7 |
| VLSKADVIQATGDAICIF | -2.2045881 | 0.30604051 | 8 | -2.1397824 | 0.48197498 | 3 | 82.2 | 81.5 | -0.7 |
| SVPKVDDEILGFISEATPL | -5.7778997 | 1.45978142 | 6 | -5.3106757 | 2.13005238 | 2 | 98.2 | 97.5 | -0.7 |
| ALRSVLGEADQKGSL | -4.8599514 | 1.45074207 | 7 | -4.5836574 | 1.33290157 | 2 | 96.7 | 96.0 | -0.7 |
| SFEFTEQQKEF | -5.2196392 | 1.85076452 | 3 | -4.8783588 | 0.15186653 | 3 | 97.4 | 96.7 | -0.7 |
| ILNESLTDPAIVKVF | -4.9474834 | 1.27421691 | 7 | -4.6558275 | 0.32529959 | 5 | 96.9 | 96.2 | -0.7 |
| VFDVELLKLE | -7.4247288 | 1.7293402 | 5 | -6.2911212 | 0.80115907 | 6 | 99.4 | 98.7 | -0.7 |
| LDFKVVEGSFVY | -5.0015183 | 1.10436868 | 6 | -4.6981057 | 0.82294896 | 6 | 97.0 | 96.3 | -0.7 |
| ESHKDEIFQVHW | -7.2055778 | 2.72800819 | 3 | -6.1838379 | 1.73493039 | 3 | 99.3 | 98.6 | -0.7 |
| TVKHEQNIDCGGGY | 1.53002296 | 0.23716121 | 5 | 1.58365491 | 0.10219491 | 5 | 25.7 | 25.0 | -0.7 |
| TILIKYGGDEIPFSPY | -5.801507 | 0.46882869 | 4 | -5.302237 | 2.10707033 | 4 | 98.2 | 97.5 | -0.7 |
| SGETAKGDYPLEAVRMQHL | -2.8859185 | 0.71003652 | 6 | -2.7896361 | 0.82694448 | 2 | 88.1 | 87.4 | -0.7 |
| DGIILPGK | -6.8423114 | 1.69381858 | 9 | -5.9435902 | 2.45175738 | 6 | 99.1 | 98.4 | -0.7 |
| NSGKVDIVAINDPFIDLNYMVY | -3.2306962 | 0.14170907 | 11 | -3.1125931 | 0.22174453 | 8 | 90.4 | 89.6 | -0.7 |
| KGVGIISEGNETVEDIAARL | -7.3247238 | 0.42698375 | 2 | -6.1824376 | 1.89467855 | 2 | 99.4 | 98.6 | -0.7 |
| SLQKTTLIPGGSESLVY | -6.6118792 | 0.8238391 | 9 | -5.808807 | 0.58589049 | 3 | 99.0 | 98.2 | -0.7 |
| LISGADDRLVKIW | -4.5704131 | 1.4360625 | 7 | -4.3154008 | 1.92544183 | 3 | 96.0 | 95.2 | -0.7 |
| KADGGTQVIDTKNIL | -6.7206541 | 1.22024056 | 3 | -5.8683258 | 6.15583466 | 3 | 99.1 | 98.3 | -0.7 |
| LAKAVANQTSATF | -4.8886813 | 1.07206322 | 2 | -4.5781097 | 0.8493139 | 2 | 96.7 | 96.0 | -0.8 |
| ELRDNDKTRY | -7.1947791 | 0.13142749 | 2 | -6.1019266 | 1.32402532 | 3 | 99.3 | 98.6 | -0.8 |
| KGQPAIIDGELY | -3.6147887 | 0.24541483 | 4 | -3.4648303 | 1.48876204 | 2 | 92.5 | 91.7 | -0.8 |
| SLEEIKQATGIEDGELRRTL | -2.7917788 | 0.33935589 | 2 | -2.6947542 | 0.19914937 | 2 | 87.4 | 86.6 | -0.8 |
| KQPAENVNQY | -7.0388015 | 0.41599646 | 4 | -6.0223099 | 1.06138537 | 3 | 99.2 | 98.5 | -0.8 |
| ISPFHDIPIYADKDVF | -6.7458425 | 1.25512291 | 4 | -5.8659011 | 0.61742166 | 3 | 99.1 | 98.3 | -0.8 |
| TFPRPVTVEPMDQLDDEEGLPEKL | -3.0007463 | 0.18199549 | 3 | -2.8916726 | 0.32064147 | 5 | 88.9 | 88.1 | -0.8 |
| IGAIAIGDLVKSTL | -7.3742825 | 2.07631822 | 7 | -6.1697741 | 1.80921761 | 5 | 99.4 | 98.6 | -0.8 |
| ELSGSSSEDSEKVIAGLY | -5.1379603 | 3.70956318 | 2 | -4.7710676 | 1.29645479 | 2 | 97.2 | 96.5 | -0.8 |
| ALKLLDEDEIRGY | -4.7035543 | 1.32598655 | 8 | -4.4184438 | 0.71882613 | 4 | 96.3 | 95.5 | -0.8 |
| GSPKVTKDGVTVA | -6.6629011 | 0.34773486 | 4 | -5.811589 | 1.50815868 | 3 | 99.0 | 98.3 | -0.8 |
| KVFDFELSSQDMTTLL | -2.4367267 | 0.25470296 | 6 | -2.3536272 | 0.2242821 | 4 | 84.4 | 83.6 | -0.8 |
| ILIDEVDKIGRGY | -5.6823832 | 2.21915397 | 3 | -5.1800778 | 0.49979879 | 3 | 98.1 | 97.3 | -0.8 |
| VGGLSPDTPEEKIREY | -6.1911297 | 0.41095663 | 9 | -5.5215935 | 1.31827645 | 3 | 98.6 | 97.9 | -0.8 |
| AITIEAMKSDIDEVALQGIEFW | -3.9060251 | 0.07725527 | 3 | -3.7234514 | 0.37170868 | 5 | 93.7 | 93.0 | -0.8 |
| ASPAHAVDAVKNADGY | -6.3375446 | 0.41597595 | 3 | -5.6047528 | 2.21185617 | 2 | 98.8 | 98.0 | -0.8 |
| ITQAQEQQIELLEIPEHSDIKQIAQPGAAY | -2.7859079 | 0.5226393 | 2 | -2.6846754 | 0.27934298 | 4 | 87.3 | 86.5 | -0.8 |
| RTVSLGAGAKDELHIVEAEAMNY | -2.6615249 | 0.13492436 | 3 | -2.5661113 | 0.44254147 | 7 | 86.4 | 85.6 | -0.8 |
| VIPNTLAVNAAQDSTDLVAKL | -3.6588087 | 0.3910197 | 7 | -3.4967418 | 0.21107166 | 3 | 92.7 | 91.9 | -0.8 |
| YCANETVHGVEFDFIPDVKGAVL | -3.9631351 | 1.72496824 | 4 | -3.7698012 | 1.73020508 | 3 | 94.0 | 93.2 | -0.8 |
| GAQGPKGGSGSGPTIEEVD | -6.263369 | 0.78988643 | 13 | -5.5495019 | 0.91746751 | 11 | 98.7 | 97.9 | -0.8 |
| KQVLGQM(15.9949)VIDEELLGDGHSY | -3.599975 | 0.02090447 | 2 | -3.4418393 | 0.20932536 | 3 | 92.4 | 91.6 | -0.8 |
| LRELISNSSDALDKIRY | -6.6266257 | 1.26471413 | 8 | -5.7572787 | 1.43861575 | 7 | 99.0 | 98.2 | -0.8 |
| ACIGGTNVGEDIRKL | -5.6804037 | 1.37634414 | 5 | -5.1552569 | 1.87683279 | 7 | 98.1 | 97.3 | -0.8 |
| EFDIKVPL | -2.0784878 | 0.3353718 | 5 | -2.0033761 | 1.502418 | 10 | 80.9 | 80.0 | -0.8 |
| LSVEGLVSSVQEGIKF | -5.5232665 | 1.38745315 | 6 | -5.0357196 | 0.66320105 | 6 | 97.9 | 97.0 | -0.8 |
| ACIGGTNVGEDIRKLDY | -5.4521343 | 2.0349646 | 3 | -4.9835752 | 1.92414728 | 4 | 97.8 | 96.9 | -0.8 |
| CLEHGIQPDGQMPSDKTIGGGDDSFNTFF | -2.4732619 | 0.29703038 | 4 | -2.3824406 | 0.41104681 | 2 | 84.7 | 83.9 | -0.8 |
| GQDEMIDVIGVTKGKGY | -6.6967775 | 0.75622108 | 6 | -5.7788417 | 1.03583716 | 6 | 99.0 | 98.2 | -0.8 |
| VINVTLPDEKQNY | -5.689857 | 0.99523667 | 9 | -5.1509688 | 1.1187954 | 7 | 98.1 | 97.3 | -0.8 |
| GFEKPSAIQQ | -6.5433386 | 1.27595183 | 4 | -5.687691 | 1.16466216 | 5 | 98.9 | 98.1 | -0.8 |
| TLVVKWGDEHIPGSPY | -4.461799 | 1.05259072 | 7 | -4.1887194 | 4.542598 | 4 | 95.7 | 94.8 | -0.9 |
| KEREENVKADVF | -8.25258 | 1.70643157 | 3 | -6.3823061 | 0.3455501 | 2 | 99.7 | 98.8 | -0.9 |
| IMPKDIQL | -4.1566886 | 1.05064726 | 14 | -3.9272123 | 1.78852043 | 6 | 94.7 | 93.8 | -0.9 |
| VGNLPHDIDENELKEFF | -6.7843675 | 1.32836241 | 3 | -5.8016935 | 1.16568421 | 3 | 99.1 | 98.2 | -0.9 |
| NVWDTAGQEKF | -6.7429372 | 0.84845595 | 4 | -5.7800916 | 1.48789555 | 4 | 99.1 | 98.2 | -0.9 |
| KVGENADSQIKL | -6.9008706 | 0.92272916 | 12 | -5.8551477 | 2.34266645 | 8 | 99.2 | 98.3 | -0.9 |
| VARGDLGIEIPAEKVF | -6.3396318 | 1.19467208 | 7 | -5.5337241 | 1.21973648 | 8 | 98.8 | 97.9 | -0.9 |
| RITESEEVVSREVSGIKAAY | -3.4905772 | 0.54656653 | 4 | -3.3230381 | 0.50321685 | 3 | 91.8 | 90.9 | -0.9 |
| TKIPDPSTW | -6.5726146 | 0.85780078 | 3 | -5.6453676 | 0.56316655 | 2 | 99.0 | 98.0 | -0.9 |
| QAAAPVLTPAKV | -4.7879615 | 0.58716486 | 5 | -4.4367163 | 0.16387232 | 3 | 96.5 | 95.6 | -0.9 |
| TLGVKQLIVGVN | -7.1226119 | 1.62690657 | 14 | -5.9116426 | 1.36831718 | 9 | 99.3 | 98.4 | -0.9 |
| LYGPPGTGKTL | -4.679429 | 0.67023924 | 7 | -4.3473025 | 0.66071007 | 7 | 96.2 | 95.3 | -0.9 |
| MYQVDTLKDMLLELEEQLAESRRQY | -4.3598773 | 0.3603252 | 3 | -4.0830538 | 0.42340829 | 3 | 95.4 | 94.4 | -0.9 |
| SFGCPEYGQLGHNSDGKF | -2.557481 | 0.38109102 | 6 | -2.45237 | 0.30045544 | 5 | 85.5 | 84.6 | -0.9 |
| TTFPRPVTVEPMDQLDDEEGLPEKL | -1.9126695 | 0.10702691 | 5 | -1.8331674 | 0.31101442 | 7 | 79.0 | 78.1 | -0.9 |
| EGERAMTKDNNLLGKF | -8.080667 | 3.85681846 | 3 | -6.2499731 | 2.79107897 | 4 | 99.6 | 98.7 | -0.9 |
| VRLAPDYDALDVANKIGII | -3.1509383 | 0.23637046 | 6 | -3.009172 | 0.38156102 | 4 | 89.9 | 89.0 | -0.9 |
| VNLGIEPPKGVL | -5.6706567 | 1.30225027 | 5 | -5.0884481 | 0.81905838 | 4 | 98.1 | 97.1 | -0.9 |
| SQFQKQLLEDYGESHF | -5.4460724 | 0.81429087 | 2 | -4.9317548 | 2.12374217 | 2 | 97.8 | 96.8 | -0.9 |

| | | | | | | | | | |
|---|---|---|---|---|---|---|---|---|---|
| GFSHLEALLDDSKELQRF | -2.6620957 | 0.52927119 | 4 | -2.5509691 | 0.45785172 | 2 | 86.4 | 85.4 | -0.9 |
| ADIAQKLQLDSPEDAEF | -4.700588 | 1.20959034 | 3 | -4.3618233 | 0.20625097 | 2 | 96.3 | 95.4 | -0.9 |
| VGLDAAGKTTILY | -1.0949397 | 0.31607169 | 13 | -1.0325266 | 0.31107472 | 7 | 68.1 | 67.2 | -0.9 |
| SGGTTMYPGIADRMQKEITALAPSTM | -2.1876089 | 0.13562209 | 11 | -2.0963506 | 0.26883369 | 8 | 82.0 | 81.0 | -1.0 |
| LLDSFLQPELVKL | -7.7603016 | 2.61859303 | 4 | -6.1240413 | 0.32941803 | 2 | 99.5 | 98.6 | -1.0 |
| KTEGDEEAEEEQEENLEASGDY | -3.3306764 | 0.22676545 | 2 | -3.1701332 | 0.11005994 | 4 | 91.0 | 90.0 | -1.0 |
| LLEEDKPEEPTAHAF | -5.1294135 | 1.63221973 | 7 | -4.685866 | 1.2144638 | 4 | 97.2 | 96.3 | -1.0 |
| LIAGIQHSCQDIGAKSL | -5.7292409 | 0.60039429 | 3 | -5.1106421 | 0.35004042 | 2 | 98.1 | 97.2 | -1.0 |
| EWDVAEARKIW | -5.8585111 | 1.14978403 | 2 | -5.1946761 | 1.49968159 | 5 | 98.3 | 97.3 | -1.0 |
| LAEPVPGIKAEPDESNARYF | -3.0393316 | 0.23795892 | 6 | -2.8976339 | 0.28250318 | 2 | 89.2 | 88.2 | -1.0 |
| CDKSDEDDWSKPLPPSERLEQELF | -6.3467748 | 0.43477693 | 4 | -5.4694891 | 1.66396664 | 5 | 98.8 | 97.8 | -1.0 |
| IANSLATAGDGLIELRKLEAAEDIAY | -4.672336 | 1.10485089 | 9 | -4.3197054 | 1.79506118 | 4 | 96.2 | 95.2 | -1.0 |
| RSCGSSTPDEFPTDIPGTKGNF | -2.9670349 | 0.39986341 | 5 | -2.828986 | 0.41055325 | 2 | 88.7 | 87.7 | -1.0 |
| EPPQIGTQNSLELLEDPKAEVVDEIAAKL | -5.1078292 | 1.34228975 | 2 | -4.6547067 | 1.30597672 | 5 | 97.2 | 96.2 | -1.0 |
| LEKRYNEDLELEDAIHTAIL | -2.5235969 | 0.35166065 | 5 | -2.4119266 | 0.25278344 | 5 | 85.2 | 84.2 | -1.0 |
| SAIRDGETPDPEDPSRKIY | -3.1837705 | 0.15159359 | 8 | -3.0284288 | 0.42838656 | 4 | 90.1 | 89.1 | -1.0 |
| AGRLKPDEGGEVPVLN | -5.6046196 | 0.33219287 | 4 | -5.0045846 | 1.9011067 | 4 | 98.0 | 97.0 | -1.0 |
| SFTVTDEPVYIDLTLSENKEEPVKL | -5.5722616 | 0.72728386 | 3 | -4.9822016 | 1.75585434 | 2 | 97.9 | 96.9 | -1.0 |
| KATAVMPDGQFKDISLSDY | -6.3820844 | 0.43321105 | 4 | -5.4782082 | 1.69086069 | 4 | 98.8 | 97.8 | -1.0 |
| KLDRDPASGTALQEISFW | -4.8083543 | 1.18393132 | 4 | -4.4222466 | 1.56873494 | 3 | 96.6 | 95.5 | -1.0 |
| KFSGDLDDQTCREDLHILF | -4.0526005 | 0.5232551 | 2 | -3.8008708 | 0.61223519 | 2 | 94.3 | 93.3 | -1.0 |
| DIAQVNLKY | -7.8949797 | 3.26442873 | 3 | -6.0944281 | 3.12743654 | 2 | 99.6 | 98.6 | -1.0 |
| NDTSDQIISGGIDNDIKVW | -2.5873557 | 0.50282319 | 3 | -2.4692949 | 0.04327007 | 2 | 85.7 | 84.7 | -1.0 |
| DFEQEMATAASSSSLEKSYELPDGQVITIGNERF | -2.0742476 | 0.51643647 | 3 | -1.9800692 | 0.1113047 | 2 | 80.8 | 79.8 | -1.0 |
| LEQQNKILLAELEQL | -7.2557117 | 2.01682404 | 8 | -5.8613327 | 1.53634806 | 4 | 99.3 | 98.3 | -1.0 |
| ESIQKDSSSTNLESMDTS | -2.5198293 | 0.33184619 | 2 | -2.4043049 | 0.11302695 | 2 | 85.2 | 84.1 | -1.0 |
| AIAQELGSKVPF | -6.2795533 | 1.53398108 | 5 | -5.3923231 | 1.45834153 | 2 | 98.7 | 97.7 | -1.1 |
| QSITFPELSNKGSY | -1.9772202 | 2.43003454 | 4 | -1.8847068 | 4.95194799 | 3 | 79.7 | 78.7 | -1.1 |
| ALSLGDKINL | -6.9589737 | 1.84954579 | 8 | -5.7248644 | 0.8039727 | 2 | 99.2 | 98.1 | -1.1 |
| KNTPWPEAEAIAPQVGNDAVF | -2.9142323 | 0.37913579 | 4 | -2.7710895 | 0.34316775 | 3 | 88.3 | 87.2 | -1.1 |
| TFGADVVSKF | -6.7642118 | 1.34436686 | 2 | -5.6304281 | 1.34444576 | 4 | 99.1 | 98.0 | -1.1 |
| KTVIGELPPASSGSAL | -4.3992859 | 1.77738992 | 4 | -4.0773117 | 3.07839612 | 3 | 95.5 | 94.4 | -1.1 |
| SAYIKEVDEKPASTPW | -4.6089473 | 1.1204321 | 8 | -4.2450027 | 0.89504803 | 2 | 96.1 | 95.0 | -1.1 |
| SFGCPEYGQLGHNSDGKF | -2.5713802 | 0.47060004 | 4 | -2.4492494 | 0.99402117 | 3 | 85.6 | 84.5 | -1.1 |
| DVIAQAQSGTGKTATF | -4.053331 | 1.14071849 | 7 | -3.784977 | 1.47422075 | 8 | 94.3 | 93.2 | -1.1 |
| IQENLELVEKGFSNL | -6.0549238 | 0.28662332 | 2 | -5.2469302 | 2.36263094 | 3 | 98.5 | 97.4 | -1.1 |
| TTNLTEEEEKSKSLAKL | -4.9249931 | 0.02417211 | 2 | -4.485106 | 1.11355469 | 3 | 96.8 | 95.7 | -1.1 |
| AYVQFEDVRDAEDALHNLDRKW | -3.4698409 | 0.22705624 | 2 | -3.2713208 | 0.49711763 | 3 | 91.7 | 90.6 | -1.1 |
| VGGLKGDVAEGDLIEHFSQF | -2.8594147 | 0.33791978 | 10 | -2.7144591 | 0.35190493 | 5 | 87.9 | 86.8 | -1.1 |
| ISQCTPKVDFPQDQLTAL | -1.9969771 | 0.19378559 | 11 | -1.8982382 | 0.38077455 | 4 | 80.0 | 78.8 | -1.1 |
| VAIIGEQELKDGVIKL | -6.6687652 | 2.55777527 | 5 | -5.5478387 | 1.42952554 | 5 | 99.0 | 97.9 | -1.1 |
| ALKEVEEISLLQPQVEESVLNLGKF | -5.9445822 | 0.24070292 | 2 | -5.1544911 | 0.75000956 | 2 | 98.4 | 97.3 | -1.1 |
| TNVPRASVPDGFLSELTQQLAQATGKPPQY | -3.4133517 | 0.00586913 | 2 | -3.214216 | 0.73687207 | 3 | 91.4 | 90.3 | -1.1 |
| LTTTPRPVIVEPLEQLDDEDGLPEKL | -3.0618823 | 0.08150281 | 4 | -2.8925604 | 0.44437538 | 4 | 89.3 | 88.1 | -1.2 |
| LTVKDNQVVQLHPSTVL | -6.4520618 | 0.60042322 | 2 | -5.4059058 | 0.49006908 | 2 | 98.9 | 97.7 | -1.2 |
| KFVEGLPINDF | -5.4659759 | 0.68506344 | 5 | -4.8269744 | 1.8815049 | 2 | 97.8 | 96.6 | -1.2 |
| SGWYDADLSPAGHEEAKRGGQ | -3.3472721 | 0.09370104 | 3 | -3.14588 | 0.65612768 | 3 | 91.1 | 89.8 | -1.2 |
| IFNKQQVPSGESAIL | -4.2211492 | 0.9473656 | 11 | -3.8952996 | 0.76545824 | 5 | 94.9 | 93.7 | -1.2 |
| NAAKVPADTEVVCAPPTAY | -2.5794455 | 0.2173699 | 12 | -2.4418676 | 0.27641175 | 11 | 85.7 | 84.5 | -1.2 |
| KYDGSTIVPGEQGAEYQHF | -3.1632456 | 0.26316922 | 4 | -2.9781098 | 0.20303172 | 2 | 90.0 | 88.7 | -1.2 |
| KAVNPDEAVAIGAAIQGGVL | -4.1314578 | 2.34364317 | 2 | -3.8173253 | 1.21711654 | 3 | 94.6 | 93.4 | -1.2 |
| GNVEKVKF | -6.1449402 | 0.92102226 | 9 | -5.2116903 | 1.80738714 | 5 | 98.6 | 97.4 | -1.2 |
| RIDFYFDENPYFENKVL | -2.5635703 | 0.24336778 | 3 | -2.4236501 | 0.14240601 | 4 | 85.5 | 84.3 | -1.2 |
| SSTSHVPEVDPGSAELQKVL | -5.3052008 | 2.06770649 | 2 | -4.6977446 | 0.95584101 | 2 | 97.5 | 96.3 | -1.2 |
| QAAIQQLAEAQPEATAKNLL | -3.7236517 | 0.02264075 | 2 | -3.4682522 | 0.30266741 | 3 | 93.0 | 91.7 | -1.2 |
| APVNVTTEVKSVEMHHEAL | -1.8538563 | 0.43668677 | 4 | -1.7483283 | 0.51035612 | 5 | 78.3 | 77.1 | -1.3 |
| KEGIPALDNFLDKL | -8.7601908 | 1.53212967 | 4 | -6.0269283 | 1.94308766 | 4 | 99.8 | 98.5 | -1.3 |
| KNLVLVDELDSLSPILF | -5.3985119 | 1.13755532 | 5 | -4.741753 | 1.44186501 | 7 | 97.7 | 96.4 | -1.3 |
| NSLPAERIQEIQKAIELF | -2.5347037 | 0.01536918 | 2 | -2.3906826 | 0.16683062 | 2 | 85.3 | 84.0 | -1.3 |
| AEIAKAELDGTIL | -4.2769311 | 1.37359793 | 5 | -3.9173629 | 0.48080346 | 3 | 95.1 | 93.8 | -1.3 |
| SDPFVEAEKSNLAYDIVQLPTGLTGIKVTY | -7.3231225 | 2.48441498 | 3 | -5.6658381 | 0.75727268 | 2 | 99.4 | 98.1 | -1.3 |
| TVNVEGTKTLPDDQTEVVIY | -4.5531843 | 0.221104 | 2 | -4.1314244 | 0.45270864 | 3 | 95.9 | 94.6 | -1.3 |
| VLNKIDQISIEELDIIY | -5.6076194 | 1.39133283 | 2 | -4.8626987 | 2.12377942 | 6 | 98.0 | 96.7 | -1.3 |
| TDHSDAGADELGEVIKDDIWPNPLQY | -4.5758579 | 0.26045828 | 3 | -4.1467143 | 0.3614825 | 2 | 96.0 | 94.7 | -1.3 |
| EQNDRLTPKIGFPW | -6.3905012 | 1.97174062 | 4 | -5.2862067 | 1.44355667 | 3 | 98.8 | 97.5 | -1.3 |
| TLIIDDPSGNSFVENPHAPQKDDALVITHY | -2.1651717 | 0.07215543 | 3 | -2.0402958 | 0.2204972 | 2 | 81.8 | 80.4 | -1.3 |
| NPNKIFGVTTL | -7.0755773 | 1.05400966 | 8 | -5.5694918 | 2.08008874 | 3 | 99.3 | 97.9 | -1.3 |
| DLTPVDKF | -4.6231335 | 1.23519675 | 2 | -4.180151 | 7.69523775 | 3 | 96.1 | 94.8 | -1.3 |
| APVNVTTEVKSVEMHHEALSEALPGDNVGF | -1.3971454 | 0.25210975 | 3 | -1.302403 | 0.09555039 | 2 | 72.5 | 71.2 | -1.3 |
| SSSSLEKSYELPDGQVITIGNERF | -3.6589513 | 1.46047565 | 5 | -3.3972689 | 0.1696275 | 2 | 92.7 | 91.3 | -1.3 |
| VGNPDGEGEATKGYLDDPTVPRGSTTATF | -3.7418859 | 0.09852255 | 3 | -3.4678214 | 0.28497177 | 2 | 93.0 | 91.7 | -1.3 |
| SVTGAEEVESNGSLQKL | -5.3442525 | 0.32804797 | 2 | -4.68623 | 0.47730986 | 2 | 97.6 | 96.3 | -1.3 |

| | | | | | | | | | |
|---|---|---|---|---|---|---|---|---|---|
| REEGKAEIIPGVLF | -5.2799847 | 1.61069298 | 4 | -4.6439207 | 2.15281648 | 4 | 97.5 | 96.2 | -1.3 |
| LDGKHVVF | -5.9944166 | 1.04979817 | 22 | -5.0744557 | 1.22991435 | 12 | 98.5 | 97.1 | -1.3 |
| GVSGYPTLKIF | -5.8052166 | 0.81662573 | 8 | -4.9685477 | 1.19266771 | 5 | 98.2 | 96.9 | -1.3 |
| APVISAEKAYHEQL | -4.9808279 | 1.16541458 | 27 | -4.4358522 | 0.56109242 | 16 | 96.9 | 95.6 | -1.3 |
| LVKTGTITTF | -5.5039279 | 1.07230574 | 12 | -4.7807455 | 1.16296961 | 10 | 97.8 | 96.5 | -1.4 |
| LFRDGDILGKYVD | -3.9710449 | 1.03867785 | 6 | -3.656082 | 1.34432175 | 4 | 94.0 | 92.7 | -1.4 |
| IVYDIAQVNLKY | -4.4357702 | 0.04989127 | 2 | -4.0289073 | 0.704634 | 2 | 95.6 | 94.2 | -1.4 |
| KILNPEEIEKY | -7.5976842 | 1.48726546 | 7 | -5.7139782 | 1.75977288 | 6 | 99.5 | 98.1 | -1.4 |
| TIEFTEEYPNKPPTVRF | -2.8511409 | 0.24207309 | 4 | -2.6755736 | 0.33059709 | 6 | 87.8 | 86.5 | -1.4 |
| TIEIDFLQKKSIDSNPY | -5.0360543 | 1.23604203 | 2 | -4.4648591 | 0.28294073 | 2 | 97.0 | 95.7 | -1.4 |
| VLQELDNPGAKRIL | -7.4506162 | 3.55698475 | 2 | -5.65556 | 1.4543351 | 4 | 99.4 | 98.1 | -1.4 |
| SGETAKGDYPLEAVRMQHL | -2.8705957 | 0.69712511 | 8 | -2.691165 | 0.92054828 | 4 | 88.0 | 86.6 | -1.4 |
| TTDEDLTEAVHSLGVNDILEIKFF | -3.4247905 | 0.94884543 | 8 | -3.1853162 | 0.38779728 | 3 | 91.5 | 90.1 | -1.4 |
| NENLVKEGVSAAF | -4.8574682 | 1.70043284 | 2 | -4.3312695 | 0.887798 | 2 | 96.7 | 95.3 | -1.4 |
| YELSENDLNFIKQSKDGAGFL | -5.8098415 | 0.86686101 | 8 | -4.9423978 | 1.22642121 | 3 | 98.2 | 96.9 | -1.4 |
| VGGLSPDTPEEKIREYF | -6.2033231 | 1.20589198 | 6 | -5.1451108 | 1.61779803 | 3 | 98.7 | 97.3 | -1.4 |
| ATEFDLEAEEYVPLPKGDVH | -2.6093491 | 0.27011842 | 5 | -2.4455352 | 0.20799053 | 4 | 85.9 | 84.5 | -1.4 |
| EVADLQPQLKIDKAVAF | -5.3193334 | 1.8959596 | 3 | -4.6327915 | 0.67623886 | 3 | 97.6 | 96.1 | -1.4 |
| VGGLSPDTPEEKIREYF | -5.4874406 | 1.10598391 | 6 | -4.7377194 | 0.89600456 | 4 | 97.8 | 96.4 | -1.4 |
| ELKADVVPKTAENF | -7.1517804 | 1.68176578 | 5 | -5.5179299 | 0.72330243 | 2 | 99.3 | 97.9 | -1.4 |
| KLESDYEILERFPGAYL | -2.7984065 | 0.47219962 | 3 | -2.6179421 | 0.47288464 | 2 | 87.4 | 86.0 | -1.4 |
| NILGTNTIMDKMM(15.9949)VAGF | -3.2099288 | 0.99992378 | 3 | -2.9872894 | 0.33505965 | 3 | 90.2 | 88.8 | -1.4 |
| VNHPQVSALLGEEDEEALHYLTRVEVTEFEDIKSG | -2.0783577 | 0.05815439 | 2 | -1.9472883 | 0.1820818 | 4 | 80.9 | 79.4 | -1.4 |
| LATAADDSSVKLW | -3.9660541 | 1.5487383 | 4 | -3.6312673 | 4.51766939 | 5 | 94.0 | 92.5 | -1.5 |
| SVSQAQKDELILEGNDIELVSNSAAL | -3.2483973 | 0.08489915 | 2 | -3.0191875 | 0.34236254 | 4 | 90.5 | 89.0 | -1.5 |
| KEVDEQMLNVQNKNSSYF | -6.4717473 | 2.31808031 | 8 | -5.2422955 | 1.323715 | 3 | 98.9 | 97.4 | -1.5 |
| SATMPSDVLEVTKKF | -7.596422 | 0.27751802 | 2 | -5.6201411 | 1.2210736 | 5 | 99.5 | 98.0 | -1.5 |
| KDDIAQVDYVEPSQNTISL | -2.6241075 | 0.10403126 | 3 | -2.453659 | 0.9518329 | 2 | 86.0 | 84.6 | -1.5 |
| AALADVPELARLLEIDPYLKPY | -3.4455502 | 0.75590553 | 2 | -3.1856304 | 0.79832012 | 3 | 91.6 | 90.1 | -1.5 |
| VGGIKEDTEEHHLRDYFEQY | -3.1306972 | 0.58522232 | 3 | -2.9096045 | 0.45783247 | 3 | 89.8 | 88.3 | -1.5 |
| SFGGKLVTF | -7.4371152 | 2.2700721 | 5 | -5.5581525 | 2.22273033 | 2 | 99.4 | 97.9 | -1.5 |
| AISILQQLEIEFKETQAL | -2.8506718 | 0.35448415 | 3 | -2.6573574 | 0.33605758 | 5 | 87.8 | 86.3 | -1.5 |
| TGSGVVGVKVPQFSF | -5.1763196 | 0.87043712 | 4 | -4.5119804 | 0.85729668 | 3 | 97.3 | 95.8 | -1.5 |
| TLPEVAECFDEITYVELQKEEAQKLLEQY | -5.9830191 | 2.14322042 | 2 | -4.981204 | 1.78086284 | 4 | 98.4 | 96.9 | -1.5 |
| SNQTVDIPENVDITLKGR | -7.5105866 | 1.70124415 | 5 | -5.5674197 | 1.77156711 | 2 | 99.5 | 97.9 | -1.5 |
| SLEKSYELPDGQVITIGNERF | -2.6199584 | 0.43856033 | 6 | -2.4452862 | 0.67563034 | 4 | 86.0 | 84.5 | -1.5 |
| GFPVKVPY | -6.0326997 | 0.97972516 | 2 | -4.9973808 | 1.03002627 | 4 | 98.5 | 97.0 | -1.5 |
| LQTPKIVADKDY | -6.4839571 | 2.21103945 | 2 | -5.1838508 | 1.01481694 | 4 | 98.9 | 97.3 | -1.6 |
| EVLSDSEKRKQY | -5.3172539 | 3.14482274 | 4 | -4.5735556 | 0.20031777 | 2 | 97.6 | 96.0 | -1.6 |
| LYVDKNFINNPLAQADW | -2.7955825 | 0.63819997 | 8 | -2.5970572 | 0.38790374 | 5 | 87.4 | 85.8 | -1.6 |
| VRDKDTDKF | -5.449749 | 0.80511801 | 2 | -4.6488693 | 1.67607066 | 2 | 97.8 | 96.2 | -1.6 |
| FKDDANNDPQWSEEQLIAAKF | -4.1052856 | 1.34060769 | 3 | -3.7123558 | 0.32045964 | 3 | 94.5 | 92.9 | -1.6 |
| KEIEYEVVRDAY | 0.1040904 | 0.333358 | 2 | 0.19663634 | 0.20468534 | 6 | 48.2 | 46.6 | -1.6 |
| KAIVAGDQNVEY | -4.6339849 | 0.1828732 | 5 | -4.1053289 | 1.38936072 | 3 | 96.1 | 94.5 | -1.6 |
| ISKIISSDRDLL | -7.9189786 | 1.78396852 | 8 | -5.5864911 | 0.82935128 | 5 | 99.6 | 98.0 | -1.6 |
| AYVEFENPDEAEKAL | -4.5218403 | 1.35051282 | 3 | -4.0196836 | 0.52366689 | 3 | 95.8 | 94.2 | -1.6 |
| LLPGELAKH | -6.2770647 | 0.97535526 | 7 | -5.0575135 | 0.94910999 | 5 | 98.7 | 97.1 | -1.6 |
| KLVICPDEGFY | -6.4113595 | 1.46540142 | 8 | -5.1142971 | 1.58865524 | 4 | 98.8 | 97.2 | -1.6 |
| AFVTFDDHDTVDKIVVQKY | -5.5456927 | 1.24144404 | 2 | -4.6842609 | 0.16976048 | 2 | 97.9 | 96.3 | -1.6 |
| KMLVDDIGDVTITNDGATIL | -3.1583562 | 0.75731516 | 8 | -2.9115158 | 1.31520997 | 7 | 89.9 | 88.3 | -1.7 |
| VALDFEQEMATAASSSSLEKSYELPDGQVITIGNE | -2.2165689 | 0.06489989 | 7 | -2.0564233 | 0.35238007 | 5 | 82.3 | 80.6 | -1.7 |
| VDIEKLPW | -5.6518521 | 1.00903339 | 7 | -4.7309582 | 1.19085139 | 8 | 98.0 | 96.4 | -1.7 |
| RAIKQVYEEEYGSSLEDDVVGDTSGY | -3.0854259 | 0.09572159 | 3 | -2.842938 | 0.32419621 | 2 | 89.5 | 87.8 | -1.7 |
| GVTKIGVTVL | -7.7317301 | 1.81022481 | 10 | -5.486441 | 1.34738616 | 5 | 99.5 | 97.8 | -1.7 |
| KEQISDIDDAVRKL | -4.9030268 | 1.95983573 | 2 | -4.2610516 | 1.06361385 | 3 | 96.8 | 95.0 | -1.7 |
| LIIDITTTFPKDPVY | -7.1397173 | 2.50770784 | 3 | -5.3279574 | 1.05289586 | 5 | 99.3 | 97.6 | -1.7 |
| VVINQKGIDPF | -6.6781465 | 1.6964415 | 14 | -5.1711583 | 1.31198755 | 9 | 99.0 | 97.3 | -1.7 |
| ALAAVAGGAPSVGIKAAN | -4.5909715 | 1.62612843 | 2 | -4.0422099 | 0.55590765 | 3 | 96.0 | 94.3 | -1.7 |
| TASAGIQVVGDDLTVTNPKRIA | -4.1697929 | 0.25542404 | 10 | -3.7308134 | 0.32812589 | 5 | 94.7 | 93.0 | -1.7 |
| AAMEELVDEGLVKAIGISNF | -2.5275121 | 0.33789758 | 9 | -2.3357967 | 0.29371857 | 5 | 85.2 | 83.5 | -1.8 |
| LEKEELPRAVGTQTL | -4.3452533 | 0.83341338 | 2 | -3.8553767 | 1.45425605 | 3 | 95.3 | 93.5 | -1.8 |
| LSVEGLVSSVQEGIKF | -6.070382 | 0.59809147 | 2 | -4.89924 | 1.1451766 | 5 | 98.5 | 96.8 | -1.8 |
| VTAKDAERAIN | -5.4141844 | 0.91039682 | 2 | -4.5575751 | 0.69053824 | 3 | 97.7 | 95.9 | -1.8 |
| VLGDQQHCDEAKAVDIPHMDIEAL | -2.1320512 | 0.40150933 | 2 | -1.9668003 | 0.14765624 | 2 | 81.4 | 79.6 | -1.8 |
| LTNSLELEHIEVKF | -5.3247072 | 0.76962101 | 4 | -4.5001334 | 0.60030081 | 5 | 97.6 | 95.8 | -1.8 |
| IGGLPNYLNDDQVKELLTSF | -3.3388737 | 0.25265642 | 8 | -3.0456512 | 0.27645447 | 2 | 91.0 | 89.2 | -1.8 |
| DVGGQDKIRPL | -6.3347655 | 1.15691628 | 4 | -4.9988899 | 0.61771077 | 5 | 98.8 | 97.0 | -1.8 |
| SIYPHGSTDKLPY | -3.0875858 | 1.08505957 | 12 | -2.828071 | 1.03794282 | 3 | 89.5 | 87.7 | -1.8 |
| VHGGLDSNGKPADAVY | -5.0072008 | 1.32235179 | 7 | -4.2896887 | 2.17433239 | 4 | 97.0 | 95.1 | -1.8 |
| VTGIKVVDLLAPY | 1.27450705 | 0.33615998 | 9 | 1.40653522 | 0.31317352 | 4 | 29.2 | 27.4 | -1.9 |
| LEKRYNEDLELEDAIHTAIL | -2.9202174 | 0.13314946 | 4 | -2.6753628 | 0.06494164 | 3 | 88.3 | 86.5 | -1.9 |
| GYDHLILNVEWAKPSTN | -5.547161 | 0.76118562 | 3 | -4.5994286 | 0.72961954 | 3 | 97.9 | 96.0 | -1.9 |

| | | | | | | | | | |
|---|---|---|---|---|---|---|---|---|---|
| VGNLPTDITEEDFKRLFERYGEPSEVF | -3.0705403 | 0.04935878 | 2 | -2.8063932 | 0.20725189 | 3 | 89.4 | 87.5 | -1.9 |
| DVGGQDKIRPLW | -4.6351646 | 0.67089094 | 6 | -4.0333717 | 0.4842949 | 4 | 96.1 | 94.2 | -1.9 |
| NILGTNTIMDKMMVAGF | -3.3836263 | 0.17974656 | 8 | -3.0712142 | 0.50327178 | 4 | 91.3 | 89.4 | -1.9 |
| NAIVIKETKDW | -4.7446757 | 1.15284399 | 6 | -4.1052587 | 0.9719076 | 3 | 96.4 | 94.5 | -1.9 |
| LTMEINPKVPVNLL | -5.0187334 | 0.2769105 | 3 | -4.2810709 | 0.67653537 | 2 | 97.0 | 95.1 | -1.9 |
| GAGAKDELHIVEAEAMNY | -2.052332 | 0.67871556 | 4 | -1.8831407 | 0.53498973 | 4 | 80.6 | 78.7 | -1.9 |
| IQATCATSGDGLYEGLDWLSNQLRNQK | -2.1495298 | 0.24796075 | 9 | -1.9704976 | 0.40218863 | 4 | 81.6 | 79.7 | -1.9 |
| ADDQIAQSLCGEDLIIKGISVH | -3.9140945 | 0.99163603 | 2 | -3.4921172 | 0.23741666 | 2 | 93.8 | 91.8 | -1.9 |
| KVADIGLAAW | -6.6966951 | 1.74439015 | 9 | -5.053623 | 2.43438087 | 5 | 99.0 | 97.1 | -2.0 |
| NQVNVVLKDAEGILEDLQSY | -2.6048379 | 0.01365254 | 2 | -2.3830457 | 0.48183355 | 2 | 85.9 | 83.9 | -2.0 |
| HAIIVDGHSVEELCKAF | -2.5433488 | 0.33049245 | 6 | -2.3268087 | 0.10547915 | 4 | 85.4 | 83.4 | -2.0 |
| KEFLPEGQDIGAF | -5.277639 | 1.22891133 | 4 | -4.4060277 | 1.44691318 | 3 | 97.5 | 95.5 | -2.0 |
| DLDFKSPDDPSRY | -1.8006561 | 0.10285597 | 5 | -1.6394882 | 0.53407209 | 10 | 77.7 | 75.7 | -2.0 |
| RIQAGEIGEMKDGVPEGAQL | -2.4383908 | 0.02659629 | 2 | -2.2297271 | 0.65981044 | 2 | 84.4 | 82.4 | -2.0 |
| FQFQEEGKEGENRAVIHY | -2.8710604 | 0.31557495 | 4 | -2.6141417 | 0.27345059 | 3 | 88.0 | 86.0 | -2.0 |
| VALDFEQEM(15.9949)ATAASSSSLEKSYELPDG( | -2.3584929 | 0.22412268 | 8 | -2.1550407 | 0.20652451 | 4 | 83.7 | 81.7 | -2.0 |
| GMKTIGYDPIISPEVSASF | -3.4571855 | 0.48577342 | 3 | -3.1096674 | 0.32147601 | 4 | 91.7 | 89.6 | -2.0 |
| DFWKDIVAAIQHNY | -7.9258819 | 2.88485761 | 2 | -5.3164593 | 0.41860968 | 2 | 99.6 | 97.6 | -2.0 |
| SQPKMDELQLF | -5.818538 | 2.57315149 | 3 | -4.6659742 | 0.98756601 | 4 | 98.3 | 96.2 | -2.0 |
| VALVPQEEELDDQKIQVTPPGF | -2.8231585 | 0.35291276 | 3 | -2.5677815 | 0.63907458 | 3 | 87.6 | 85.6 | -2.1 |
| KLAVEALSSLDGDLAGRYY | -6.6542899 | 1.08495681 | 10 | -4.9963669 | 0.67987404 | 6 | 99.0 | 97.0 | -2.1 |
| YLTRVEVTEFEDIKSGY | -2.6745445 | 0.0754679 | 3 | -2.4360335 | 0.19280831 | 5 | 86.5 | 84.4 | -2.1 |
| QEALDAAGDKLVVVDFSATW | -3.2842709 | 1.94446954 | 7 | -2.9629803 | 0.85991556 | 11 | 90.7 | 88.6 | -2.1 |
| VLAADKVASVASTLETTFETISTL | -2.5323323 | 0.33673864 | 5 | -2.3076984 | 0.5361595 | 3 | 85.3 | 83.2 | -2.1 |
| VQSEIFPLETPAFAIKEQGF | -2.6850941 | 0.51486943 | 8 | -2.4425516 | 0.64788367 | 3 | 86.5 | 84.5 | -2.1 |
| SVAKGSDEPPVFLEIHY | -3.7701248 | 0.5065028 | 6 | -3.3530857 | 0.477203 | 5 | 93.2 | 91.1 | -2.1 |
| QAQIQEQHVQIDVDVSKPDLTAAL | -3.1607418 | 0.06923028 | 2 | -2.8545493 | 2.50774042 | 3 | 89.9 | 87.9 | -2.1 |
| VASVHQDLSDDDIKSVF | -5.0173276 | 1.21406268 | 4 | -4.2042943 | 1.12529775 | 2 | 97.0 | 94.9 | -2.2 |
| VRNLATTVTEEILEKSFSEF | -2.9544857 | 0.26988004 | 5 | -2.6698467 | 0.08719379 | 4 | 88.6 | 86.4 | -2.2 |
| NTGVEAGETACKL | -4.0707495 | 3.31510625 | 2 | -3.5689534 | 2.28958561 | 4 | 94.4 | 92.2 | -2.2 |
| ALKEVEEISLLQPQVEESVL | -3.1890752 | 0.18456973 | 5 | -2.8689301 | 1.55237231 | 6 | 90.1 | 88.0 | -2.2 |
| ISDKDASIVGFFDDSFSEAHSEFL | -3.4028446 | 0.34589777 | 10 | -3.043591 | 0.32797864 | 2 | 91.4 | 89.2 | -2.2 |
| MQDPMEIFVDDETKLTLHGLQQYY | -2.1587674 | 0.08223138 | 3 | -1.9551379 | 0.06573105 | 2 | 81.7 | 79.5 | -2.2 |
| EIQDIYENSWTKLTERFF | -4.468335 | 0.26655245 | 3 | -3.8396743 | 0.38059918 | 2 | 95.7 | 93.5 | -2.2 |
| DLKATQAL | -4.3743571 | 1.67702231 | 3 | -3.7747382 | 2.9990616 | 2 | 95.4 | 93.2 | -2.2 |
| CVECSEPLCETCVEAHQRVKY | -2.5901186 | 0.53721929 | 5 | -2.3424975 | 0.64100872 | 2 | 85.8 | 83.5 | -2.2 |
| AALAALEKLFPDTPL | -5.3405929 | 0.89807426 | 6 | -4.3614556 | 1.03139788 | 4 | 97.6 | 95.4 | -2.2 |
| IIDPNGVIKHL | -6.2541902 | 3.19247144 | 2 | -4.7706421 | 0.60309714 | 6 | 98.7 | 96.5 | -2.2 |
| NMDQFTPVKIEGYEDQVLITEHGDLGNSRF | -2.2702172 | 0.62194723 | 2 | -2.052228 | 0.24451537 | 2 | 82.8 | 80.6 | -2.3 |
| VTTVTEIGKDVIGL | -8.1362793 | 1.64909355 | 3 | -5.2009357 | 0.90181728 | 4 | 99.6 | 97.4 | -2.3 |
| VAVEEDAES(79.9663)EDEEEEDVKLL | -3.1072276 | 1.29697478 | 11 | -2.7819422 | 0.13597891 | 12 | 89.6 | 87.3 | -2.3 |
| TEGAELVDSVLDVVRKEAESCDCLQGF | -2.6644412 | 0.18242997 | 9 | -2.3999563 | 0.20234252 | 7 | 86.4 | 84.1 | -2.3 |
| VSGAGDIKLTKDGNVL | -6.9038516 | 3.97800591 | 3 | -4.9389762 | 1.10412534 | 3 | 99.2 | 96.8 | -2.3 |
| VSASQDGKLIIWDSY | -4.8721787 | 2.27613068 | 2 | -4.0661487 | 0.72378123 | 2 | 96.7 | 94.4 | -2.3 |
| TAAEVVVPRDQTPDENDQVIVKIIGHFY | -2.7923916 | 0.30546256 | 4 | -2.5068878 | 0.35757861 | 5 | 87.4 | 85.0 | -2.3 |
| DVSNADRLGFSEVELVQMVVDGVKL | -3.2250256 | 0.03047039 | 3 | -2.8718327 | 0.36489184 | 6 | 90.3 | 88.0 | -2.4 |
| CLEHGIQPDGQM(15.9949)PSDKTIGGGDDSFN | -2.3624059 | 0.44182716 | 4 | -2.1251984 | 0.40433345 | 5 | 83.7 | 81.4 | -2.4 |
| LFGGLANDSEDPKNNIPRYLNDLY | -2.7187749 | 0.4560158 | 3 | -2.4405226 | 0.59794857 | 2 | 86.8 | 84.4 | -2.4 |
| GVVTAIDLLNFVAAQERDQK | -2.6130558 | 0.26532612 | 9 | -2.3463762 | 0.09626169 | 4 | 86.0 | 83.6 | -2.4 |
| TLVCSSAPGPLELDLTGDLESFKKQSF | -4.5625005 | 1.98874706 | 4 | -3.8585792 | 1.96994233 | 2 | 95.9 | 93.6 | -2.4 |
| SNTGEDWYVLVGVAKDLILNPRSVAGGF | -1.7818675 | 0.16775279 | 6 | -1.5907981 | 0.35280641 | 3 | 77.5 | 75.1 | -2.4 |
| DLERKVESLQEEIAFL | -6.3235491 | 0.6517235 | 5 | -4.7275714 | 1.96107424 | 3 | 98.8 | 96.4 | -2.4 |
| GIVDYMIEQSGPPSKEIL | -2.5551941 | 0.32508618 | 6 | -2.2927629 | 0.24017833 | 2 | 85.5 | 83.1 | -2.4 |
| IMTVGCVAGDEESYEVFKDLFDPIIEDRHGGY | -1.7676425 | 0.38251972 | 2 | -1.5745674 | 0.03563916 | 2 | 77.3 | 74.9 | -2.4 |
| DLSDKSINPLGGFVHY | -5.0821527 | 1.09818687 | 8 | -4.1575553 | 0.77764028 | 3 | 97.1 | 94.7 | -2.4 |
| GVDVTTKEIVLADVIDNDSW | -2.5228236 | 0.2129659 | 4 | -2.2586637 | 0.01347383 | 2 | 85.2 | 82.7 | -2.5 |
| AISILQQLEIEFKETQAL | -2.7025909 | 0.1333188 | 2 | -2.4142361 | 0.24146195 | 5 | 86.7 | 84.2 | -2.5 |
| ANLGVAEKTQLL | -5.8309265 | 3.76290542 | 3 | -4.4975645 | 1.31478768 | 4 | 98.3 | 95.8 | -2.5 |
| RFDDTNPEKEEAKFF | -5.6726642 | 2.48838893 | 2 | -4.4278202 | 0.71859209 | 3 | 98.1 | 95.6 | -2.5 |
| QTGSTEPVEVPPELCHGIQGKF | -2.6769426 | 0.26598588 | 6 | -2.3859571 | 0.47945148 | 6 | 86.5 | 83.9 | -2.5 |
| VLQELDNPGAKRILELDQF | -2.6370218 | 0.46420382 | 5 | -2.3438376 | 0.44293879 | 6 | 86.2 | 83.5 | -2.6 |
| LIVDRDEAWVLETIGKYW | -3.0446157 | 0.60292622 | 3 | -2.6898357 | 0.21202525 | 3 | 89.2 | 86.6 | -2.6 |
| FVGNLPADITEDEFKRLF | -3.5440979 | 0.53923757 | 5 | -3.0906655 | 0.48289505 | 4 | 92.1 | 89.5 | -2.6 |
| EFKGEDLTEEEDGGIIRR | -6.5181637 | 1.09014118 | 3 | -4.6993385 | 1.02941685 | 2 | 98.9 | 96.3 | -2.6 |
| QPPTVVPGGDLAKVQ | -5.0857197 | 1.67963786 | 7 | -4.1036805 | 0.1752694 | 6 | 97.1 | 94.5 | -2.6 |
| GLVPAKGLATSVL | -3.8719396 | 1.61760892 | 4 | -3.3316717 | 0.57513022 | 3 | 93.6 | 91.0 | -2.6 |
| SGRDLNCVPEIADTLGAVAKQGFDFL | -2.1520355 | 0.00131488 | 2 | -1.9060127 | 0.46018597 | 2 | 81.6 | 78.9 | -2.7 |
| CFITYTDEEPVKKLL | -4.7976343 | 1.68130691 | 4 | -3.9266672 | 0.83142463 | 2 | 96.5 | 93.8 | -2.7 |
| SLEKSYELPDGQVITIGNERF | -2.6633947 | 0.39625762 | 10 | -2.3508555 | 0.45696938 | 5 | 86.4 | 83.6 | -2.8 |
| AQRALDEATKY | -3.8702607 | 0.21042465 | 2 | -3.3029435 | 1.74678993 | 4 | 93.6 | 90.8 | -2.8 |
| IFAGKQLEDGRTL | -5.3239941 | 1.32967957 | 5 | -4.1730172 | 0.48165371 | 4 | 97.6 | 94.7 | -2.8 |
| GFVEFEDPRDADDAVYELDGKEL | -4.2576359 | 0.91166633 | 2 | -3.5611537 | 0.00072248 | 2 | 95.0 | 92.2 | -2.8 |

| | | | | | | | | | |
|---|---|---|---|---|---|---|---|---|---|
| QNSESGEKNEGSESAPEGQAQQRRPY | -3.2595336 | 0.11890016 | 4 | -2.8293396 | 0.775564 | 4 | 90.5 | 87.7 | -2.9 |
| GGQKCSVIRDSLLQDGEF | -2.7925362 | 0.42716863 | 10 | -2.4458181 | 0.46288126 | 6 | 87.4 | 84.5 | -2.9 |
| KLDDPSCPRPECY | -2.8083813 | 0.38011056 | 4 | -2.4583234 | 0.68328602 | 5 | 87.5 | 84.6 | -2.9 |
| AAPSTDAPDKGYVVPNVDLPPLCSSRF | -1.9874219 | 0.14655058 | 2 | -1.7389612 | 1.1651653 | 2 | 79.9 | 76.9 | -2.9 |
| ALEVAAKTLPF | -4.7911148 | 0.86807109 | 4 | -3.8695411 | 1.17265816 | 6 | 96.5 | 93.6 | -2.9 |
| AFVTFDDHDTVDKIVVQKY | -5.6866106 | 1.99279873 | 5 | -4.3026053 | 1.5597867 | 5 | 98.1 | 95.2 | -2.9 |
| SGGTTMYPGIADRMQKEITALAPSTM(15.9949) | -2.3070041 | 0.00333646 | 3 | -2.0214296 | 0.12714797 | 6 | 83.2 | 80.2 | -3.0 |
| VLQELDNPGAKRILELDQF | -2.5726806 | 0.51106627 | 4 | -2.2521495 | 0.45668609 | 6 | 85.6 | 82.7 | -3.0 |
| IGGLSFETTEESLRNYYEQWGKLTDCVVM | -2.8377159 | 0.20533145 | 4 | -2.4727091 | 0.51374608 | 4 | 87.7 | 84.7 | -3.0 |
| ITFLEDLKSF | -2.0490768 | 0.60181547 | 9 | -1.7867104 | 0.47083971 | 5 | 80.5 | 77.5 | -3.0 |
| AANPNQNKNVALL | -2.3975095 | 2.25338887 | 3 | -2.0938379 | 5.2697401 | 2 | 84.0 | 81.0 | -3.0 |
| DEVAAQGEVVRKL | -3.9434202 | 1.2444382 | 2 | -3.3145295 | 2.01519325 | 3 | 93.9 | 90.9 | -3.0 |
| GMKTIGYDPIISPEVSASF | -3.5561038 | 0.65874194 | 11 | -3.0355109 | 0.45049303 | 6 | 92.2 | 89.1 | -3.0 |
| KEIVVIHQDPEAL | -4.4391493 | 0.93649383 | 2 | -3.6346124 | 3.49874391 | 3 | 95.6 | 92.5 | -3.0 |
| VISDRKELEEDFIKSEL | -7.1559223 | 0.84690156 | 2 | -4.670023 | 1.99899359 | 4 | 99.3 | 96.2 | -3.1 |
| VALDFEQEMATAASSSSLEKSYELPDGQVITIGNE | -2.1099014 | 0.19696814 | 3 | -1.8346749 | 0.82939578 | 3 | 81.2 | 78.1 | -3.1 |
| IKDYPVVSIEDPFDQDDW | -2.2062371 | 0.42395632 | 5 | -1.9167969 | 0.48790213 | 5 | 82.2 | 79.1 | -3.1 |
| SMSPVDDTFISGSLDKTIRLW | -2.8945569 | 0.44460627 | 3 | -2.5033597 | 0.34104812 | 3 | 88.1 | 85.0 | -3.1 |
| IGGLSFETTDDSLREHFEKW | -2.3950446 | 0.11837405 | 3 | -2.0778109 | 0.41934564 | 6 | 84.0 | 80.8 | -3.2 |
| MAVLNEQVKEAEGSSAEY | -3.2081823 | 0.87619443 | 3 | -2.7440063 | 0.38963707 | 2 | 90.2 | 87.0 | -3.2 |
| SRILDKSLPADILYEDQQCL | -2.8998885 | 0.20288318 | 2 | -2.4912344 | 0.88590263 | 2 | 88.2 | 84.9 | -3.3 |
| SQQAASVVKQEGGDNDLIERIQVDAY | -3.1753854 | 0.33882713 | 4 | -2.7101204 | 1.95649492 | 4 | 90.0 | 86.7 | -3.3 |
| KFDVFEDFISPTTAAQTLLF | -4.6409675 | 0.27128683 | 2 | -3.6997352 | 0.22159102 | 2 | 96.1 | 92.9 | -3.3 |
| KLLLQGEADQSLL | -7.4151953 | 2.16221003 | 6 | -4.6184279 | 0.36514603 | 2 | 99.4 | 96.1 | -3.3 |
| VKLFPNSLDQTDMHGDSEY | -1.6982559 | 1.16430561 | 8 | -1.4401458 | 0.803595 | 5 | 76.4 | 73.1 | -3.4 |
| AFGPDATGPNILVDDTLPSEVDKALL | -5.4598422 | 2.44875855 | 4 | -4.0742481 | 0.16958499 | 2 | 97.8 | 94.4 | -3.4 |
| MIQGGDFTRGDGTGGKSIY | -2.7590166 | 0.3030796 | 6 | -2.3643642 | 0.44328284 | 6 | 87.1 | 83.7 | -3.4 |
| SKQILEILNSTGVEYETF | -2.4895379 | 0.45046621 | 10 | -2.1337837 | 0.20266927 | 5 | 84.9 | 81.4 | -3.4 |
| IVAAGVGEFEAGISKNGQTREH | -3.3529885 | 0.42386224 | 3 | -2.8253829 | 0.38747739 | 2 | 91.1 | 87.6 | -3.4 |
| LVQLPLDSENSINTEEVINAIAPEKDVDGLTSINAG | -2.3048404 | 0.01703716 | 2 | -1.9709425 | 0.57053848 | 2 | 83.2 | 79.7 | -3.5 |
| IGDSGVGKSNLL | 1.28107827 | 0.39113878 | 2 | 1.53620677 | 0.40945127 | 3 | 29.2 | 25.6 | -3.5 |
| TAIEAPKGEFGVY | 0.96992447 | 0.44735767 | 7 | 1.20431508 | 0.25741045 | 6 | 33.8 | 30.3 | -3.5 |
| SASPLEEATLSELKTVL | -4.6188019 | 0.67892777 | 2 | -3.6214587 | 0.43560312 | 4 | 96.1 | 92.5 | -3.6 |
| EIIDDHRAGKIVVNL | -6.6679847 | 0.35096368 | 2 | -4.3642245 | 1.87600743 | 4 | 99.0 | 95.4 | -3.7 |
| AVELFEGLKAF | -3.9669297 | 0.27276368 | 4 | -3.217025 | 0.758012 | 2 | 94.0 | 90.3 | -3.7 |
| VASLAEPDFVVTDFAKF | 1.71952016 | 0.19326115 | 3 | 2.03800304 | 0.04174485 | 3 | 23.3 | 19.6 | -3.7 |
| ENLFPGLKEAF | -6.935015 | 2.82156168 | 2 | -4.3996701 | 1.41357274 | 4 | 99.2 | 95.5 | -3.7 |
| GFGSGDDPYSSAEPHVSGVKR | -2.9299634 | 0.06751378 | 3 | -2.4669078 | 0.38982378 | 2 | 88.4 | 84.7 | -3.7 |
| TIKDPAVGFLETISPGY | -5.4456928 | 0.76442711 | 3 | -3.9673782 | 0.60690178 | 3 | 97.8 | 94.0 | -3.8 |
| QIATVTEKGVEIEGPLSTETNWDIAHM(15.9949) | -2.528672 | 0.50176392 | 3 | -2.1345137 | 0.19357448 | 2 | 85.2 | 81.5 | -3.8 |
| EVADLQPQLKIDKAVAF | -4.8186836 | 0.37966118 | 3 | -3.6872314 | 0.42800842 | 2 | 96.6 | 92.8 | -3.8 |
| AALGDPKAPGLGAF | -5.8579217 | 1.73709559 | 9 | -4.1085891 | 2.91689824 | 5 | 98.3 | 94.5 | -3.8 |
| VEKNPTIVNFPITNVDLREY | 2.26864092 | 0.14054909 | 4 | 2.70473134 | 1.0654276 | 2 | 17.2 | 13.3 | -3.9 |
| LAVQVQNEEGKCEVTEVSKL | -8.3312677 | 2.31155551 | 2 | -4.4669809 | 0.47533132 | 2 | 99.7 | 95.7 | -4.0 |
| VTEKVLAAVY | -2.8734373 | 0.21627531 | 12 | -2.3877712 | 0.74461104 | 13 | 88.0 | 84.0 | -4.0 |
| KQVLGQMVIDEELLGDGHSY | -3.3041284 | 0.46168218 | 7 | -2.7121515 | 0.33930596 | 4 | 90.8 | 86.8 | -4.0 |
| HTELERLPAKDIQTNVY | -3.9375388 | 1.2752126 | 5 | -3.1368328 | 1.23511716 | 2 | 93.9 | 89.8 | -4.1 |
| LQDKDVIAINQDPLGKQGY | -7.7676086 | 1.03733611 | 2 | -4.3585613 | 0.79719716 | 2 | 99.5 | 95.4 | -4.2 |
| RAEVDLVVQDLKQAVAQLEDQA | -3.2383451 | 0.16476726 | 3 | -2.642504 | 0.96227949 | 2 | 90.4 | 86.2 | -4.2 |
| YLDFLAPAKATL | -4.8453509 | 1.44529811 | 2 | -3.6066046 | 2.74497406 | 2 | 96.6 | 92.4 | -4.2 |
| ALEWPSLTAQWLPDVTRPEGKDF | -2.5498265 | 0.14038751 | 2 | -2.1075782 | 0.61123749 | 3 | 85.4 | 81.2 | -4.2 |
| VVGESGLGKSTL | 0.01737614 | 0.02822727 | 4 | 0.27217176 | 0.21629316 | 9 | 49.7 | 45.3 | -4.4 |
| AALETLDNGKPY | -2.5368743 | 0.27093223 | 3 | -2.0784677 | 0.24383624 | 3 | 85.3 | 80.9 | -4.4 |
| TVEAAEVVFPNGKSHTFPELAY | -2.502512 | 0.37632256 | 4 | -2.0415688 | 0.81767288 | 3 | 85.0 | 80.5 | -4.5 |
| QYLKQVLGQMVIDEELLGDGHSY | -1.7461707 | 0.87006072 | 3 | -1.3927157 | 0.47992018 | 3 | 77.0 | 72.4 | -4.6 |
| KGLVYETSVLDPDEGIRF | -2.5953408 | 1.04190978 | 6 | -2.1065356 | 1.7662991 | 2 | 85.8 | 81.2 | -4.6 |
| GDPVVQSDMKHWPF | -5.7774613 | 2.58110526 | 5 | -3.8381405 | 2.18934624 | 2 | 98.2 | 93.5 | -4.7 |
| TDKVVIGM(15.9949)DVAASEFF | -5.8741536 | 1.61709231 | 4 | -3.845826 | 2.50917798 | 4 | 98.3 | 93.5 | -4.8 |
| VKLFPNSLDQTDMHGDSEY | -1.7147721 | 1.1908556 | 8 | -1.3491602 | 0.70422058 | 6 | 76.6 | 71.8 | -4.8 |
| YTEPVIAGLDPKTF | -4.5527657 | 0.29230083 | 2 | -3.3441186 | 2.677165 | 3 | 95.9 | 91.0 | -4.9 |
| VGETGSGKTTQIPQWCVEY | -3.6303972 | 0.55262219 | 2 | -2.8085993 | 0.36027035 | 2 | 92.5 | 87.5 | -5.0 |
| IAKADTLTPEECQQF | -1.6323864 | 1.004748 | 5 | -1.2576877 | 0.10751888 | 5 | 75.6 | 70.5 | -5.1 |
| AYEPVWAIGTGKTATPQ | -5.3431812 | 2.20902236 | 3 | -3.615871 | 4.68192589 | 9 | 97.6 | 92.5 | -5.1 |
| QSVDDAAIVIKNTKEPPLSL | -5.921984 | 2.20323841 | 3 | -3.781042 | 0.56195481 | 2 | 98.4 | 93.2 | -5.2 |
| KNLVLVDELDSLSPILF | -6.1186733 | 0.27280837 | 2 | -3.8185735 | 0.27930987 | 3 | 98.6 | 93.4 | -5.2 |
| SLVDPEKAKAL | -4.3544013 | 2.39935426 | 3 | -3.1841517 | 0.33306123 | 2 | 95.3 | 90.1 | -5.3 |
| TADMLGSAELAEEVNLNGSGKLL | -0.2434363 | 3.25356759 | 2 | 0.06190987 | 5.87169321 | 2 | 54.2 | 48.9 | -5.3 |
| TNEVLTKTY | -1.2732112 | 0.05679951 | 3 | -0.9135836 | 0.00903175 | 2 | 70.7 | 65.3 | -5.4 |
| TNTCPSDKEVEIAY | -4.2129335 | 0.86386658 | 2 | -3.0837824 | 1.85393772 | 2 | 94.9 | 89.4 | -5.4 |
| NNNEDFQVGQAKVVL | 1.42817693 | 0.09853992 | 9 | 1.86354961 | 0.40568352 | 11 | 27.1 | 21.6 | -5.5 |
| LVAEKVTVITKHNDDEQY | -2.4208481 | 0.16332415 | 2 | -1.880229 | 1.46655914 | 2 | 84.3 | 78.6 | -5.6 |
| GVVTAIDLLNFVAAQERDQK | -2.3656746 | 0.29785178 | 5 | -1.8363753 | 1.51468585 | 3 | 83.8 | 78.1 | -5.6 |

| | | | | | | | | | |
|---|---|---|---|---|---|---|---|---|---|
| FVKGLSEDTTEETLKESF | -4.2317954 | 0.62674302 | 3 | -3.0565594 | 3.89446053 | 2 | 94.9 | 89.3 | -5.7 |
| DLDFKSPDDPSRYISPDQLADLY | -1.6428099 | 0.21738803 | 7 | -1.2231566 | 0.1675705 | 2 | 75.7 | 70.0 | -5.7 |
| LLGLDNAGKTTLL | 1.91665806 | 0.06722019 | 3 | 2.48896397 | 0.30827055 | 5 | 20.9 | 15.1 | -5.8 |
| QVEYTPFEKGLHVVEVTYDDVPIPNSPF | -2.1869721 | 0.01755861 | 2 | -1.6737157 | 0.1333763 | 2 | 82.0 | 76.1 | -5.9 |
| AALEKLFPDTPL | -4.2449446 | 4.37116366 | 7 | -3.0277003 | 4.19541117 | 5 | 95.0 | 89.1 | -5.9 |
| QFLNAKCESAFL | -4.9856344 | 1.85383982 | 4 | -3.3281108 | 5.93630079 | 3 | 96.9 | 90.9 | -6.0 |
| DLDFKSPDDPSRY | -1.8356024 | 0.1586121 | 9 | -1.3638752 | 0.14554647 | 12 | 78.1 | 72.0 | -6.1 |
| IIFDEPPVDKTY | -4.3958433 | 0.42660643 | 2 | -3.062713 | 2.15205009 | 3 | 95.5 | 89.3 | -6.2 |
| VLLGESAVGKSSL | -0.6096688 | 0.47906255 | 8 | -0.2434736 | 1.40993245 | 4 | 60.4 | 54.2 | -6.2 |
| QAGELKWGTDEEKFITIF | -1.4372852 | 3.66884228 | 5 | -1.0052019 | 5.59019727 | 5 | 73.0 | 66.7 | -6.3 |
| SAEAPRLIEESSVDEVVVTNTVPHEVQKL | -2.8237098 | 0.30759065 | 2 | -2.1049518 | 0.4396109 | 3 | 87.6 | 81.1 | -6.5 |
| KVIAINVDDPDAANY | -0.2269037 | 0.66024222 | 11 | 0.16328547 | 1.225332 | 7 | 53.9 | 47.2 | -6.8 |
| TDSLVCKAGIIASARAGETRF | -3.4836989 | 0.11098291 | 2 | -2.5055112 | 0.72328924 | 3 | 91.8 | 85.0 | -6.8 |
| GGLDMLAEKLPNLTHL | -5.4210259 | 0.33460487 | 2 | -3.3198709 | 2.00026963 | 2 | 97.7 | 90.9 | -6.8 |
| GGLTQAPGNPVLAVQINQDKNF | -6.7238764 | 4.5847501 | 2 | -3.4401665 | 0.21221266 | 2 | 99.1 | 91.6 | -7.5 |
| LGETNPADSKPGTIRGDF | -2.4072935 | 1.22196276 | 5 | -1.7106431 | 1.85475703 | 4 | 84.1 | 76.6 | -7.5 |
| DLLKTNLYQDDAVTGEAAGL | -2.3534577 | 0.22755609 | 4 | -1.6417965 | 1.6028196 | 4 | 83.6 | 75.7 | -7.9 |
| GIEKTDPTTLTDEEINRF | 0.01873098 | 0.28758456 | 7 | 0.50275813 | 0.59472604 | 7 | 49.7 | 41.4 | -8.3 |
| ILGLDGAGKTTILY | -1.1124335 | 0.6976404 | 8 | -0.5878758 | 0.86156736 | 3 | 68.4 | 60.0 | -8.3 |
| GAGAKDELHIVEAEAM(15.9949)NY | -2.3199513 | 1.39966125 | 3 | -1.5577225 | 0.61833584 | 4 | 83.3 | 74.6 | -8.7 |
| LGETNPADSKPGTIRGDF | -2.5741052 | 1.08948038 | 14 | -1.7369281 | 1.74545029 | 14 | 85.6 | 76.9 | -8.7 |
| TLMVVGESGLGKSTL | -0.3898158 | 0.32969387 | 6 | 0.11674406 | 0.09151579 | 6 | 56.7 | 48.0 | -8.7 |
| NLVPLAESVVKL | -2.773842 | 1.62016454 | 3 | -1.8660533 | 1.58782896 | 3 | 87.2 | 78.5 | -8.8 |
| TIVGHTDVVKDVAW | -0.0030866 | 0.40953612 | 8 | 0.53702293 | 0.87152416 | 5 | 50.1 | 40.8 | -9.3 |
| LFRDGDILGKYVD | -4.7631812 | 1.50602262 | 5 | -2.7343954 | 0.13452494 | 3 | 96.4 | 86.9 | -9.5 |
| VGDLSPEITTEDIKSAFAPF | -3.0505112 | 0.40344457 | 3 | -1.9692566 | 2.08286632 | 2 | 89.2 | 79.7 | -9.6 |
| IQVTPQEKEAIERL | -1.5362379 | 2.48414451 | 2 | -0.8792969 | 3.63427852 | 4 | 74.4 | 64.8 | -9.6 |
| SRLAEISDVWEEMKTTL | -3.7536685 | 0.02902555 | 3 | -2.3184208 | 0.40476093 | 2 | 93.1 | 83.3 | -9.8 |
| LADKSYIEGYVPSQADVAVF | -1.2671708 | 0.08680659 | 3 | -0.6343864 | 0.19894244 | 3 | 70.6 | 60.8 | -9.8 |
| VAEDVDAGQIILQEAVPVKRGDTVATL | -2.8601095 | 0.54470955 | 7 | -1.7946408 | 2.26172464 | 4 | 87.9 | 77.6 | -10.3 |
| NVADVVIKFPEEEAPSTVL | -2.6815729 | 0.24211888 | 6 | -1.6561104 | 2.85857649 | 5 | 86.5 | 75.9 | -10.6 |
| GTADVHFERKADAL | -3.0970267 | 0.71795142 | 4 | -1.8405706 | 2.51558698 | 2 | 89.5 | 78.2 | -11.4 |
| VEIQTPKIISAASEGGANVF | -3.405089 | 0.58927141 | 4 | -1.9879192 | 0.8029837 | 3 | 91.4 | 79.9 | -11.5 |
| SEAAKVAANAPKGIL | -7.7335155 | 0.2533669 | 2 | -2.8279323 | 10.0944471 | 2 | 99.5 | 87.7 | -11.9 |
| EAADLGNEERKQKF | -5.6633331 | 1.75947212 | 2 | -2.6333124 | 7.34598763 | 3 | 98.1 | 86.1 | -11.9 |
| GFVTFDDHDPVDKIVLQ | -5.3114613 | 1.97421984 | 3 | -2.5344687 | 3.6533643 | 2 | 97.5 | 85.3 | -12.3 |
| EIQVNGGTVAEKLDW | -2.4976707 | 0.49569659 | 5 | -1.3611637 | 1.44527734 | 5 | 85.0 | 72.0 | -13.0 |
| APVNVTTEVKSVEMHHEAL | -2.4245375 | 0.09815916 | 2 | -1.30505 | 0.81831762 | 3 | 84.3 | 71.2 | -13.1 |
| LLSGAGEHLKTDLLLEPY | -5.5246674 | 0.77084597 | 4 | -2.3770137 | 4.74625603 | 2 | 97.9 | 83.9 | -14.0 |
| DAGAGIALNDHFVKL | -1.1622723 | 1.0743343 | 12 | -0.2785962 | 0.868369 | 15 | 69.1 | 54.8 | -14.3 |
| CANETVHGVEFDFIPDVKGAVL | -2.2321811 | 0.56319831 | 5 | -1.0037507 | 2.47687097 | 4 | 82.5 | 66.7 | -15.7 |
| DFKTLESSIQGL | -4.4738801 | 0.97832582 | 2 | -1.8436423 | 3.16782156 | 2 | 95.7 | 78.2 | -17.5 |
| LTPEQEKDKSHF | -5.7401706 | 0.91759207 | 4 | -1.9675187 | 2.27942875 | 2 | 98.2 | 79.6 | -18.5 |
| LGCVDIKDLPVSEQQERAF | -2.2940403 | 0.32868228 | 5 | -0.8310012 | 3.41709003 | 4 | 83.1 | 64.0 | -19.0 |
| DAGAGIALNDHFVKLISWYDNEFGY | -2.6934234 | 1.27478312 | 5 | -1.028913 | 0.02453956 | 3 | 86.6 | 67.1 | -19.5 |
| IGNLNESVTPADLEKVFAEH | -2.5270807 | 0.36770815 | 4 | -0.7311176 | 4.10284693 | 3 | 85.2 | 62.4 | -22.8 |
| DAGAGIALNDHFVKL | -1.4406897 | 1.40650707 | 5 | 0.06321371 | 0.81640133 | 12 | 73.1 | 48.9 | -24.2 |
| SAVIKFLELLSDDSNFGQF | -2.6891616 | 0.19674415 | 2 | -0.7056658 | 2.78210419 | 2 | 86.6 | 62.0 | -24.6 |
| IQSQEKPDRVL | -5.4838849 | 2.45433447 | 7 | -1.4108522 | 6.69652913 | 3 | 97.8 | 72.7 | -25.1 |
| TLVVKWGDEHIPGSPY | -4.8758096 | 1.06992391 | 5 | -1.1537216 | 4.85713403 | 5 | 96.7 | 69.0 | -27.7 |
| NTSTGGLLLPSDTKR | -3.8488977 | 0.46483222 | 3 | -0.3934666 | 7.72697049 | 2 | 93.5 | 56.8 | -36.7 |
| HVDGSLEKDRTL | -0.8049333 | 3.21006402 | 3 | 2.5186532 | 1.51063971 | 2 | 63.6 | 14.9 | -48.7 |
| RSLEDALAEAQRVNTKSQSAF | -2.684862 | 0.56810483 | 3 | 1.25929439 | 5.77130699 | 2 | 86.5 | 29.5 | -57.1 |
