## Extended data table 3 for "Covalent Protein Painting Reveals Structural Changes in the Proteome in Alzheimer Disease"

| Lysine site | P value | Mean1 | Mean2 | Difference | of differen | t ratio | df | q value | Entry | Entry name | Gene names |
|---|---|---|---|---|---|---|---|---|---|---|---|
| O43242#273-O43242-2#135 | 2.03E-10 | -3.476 | 0.7785 | -4.255 | 0.6688 | 6.362 | 24024 | 6.01E-07 | O43242 | PSMD3_HUMAN | PSMD3 |
| M0QXG8#248-Q92990#507 | 8.86E-09 | 0.0659 | -4.801 | 4.867 | 0.8459 | 5.753 | 24024 | 1.14E-05 | M0QXG8 | M0QXG8_HUMAN | GLMN |
| P14868#40 | 1.16E-08 | -6.619 | -2.802 | -3.817 | 0.6688 | 5.708 | 24024 | 1.14E-05 | P14868 | SYDC_HUMAN | DARS PIG40 |
| A0A087WTP3#448-Q92945#448 | 2.07E-07 | -0.8039 | -7.172 | 6.368 | 1.226 | 5.195 | 24024 | 0.000153 | Q92945 | FUBP2_HUMAN | KHSRP FUBP2 |
| P09874#528 | 2.89E-07 | -3.215 | -4.899 | 1.683 | 0.328 | 5.132 | 24024 | 0.000168 | P09874 | PARP1_HUMAN | PARP1 ADPRT PPOL |
| Q14690#1402 | 3.41E-07 | 1.662 | -5.16 | 6.822 | 1.338 | 5.101 | 24024 | 0.000168 | Q14690 | RRP5_HUMAN | PDCD11 KIAA0185 |
| P61353#27 | 5.64E-07 | -2.608 | -5.433 | 2.824 | 0.5644 | 5.005 | 24024 | 0.000223 | P61353 | RL27_HUMAN | RPL27 |
| D6RC14#83-E7ETU7#215-E9PF06#155-H0Y9G6#203-P09001#188 | 6.05E-07 | -0.7723 | -5.054 | 4.282 | 0.8579 | 4.991 | 24024 | 0.000223 | P09001 | RM03_HUMAN | MRPL3 MRL3 RPML3 |
| A0A0A0MTS2#469-P06744#454 | 2.54E-06 | -1.226 | -5.264 | 4.038 | 0.8579 | 4.706 | 24024 | 0.000834 | P06744 | G6PI_HUMAN | GPI |
| H0YFX9#59-P04908#96-P0C0S8#96-P16104#96-P20671#96-Q16777#96-Q6FI13#96 | 5.15E-06 | -5.381 | -7.305 | 1.924 | 0.422 | 4.56 | 24024 | 0.001523 | P04908 | H2A1B_HUMAN | HIST1H2AB H2AFM; HIST1H2AE H2AFA |
| A0A087X2B5#33-P35613#152-P35613-2#36 | 1.04E-05 | -4.711 | -8.305 | 3.595 | 0.8152 | 4.41 | 24024 | 0.002795 | P35613 | BASI_HUMAN | BSG UNQ6505/PRO21383 |
| P00390#501-P00390-2#458-P00390-3#472-P00390-4#448-P00390-5#419 | 1.65E-05 | 0.3538 | -2.43 | 2.784 | 0.6461 | 4.308 | 24024 | 0.004067 | P00390 | GSHR_HUMAN | GSR GLUR GRD1 |
| Q15084#102-Q15084-2#154-Q15084-3#99-Q15084-4#107-Q15084-5#150 | 2.24E-05 | -5.995 | -8.71 | 2.715 | 0.6402 | 4.241 | 24024 | 0.005087 | Q15084 | PDIA6_HUMAN | PDIA6 ERP5 P5 TXNDC7 |
| Q9UKD2#177 | 3.37E-05 | -0.4122 | -3.475 | 3.062 | 0.7384 | 4.147 | 24024 | 0.007127 | Q9UKD2 | MRT4_HUMAN | MRTO4 C1orf33 MRT4 |
