## Extended data table 1 for "Covalent Protein Painting Reveals Structural Changes in the Proteome in Alzheimer Disease"

| Sequence | Proteins | QuantSitePos | Quant site | INF | SITES | QUANT | rel.QUANT | rel.Position |
|---|---|---|---|---|---|---|---|---|
| AADVGKAGAEREF | P05141 | 6 | K | 1 | 2 | 2 | 1 | 7 |
| LPALVLKMQLF | Q8NDW8-6 | 7 | K | 1 | 1 | 1 | 1 | 4 |
| AIEGPSQAKIEY | O75369-3 | 9 | K | 1 | 1 | 1 | 1 | 3 |
| DSCSTSEHKL | F8VSZ4 | 9 | K | 1 | 1 | 1 | 1 | 1 |
| TFKDLVSCAY | P22607-3 | 3 | K | 1 | 1 | 1 | 1 | 7 |
| VKQHIDSFNY | Q9NW08 | 2 | K | 1 | 1 | 1 | 1 | 8 |
| GIQKELQF | P00338-4 | 4 | K | 1 | 1 | 1 | 1 | 4 |
| SGLEIVPNGITLPVDPEGKITGEAF | P52597 | 19 | K | 1 | 2 | 2 | 1 | 6 |
| KTSDLVPSF | Q8TEP8 | 1 | K | 1 | 2 | 2 | 1 | 8 |
| INGSLLEALKGVF | O75618 | 10 | K | 1 | 1 | 1 | 1 | 3 |
| VLDKLGDDEVRTDL | Q14444 | 4 | K | 1 | 1 | 1 | 1 | 10 |
| HLARLAKDSAAFF | A6NNB3 | 7 | K | 1 | 1 | 1 | 1 | 6 |
| PTYQKWISAASKLY | Q8ND04-3 | 12 | K | 1 | 2 | 2 | 1 | 2 |
| **QVSLQDKTGF** | **Q6GPI1** | **7** | **K** | **1** | **34** | **32** | **0.94** | **3** |
| VLEGKELEFY | Q5JR95 | 5 | K | 0 | 3 | 1 | 0.33 | 5 |
| GYVDFESAEDLEKALEL | P19338 | 13 | K | -1 | 3 | 1 | 0.33 | 4 |
| NDPSVQQDIKFLPF | P11021 | 10 | K | 1 | 2 | 0 | 0 | 4 |
| CFGPDGTGPNILTDITKGVQY | P13639 | 17 | K | -1 | 2 | 0 | 0.00 | 4 |
| **DAGAGIALNDHFVKL** | **P04406** | **14** | **K** | **-1** | **105** | **-47** | **-0.45** | **1** |
| IVLKEPISVSSEQVL | P00918 | 4 | K | 1 | 4 | -2 | -0.50 | 11 |
| VTGVHEEATEEDIHDKFAEY | Q9Y5S9 | 16 | K | 1 | 4 | -2 | -0.50 | 4 |
| LKDLLPGPVTL | Q86U90 | 2 | K | 1 | 5 | -3 | -0.60 | 9 |
| KVIAINVDDPDAANY | Q5SQT6 | 1 | K | 1 | 13 | -8 | -0.62 | 14 |
| KDQIYDIF | P60842 | 1 | K | 0 | 3 | -2 | -0.67 | 7 |
| GLDVEDVKF | Q92841-2 | 8 | K | -1 | 10 | -7 | -0.70 | 1 |
| KSLVASLAEPDFVVTDFAKF | P22314 | 19 | K | -1 | 7 | -5 | -0.71 | 1 |
| IVPDNPPYDKGAF | P68036 | 10 | K | -1 | 7 | -5 | -0.71 | 3 |
| SLLPATKL | E9PRQ6 | 7 | K | -1 | 11 | -8 | -0.73 | 1 |
| VLAPEGSVANKF | E5RIZ6 Q132 | 11 | K | -1 | 26 | -19 | -0.73 | 1 |
| VVKKSDVEAIF | P07910-2 | 4 | K | 1 | 8 | -6 | -0.75 | 7 |
| SEVELVQMVVDGVKL | P12277 | 14 | K | 1 | 8 | -6 | -0.75 | 1 |
| DIAHTPGVAADLSHIETKAAVKGY | P40926 | 22 | K | -1 | 8 | -6 | -0.75 | 2 |
| SEKGESSGKNVTLPAVF | H3BTP7 P365 | 3 | K | 1 | 9 | -7 | -0.78 | 14 |
| DLSKIGEEQSPEDAEDGPPELLF | Q09028-3 | 4 | K | -1 | 9 | -7 | -0.78 | 19 |
| KLLPQLTY | Q92688-2 | 1 | K | -1 | 9 | -7 | -0.78 | 7 |
| KILPTLEAVAAL | Q12905 | 1 | K | -1 | 10 | -8 | -0.80 | 11 |
| SVDIPLDKTVVNKDVF | K7ELC7 | 8 | K | -1 | 5 | -4 | -0.80 | 8 |
| DGIILPGK | P62913 | 8 | K | -1 | 11 | -9 | -0.82 | 0 |
| STSGSSNTDTGKVTGTLETKY | P45880-2 | 20 | K | 1 | 12 | -10 | -0.83 | 1 |
| VALDFEQEMATAASSSSLEKSY | P63261 | 20 | K | 1 | 12 | -10 | -0.83 | 2 |
| YPLEIDYGQDEEAVKKL | P09874 | 16 | K | 0 | 6 | -5 | -0.83 | 1 |
| KIGGIGTVPVGRVETGVL | Q5VTE0 | 1 | K | -1 | 12 | -10 | -0.83 | 17 |
| SVESTGVLPPDVLVSEAIKVL | E7EQB9 | 19 | K | 1 | 14 | -12 | -0.86 | 2 |
| DDHDSVDKIVIQKY | P09651-3 | 13 | K | -1 | 14 | -12 | -0.86 | 1 |
| GKESQAKDVIEEYF | P25398 | 2 | K | -1 | 15 | -13 | -0.87 | 12 |
| STDVSVDEVKAL | P49448 | 10 | K | 0 | 8 | -7 | -0.88 | 2 |
| EYIEENKY | Q00839 | 7 | K | -1 | 19 | -17 | -0.89 | 1 |
| ITFLEDLKSF | I3L3B0 | 8 | K | -1 | 29 | -26 | -0.90 | 2 |
| IAGKCGLVPVLAENY | J3KN47 | 4 | K | -1 | 20 | -18 | -0.90 | 11 |
| SGANKEKLEATINELV | P10599 | 5 | K | -1 | 21 | -19 | -0.90 | 11 |
| VKGEDFPANNIVKF | C9IZQ1 | 2 | K | 1 | 23 | -21 | -0.91 | 12 |
| KDLFDPIIEDRHGGY | G3V4N7 | 1 | K | -1 | 69 | -63 | -0.91 | 14 |
| TSFIGAIAIGDLVKSTL | F5GWF6 | 14 | K | -1 | 24 | -22 | -0.92 | 3 |
| QVKSGTIFDNF | K7EL50 | 3 | K | -1 | 24 | -22 | -0.92 | 8 |
| NVDLIPKFL | P31150 P503 | 7 | K | -1 | 13 | -12 | -0.92 | 2 |
| NVTEQEKIDKL | P06733 | 10 | K | -1 | 14 | -13 | -0.93 | 1 |
| AGKQLEDGRTLSDY | P0CG48 | 3 | K | -1 | 29 | -27 | -0.93 | 11 |
| KEGIPALDNFLDKL | P13639 | 13 | K | -1 | 228 | -213 | -0.93 | 1 |
| AIRNDEELNKL | P20671 | 10 | K | -1 | 122 | -114 | -0.93 | 1 |
| AKTAFDEAIAELDTLSEESY | P63104 | 2 | K | -1 | 31 | -29 | -0.94 | 18 |
| FEYIEENKY | Q00839 | 8 | K | -1 | 31 | -29 | -0.94 | 1 |
| ELELNGTEAKL | I3L3B0 | 10 | K | -1 | 79 | -75 | -0.95 | 1 |
| TTTPRPVIVEPLEQLDDEDGLPEKL | P23246 | 24 | K | 0 | 22 | -21 | -0.95 | 1 |

| | | | | | | | | |
|---|---|---|---|---|---|---|---|---|
| ESLTDPSKL | P07900 P082 | 8 | K | -1 | 24 | -23 | -0.96 | 1 |
| NGKNIEDVIAQGIGKL | H0YDD8 | 15 | K | -1 | 54 | -52 | -0.96 | 1 |
| EALDCILPPTRPTDKPL | Q5VTE0 | 15 | K | -1 | 54 | -52 | -0.96 | 2 |
| VKGLSEDTTEETLKESF | P19338 | 2 | K | -1 | 79 | -77 | -0.97 | 15 |
| TTVEDLGSKIL | P35613 | 9 | K | -1 | 42 | -41 | -0.98 | 2 |
| LGKVTIAQGGVLPNIQAVL | P20671 | 3 | K | -1 | 72 | -71 | -0.99 | 16 |
| DVTEESIKEFF | E7EX17 P235 | 8 | K | -1 | 1 | -1 | -1 | 3 |
| QVINDGDKPKVQVSY | P0DMV9 | 8 | K | -1 | 1 | -1 | -1 | 7 |
| EEIVKEVSTY | Q5VTE0 | 5 | K | -1 | 1 | -1 | -1 | 5 |
| DLERKVESLQEEIAFL | B0YJC4 | 5 | K | -1 | 4 | -4 | -1.00 | 11 |
| QEALDAAGDKLVVVDF | P10599 | 10 | K | -1 | 2 | -2 | -1 | 6 |
| GYVDFESAEDLEKALELTGL | P19338 | 13 | K | -1 | 2 | -2 | -1 | 7 |
| SELAEDKENY | P08238 | 7 | K | -1 | 1 | -1 | -1 | 3 |
| AIRNDEELNKLLGGVTIAQGGVLPNIQAVL | Q96QV6 | 10 | K | -1 | 1 | -1 | -1 | 20 |
| AKTAFDEAIAELDTLNEESY | P31946 | 2 | K | -1 | 4 | -4 | -1.00 | 18 |
| SKTPELNLDQFHDKTPY | P27824 | 2 | K | -1 | 3 | -3 | -1.00 | 15 |
| VEKLTEVSISSDAFFPF | P31939 | 3 | K | -1 | 2 | -2 | -1 | 14 |
| QLALKDCEECIQLEPTFIKGY | P31948 | 19 | K | -1 | 2 | -2 | -1 | 2 |
| KFIDTTSKF | Q92901 | 8 | K | -1 | 2 | -2 | -1 | 1 |
| YTEPVIAGLDPKTF | J3QKR3 | 12 | K | -1 | 1 | -1 | -1 | 2 |
| VSISDLLVPKDLGTESQIF | P50395 | 10 | K | -1 | 1 | -1 | -1 | 9 |
| NLPKTCDISFSDPDDLLNF | P61081 | 4 | K | -1 | 1 | -1 | -1 | 15 |
| FVMGVNHEKY | P04406 | 9 | K | -1 | 3 | -3 | -1.00 | 1 |
| FHSLSEKY | P10599 | 7 | K | -1 | 2 | -2 | -1 | 1 |
| IAKIPNFW | Q01105-3 | 3 | K | -1 | 2 | -2 | -1 | 5 |
| VGVNLPQKAGGF | P00558 | 8 | K | -1 | 1 | -1 | -1 | 4 |
| LYEDGEDQPKQVY | P34932 | 10 | K | -1 | 1 | -1 | -1 | 3 |
| ADLLLEKETL | Q99459 | 7 | K | -1 | 1 | -1 | -1 | 3 |
| IAKVATAQDDITGDGTTSNVL | B4DPJ8 | 3 | K | -1 | 7 | -7 | -1.00 | 18 |
| VKLEAEGVPEVSEKY | O76003 | 14 | K | -1 | 5 | -5 | -1.00 | 1 |
| NAVTIEDVQKL | Q9Y617 | 10 | K | -1 | 5 | -5 | -1.00 | 1 |
| DIQKDLKDL | P07195 | 7 | K | -1 | 4 | -4 | -1.00 | 2 |
| SKVTEGLTDVILY | O60506-3 | 2 | K | -1 | 3 | -3 | -1.00 | 11 |
| MFQYDSTHGKF | P04406 | 10 | K | -1 | 3 | -3 | -1.00 | 1 |
| KLDTLCDLY | P60842 | 1 | K | -1 | 3 | -3 | -1.00 | 8 |
| NLVPLAESVVKL | K7ERJ1 | 11 | K | -1 | 2 | -2 | -1 | 1 |
| ILIQDGSQNTNVDKPL | M0QYH3 | 14 | K | -1 | 2 | -2 | -1 | 2 |
| KLIINSLY | P14625 | 1 | K | -1 | 2 | -2 | -1 | 7 |
| NLKPEEVF | P49915 | 3 | K | -1 | 2 | -2 | -1 | 5 |
| KATAVMPDGQFKDISLSDY | Q06830 | 1 | K | -1 | 2 | -2 | -1 | 18 |
| VVFDDSEPVQKVL | Q13283 | 11 | K | -1 | 2 | -2 | -1 | 2 |
| SEDVSISKF | Q15029 | 8 | K | -1 | 2 | -2 | -1 | 1 |
| IANTGMDTDKIKIF | F5GWF6 | 12 | K | -1 | 1 | -1 | -1 | 2 |
| VANNVTLPAGEQRKDVY | G5E972 P421 | 14 | K | -1 | 1 | -1 | -1 | 3 |
| DFTGAVEDISKIPEQSVL | P00505 | 11 | K | -1 | 1 | -1 | -1 | 7 |
| KMSVQPTVSLGGF | P06748-3 | 1 | K | -1 | 1 | -1 | -1 | 12 |
| VVKTGVVPQLVKLLGASELPIVTPAL | P52292 | 3 | K | -1 | 1 | -1 | -1 | 23 |
| AGYIEDLKKF | P54709 | 9 | K | -1 | 1 | -1 | -1 | 1 |
| DVIAQAQSGTGKTATF | P60842 | 12 | K | -1 | 1 | -1 | -1 | 4 |
| RENTQTTIKLF | P61978-3 | 9 | K | -1 | 1 | -1 | -1 | 2 |
| ALEKLGAVF | Q15393 | 4 | K | -1 | 1 | -1 | -1 | 5 |
| KADEGISF | Q06830 | 1 | K | -1 | 5 | -5 | -1.00 | 7 |
| TLSGLVEVKEL | Q6FI81 | 9 | K | -1 | 3 | -3 | -1.00 | 2 |
| AKVDLNSNGF | P13797 | 2 | K | -1 | 2 | -2 | -1 | 8 |
| AGKQLEDGRTL | Q5PY61 | 3 | K | -1 | 2 | -2 | -1 | 8 |
| CQVGEKPY | I3L2R6 | 6 | K | -1 | 1 | -1 | -1 | 2 |
| AIPQPDLTKL | Q15366-4 | 9 | K | -1 | 9 | -9 | -1.00 | 1 |
| GTKGNVQVVIPF | P22314 | 3 | K | -1 | 6 | -6 | -1.00 | 9 |
| TSSGSANTETTKVTGSLETKY | C9JI87 | 20 | K | -1 | 5 | -5 | -1.00 | 1 |
| VEEITDLPIKL | O00232 | 10 | K | -1 | 5 | -5 | -1.00 | 1 |
| VARGDLGIEIPAEKVF | P14618-3 | 14 | K | -1 | 5 | -5 | -1.00 | 2 |
| LIPNATQPESKVF | P31946 | 11 | K | -1 | 5 | -5 | -1.00 | 2 |
| DLSKIGEEQSAEDAEDGPPELLF | Q16576 | 4 | K | -1 | 5 | -5 | -1.00 | 19 |
| LTAEVLELAGNASKDL | Q71UI9-2 | 14 | K | -1 | 5 | -5 | -1.00 | 2 |

| | | | | | | | | |
|---|---|---|---|---|---|---|---|---|
| KFIIPQIVKY | A8MW50 | 9 | K | -1 | 4 | -4 | -1.00 | 1 |
| LSSAVDHGSDEVKF | F5GWF6 | 13 | K | -1 | 4 | -4 | -1.00 | 1 |
| KVDNDENEHQLSL | P06748-3 | 1 | K | -1 | 4 | -4 | -1.00 | 12 |
| KATLPSPDKLPGF | P22314 | 1 | K | -1 | 4 | -4 | -1.00 | 12 |
| CTKEAAQEAVKLY | O60506-3 | 11 | K | -1 | 3 | -3 | -1.00 | 2 |
| LAKAIANECQANF | P55072 | 3 | K | -1 | 3 | -3 | -1.00 | 10 |
| VQYPVEHPDKF | P55072 | 10 | K | -1 | 3 | -3 | -1.00 | 1 |
| AAVPVQDLGSTVIKEVL | Q9BWD1 | 14 | K | -1 | 3 | -3 | -1.00 | 3 |
| KYDYNSGEELESY | B0AZV0 | 1 | K | -1 | 2 | -2 | -1 | 12 |
| AAAGGVSHDELLDLAKF | E7ERZ4 | 16 | K | -1 | 2 | -2 | -1 | 1 |
| KNLVLVDELDSLSPILF | J3QKV4 | 1 | K | -1 | 2 | -2 | -1 | 16 |
| TVGKEPFPTIY | O43776 | 4 | K | -1 | 2 | -2 | -1 | 7 |
| ANTVTEEILEKAF | O60506-3 | 11 | K | -1 | 2 | -2 | -1 | 2 |
| CVQQLKEF | P07900 | 6 | K | -1 | 2 | -2 | -1 | 2 |
| VLVLYDELKKVI | P12236 | 10 | K | -1 | 2 | -2 | -1 | 2 |
| NLVKDSATGL | P26368 | 4 | K | -1 | 2 | -2 | -1 | 6 |
| VFKEDGQEY | P47813 | 3 | K | -1 | 2 | -2 | -1 | 6 |
| SATMPSDVLEVTKKF | P60842 | 14 | K | -1 | 2 | -2 | -1 | 1 |
| AHELPKYGVKVGL | Q5T7N0 | 10 | K | -1 | 2 | -2 | -1 | 3 |
| TVKVEDLTF | Q99873-3 | 3 | K | -1 | 2 | -2 | -1 | 6 |
| LFRDGDILGKYVD | B8ZZ54 | 10 | K | -1 | 1 | -1 | -1 | 3 |
| IIAIPTGVKVF | P00395 | 9 | K | -1 | 1 | -1 | -1 | 2 |
| ELFTELAEDKENY | P07900 | 10 | K | -1 | 1 | -1 | -1 | 3 |
| QVVAVKAPGF | P10809 | 6 | K | -1 | 1 | -1 | -1 | 4 |
| EVAKDFPEYTF | P13667 | 4 | K | -1 | 1 | -1 | -1 | 7 |
| AKTAFDEAIAELDTLNEDSYKDSTL | P27348 | 2 | K | -1 | 1 | -1 | -1 | 23 |
| KHTGPNSPDTANDGFVRL | P55795 | 1 | K | -1 | 1 | -1 | -1 | 17 |
| MLDKLTGVF | P62701 | 4 | K | -1 | 1 | -1 | -1 | 5 |
| ELFADIVPKTAENF | Q08752 | 9 | K | -1 | 1 | -1 | -1 | 5 |
| SIYPHGSTDKLPY | Q5TBG5 | 10 | K | -1 | 1 | -1 | -1 | 3 |
| FLKVSELF | Q9UNH7 | 3 | K | -1 | 1 | -1 | -1 | 5 |
| SSKIIGINGDFF | E7EQR6 | 3 | K | -1 | 4 | -4 | -1.00 | 9 |
| LVLQEILESEEKGDPNKPSGF | O00151 | 17 | K | -1 | 4 | -4 | -1.00 | 4 |
| GYPNLKSVNELIY | A8MUD9 | 6 | K | -1 | 2 | -2 | -1 | 7 |
| YVSNIDGTHIAKTL | K7ENA0 | 12 | K | -1 | 2 | -2 | -1 | 2 |
| GSSKLVPVGY | P24534 | 4 | K | -1 | 2 | -2 | -1 | 6 |
| SKLPIGDVATQY | Q99832 Q99 | 2 | K | -1 | 2 | -2 | -1 | 10 |
| GRDLLDDLKSEL | E9PHT9 | 9 | K | -1 | 1 | -1 | -1 | 3 |
| DIDEAEEGVKDL | P20042 | 10 | K | -1 | 1 | -1 | -1 | 2 |
| KVGSAADIPINISETDLSLL | P21333 | 1 | K | -1 | 1 | -1 | -1 | 19 |
| GFVTFDDHDPVDKIVL | P22626 | 13 | K | -1 | 1 | -1 | -1 | 3 |
| NFGQKEKPYFPIPEEY | Q00839 | 5 | K | -1 | 1 | -1 | -1 | 11 |
| DDSEPVQKVL | Q13283 | 8 | K | -1 | 4 | -4 | -1.00 | 2 |
| AVNEVVAGIKEY | Q9UBU8-2 | 10 | K | -1 | 3 | -3 | -1.00 | 2 |
| AALPKATIL | Q96AG4 | 5 | K | -1 | 9 | -9 | -1.00 | 4 |
| KLLDVVHPAAKTL | Q99832 | 11 | K | -1 | 9 | -9 | -1.00 | 2 |
| ISVEEVHDDGTPTSKTF | B4DXI8 | 15 | K | -1 | 6 | -6 | -1.00 | 2 |
| YVPAEPKLAF | A8MUD9 | 7 | K | -1 | 5 | -5 | -1.00 | 3 |
| AVNAAQDSTDLVAKL | E7EQR6 | 14 | K | -1 | 4 | -4 | -1.00 | 1 |
| VRNLANTVTEEILEKAF | O60506-3 | 15 | K | -1 | 4 | -4 | -1.00 | 2 |
| ILSADFPALVVKASGL | P22102 | 12 | K | -1 | 4 | -4 | -1.00 | 4 |
| VPGVVSTVVPDSAHKLF | P26368 | 15 | K | -1 | 4 | -4 | -1.00 | 2 |
| AENSGVKANEVISKLY | P50990 | 14 | K | -1 | 4 | -4 | -1.00 | 2 |
| APIKVGDAIPAVEVF | P30044-3 | 4 | K | -1 | 3 | -3 | -1.00 | 11 |
| SALDILIKNY | Q86VP6 | 8 | K | -1 | 3 | -3 | -1.00 | 2 |
| AAVPGKTF | I3L3D5 | 6 | K | -1 | 2 | -2 | -1 | 2 |
| KVNQIGSVTESL | P06733 | 1 | K | -1 | 2 | -2 | -1 | 11 |
| QLLQEAGIKTAF | E9PBS1 | 9 | K | -1 | 1 | -1 | -1 | 3 |
| LNALKSVINDPIYKENIM | O75795 | 14 | K | -1 | 1 | -1 | -1 | 4 |
| IFQEVKSSDIAKTF | P13797 | 12 | K | -1 | 1 | -1 | -1 | 2 |
| SVSDKTGLVEF | P31939 | 5 | K | -1 | 1 | -1 | -1 | 6 |
| NEDNGIIKAF | H3BM89 P36 | 8 | K | -1 | 10 | -10 | -1.00 | 2 |
| ISNSSDALDKIRY | P07900 | 10 | K | -1 | 5 | -5 | -1.00 | 3 |
| GCSKEEIVQFF | P31942 P557 | 4 | K | -1 | 4 | -4 | -1.00 | 7 |

| | | | | | | | | |
|---|---|---|---|---|---|---|---|---|
| DQLLAEEKTISAKY | P35579 | 13 | K | -1 | 4 | -4 | -1.00 | 1 |
| LLSLDDSVDETEAVKRY | Q9BXP5-3 | 15 | K | -1 | 4 | -4 | -1.00 | 2 |
| GHEGAGIVESVGEGVTKL | D6R9G2 | 17 | K | -1 | 3 | -3 | -1.00 | 1 |
| ESKSVVPGGGAVEAAL | E7EQR6 | 3 | K | -1 | 3 | -3 | -1.00 | 13 |
| VLVLYDEIKKY | P12235 | 10 | K | -1 | 3 | -3 | -1.00 | 1 |
| CIQALPEFDGKRF | P14625 | 11 | K | -1 | 3 | -3 | -1.00 | 2 |
| DANTLAEKDEF | P0DMV9 | 8 | K | -1 | 2 | -2 | -1 | 3 |
| QHGKVEIIANDQGNRTTPSYVAF | P0DMV9 P11 | 4 | K | -1 | 2 | -2 | -1 | 19 |
| SDPFVEAEKSNL | P34932 | 9 | K | -1 | 2 | -2 | -1 | 3 |
| SGNPIKVSF | P35637 | 6 | K | -1 | 2 | -2 | -1 | 3 |
| NNLVLFDKATY | P62851 | 8 | K | -1 | 2 | -2 | -1 | 3 |
| QYPPPPPPPPPSRK | Q92841-2 | 14 | K | -1 | 1 | -1 | -1 | 0 |
| GILADATEQVGQHKDAY | Q9BZZ5-3 | 14 | K | -1 | 1 | -1 | -1 | 3 |
| TPDSPTIKPSPAASKEY | P30048 | 15 | K | -1 | 7 | -7 | -1.00 | 2 |
| ISDKDASIVGFFDDSFSEAHSEF | G5EA52 | 4 | K | -1 | 6 | -6 | -1.00 | 19 |
| QLDKDGVVLF | F5H8J2 | 4 | K | -1 | 3 | -3 | -1.00 | 6 |
| TELAEDKENY | P07900 | 7 | K | -1 | 3 | -3 | -1.00 | 3 |
| FKPEELVDY | P22392 Q32( | 2 | K | -1 | 2 | -2 | -1 | 7 |
| RALSTGEKGFGY | C9J5S7 | 8 | K | -1 | 1 | -1 | -1 | 4 |
| ELFADKVPKTAENF | C9J5S7 P629 | 9 | K | -1 | 1 | -1 | -1 | 5 |
| DLDSSCKEAADGY | P31948 | 7 | K | -1 | 1 | -1 | -1 | 6 |
| SLPIKESEIIDFF | E9PMM9 E9I | 5 | K | -1 | 3 | -3 | -1.00 | 8 |
| GLDKSEDKVIAVY | P38646 | 4 | K | -1 | 3 | -3 | -1.00 | 9 |
| DAAGDKLVVVDF | P10599 | 6 | K | -1 | 2 | -2 | -1 | 6 |
| LGETNPADSKPGTIRGDF | P15531-2 P2 | 10 | K | -1 | 1 | -1 | -1 | 8 |
| TGKTITDVINIGIGGSDLGPL | K7ENA0 | 3 | K | -1 | 11 | -11 | -1.00 | 18 |
| KVFLENVIRDAVTY | P62805 | 1 | K | -1 | 11 | -11 | -1.00 | 13 |
| ALSVETDYTFPLAEKVKAF | F8VWS0 | 17 | K | -1 | 6 | -6 | -1.00 | 2 |
| LDANTLAEKDEF | P0DMV9 | 9 | K | -1 | 6 | -6 | -1.00 | 3 |
| KGFGYAEFEDLDSLL | E7EX17 | 1 | K | -1 | 5 | -5 | -1.00 | 14 |
| KQVHPDTGISSKAM | P06899 | 12 | K | -1 | 3 | -3 | -1.00 | 2 |
| FLKIIQLLDDYPKCF | F8VWS0 | 3 | K | -1 | 2 | -2 | -1 | 12 |
| SSVPPPSAPPPKKSL | Q14978-3 | 13 | K | -1 | 2 | -2 | -1 | 2 |
| LGNLPYDVTEESIKEFF | E7EX17 | 14 | K | -1 | 1 | -1 | -1 | 3 |
| SGFPFEKGSVQY | P35659 | 7 | K | -1 | 1 | -1 | -1 | 5 |
| CQLILDPIFKVF | P13639 | 10 | K | -1 | 24 | -24 | -1.00 | 2 |
| SGDKSENVQDLLLLDVTPL | P11142 | 4 | K | -1 | 21 | -21 | -1.00 | 15 |
| AKAAFDDAIAELDTLSEESY | P62258 | 2 | K | -1 | 20 | -20 | -1.00 | 18 |
| RSTLEPVEKAL | P0DMV9 | 9 | K | -1 | 12 | -12 | -1.00 | 2 |
| NDELEIIEGMKF | E7ESH4 | 11 | K | -1 | 11 | -11 | -1.00 | 1 |
| NTVQGDIDAIFKDL | Q15056 | 12 | K | -1 | 11 | -11 | -1.00 | 2 |
| VKQIESKTAF | P10599 | 2 | K | -1 | 9 | -9 | -1.00 | 8 |
| EVKSTNGDTFLGGEDFDQALL | P38646 | 3 | K | -1 | 6 | -6 | -1.00 | 18 |
| KMSVQPTVSLGGFEITPPVVL | P06748-3 | 1 | K | -1 | 5 | -5 | -1.00 | 20 |
| GAQGPKGGSGSGPTIEEVD | P0DMV9 | 6 | K | -1 | 3 | -3 | -1.00 | 13 |
| NKSAPELKTGISDVF | P19338 | 2 | K | -1 | 2 | -2 | -1 | 13 |
| LADKSYIEGYVPSQADVAVF | P24534 | 4 | K | -1 | 2 | -2 | -1 | 16 |
| IQVDIGGGQTKTF | P11021 | 11 | K | -1 | 22 | -22 | -1.00 | 2 |
| GKVTIAQGGVLPNIQAVL | P20671 | 2 | K | -1 | 17 | -17 | -1.00 | 16 |
| SASGELGNGNIKL | P12004 | 12 | K | -1 | 10 | -10 | -1.00 | 1 |
| AYVEFENPDEAEKAL | Q15287-3 | 13 | K | -1 | 9 | -9 | -1.00 | 2 |
| FDPANGKF | P13639 | 7 | K | -1 | 8 | -8 | -1.00 | 1 |
| ELAGNASKDL | Q71UI9-2 | 8 | K | -1 | 8 | -8 | -1.00 | 2 |
| ELLDSPGKVLL | P22234 | 8 | K | -1 | 4 | -4 | -1.00 | 3 |
| KVVETDPSPY | C9IZA5 | 1 | K | -1 | 2 | -2 | -1 | 9 |
| KPSDEHKTDLNPDNLQGGDDLDPNYVL | G3V4N7 | 1 | K | -1 | 1 | -1 | -1 | 26 |
| NSGKVDIVAINDPF | P04406 | 4 | K | -1 | 1 | -1 | -1 | 10 |
| GGVKLEDLIVKDGLTDVY | P24752 | 4 | K | -1 | 1 | -1 | -1 | 14 |
| KLDNLVAIL | P29401 | 1 | K | -1 | 1 | -1 | -1 | 8 |
| SSEVKEDLNGPF | Q9UQ35 | 5 | K | -1 | 1 | -1 | -1 | 7 |
| SSAVDHGSDEVKF | F5GWF6 | 12 | K | -1 | 10 | -10 | -1.00 | 1 |
| DDHDPVDKIVL | P22626 | 8 | K | -1 | 9 | -9 | -1.00 | 3 |
| SLPIKESEIIDF | E9PMM9 E9I | 5 | K | -1 | 6 | -6 | -1.00 | 7 |
| VVQEPGDYEVSVKF | P21333 | 13 | K | -1 | 2 | -2 | -1 | 1 |

| | | | | | | | | |
|---|---|---|---|---|---|---|---|---|
| KGLLPEELTPL | P13804 | 1 | K | -1 | 1 | -1 | -1 | 10 |
| TGKDVNFEFPEFQL | P62081 | 3 | K | -1 | 44 | -44 | -1.00 | 11 |
| NVEFDDSQDKAVL | P00918 | 10 | K | -1 | 20 | -20 | -1.00 | 3 |
| KLDEDEDEDDADLSKY | O95218 | 15 | K | -1 | 14 | -14 | -1.00 | 1 |
| KSLVASLAEPDFVVTDF | P22314 | 1 | K | -1 | 13 | -13 | -1.00 | 16 |
| KVAGQDGSVVQF | P61956-2 | 1 | K | -1 | 11 | -11 | -1.00 | 11 |
| SVKILAPGIEPF | I3L0P4 | 3 | K | -1 | 7 | -7 | -1.00 | 9 |
| SGLIADAKTL | P28066 | 8 | K | -1 | 7 | -7 | -1.00 | 2 |
| YVKLVEAL | P25398 | 3 | K | -1 | 3 | -3 | -1.00 | 5 |
| KPGMVVTF | Q5VTE0 | 1 | K | -1 | 2 | -2 | -1 | 7 |
| LFSCGDEGKAFLDTFTLL | E7EQD3 | 9 | K | -1 | 1 | -1 | -1 | 9 |
| TKMKEIAEAY | P0DMV9 P11 | 2 | K | -1 | 1 | -1 | -1 | 8 |
| IVLAKDFEKAY | P43686 | 9 | K | -1 | 1 | -1 | -1 | 2 |
| GEKFEDENF | P62937 Q08 | 3 | K | -1 | 26 | -26 | -1.00 | 6 |
| SGIESSSPEVKGY | Q15366-4 | 11 | K | -1 | 24 | -24 | -1.00 | 2 |
| YNEATGGKY | P68371 | 8 | K | -1 | 21 | -21 | -1.00 | 1 |
| KDISLSDY | Q06830 | 1 | K | -1 | 20 | -20 | -1.00 | 7 |
| NSGKVDIVAINDPFIDLNY | P04406 | 4 | K | -1 | 19 | -19 | -1.00 | 15 |
| NFVKGVVDSDDLPL | P14625 | 4 | K | -1 | 19 | -19 | -1.00 | 10 |
| KATLVESSTSGF | P49321-3 | 1 | K | -1 | 13 | -13 | -1.00 | 11 |
| QGLIVPDNPPYDKGAF | P68036 | 13 | K | -1 | 13 | -13 | -1.00 | 3 |
| RIDFYFDENPYFENKVL | Q01105-3 | 15 | K | -1 | 13 | -13 | -1.00 | 2 |
| AAVSAAIVKEL | Q9BWD1 | 9 | K | -1 | 12 | -12 | -1.00 | 2 |
| AEGVIPDEAKAL | Q05519 | 10 | K | -1 | 11 | -11 | -1.00 | 2 |
| GIKDDVF | P00338-4 | 3 | K | -1 | 10 | -10 | -1.00 | 4 |
| NYEGSPIKVTL | P06748-3 | 8 | K | -1 | 9 | -9 | -1.00 | 3 |
| ALELLFDQLHEGAKAL | P22061 | 14 | K | -1 | 9 | -9 | -1.00 | 2 |
| KVNFPENGFLSPDKL | P31689 | 14 | K | -1 | 9 | -9 | -1.00 | 1 |
| YFDENPYFENKVL | Q01105-3 | 11 | K | -1 | 9 | -9 | -1.00 | 2 |
| AGIDSSSPEVKGY | Q15365 | 11 | K | -1 | 9 | -9 | -1.00 | 2 |
| SGLLVTVGEVLEKSLL | Q96M27-3 | 13 | K | -1 | 8 | -8 | -1.00 | 3 |
| LFDKPVSPL | G3V4N7 | 4 | K | -1 | 7 | -7 | -1.00 | 5 |
| KVTAVPTLL | I3L2R6 | 1 | K | -1 | 7 | -7 | -1.00 | 8 |
| VVFDSIESAKKF | P05455 | 11 | K | -1 | 7 | -7 | -1.00 | 1 |
| GTVKDELTESPKY | Q8NC51 | 12 | K | -1 | 7 | -7 | -1.00 | 1 |
| NFIVPTGKTGL | Q96AE4 | 8 | K | -1 | 7 | -7 | -1.00 | 3 |
| VLVGDGGTGKTTF | B5MDF5 | 10 | K | -1 | 6 | -6 | -1.00 | 3 |
| IKAAGVNVEPF | P05386 | 2 | K | -1 | 6 | -6 | -1.00 | 9 |
| DVTEESIKEF | P23588 | 8 | K | -1 | 6 | -6 | -1.00 | 2 |
| KDLDVAIL | P40925 | 1 | K | -1 | 6 | -6 | -1.00 | 7 |
| DADGTGTIDVKEL | P41208 | 11 | K | -1 | 6 | -6 | -1.00 | 2 |
| VVKTGVVPQLVKLL | P52292 | 12 | K | -1 | 6 | -6 | -1.00 | 2 |
| KFVEGLPINDF | P40925 | 1 | K | -1 | 5 | -5 | -1.00 | 10 |
| EEVHDLERKY | P55209 Q99 | 9 | K | -1 | 5 | -5 | -1.00 | 1 |
| TSKIPALAVEMPGSADISGL | Q14157-3 | 3 | K | -1 | 5 | -5 | -1.00 | 17 |
| VTGIKVVDLLAPY | F8VPV9 | 5 | K | -1 | 4 | -4 | -1.00 | 8 |
| QVKSGTIF | K7EL50 | 3 | K | -1 | 4 | -4 | -1.00 | 5 |
| GSLVDEFKEL | P12956 | 8 | K | -1 | 4 | -4 | -1.00 | 2 |
| KAPVPTGEVY | P27824 | 1 | K | -1 | 4 | -4 | -1.00 | 9 |
| KLDTLCDLYETL | P60842 | 1 | K | -1 | 4 | -4 | -1.00 | 11 |
| KHLNEIDLF | Q15185 | 1 | K | -1 | 4 | -4 | -1.00 | 8 |
| KASGDGESLDEDSEF | Q99733 | 1 | K | -1 | 4 | -4 | -1.00 | 14 |
| TLQPKLPITVL | P31939 | 5 | K | -1 | 3 | -3 | -1.00 | 6 |
| DASKVTASGPGLSSY | O75369-3 | 4 | K | -1 | 2 | -2 | -1 | 11 |
| VKGLSEDTTEETL | P19338 | 2 | K | -1 | 2 | -2 | -1 | 11 |
| KCDACGKAFSTCTDL | Q14591 | 7 | K | -1 | 2 | -2 | -1 | 8 |
| TKILIPPKGLF | Q15459 | 2 | K | -1 | 2 | -2 | -1 | 9 |
| TDEEPVKKLL | O14979-3 | 8 | K | -1 | 1 | -1 | -1 | 2 |
| SKDIVENYF | Q13263 | 2 | K | -1 | 11 | -11 | -1.00 | 7 |
| VQTKGTGASGSF | P16401 | 4 | K | -1 | 10 | -10 | -1.00 | 8 |
| EIENNPTVKASGY | P51858-3 | 9 | K | -1 | 7 | -7 | -1.00 | 4 |
| SALILHDDEVTVTEDKINAL | P05386 | 16 | K | -1 | 1 | -1 | -1 | 4 |
| CYVEFDEVDSLKEAL | Q15056 | 12 | K | -1 | 1 | -1 | -1 | 3 |
| ESLTDPSKLDSGKEL | P07900 P082 | 13 | K | -1 | 73 | -73 | -1.00 | 2 |

| | | | | | | | | |
|---|---|---|---|---|---|---|---|---|
| LIPNASQAESKVF | P63104 | 11 | K | -1 | 41 | -41 | -1.00 | 2 |
| AIRNDEELNKLL | P20671 Q96( | 10 | K | -1 | 39 | -39 | -1.00 | 2 |
| SKFGEVVDCTL | Q14103-3 | 2 | K | -1 | 27 | -27 | -1.00 | 9 |
| AKDIGFIKLD | P62273 | 8 | K | -1 | 23 | -23 | -1.00 | 2 |
| DVSGYPTLKIF | P13667 | 9 | K | -1 | 16 | -16 | -1.00 | 2 |
| GSAGPPPTGEEDTAEKDEL | P11021 | 16 | K | -1 | 14 | -14 | -1.00 | 3 |
| QLAIRNDEELNKL | P20671 | 12 | K | -1 | 14 | -14 | -1.00 | 1 |
| DLSLSDLNEVPVKEL | I3L223 Q96A | 13 | K | -1 | 12 | -12 | -1.00 | 2 |
| KLITPAVVSERL | P62851 | 1 | K | -1 | 12 | -12 | -1.00 | 11 |
| AKTAFDEAIAELDTLNEDSY | P27348 | 2 | K | -1 | 10 | -10 | -1.00 | 18 |
| IIDPNGVIKHL | P30048 | 9 | K | -1 | 8 | -8 | -1.00 | 2 |
| IIPQIVKY | P07195 | 7 | K | -1 | 6 | -6 | -1.00 | 1 |
| GGSVTGATCKEL | P60174 | 10 | K | -1 | 6 | -6 | -1.00 | 2 |
| ATAAGSEDAEKKVL | P16989-3 | 12 | K | -1 | 2 | -2 | -1 | 2 |
| DSGFGGGAGVETGGKLL | E9PB61 | 15 | K | -1 | 55 | -55 | -1.00 | 2 |
| NAAKVPADTEVVCAPPTAY | P60174 | 4 | K | -1 | 51 | -51 | -1.00 | 15 |
| LIANATNPESKVF | P27348 | 11 | K | -1 | 46 | -46 | -1.00 | 2 |
| SATEETLQEVFEKATF | P19338 | 13 | K | -1 | 44 | -44 | -1.00 | 3 |
| VIEFTEQTAPKIF | F5H8J2 | 11 | K | -1 | 40 | -40 | -1.00 | 2 |
| TGLAAAIAGAKL | Q8N8S7 | 11 | K | -1 | 40 | -40 | -1.00 | 1 |
| GDLGGPIITTQVTIPKDL | P61978-3 | 16 | K | -1 | 39 | -39 | -1.00 | 2 |
| FDENPYFENKVL | Q01105-3 | 10 | K | -1 | 38 | -38 | -1.00 | 2 |
| AGAAVDELGKVL | P30086 | 10 | K | -1 | 35 | -35 | -1.00 | 2 |
| LVKTGTITTF | P13639 | 3 | K | -1 | 31 | -31 | -1.00 | 7 |
| TTVEDLGSKILL | P35613 | 9 | K | -1 | 30 | -30 | -1.00 | 3 |
| GVQGFPTIKIF | Q15084 | 9 | K | -1 | 30 | -30 | -1.00 | 2 |
| ETAAPAVAETPDIKLF | M0QZN2 | 14 | K | -1 | 29 | -29 | -1.00 | 2 |
| VGGLSPDTSEEQIKEY | O14979-3 | 14 | K | -1 | 29 | -29 | -1.00 | 2 |
| GEKFEDENFIL | C9J5S7 P629: | 3 | K | -1 | 28 | -28 | -1.00 | 8 |
| IGAIAIGDLVKSTL | F5GWF6 | 11 | K | -1 | 28 | -28 | -1.00 | 3 |
| STASDNQPTVTIKVY | P11021 | 13 | K | -1 | 28 | -28 | -1.00 | 2 |
| GYVDFESAEDLEKAL | P19338 | 13 | K | -1 | 26 | -26 | -1.00 | 2 |
| QEVTNNLEFAKEL | Q14444 | 11 | K | -1 | 26 | -26 | -1.00 | 2 |
| ELLDSPGKVL | P22234 | 8 | K | -1 | 21 | -21 | -1.00 | 2 |
| EVLEGEVEKEAL | P05455 | 9 | K | -1 | 20 | -20 | -1.00 | 3 |
| KLGEVSVESENY | P49321-3 | 1 | K | -1 | 20 | -20 | -1.00 | 11 |
| LTRVEVTEFEDIKSGY | Q01105-3 | 13 | K | -1 | 20 | -20 | -1.00 | 3 |
| TFPLAEKVKAF | F8VWS0 | 9 | K | -1 | 18 | -18 | -1.00 | 2 |
| ADKVPKTAENF | C9J5S7 P629: | 3 | K | -1 | 17 | -17 | -1.00 | 8 |
| AADESTGSIAKRL | P04075 | 11 | K | -1 | 16 | -16 | -1.00 | 2 |
| KVFPGSTTEDY | Q13263 | 1 | K | -1 | 16 | -16 | -1.00 | 10 |
| AGKILNDDTALKEY | Q5W0S4 Q5\ | 12 | K | -1 | 16 | -16 | -1.00 | 2 |
| SNFPISEETIKLL | Q9NR30-2 | 11 | K | -1 | 16 | -16 | -1.00 | 2 |
| GVSGYPTLKIF | G5EA52 | 9 | K | -1 | 15 | -15 | -1.00 | 2 |
| AKGHYTEGAELVDSVL | P68371 | 2 | K | -1 | 15 | -15 | -1.00 | 14 |
| DVSGYPTIKIL | P13667 | 9 | K | -1 | 14 | -14 | -1.00 | 2 |
| GGDPIPKSPF | O75369-3 P2 | 7 | K | -1 | 13 | -13 | -1.00 | 3 |
| TNKGTEDFIVESLDASF | C9IZQ1 | 3 | K | -1 | 10 | -10 | -1.00 | 14 |
| SLLATEDKEAL | P09874 | 8 | K | -1 | 10 | -10 | -1.00 | 3 |
| KLGEIVTTIPTIGF | P61204 | 1 | K | -1 | 10 | -10 | -1.00 | 13 |
| DLKNPSDSAVHSPF | Q14157-3 | 3 | K | -1 | 10 | -10 | -1.00 | 11 |
| AEKLGGSAVISLEGKPL | E9PK25 | 15 | K | -1 | 9 | -9 | -1.00 | 2 |
| AIDKSLTPVTL | O00273 | 4 | K | -1 | 8 | -8 | -1.00 | 7 |
| NNKLVTLPVSF | Q96AG4 | 3 | K | -1 | 8 | -8 | -1.00 | 8 |
| DSGFGGGAGVETGGKL | E9PB61 | 15 | K | -1 | 7 | -7 | -1.00 | 1 |
| SSIGEVESAKL | Q15717 | 10 | K | -1 | 7 | -7 | -1.00 | 1 |
| AKALESPERPF | P00558 | 2 | K | -1 | 6 | -6 | -1.00 | 9 |
| EGSPIKVTL | P06748-3 | 6 | K | -1 | 5 | -5 | -1.00 | 3 |
| SDYVGSGPPKGTGL | P30086 | 10 | K | -1 | 5 | -5 | -1.00 | 4 |
| SKVPSLVGSF | P78417 | 2 | K | -1 | 5 | -5 | -1.00 | 8 |
| GASKLVPVGY | P29692-3 | 4 | K | -1 | 4 | -4 | -1.00 | 6 |
| SLDDIIKL | E9PB61 | 7 | K | -1 | 3 | -3 | -1.00 | 1 |
| TNEVLTKTY | P55209 | 7 | K | -1 | 3 | -3 | -1.00 | 2 |
| VKETTYY | P31689 | 2 | K | -1 | 2 | -2 | -1 | 5 |

| | | | | | | | | |
|---|---|---|---|---|---|---|---|---|
| EGSPIKVTLATL | P06748-3 | 6 | K | -1 | 1 | -1 | -1 | 6 |
| QKALDLDSSCKEAADGY | P31948 | 2 | K | -1 | 1 | -1 | -1 | 15 |
