## Supplementary information for "Covalent Protein Painting Reveals Structural Changes in the Proteome in Alzheimer Disease"

to

### CPP of GAPDH in reactant-limited conditions

In order to determine whether specific lysine residues at the surface of the GAPDH homo-tetramer were preferentially dimethylated, either formaldehyde or sodium cyanoborohydride was added at varying concentrations relative to the number of primary amines and the titration evaluated by monitoring the electrophoretic mobility of dimethylated GAPDH (Supplementary Information Figure 1A). The reaction conditions used in the CPP experimental protocol ensured that GAPDH was completely dimethylated. The gel-electrophoretic mobility was different for non-modified and completely dimethylated GAPDH homotetramers. Native gel electrophoresis showed a sharp band for non-modified GAPDH and a faster migrating single band for completely dimethylated GAPDH. When reagents were sub-stoichiometric to the number of primary amines of all GAPDH molecules, lysine residues at the protein surface were sub-stochiometric and differentially modified. Partially dimethylated GAPDH slightly shifted as a single band to higher molecular weight and closer to the non-modified GAPDH homotetramer. The shift in mobility proportionally scaled with the amount of the reagent that limited the chemical reaction. Notably, incompletely labeled GAPDH molecules did not separate into distinct fractions or into an extended smear of completely non-modified and/or completely dimethylated GAPDH homotetramers indicating that the dimethylation pattern of GAPDH under sub-stochiometric reaction conditions is also influenced by the specific chemical reactivity of individual lysine sites in the molecule.

Confirming differential reactivity under sub-stochiometric reaction conditions, mass spectrometry showed that specific lysine residues were labeled with different efficiencies under reagent-limiting conditions (Supplementary Information Figure 1B). GAPDH#K107, K194, and K271 were almost as efficiently dimethylated as in the control despite limiting amounts of reactant. In contrast, lysine residues K27, K55, K117, K139, K162, K215, K263, and K334 had significantly reduced labeling efficiencies (> 2-fold + 1 $\sigma$, denoted with an asterix in supplementary information figure 1B). GAPDH#K309 remained inaccessible and thus served as a positive control, confirming that GAPDH homotetramers remained intact even following dimethylation. Thus, the reactivity of individual, solvent exposed lysine sites differed under reagent-limiting conditions and a subset of lysine residues was more likely to be chemically modified than others.

### Correlating Relative Solvent Accessibility with Lysine Reactivity in CPP

The results obtained with CPP posed the question of whether the extent of relative dimethylation scales with the solvent exposed area of the $\varepsilon$-amine in crystal structure data. To answer this question, we compared the CPP results to the solvent accessible surface area (SASA) of each lysine $\varepsilon$-amine in the crystal structure. Supplementary information Figure 2A displays the SASA of each $\varepsilon$-amine in GAPDH *versus* the relative labeling efficiency obtained with CPP. Lysine K309 showed a very low SASA (< 1 Å$^2$) in crystal structure and was the most inaccessible lysine site for chemical modification in almost all of the highly purified GAPDH molecules. We observed that the $\varepsilon$-amine of a lysine residue could be readily dimethylated as soon as it was exposed by > 20 Å$^2$ to the solvent in GAPDH *in vitro*. Overall, SASA values retrieved from crystallographic data and mass spectrometric measurements (shown here: HEK controls in heat shock experiment) correlated for only a very limited number of lysine sites although replicate measurements showed high reproducibility between biological replicates (Supplementary information figure 3), indicating that the extent of dimethylation may depend on additional parameters in addition to the SASA value of the lysine $\varepsilon$-amine determined by crystal structure or other techniques that deliver high spatial resolution (Supplementary information figure 2B). Thus, additional determinants other than the SASA might govern labeling efficiency. These might include but are not limited to the specific local environment of the $\varepsilon$-amine in case it is > 1 Å$^2$ solvent-exposed as well as dynamic protein conformation changes and protein-protein interactions *in vivo*.

### Limiting Reaction Time in CPP

Because the SASA of $\varepsilon$-amines in crystal structures did not correlate with the relative labeling efficiencies observed with CPP, it raised the question of whether the extent of dimethylation may actually be influenced by the immediate local environment of the primary amine. To address this question, we limited dimethylation reaction times in order to elucidate how reaction kinetics might be influenced at individual lysine sites by additional structural features of the protein.

GAPDH was subjected to time-limited dimethylation. Native gel-electrophoresis revealed that GAPDH homotetramers migrated as a sharp band that successively shifted towards the completely dimethylated GAPDH homotetramer with increasing reaction times (Supplementary information figure 4A). After 5 s of labeling, all GAPDH homo-tetramers moved ~$2/3^{rd}$ distance towards the fully dimethylated homo-tetramer. The final transition to fully dimethylate almost all GAPDH homo-tetramers to completeness required > 2.5 min. This indicated that only a specific subset of lysine sites in GAPDH was labeled within the first 5 s, consistent with a similar time-dependent reaction of formaldehyde in protein crosslinking that has been previously reported [1]. We used CPP to determine which lysine residues required more than 5 s to achieve complete dimethylation (Supplementary information figure 4B) and found that GAPDH#K27, K55, and K162 were dimethylated < 50 % of all GAPDH molecules after 5 s, although each of the $\varepsilon$-amines had a solvent accessible area > 20 Å$^2$.

Because solvent exposure of the primary amines did not explain the observed results and earlier studies indicated that lysine $pK_a$ values critically influence chemical reactivity of lysine residues [2], we analyzed the immediate molecular environment of each lysine residue in greater detail. Focusing on atoms that reside in close proximity to the $\varepsilon$-amine within the protein structure, we looked at the presence of oxygen atoms that are part of the carboxyl moieties of aspartate and glutamate and that might influence the nucleophilicity of the $\varepsilon$-amine in lysine. A nearby carboxyl group might "acidify" a lysine residue and thus reduce the efficiency of Schiff base formation during Michael-addition of formaldehyde. A detailed analysis of crystal structure data revealed that oxygen in the carboxyl groups of aspartate GAPDH#D326 and GAPDH#D166 was in close proximity (~1 Å distance) to the ε-amine in GAPDH#K27 and GAPDH#K162, respectively, and thus may decrease their relative dimethylation efficiencies. In addition, the close proximity (< 2 Å) of the ε-amine of GAPDH#K263 to the carboxyl group of glutamate E265 could explain its reduced labeling efficiency. The only incidence of reduced labeling efficiency under time-limiting reaction conditions which could not be attributed to a nearby carboxyl moiety was GAPDH#K55. Based on crystal structure data, histidine GAPDH#H57 resides in close proximity potentially influencing the chemical reactivity of GAPDH#K263.

In order to validate these observations, we measured the extent of monomethylation per individual lysine residue. During dimethylation, each primary amine is sequentially methylated, and if reaction efficiency is low, monomethylated intermediates may be observed. If monomethylated intermediates are dimethylated after digestion of the protein into peptides, they are converted into dimethylated lysine residues that harbor one light and one heavy isotope labeled methyl moiety, and its peptides show a distinct mass shift of +2 Da instead of +4 Da.

An in-depth analysis of the mass spectrometric data revealed that monomethylation was prevalent under reagent- or time-limited reaction conditions with distinct differences (Supplementary information figure 4). Reagent-limited conditions increased the relative amount of monomethylation, independent of the specific lysine residue in GAPDH (Supplementary information figure 4, yellow bars). In contrast, limiting reaction time increased monomethylation at selected lysine residues only (GAPDH#K27, K162, K215 and K309, Supplementary Information Figure 4, blue bars). As described above, aspartate residues in close proximity to GAPDH#K27 and K162 might slow down dimethylation under reaction time-limited conditions. The differences in mono-methylation observed between concentration- and time-limited condition suggest that the local environment might influence the reactivity of lysine residues if they are amenable to dimethylation based on a minimal solvent exposed area.

In summary, reaction conditions employed in CPP are sufficient to ensure that relative measurements of isotopic dimethyl labels reflect the relative proportion of protein molecules in which a lysine residue is solvent exposed with a surface area of > 1 $Å^2$. Rather than individual reactivity of lysine sites. Based on the results obtained with highly purified GAPDH, CPP directly assesses the quantity of lysine residues that were available for dimethylation due to exposure to the solvent *in vivo*. As described for alternative chemical labeling techniques, reaction time-limited experiments can be used to gain additional information about the local environment of a particular lysine site on a protein's surface [3].

### CPP and PTM status of lysine residues

When interpreting CPP results it is important to understand that a lysine site measured in CPP can also modified by naturally occurring (not chemically introduced) post translational modifications (PTM). In essence, any PTM can also alter its relative abundance protein molecules with lysine being available for chemical modification. With the exception of naturally occurring dimethylation, naturally occurring modifications yield peptides of different mass and biophysical property than chemically dimethylated peptides, and thereby indirectly skew the CPP ratio calculation of accessible *versus* inaccessible lysine residue. Only natural dimethylation of a protein lysine site can directly influence the quantification result because the mass and biophysical properties of naturally dimethylated peptides very closely resemble or directly overlap with the mass of peptides chemically labeled light with isotope defined labeling reagents in CPP. By altering the isotope composition of isotope-defined reagents in the CPP protocol, it is possible to incorporate or discern the extent of natural dimethylation. The following labeling scenarios results in different interpretations when also considering natural dimethylation as a possible alternative:

(1) In case isotope composition of the first labeling (*in vivo*) step in CPP is identical to the naturally occurring isotope composition of dimethylated lysine residues, natural dimethylation contributes to the proportion of protein molecules in which a lysine site was accessible for chemical modification *in vivo*.

(2) In case the natural isotope composition of methyl groups is used in the second labeling reaction, any natural dimethylation of the lysine site directly contributes to the pool of lysine

residues that are determined as inaccessible to chemical modification *in vivo*. In this case CPP does not reveal whether the site was inaccessible due to natural dimethylation (PTM) or steric hinderance during the initial chemical labeling reaction *in vivo*. The possibility that a lysine site of interest was in fact PTM modified by natural dimethylation needs to be confirmed in an additional experiment.

(3) First and second labeling step use non-natural isotope compositions. Here, chemical dimethylation as well as natural dimethylation can be identified and discerned in the same experiment. However, mass differences between different isotope labeled peptides are +2 Da instead of +4 Da making it necessary to correct for a potential overlap between the second isotopic peak of the lighter isotope labeled peptide within the quantification of the heavier isotope labeled peptide [4]. Furthermore, isotope impurities of labeling reagents (typically 2 % of the first neighboring isotope) may result in stronger artificial signal for dimethylation at the additional masses of the isotope-defined peptides.

(4) If a mass spectrometer with high mass resolving power is available, isotope-defined labeling reagents can be used in a labeling scheme which employs pairs of isotopologue dimethyl moieties that are isobaric as demonstrated here and further described in [5].

Here, labeling solvent exposed lysine sites first with heavy isotope combinations and then with light isotopes was preferred because lysine sites are far more likely to be accessible than inaccessible. Labeling accessible lysine sites with heavy isotopes first also reduces interference by isotope impurities when quantifying the typically less abundant proportion of inaccessible lysine sites that are measured as light labeled peptide peaks [4].

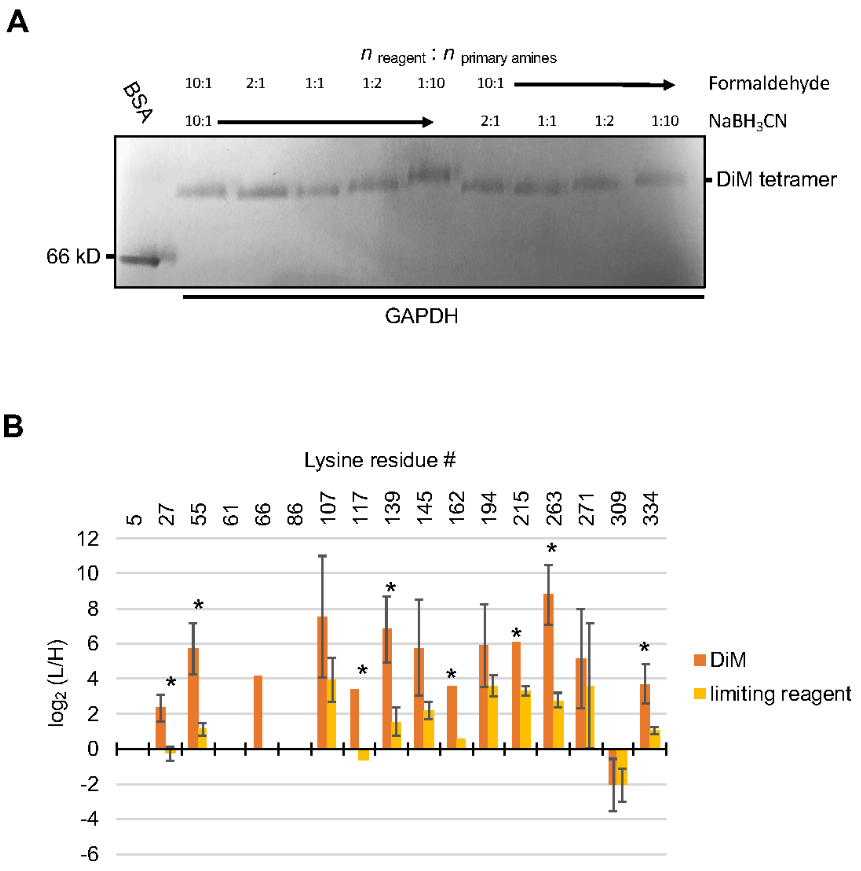


**Supplementary information figure 1: Dimethylating GAPDH with limited amounts of reactant reveals differential chemical reactivity of $\varepsilon$-amines in lysine.** (**A**) GAPDH was dimethylated in different sub-molar ratios of either formaldehyde or sodium cyanoborohydride relative to all lysine sites available in GAPDH in the sample (n : n, reactant/primary amines) and separated by native Blue gel electrophoresis. (**B**) The bar graph compares the relative labeling of each GAPDH lysine residues in reactant-reduced conditions (1 : 1, formaldehyde/primary amines, yellow) with control (10 : 1, reactants/primary amines, orange). Measurements were significantly different ($\Delta > (\log_2 2 + \sigma_1 + \sigma_2)$) for a subset of lysine residues as indicated by an asterix. Error bars are standard deviation ($\sigma$).

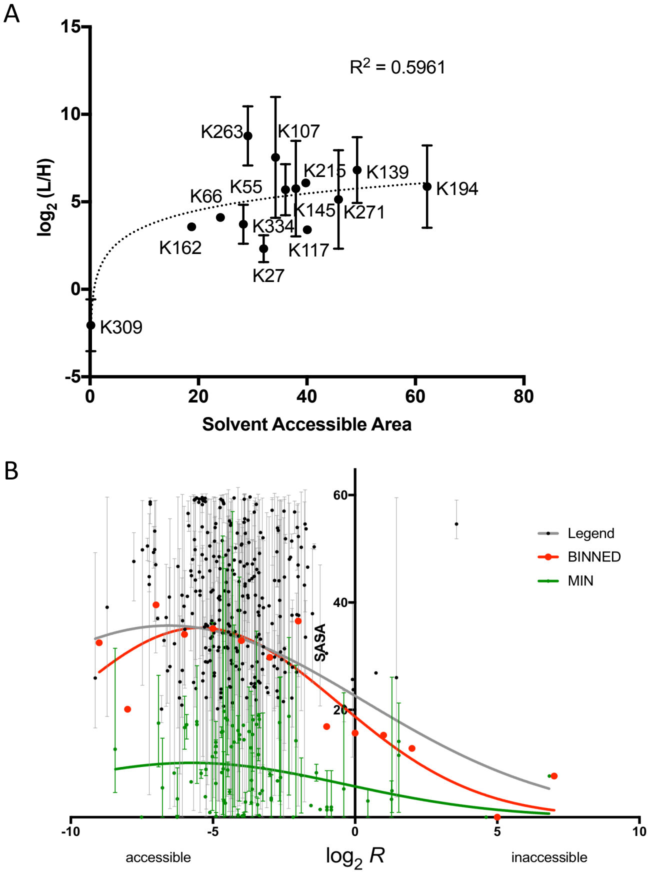


**Supplementary information figure 2: CPP and ε-amine atoms in lysine residues of GAPDH homotetramers and additional protein displayed different solvent exposed areas. (A)** The scatter plot indicates the absolute area (x-axis in Å$^2$) at which each ε-amine of lysine is exposed to solvent (water) in the GAPDH homotetramer (PDB: 4wnc, blue partial spheres in Figure 2C). The y-axis is the relative surface accessibility measured by CPP. The curved fit (dotted line) represents the best fit assuming a logarithmic relationship between solvent area and labeling efficiency. **(B)** The scatter plot shows the ε-amine's accessibility to CPP labeling *versus* its solvent accessible surface area (SASA) for determined by crystal structure in HEK control cell lysate (n = 468). Lines are Gaussian fitted to all data (grey), average per integer bin (red), and all sites with an average SASA value below 20 Å$^2$ (green). The outlier value (bin x = 4, y = 56) was removed for curve fitting.

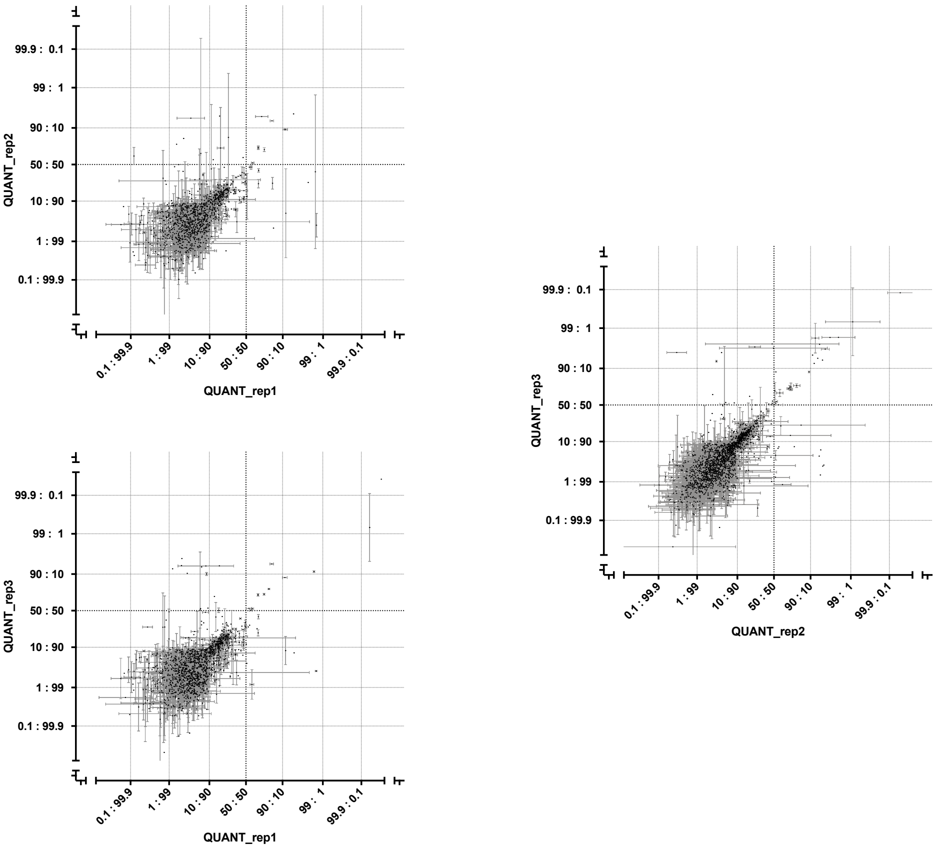


**Supplementary information figure 3: Direct comparison of CPP biological replicate measurements in HEK293 control cells.** The scatter plots indicate the reproducibility between measurements for the same site (individual dots) with error bars in a direct comparison of control HEK 293 cells measured in biological triplicates. Each scale at both axes are the percentage of chemical reactivity per site with a 100 % reactivity at the origin. Replicate 2 and 3 are more similar (average difference $\Delta \log_2 R$ = 0.93, $\sigma$ = 1.14, or ΔR = 1.9-fold) to each other than to replicate 1 (average difference between replicate 1 and 2: $\Delta \log_2 R$ = 1.3, $\sigma$ = 1.24, or ΔR = 2.47-fold and average difference between replicate 1 and 3: $\Delta \log_2 R$ = 1.39, $\sigma$ = 1.27, or ΔR = 1.62-fold).

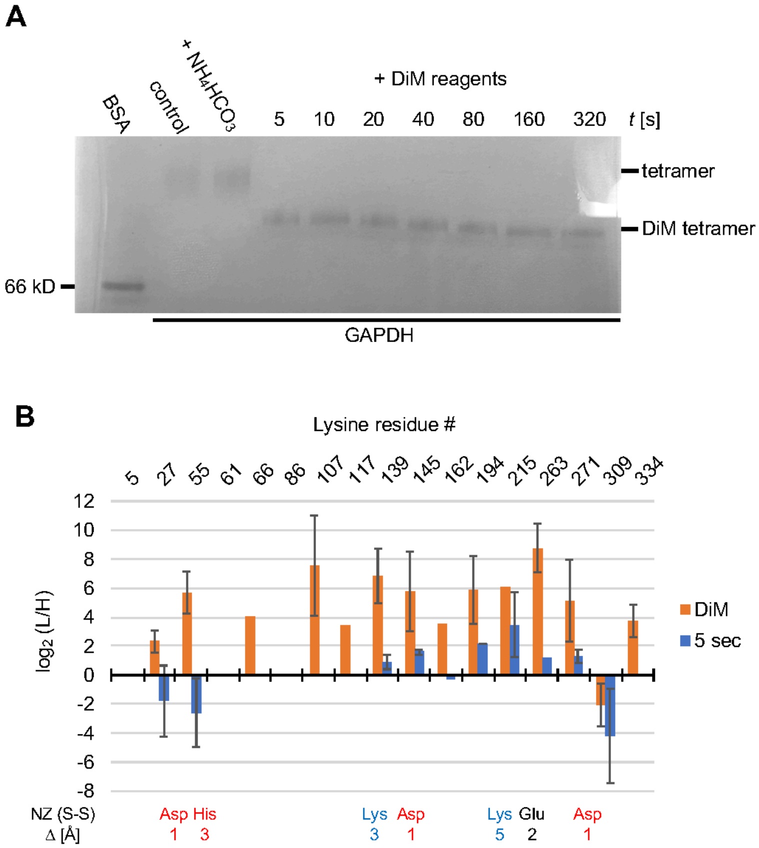


**Supplementary information figure 4: Limiting reaction time revealed differential dimethylation efficiencies of lysine ε-amines in GAPDH.** (**A**) GAPDH homo-tetramers were subjected to different incubation times (in seconds) and separated by native gel electrophoresis. (**B**) The bar graph indicates the relative efficiency in dimethyl labeling of GAPDH after 5 s ("5 s", blue) in comparison to control ("DiM", orange). The amino acid as well as the distance (Δ) of the lysine ε-amine (NZ) to the next closest charged side amino acid side chain (S-S) are shown below the bar graph.

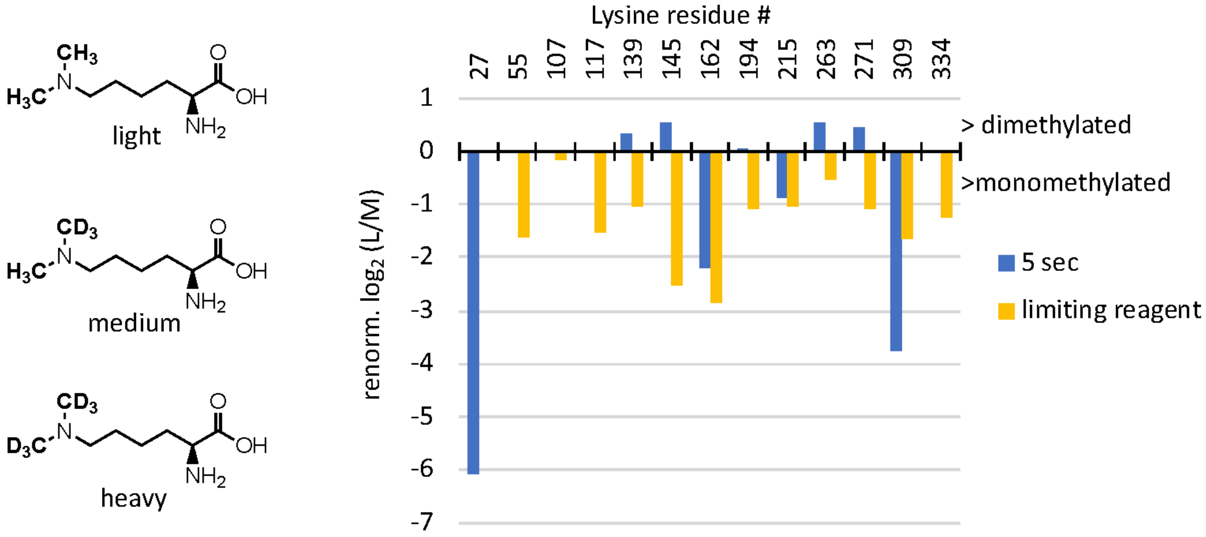


**Supplementary information figure 4: The extent of monomethylation showed reduced labeling efficiency at select lysine residues of GAPDH.** Chemical structures to the left indicate the three distinct methylations patterns of the ε-amine in lysine (K) depending on the completeness of dimethylation during the first labeling step. Lysine residues were labeled light (“L”) or heavy (“H”) only, or intermediate (“M”) wherein one methyl moiety is “light” ($CH_3$) and the second methyl moiety is “heavy” ($CD_3$). The bar graph to the right shows the abundance of monomethylated intermediates relative to dimethylated products (“M/L”) for each lysine residue in GAPDH following normalization to control. Bars depict the “M/L” ratio following either 5 s of labeling (“5 sec”, blue) or upon reduced availability of labeling reagent (“limiting reagent”, yellow, see Figure 4). Positive $\log_2$-transformed ratio values indicate increased presence of dimethylation whereas negative values show increased relative levels of monomethylation. Red boxes highlight lysine residues within 1 Å to carboxyl groups of aspartate.
